# Conserved DNA Methylation Signatures in The Prefrontal Cortex of Newborn and Juvenile Guinea Pigs Following Antenatal Corticosteroid Exposure

**DOI:** 10.1101/2024.03.26.586671

**Authors:** Bona Kim, Alisa Kostaki, Stephen G. Matthews

## Abstract

Antenatal corticosteroids (ACS) are provided to improve perinatal survival when there is risk of preterm birth. Though evidence suggests increased risk of developing neurobehavioural disorders in exposed offspring, the mechanisms that mediate this relationship remain largely unknown. Here, we investigated the DNA methylation patterns in the prefrontal cortex (PFC) of exposed offspring. We hypothesized that differential methylation will be evident at both newborn and juvenile ages.

Pregnant guinea pigs were administered saline or betamethasone (1mg/kg) on gestational days 50/51 to mimic a single course of ACS. gDNA was isolated from the PFC of term-born offspring on postnatal day 1 (PND1) and PND14 to identify differentially methylated CpG sites (DMCs) using reduced representative bisulfite sequencing.

In the PND1 PFC, 1521 DMCs, annotating to 145 genes were identified following ACS. Identified genes were involved in pathways regulating ‘developmental cellular process’. In the PND14 PFC, 776 DMCs representing 46 genes were identified, and were enriched in ‘synaptic signalling’ pathways. Though no individual DMCs were identified at both PND1 and PND14, differential methylation was consistently observed at the binding sites of transcription factors PLAGL1, TFAP2C, ZNF263, and SP1 at both ages.

In this study, we identified an altered DNA methylome in the PFC of ACS-exposed guinea pig offspring at both newborn and juvenile ages. Notably, a unique methylation signature was consistently observed at four key transcription factor binding sites at multiple post-natal time points, indicating a persistent change which may predispose the development of altered neurobehavioural phenotypes that have been described in exposed offspring.

## INTRODUCTION

Antenatal corticosteroids (ACS) are synthetic glucocorticoids provided in the form of betamethasone or dexamethasone to pregnant persons at risk of preterm birth to accelerate fetal lung development and improve perinatal survival [1–3]. The current clinical standard is to provide a single course of treatment between gestational weeks 23^+0^ to 33^+6^ in the form of betamethasone (2×12mg intramuscular injections, 24hours apart) or dexamethasone (4×6mg intramuscular injections, 12 hours apart). While multiple or repeat courses were recommended in the past, providing weekly treatment until birth or term (37^+0^ weeks), studies have indicated no additional benefit in comparison to a single course whilst negative consequences increased [4–7]. Nonetheless, single course treatments are still associated with negative long-term consequences, leading to increased risk for cardiometabolic, immune, and neurodevelopmental disorders [8–11]. Räikkönen *et al.* reported that ACS exposure increases the risk of developing mental or behavioural disorders in term-born children at as early as 5 years [11], while Alexander and colleagues demonstrated heightened stress responses in term-born children ages 6-11 [12], and altered phenotypes of higher-order cognitive and decision-making capacities [13, 14] in term-born adolescents exposed to ACS. Despite evidence of altered behavioural phenotypes following ACS treatment, the molecular mechanisms that underlie these long-lasting changes remain largely unknown.

Epigenetic modification such as DNA methylation patterns are dynamic changes to the chromatin structure that can influence gene expression patterns following an environmental stimulus. The relatively stable nature of DNA methylation makes it a good candidate to study programming mechanisms behind long-lasting or late-onset phenotypic changes in response to acute exposures like ACS. Indeed, previous work from our group has demonstrated that prenatal exposure to repeated courses of synthetic glucocorticoids can leave both acute [15, 16] and long-term [17] changes to the DNA methylome in the hippocampus.

The prefrontal cortex (PFC) is a critical region of the brain that is involved in regulating numerous cognitive functions such as attention, inhibitory control, habit formation and memory (working, spatial or long-term) [18]. In response to ACS treatment, changes in expression of the glucocorticoid receptor have been observed in the neonatal marmoset monkey PFC [19], while multiple courses of ACS were associated with genome-wide transcriptomic changes in the guinea pig PFC at post-natal day (PND) 40 [17]. The PFC is also one of the last brain structures to fully mature [20], demonstrating a longer period of sensitivity and a wider opportunity for intervention. As such, we investigated for the first time, the effects of a single-course of ACS on genome-wide methylation patterns in the newborn PFC and the longitudinal shift in the methylation landscape over the first weeks of life. We hypothesized that a single-course of ACS exposure results in genome-wide changes to the DNA methylome in the newborn guinea pig PFC. We further hypothesize that a subset of these changes will be sustained across early post-natal life, setting exposed offspring on an altered developmental trajectory.

## METHODS

### Animal Cohort Development

Female Dunkin-Hartley guinea pigs (Charles River) were singly housed and mated while food and water were available *ad libitum* on a 12-hour light-dark cycle under approval by the Animal Care Committee at the University of Toronto as previously described [21]. Pregnant dams were subcutaneously injected with a single course of betamethasone (ACS: 1mg/kg) or saline (Veh: equi-volume) on gestational days (GD) 50/51 and left undisturbed outside of routine cage care until term delivery (GD70). Offspring were sacrificed on PND1 (n=7/gp) or PND14 (n=9/gp) by decapitation under isofluorane. The PFC was isolated as we have described previously [17], frozen on dry ice and stored in −80°C until use.

### PFC Sample Preparation (DNA, RNA)

Frozen PFC regions were mounted in a cryostat to isolate the medial PFC region using a 1mm diameter punch needle. Tissue punches were homogenized using the TissueLyser II (Qiagen), and aliquoted into separate tubes for DNA and RNA extractions. ***RNA*** (260/280>1.8) was extracted using TRIZOL and converted to cDNA using the SensiFast cDNA synthesis kit (Agilent). qRT-PCR was performed against target genes and quantified using a CFX96 Real-Time System (Bio-Rad). Target gene expression was calculated using ***2^−ΔΔct^***with *Ywhaz* and *18s* as reference genes (Table S1). ***Genomic DNA*** was extracted using the AllPrep DNA/RNA/miRNA Universal Kit (Qiagen). Samples which met quantity (11.75ng/µL) and quality (DNA Integrity Number > 6) thresholds were used to prepare Reduced Representation Bisulfite Sequencing (RRBS) libraries using the Ovation RRBS Methyl-Seq System 1-16 (NuGen) and the EpiTect Fast DNA Bisulfite kit (Qiagen). Libraries were sequenced using the NextSeq500 platform (Illumina, Donnelly Sequencing Centre, University of Toronto) using single-end reads of 75bps as per manufacturer protocol. Samples were sequenced in pooled multiplexes of 10, balanced for treatment conditions to minimize potential batch effects.

### Bioinformatic analysis

#### Identification of differentially methylated sites and genes & pathway enrichment

Sequenced data were trimmed to remove low-quality reads and accessory sequences from adaptors with Phred quality scores <30 using *Trim Galore (v0.6.4),* which is a wrapper script around CutAdapt [22]. Reads without a 5’ end MspI site signature were removed using a Python script provided by NuGEN (Github: trimRRBSdiversityAdaptCustomers.py). Trimmed reads were aligned to the guinea pig genome (CavPor3.0) using *Bismark (v0.16.0)* [23] and *Bowtie2* (v2.3.4.3) [24]. Differentially methylated CpG sites (DMC) were identified using *MethPipe* [25, 26], to compare between ACS-treated and untreated samples. *methpipe/methcounts* obtained methylation levels at individual cytosines, and beta-binomial regression analyses were performed with *methpipe/radmeth* under the *regression* option. False discovery rate (FDR) correction was performed using the *adjust* option (bins 1:200:1) in *methpipie/radmeth* based on neighbouring sites. Subsequent analyses of DMCs were limited to sites with ≥10 reads, ≥5% methylation difference, and FDR≤0.05. DMCs were annotated using CompEpiTools [27] (R pkg) to call genes based on the Ensembl database. DMCs were localized to genomic features (intragenic – within gene body, promoter, or intergenic). Promoter regions were defined as areas 1000bps up/downstream of a transcription start site (TSS), and intergenic regions were defined as areas further than 1000bps. Computations were performed on the Niagara supercomputer at the SciNet HPC Consortium [28].

#### Gene Set Enrichment Analysis

DMCs were annotated to known genes to understand the functional implications of the altered methylome. Gene set enrichment analysis was performed on g:Profiler [29] to determine differentially methylated gene networks in response to ACS exposure. Searches were performed against the guinea pig genomic database using “All known genes” as the statistical domain scope, which is more extensively documented.

#### Transcription Factor Motif Search

To determine if DMCs were localized within transcription factor binding sites (TFBS), a motif enrichment search was performed using MEME [30]. DMCs were expanded by 100bps (50bps up/downstream) using BEDTools ‘slop’ [31] and converted into a .bed file as input for MEME Suite. Parameters were specified for motif distribution (any number of repetitions) to indicate that any number of non-overlapping motifs may be present per sequence, and for the background model (1^st^ order model of sequences) to adjust for dimer biases (such as high GC content). Output motifs were examined using TOMTOM [32] to identify transcription factors whose binding sequence aligned to the motif. Targeted analysis of known TFBS was performed using position-weight-matrices (PWM) obtained from JASPAR [33] using FIMO [34] on MEME Suite.

## RESULTS

### Differential DNA methylation in newborn PFC following prenatal exposure to betamethasone

In the PND1 guinea pig PFC following ACS exposure, 1521 differentially methylated sites were identified (10x coverage, FDR<0.05, ≥5% methylation difference), of which 797 were hypermethylated and 724 were hypomethylated (Fig. 1a, Table S2). DMCs were localized to promoter (110 sites), intragenic (777 sites), or intergenic (634 sites) regions of the genome (Fig. 1b), which annotated to 145 genes (Fig. 1c), of which 70 were hypomethylated and 75 were hypermethylated.

**Figure 1.**
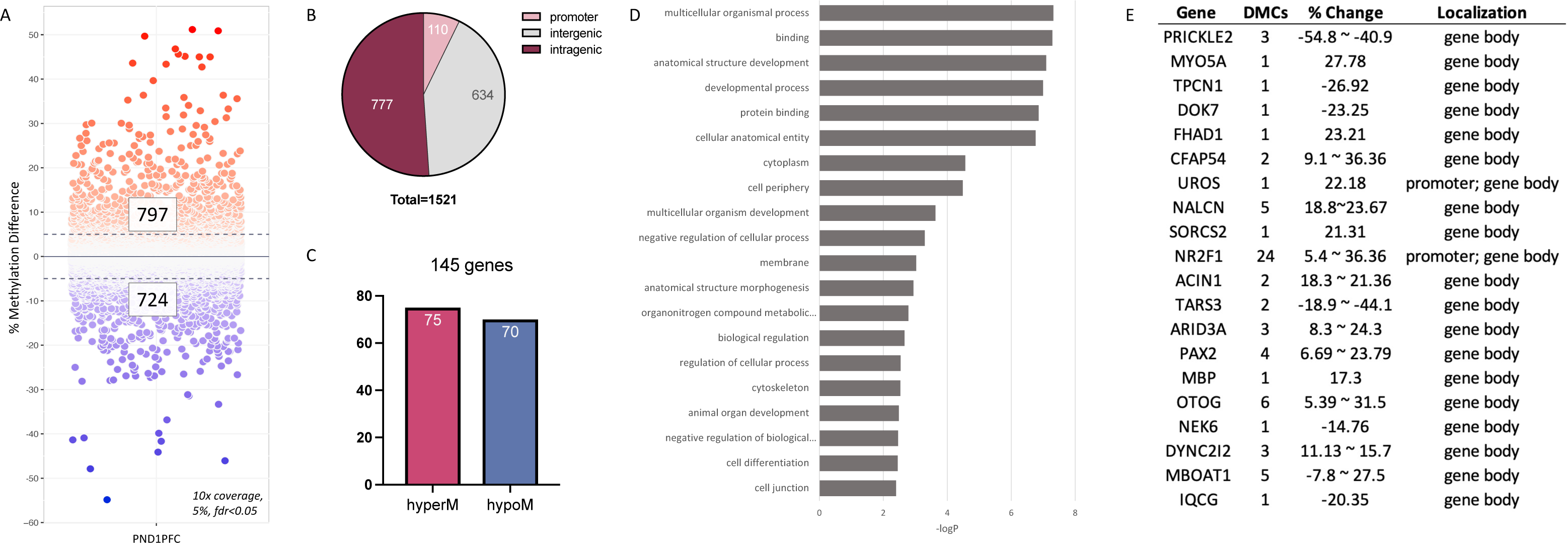
Overview of differentially methylated CpG sites (DMC) in the prefrontal cortex of newborn guinea pigs exposed to ACS. A) Scatterplot of individual CpG sites that were significantly differentially methylated in ACS-exposed animals as compared to unexposed controls. 797 sites were hypermethylated, 724 sites were hypomethylated. >10x reads; ≥5% methylation difference; FDR≤0.05 B) DMCs were localized to various genomic features. C) DMCs were annotated to known genes. 145 genes were identified in total, of which 75 genes were hypermethylated and 70 genes were hypomethylated. D) The differentially methylated genes discovered in the PND1PFC (145 genes) were used to perform a gene set enrichment analysis to identify functional gene networks. Enriched terms are represented from most to least significant. E) List of top 20 differentially methylated genes in the PND1 PFC following ACS exposure.

Gene set enrichment analysis of all 145 genes identified terms that related to ‘protein binding’, ‘developmental process’, and ‘regulation of cellular process’ (Fig. 1d, Table S3). Secondary analyses separately examining the set of hypo- and hypermethylated genes highlighted networks of ‘protein binding’ and ‘developmental process’ among the hypomethylated genes (Fig. S1a), while hypermethylated genes were part of pathways that regulate ‘cytoskeleton’, ‘synapse’, ‘cell adhesion’, and ‘microtubule-based transport’ (Fig. S1b). One gene that was frequently observed among the hypermethylated terms was *Mbp* (myelin basic protein).

*Mbp*, hypermethylated by 17.31% at one DMC in the gene body, was one of the top 20 differentially methylated genes (Fig. 1e). Though a gene typically known for its role in myelination, studies have also examined its roles in regulating neurite outgrowth and microtubule stability during development [35–37]. To determine if DNA methylation changes were associated with changes in mRNA levels, qRT-PCR experiments were performed to indicate significantly decreased levels of *Mbp* (FC: 0.558, p=0.048) in the PFC of ACS-exposed animals (Fig. 2a).

**Figure 2.**
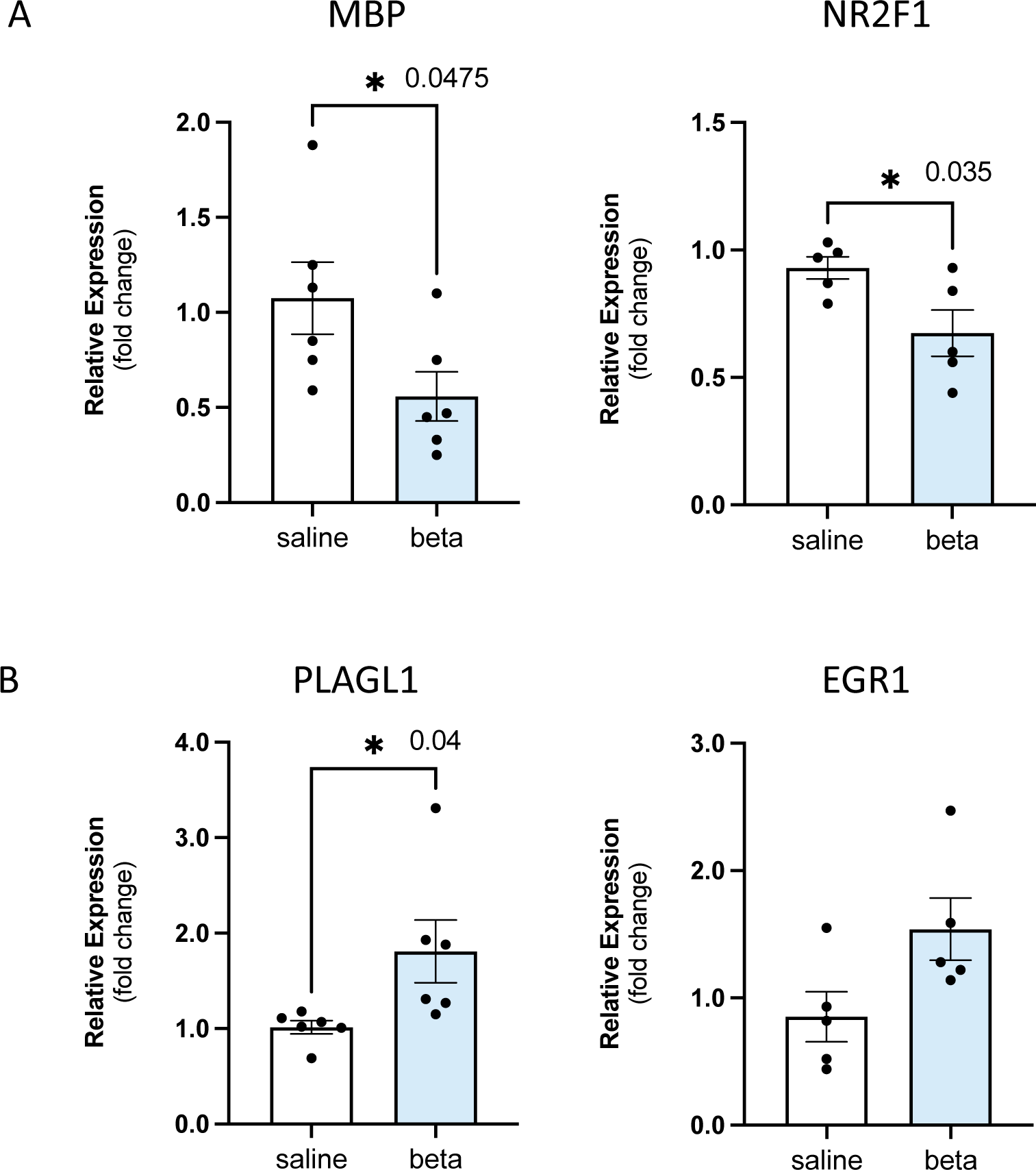
mRNA levels of genes of interest in the PND1PFC following ACS exposure. qRT-PCR were performed for key genes of interest. Values are depicted as mean ± SEM of the relative gene expression. ywhaz and 18s were used as reference genes. *: p<0.05.

Another gene that was among the top 20 differentially methylated genes was *Nr2f1* (nuclear receptor subfamily 2 group F member 1), an important transcription factor in early development that is known for its involvement in cell proliferation, differentiation, and neuronal migration [38]. Unlike other genes in the list of top 20 differentially methylated genes which all had less than 10 DMCs annotated to the gene (Fig. 1e), *Nr2f1* was associated with 24 hypermethylated DMCs that were localized to two distinct clusters; one cluster of 18 DMCs (11-36%) in the promoter region, and another set of 6 DMCs (5-15%) within the gene body (Fig. 3a). Assessing mRNA levels, *Nr2f1* was significantly downregulated in response to ACS (FC: 0.67, p=0.035; Fig. 2a). Observing these relationships between differentially methylated sites and altered mRNA levels, we sought to investigate potential mechanisms which might mediate this association.

**Figure 3.**
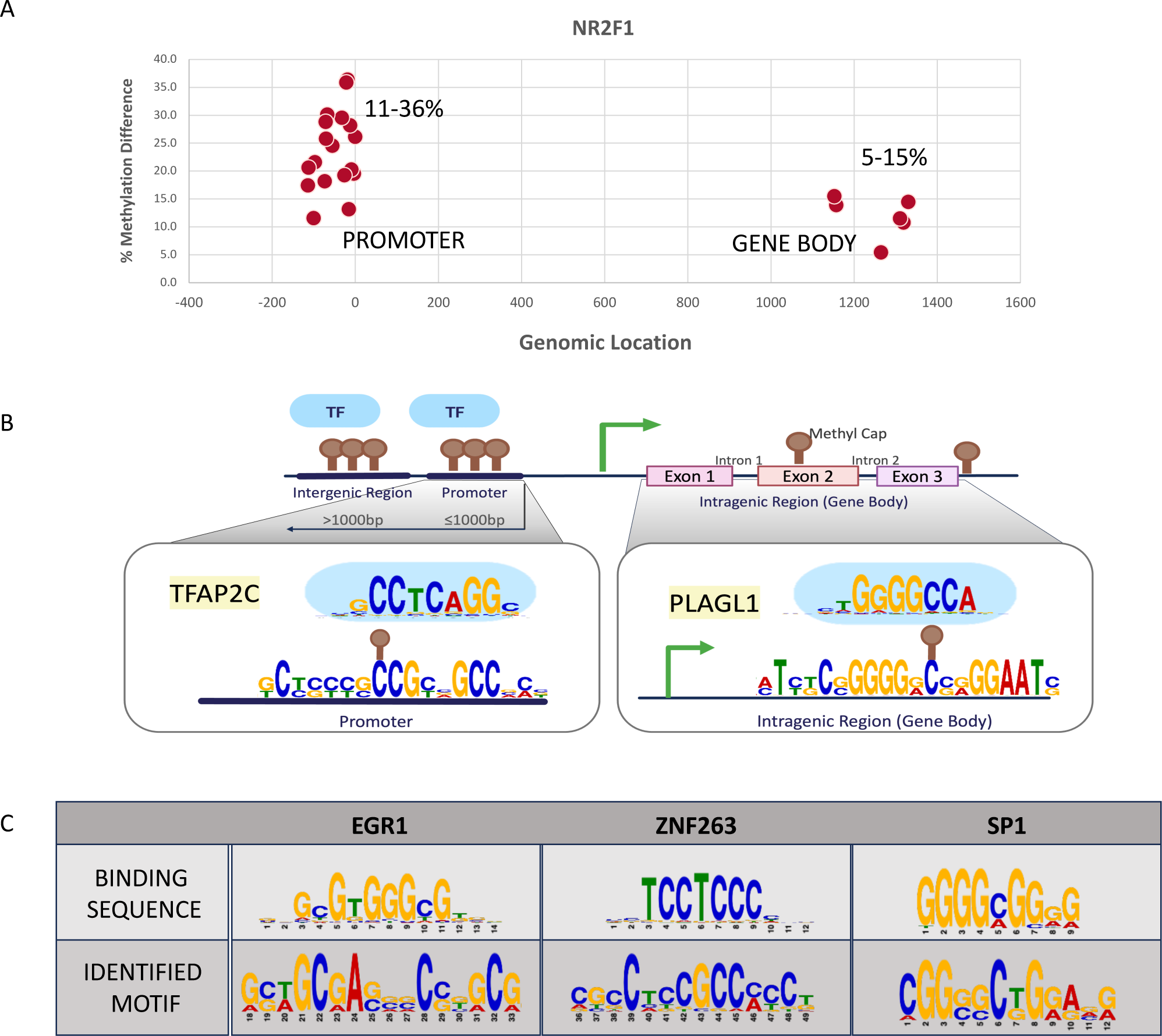
Schematic of NR2F1 methylation and associated transcription factor binding sites following ACS exposure. A) NR2F1 was differentially methylated at two distinct regions, one in the promoter region and another in the gene body. 18 sites (11.52 – 36.36%) were within the promoter while 6 DMCs (5.37-14.45%) were found in the gene body. B) The enriched motif sequence differentially methylated within the promoter region of NR2F1 significantly aligned to the binding sequence for TFAP2C. The differentially methylated cytosine within the sequence is demonstrated with a brown peg. The consensus sequence observed in the gene body significantly matched to the binding sequence for PLAGL1 C) In addition to PLAGL1 and TFAP2C (B), binding motifs for EGR1, ZNF263, and SP1 were identified to be significantly differentially methylated in the PND1 guinea pig PFC following ACS exposure. The sequences indicated on the top represent the binding sequence of the transcription factor (position weight matrix), while the sequence below represents the enriched motif from the dataset. Each nucleotide is represented in a different colour. The size of each nucleotide correlates to the likelihood of finding the specified nucleotide at the position, where the larger the letter the more likely it is to find the nucleotide at the position. Images are adapted from TOMTOM.

### Differential DNA methylation at key transcription factor binding sites in the PND1PFC

DNA methylation can alter gene expression patterns by interfering with DNA binding of transcription factors [39]. To examine if the DMCs identified following ACS were within binding sites of known transcription factors, motif enrichment analysis was performed using MEME Suite v5.5.3 [30]. Motif enrichment seeks to identify fixed-length patterns of sequence that are repeatedly observed among the provided input at a default significance threshold of E-value ≤ 0.05. Investigation of the 24 DMCs annotated to *Nr2f1* identified enriched sequences in both the promoter and gene body (Fig. 3a). Motifs were used as input for TOMTOM [32] to compare against an online database of known binding site motifs. Results identified that the motif found in the *Nr2f1* promoter region was a binding site for the transcription factor TFAP2C while the motif within the gene body was a binding site for PLAGL1 (Fig. 3b). PLAGL1 (Pleiomorphic Adenoma Gene-Like 1) and TFAP2C (Transcription Factor AP-2) binding sites were likewise significantly differentially methylated in the entire dataset of PND1PFC DMCs, in addition to binding sites for transcription factors EGR1 (Early growth response factor 1, NGFI-A), ZNF263 (Zinc Finger Protein 263), and SP1 (Specificity Protein 1). Figure 3b and 3c demonstrate the enriched consensus sequences that were identified, as well as the matched TFBS.

While MEME and TOMTOM function to identify repeating motif sequences, these tools do not specify the genomic location at which they occur. To identify the specific loci at which the TFBS exist in our dataset, we performed a targeted sequence analysis using FIMO [34] on MEME Suite. Briefly, position weight matrixes of the TFBS of interest (identified above, Fig. 3b, 3c) were retrieved from JASPAR 2022 Core [33] to compare against the DNA sequence surrounding the DMCs. PLAGL1 binding sites overlapped at 238 of the 1521 DMCs identified in the PND1 PFC following ACS exposure. These 238 sites annotated to 50 genes (Table S4) that were enriched in pathways of ‘transcription regulation’ and ‘DNA binding’. Binding sites for TFAP2C overlapped at 311 DMCs, annotating to 62 genes (Table S5) enriched in pathways of ‘protein binding’, ‘developmental process’, and ‘inhibitory synapse’. EGR1 binding sites were observed at 264 DMCs, annotating to 50 genes (Table S6) enriched in ‘protein binding’, ‘neuron projection’, ‘cell junction’, and ‘synapse’. The binding sequence for SP1 was found within 189 DMCs annotating to 41 genes (Table S7) enriched in ‘cytokine receptor activity’, ‘ciliary neurotrophic factor-mediated signalling pathway’, and ‘T cell selection’, while the binding sequence for ZNF263 was found within 172 DMCs and annotated to 44 genes (Table S8) enriched in ‘animal organ development’, and ‘microtubule cytoskeleton’. To determine whether these transcription factors (TFs) were also differentially expressed following ACS, qRT-PCR experiments were conducted to determine the mRNA levels of target genes. It was observed that *Plagl1* mRNA was increased (FC: 1.8, p=0.04) among the ACS-exposed group in the PND1PFC as compared to the controls (Fig. 2b), while mRNA levels of *Egr1* demonstrated a strong increasing trend (FC: 1.54, p=0.0598) following ACS exposure (Fig. 2b).

### Differential methylation at transcription factor binding sites was conserved in the juvenile PFC (PND14)

Genome-wide DNA methylation differences observed in the PND1PFC prompted us to investigate whether similar methylation changes are consistently affected in response to ACS at later post-natal timepoints. Examining differential DNA methylation in the PFC of PND14 offspring following ACS exposure, 776 DMCs were identified, of which 349 sites (45%) were hypermethylated and 427 sites were hypomethylated (Fig. 4a, Table S9). Annotating to known genomic features, 34 sites were found within promoter regions, 343 sites were localized to intragenic (gene body) regions, and 399 sites were intergenic (Fig. 4b). Gene annotations identified 46 genes, of which 18 were hypermethylated and 28 were hypomethylated (Fig. 4c). A list of the top 20 differentially methylated genes is represented in Figure 4d. To our surprise, no DMCs previously identified in the PFC at PND1 were also identified at PND14 following ACS. Altered DNA methylation was identified in common at four genes (*Sorcs2, Dok7, Adam11, Timp3)* between the two timepoints, though the associated DMCs were at distinct loci.

**Figure 4.**
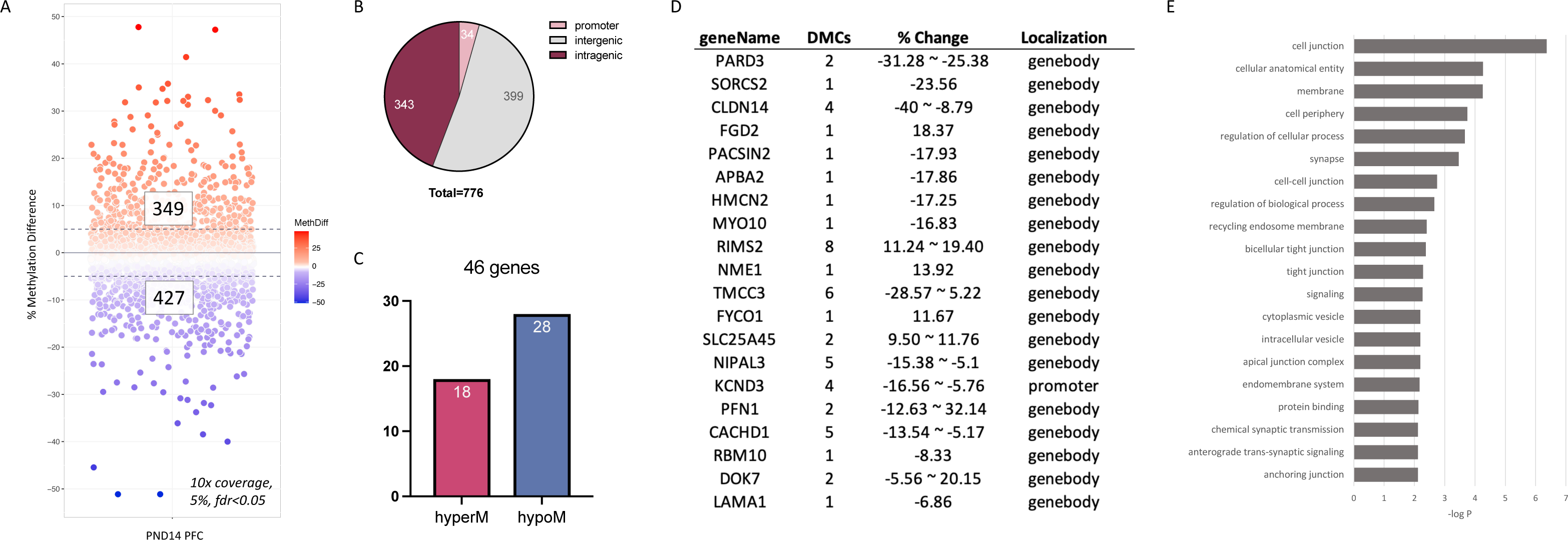
Overview of differentially methylated CpG sites (DMC) in the prefrontal cortex of juvenile guinea pigs exposed to ACS. A) Scatterplot of individual CpG sites that were significantly differentially methylated in ACS-exposed animals as compared to unexposed controls. 349 sites were hypermethylated, 427 sites were hypomethylated. >10x reads; ≥5% methylation difference; FDR≤0.05 B) DMCs were localized to various genomic features. C) DMCs were annotated to known genes. 46 genes were identified in total, of which 18 genes were hypermethylated and 28 genes were hypomethylated D) List of the top 20 differentially methylated genes in the guinea pig PND14PFC following ACS exposure. E) Differentially methylated genes identified in the PND1PFC (145 genes) were used to perform a gene set enrichment analysis to identify functional gene networks. Enriched terms are represented from most to least significant.

Gene set enrichment analysis of the 46 differentially methylated genes identified in the PND14PFC following ACS identified multiple significant terms involved in the regulation of synaptic signalling, including ‘anterograde trans-synaptic signalling’, ‘chemical synaptic transmission’, ‘trans-synaptic signalling’, and ‘synaptic signalling’ (Fig. 4e, Table S10). Four genes (*Sorcs2, Pacsin2, Apba2, Rims2*) were observed among all pathways related to synaptic signalling. qRT-PCR was performed to determine if the observed methylation differences also correlated to gene expression changes.

While mRNA levels of *Sorcs2* (FC: 1.66, p=0.05), and *Apba2* (FC: 1.3, p=0.01) were increased in the ACS-exposed group, *Pacsin2* (FC: 1.3, p=0.2) and *Rims2* levels (FC: 1.16, p=0.14) were not altered (Fig. 5a). To determine if altered gene expression of differentially methylated genes was associated with changes in methylation status at transcription factor binding sites, we performed TFBS analysis using FIMO [34] as described above. It was found that the DMCs annotated to *Sorcs2* and *Apba2*, genes which showed altered expression, were within binding motif for transcription factors PLAGL1 or TFAP2C, while DMCs annotated to *Pacsin2* and *Rims2* were not within any known TFBS. Extending TFBS analysis to the entire dataset, it was found that binding sites for PLAGL1, TFAP2C, ZNF263, and SP1 were enriched (Fig. 6) among the DMCs in the PND14 PFC following ACS, consistent with changes observed at PND1 (Fig. 3b, 3c). PLAGL1 binding sites were found surrounding 129 DMCs, which annotated to 23 genes enriched in vesicles (Table S11), while TFAP2C binding sites were found within 75 DMCs, annotating to 15 genes enriched in ‘nucleoside kinase activity’, and ‘UTP, GTP, CTP biosynthesis and metabolic processes’ (Table S12). Binding sites for SP1 were identified surrounding 81 DMCs that annotated to 16 genes enriched for ‘protein binding’ and ‘membrane’ (Table S13), and ZNF263 binding sites were found among 87 DMCs, which annotated to 13 genes enriched in the ‘membrane’ (Table S14).

**Figure 5.**
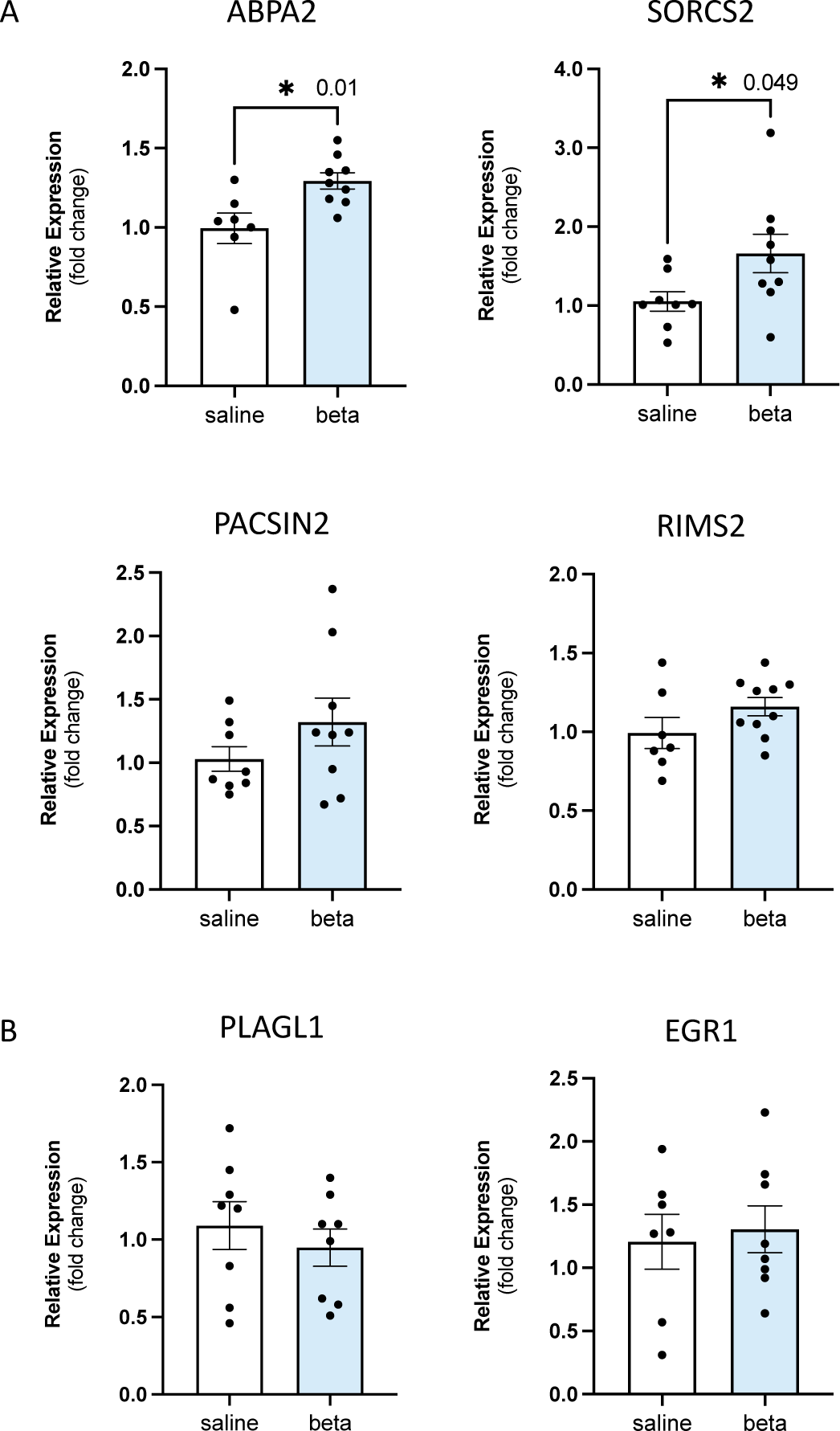
Transcription levels of target genes in the PND14PFC. A) qRT-PCR experiments were performed for genes involved in synaptic signalling. APBA2 and SORCS2 were significantly increased following ACS exposure. B) mRNA levels of transcription factors PLAGL1 and EGR1 were not altered in response to ACS at PND14. *: p<0.05. 18s and ywhaz were used as reference genes.

**Figure 6.**
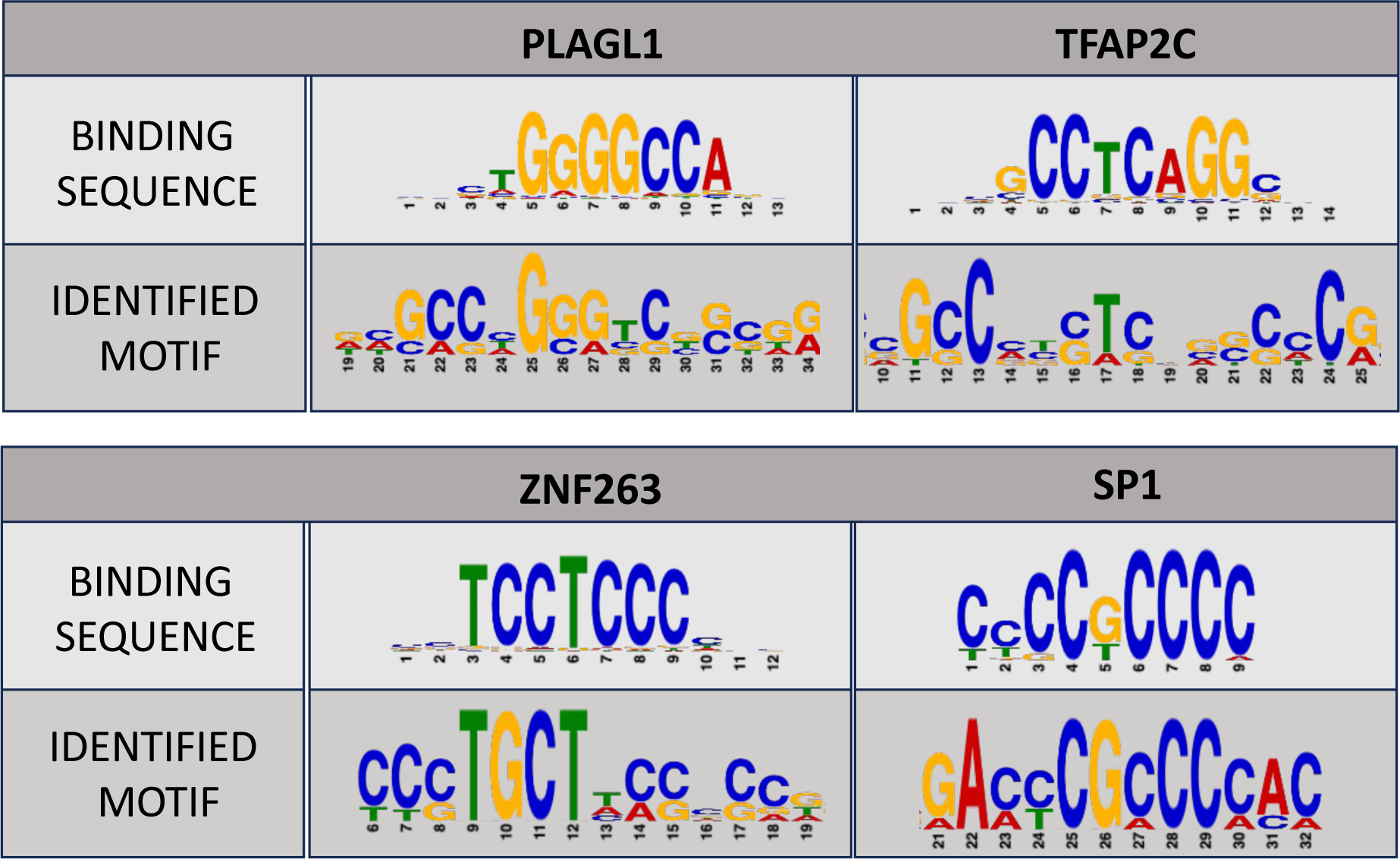
Transcription factor binding site motifs significantly differentially methylated in PND14 guinea pig PFC following ACS exposure. Binding motifs for PLAGL1, TFAP2C, ZNF263, and SP1 were identified to be significantly differentially methylated in the PND14 guinea pig PFC following ACS exposure. The sequences indicated on the top represent the binding sequence of the transcription factor, while the sequence below represents the enriched motif from the dataset.

Unlike in the PND1 PFC, mRNA levels of *Plagl1* were not significantly affected (FC: 0.95, p=0.48l) by ACS treatment at PND14 (Fig. 5b). This suggests that though the effects of ACS on mRNA expression may be transient, the associated methylation changes are long-lasting, highlighting the importance of conserved DNA methylation changes at key transcription factor binding sites following ACS exposure across post-natal development.

## DICSUSSION

In this study, we report that a single course of ACS, a treatment provided to pregnant people at risk of preterm birth to improve perinatal survival, leads to genome-wide alterations to DNA methylation patterns in the PFC of term-born guinea pigs at both PND1 and PND14. We further highlighted that this robust methylation signature is present at the binding sites for four transcription factors (PLAGL1, TFAP2C, ZNF263, and SP1) at both post-natal timepoints, suggesting that ACS exposure can lead to long-term changes in methylation status in the developing brain.

In the PND1 PFC following a single-course of ACS, we observed altered methylation among genes that were enriched in ‘regulation of cellular processes’ (GO: 0009987), which includes events such as ensheathment of neurons. Neuron ensheathment or myelination, has been reported to be dysregulated in association with ACS exposure. Sheep fetuses exposed to ACS demonstrated dysfunctional myelination, evidenced by reduced number of myelinated axons, reduced axon diameter, thinner myelination in the corpus callosum [40] and optic nerve [41], and improper myelin compaction in the auditory nerve [42]. *Mbp* is one gene that regulates myelination that has been shown to be downregulated at 24h, as well as 20 days following ACS treatment [43]. *Mbp* mRNA levels were also decreased following ACS in this study, in the PND1PFC. Similar to previous reports, we also observed a strong trend towards decreased *Mbp* mRNA levels at PND14, indicating prolonged effects of a single course of ACS on brain development. We also reported that the decreased *Mbp* levels were associated with hypermethylation of the gene in the PND1PFC following ACS. Though this site was not within any known TFBS, published ChIP-seq analyses [44] have reported that a glucocorticoid receptor TFBS exists within the *Mbp* promoter region, suggesting a possible mechanism by which *Mbp* expression is regulated following ACS.

PLAGL1 and TFAP2C are zinc finger transcription factors whose binding sites were enriched among the DMCs in both the PND1 and PND14 PFCs following ACS exposure. These factors regulate cell proliferation and differentiation, affecting neurogenesis and neuronal progenitor differentiation [45–47]. The effect of glucocorticoids to promote differentiation and suppress proliferation has been well established in various tissue types, including the brain [48]. In rodents, ACS has been associated with reduced neuronal proliferation [49, 50], while in children, ACS exposure was associated with a thinner cortex as compared to non-exposed children [51]. In the guinea pig at PND1, *Plagl1* mRNA levels were increased in the PFC following ACS exposure, which was associated with genome-wide methylation differences at its binding sequences. We further identified that 44% of the 32 differentially methylated genes involved in cellular differentiation at PND1 contained DMCs within the binding sites for PLAGL1 and TFAP2C, suggesting that these transcription factors may play a key role in mediating the relationship between glucocorticoids and the regulation of cellular proliferation and differentiation. Furthermore, dysregulation of *PLAGL1* in humans is associated with reduced growth rate and intellectual disabilities [52]. As ACS exposure has also been associated with increased risk of intellectual disabilities [11], we suggest that further studies to delineate the role of PLAGL1 in moderating the relationship between ACS and phenotypic outcomes are required.

Despite consistent changes in the methylation status of PLAGL1 binding sites in the PFC at both PND1 and PND14, its mRNA levels were only upregulated at PND1 and no longer significant at PND14. This indicated that altered methylation at TFBSs can persist long-term, even after the associated mRNA changes disappear. This is especially important in the case of EGR1 (NGFI-A), an immediate early response gene (IEG) where the binding site was differentially methylated in the PFC at PND1, even though only a trend towards increased *Egr1* mRNA was observed at the same timepoint following a single course of ACS. We and others have reported increased *Egr1* expression following acute stress or glucocorticoid exposure in various regions of the brain [53, 54], specifically in the cingulate cortex 24-hours after multiple courses of ACS [54]. Since IEGs are rapidly and transiently induced by external stimuli to activate a genomic response [55], it is probable that its expression levels are no longer significantly increased at PND1. Nonetheless, persistent differential methylation among EGR1 binding sites in the PND1 PFC, indicates that its target genes are programmed in a poised state, such that future environmental triggers may elicit a differential genomic response when compared to non-programmed genes. Indeed, DMCs among EGR1 binding sites in the PND1PFC were annotated to genes involved in neuron projection and synapse, supporting previously known roles of EGR1 and glucocorticoids in regulating cell growth and synaptic activation [56–58]. Our finding suggests that EGR1 may play an essential role in mediating the relationship between ACS exposure and synaptic development.

ZNF263 and SP1 binding sites were also differentially methylated following ACS exposure in the PFC at both ages. Previous studies from our group have also identified differential methylation at ZNF263, SP1 and EGR1 binding sites in the PND14 guinea pig hippocampus and peripheral whole blood following multiple courses of ACS [59], suggesting that these may represent a group of transcription factors with binding sites that are sensitive to glucocorticoid exposure, making them attractive biomarkers of ACS exposure. Furthermore, a significant correlation has also been reported between cortisol concentrations measured in the hair and differential methylation at ZNF263, SP1, and EGR1 binding sites in the peripheral blood of 5-year-old children [60]. Differential methylation of ZNF263, SP1, and EGR1 binding sites in multiple studies and multiple tissue types highlights these transcription factors and their binding sequences as important targets of differential methylation following glucocorticoid exposure. Future studies to examine if altered methylation at the TFBS of these three factors as well as PLAGL1 and TFAP2C are associated with altered neurobehavioural phenotypes following ACS exposure will be essential in establishing this methylation signature as a predictive biomarker of developmental adversity following ACS.

We acknowledge that there are key limitations to the study. The DNA was derived from whole tissue punches of the medial PFC, which can include various cell populations including neurons, astrocytes, oligodendrocytes, and endothelial cells. Mixed cell populations can introduce background noise to the DNA methylation data, as epigenetic changes often occur in a cell-type specific manner [61]. Future studies to isolate homogenous cell populations to delineate cell-type specific methylation profiles will be important. It is also a limitation that the TFBS analysis was highly driven by bioinformatic tools, as this approach identifies preferred binding sites of the transcription factors but does not indicate the DNA-protein interaction. Transcription factor binding is dependent on a variety of factors such as environmental triggers and developmental timing [62] and will require ChIP-seq studies to validate the physical interaction. Nonetheless, differential methylation at TFBS indicates an important alteration to the chromatin structure that can persistently interfere with the binding of the transcription factor, programming downstream genes to be in a poised state and elicit an altered genomic response to future environmental triggers. In conclusion, we have identified a unique DNA methylation signature in the guinea pig PFC following a single course ACS at key transcription factor binding sites. This signature was observed in newborn and juvenile offspring, highlighting differential programming of networks involved in neuronal proliferation, differentiation, migration, and synaptic regulation. Future studies to examine methylation patterns in peripheral tissues such as blood following ACS will be important establish whether this signature will be an appropriate surrogate biomarker for changes that occur in the developing brain.

## Acknowledgements

This research was funded by the Canadian Institutes of Health Research (FDN-148368) and by a Canada Research Chair (Tier 1, CRC-2019-00386) awarded to Dr. Stephen G. Matthews. Animal work was performed by Ms. Alisa Kostaki. Bioinformatic computations were performed on the Niagara supercomputer at the SciNet HPC Consortium, funded by the Canadian Foundation for Innovation; the Government of Ontario; Ontario Research Fund – Research Excellence; and the University of Toronto.

## Data Availability Statement

The datasets generated in this study are available from the corresponding author on reasonable request.

