## Supplemental Table 1 for "Conserved DNA Methylation Signatures in The Prefrontal Cortex of Newborn and Juvenile Guinea Pigs Following Antenatal Corticosteroid Exposure"

**Supplementary Table 1. List of Primer Sequences that were used for qRT-PCR experiments**

| Gene | Forward Primer Sequence | Reverse Primer Sequence |
|---|---|---|
| *Mbp* | CAG GAT TTG GCT ATG GAG GCA | GGC TGT CTC TTC CTC CCA GTT |
| *Nr2f1* | TAA TGT TTG GCT ACT CGG TTC A | GCG AGA TGT AGC CGG ACA G |
| *Plagl1* | CCA GAT ACA AAC TGA TGA GGC AC | AGG TGG TCT TTC CGG TTG AA |
| *Egr1* | ACC TAA CCG CAG AGT CCT T | GGT TTG GCT GGG GTA ACT TGT |
| *Apba2* | CAA CTC GGT CGG GGA GAT TC | GTA CAG TGG GGT CTC TTG GC |
| *Pacsin2* | GCG CAT CGA GAA GTC CTA CG | TCC CAT ACT GTG GCC CCT T |
| *Rims2* | ATA CGG TTG GCC GAA TGG AG | GCT GCT TCA CAA TCC CTG TC |
| *Sorcs2* | CTG GGG CTT CCG CTA TCT TT | ACC CCA AAA AGG GAC CAC AG |
| *Gapdh* | TGT ACT GGA GGT CAA TGA AGG | GTC GGA GTG AAC GGA TTT G |
| *Ywhaz* | TGG CCC ATC ATG ACA TTG GG | GCA CAT GGC CAC CAA ATA GG |
| *18s* | CCT GCG GCT TAA TTT GAC TC | CGG ACA TCT AAG GGC ATC AC |
