## Supplemental Table 2 for "Conserved DNA Methylation Signatures in The Prefrontal Cortex of Newborn and Juvenile Guinea Pigs Following Antenatal Corticosteroid Exposure"

**Supplemental Table 2. Full list of differentially methylated cytosines in the guinea pig PND1**

| | seqnames | start | location | MethDiff | Gene name |
|---|---|---|---|---|---|
| 1 | scaffold_0 | 3190833 | intergenic | 14.9099 | |
| 2 | scaffold_0 | 3190835 | intergenic | 14.5480 | |
| 3 | scaffold_0 | 3190849 | intergenic | 15.7848 | |
| 4 | scaffold_0 | 3190902 | intergenic | 21.3345 | |
| 5 | scaffold_0 | 4938109 | intergenic | 19.7743 | |
| 6 | scaffold_0 | 4938141 | intergenic | 19.2302 | |
| 7 | scaffold_0 | 4938157 | intergenic | 21.2032 | |
| 8 | scaffold_0 | 4938197 | intergenic | 6.9170 | |
| 9 | scaffold_0 | 4938208 | intergenic | 8.9113 | |
| 10 | scaffold_0 | 4938239 | intergenic | 5.2588 | |
| 11 | scaffold_0 | 4938243 | intergenic | 9.5022 | |
| 12 | scaffold_0 | 6252132 | promoter | 6.6659 | |
| 13 | scaffold_0 | 56901510 | intergenic | 15.1370 | |
| 14 | scaffold_0 | 62694867 | genebody | -40.9091 | PRICKLE2 |
| 15 | scaffold_0 | 62694881 | genebody | -47.8788 | PRICKLE2 |
| 16 | scaffold_0 | 62694884 | genebody | -54.8485 | PRICKLE2 |
| 17 | scaffold_0 | 81521526 | intergenic | -6.9429 | |
| 18 | scaffold_0 | 81521604 | intergenic | 6.4516 | |
| 19 | scaffold_0 | 81521644 | intergenic | 12.5325 | |
| 20 | scaffold_0 | 81521658 | intergenic | 9.6180 | |
| 21 | scaffold_0 | 81521673 | intergenic | 13.0109 | |
| 22 | scaffold_0 | 81521679 | intergenic | 14.7116 | |
| 23 | scaffold_0 | 81521686 | intergenic | 13.3857 | |
| 24 | scaffold_0 | 81521694 | intergenic | 11.8648 | |
| 25 | scaffold_0 | 81521703 | intergenic | 10.3668 | |
| 26 | scaffold_0 | 81521735 | intergenic | 25.9259 | |
| 27 | scaffold_0 | 81521743 | intergenic | 7.2595 | |
| 28 | scaffold_0 | 81521750 | intergenic | 6.7669 | |
| 29 | scaffold_0 | 81521759 | intergenic | -8.3485 | |
| 30 | scaffold_0 | 81521768 | intergenic | -8.5299 | |
| 31 | scaffold_0 | 81521783 | intergenic | 23.0490 | |
| 32 | scaffold_0 | 81521785 | intergenic | 5.2632 | |
| 33 | scaffold_0 | 81521788 | intergenic | 5.2632 | |
| 34 | scaffold_0 | 81521791 | intergenic | 21.0526 | |
| 35 | scaffold_0 | 81521795 | intergenic | 5.2632 | |
| 36 | scaffold_0 | 81521799 | intergenic | -15.4265 | |
| 37 | scaffold_1 | 12188172 | genebody | 14.4483 | NR2F1 |
| 38 | scaffold_1 | 12188184 | genebody | 10.7448 | NR2F1 |
| 39 | scaffold_1 | 12188192 | genebody | 11.4828 | NR2F1 |
| 40 | scaffold_1 | 12188238 | genebody | 5.3701 | NR2F1 |
| 41 | scaffold_1 | 12188346 | genebody | 13.8646 | NR2F1 |
| 42 | scaffold_1 | 12188350 | genebody | 15.4777 | NR2F1 |
| 43 | scaffold_1 | **12189503** | promoter | 26.1029 | NR2F1 |

| | | | | | |
|---|---|---|---|---|---|
| 44 | scaffold_1 | 12189506 | promoter | 19.4989 | NR2F1 |
| 45 | scaffold_1 | 12189512 | promoter | 20.2381 | NR2F1 |
| 46 | scaffold_1 | 12189515 | promoter | 28.1746 | NR2F1 |
| 47 | scaffold_1 | 12189518 | promoter | 13.1250 | NR2F1 |
| 48 | scaffold_1 | 12189521 | promoter | 36.3636 | NR2F1 |
| 49 | scaffold_1 | 12189524 | promoter | 35.8586 | NR2F1 |
| 50 | scaffold_1 | 12189529 | promoter | 19.1919 | NR2F1 |
| 51 | scaffold_1 | **12189535** | promoter | 29.5455 | NR2F1 |
| 52 | scaffold_1 | 12189558 | promoter | 24.4881 | NR2F1 |
| 53 | scaffold_1 | **12189570** | promoter | 30.0676 | NR2F1 |
| 54 | scaffold_1 | 12189573 | promoter | 25.7353 | NR2F1 |
| 55 | scaffold_1 | 12189574 | promoter | 28.8007 | NR2F1 |
| 56 | scaffold_1 | **12189576** | promoter | 18.1373 | NR2F1 |
| 57 | scaffold_1 | 12189600 | promoter | 21.5686 | NR2F1 |
| 58 | scaffold_1 | 12189603 | promoter | 11.5196 | NR2F1 |
| 59 | scaffold_1 | 12189615 | promoter | 20.5882 | NR2F1 |
| 60 | scaffold_1 | 12189617 | promoter | 17.4020 | NR2F1 |
| 61 | scaffold_1 | 22073022 | intergenic | -9.9795 | |
| 62 | scaffold_1 | 22073034 | intergenic | -18.1476 | |
| 63 | scaffold_1 | 22073043 | intergenic | -13.6364 | |
| 64 | scaffold_1 | 22073062 | intergenic | -12.8938 | |
| 65 | scaffold_1 | 22073083 | intergenic | -15.3031 | |
| 66 | scaffold_1 | 22073091 | intergenic | -13.9891 | |
| 67 | scaffold_1 | 22073209 | intergenic | 43.3566 | |
| 68 | scaffold_1 | 22073210 | intergenic | -7.0011 | |
| 69 | scaffold_1 | 22333270 | intergenic | 30.4233 | |
| 70 | scaffold_1 | 22333304 | intergenic | 46.7980 | |
| 71 | scaffold_1 | 22333310 | intergenic | 43.5961 | |
| 72 | scaffold_1 | 22333317 | intergenic | -8.9606 | |
| 73 | scaffold_1 | 22333318 | intergenic | 39.6552 | |
| 74 | scaffold_1 | 22333345 | intergenic | 13.1720 | |
| 75 | scaffold_1 | 22333360 | intergenic | 10.0493 | |
| 76 | scaffold_1 | 42320161 | promoter | 22.1803 | UROS |
| 77 | scaffold_1 | 58427277 | intergenic | 10.2721 | |
| 78 | scaffold_1 | 58427417 | intergenic | -7.5682 | |
| 79 | scaffold_1 | 67622577 | genebody | 11.1722 | MCPH1 |
| 80 | scaffold_1 | 67622582 | genebody | 10.7028 | MCPH1 |
| 81 | scaffold_1 | 67622586 | genebody | 18.7042 | MCPH1 |
| 82 | scaffold_1 | 67622588 | genebody | 12.7957 | MCPH1 |
| 83 | scaffold_1 | 67622611 | genebody | 14.9232 | MCPH1 |
| 84 | scaffold_1 | 67622618 | genebody | 20.3838 | MCPH1 |
| 85 | scaffold_1 | 67622633 | genebody | 14.8939 | MCPH1 |
| 86 | scaffold_1 | 67622634 | genebody | 12.8364 | MCPH1 |
| 87 | scaffold_1 | 67622640 | genebody | 8.4939 | MCPH1 |
| 88 | scaffold_1 | 67622664 | genebody | 11.2228 | MCPH1 |

| | | | | | |
|---|---|---|---|---|---|
| 89 | scaffold_1 | 67622686 | genebody | 16.8363 | MCPH1 |
| 90 | scaffold_1 | 67622719 | genebody | 12.5307 | MCPH1 |
| 91 | scaffold_1 | 73507716 | intergenic | 11.4618 | |
| 92 | scaffold_1 | 73507741 | intergenic | 26.6611 | |
| 93 | scaffold_1 | 73507755 | intergenic | 20.7641 | |
| 94 | scaffold_1 | 73507764 | intergenic | 27.3810 | |
| 95 | scaffold_1 | 73507778 | intergenic | 33.4718 | |
| 96 | scaffold_1 | 73765934 | promoter | 9.7214 | |
| 97 | scaffold_1 | 73765935 | promoter | 12.1448 | |
| 98 | scaffold_1 | 73765940 | promoter | 9.8005 | |
| 99 | scaffold_1 | 73765941 | promoter | 9.2697 | |
| 100 | scaffold_1 | 73765942 | promoter | 5.8884 | |
| 101 | scaffold_1 | 73765943 | promoter | 15.5836 | |
| 102 | scaffold_10 | 15327268 | genebody | 5.5074 | IL6R |
| 103 | scaffold_10 | 15327363 | genebody | 13.0848 | IL6R |
| 104 | scaffold_10 | 15327371 | genebody | 14.2178 | IL6R |
| 105 | scaffold_10 | 15327383 | genebody | 11.3562 | IL6R |
| 106 | scaffold_10 | 15327400 | genebody | 13.4804 | IL6R |
| 107 | scaffold_10 | 15327412 | genebody | 11.2555 | IL6R |
| 108 | scaffold_10 | 15327420 | genebody | 12.7178 | IL6R |
| 109 | scaffold_10 | 15327447 | genebody | 11.2500 | IL6R |
| 110 | scaffold_10 | 15327463 | genebody | 25.0000 | IL6R |
| 111 | scaffold_10 | 32971346 | intergenic | -18.9423 | |
| 112 | scaffold_10 | 32971348 | intergenic | -14.0687 | |
| 113 | scaffold_10 | 32971370 | intergenic | -13.2252 | |
| 114 | scaffold_10 | 32971389 | intergenic | -15.1564 | |
| 115 | scaffold_10 | 35166305 | intergenic | 45.6140 | |
| 116 | scaffold_10 | 35166327 | intergenic | 50.8772 | |
| 117 | scaffold_10 | 35166332 | intergenic | 51.1696 | |
| 118 | scaffold_10 | 35166362 | intergenic | 49.6732 | |
| 119 | scaffold_10 | 45822346 | intergenic | -18.8235 | |
| 120 | scaffold_10 | 45822371 | intergenic | -19.3590 | |
| 121 | scaffold_10 | 45822383 | intergenic | -18.4615 | |
| 122 | scaffold_10 | 45822409 | intergenic | -17.6923 | |
| 123 | scaffold_10 | 47593463 | genebody | -18.8811 | TARS3 |
| 124 | scaffold_10 | 47593483 | genebody | -44.1176 | TARS3 |
| 125 | scaffold_100 | 3834844 | genebody;ge | 11.7035 | TNIK |
| 126 | scaffold_100 | 3834848 | genebody;ge | 11.0834 | TNIK |
| 127 | scaffold_100 | 3834863 | genebody;ge | 12.1525 | TNIK |
| 128 | scaffold_100 | 3834889 | genebody;ge | 16.4347 | TNIK |
| 129 | scaffold_102 | 1531127 | intergenic | 8.1801 | |
| 130 | scaffold_102 | 2182012 | genebody | 45.0000 | |
| 131 | scaffold_102 | 2182014 | genebody | 42.7273 | |
| 132 | scaffold_102 | 2182021 | genebody | -13.3333 | |
| 133 | scaffold_102 | 2182022 | genebody | 45.0000 | |

| | | | | | |
|---|---|---|---|---|---|
| 134 | scaffold_102 | 2182027 | genebody | -15.4158 | |
| 135 | scaffold_102 | 2182029 | genebody | -15.4158 | |
| 136 | scaffold_102 | 2182033 | genebody | -15.4158 | |
| 137 | scaffold_102 | 2182038 | genebody | -11.9675 | |
| 138 | scaffold_102 | 2182044 | genebody | -21.2982 | |
| 139 | scaffold_102 | 2182048 | genebody | -11.9675 | |
| 140 | scaffold_102 | 2182054 | genebody | -15.4158 | |
| 141 | scaffold_102 | 2182066 | genebody | -15.4158 | |
| 142 | scaffold_102 | 2182077 | genebody | -11.6071 | |
| 143 | scaffold_102 | 2182143 | genebody | -7.6993 | |
| 144 | scaffold_102 | 2182163 | genebody | -5.4740 | |
| 145 | scaffold_102 | 2182170 | genebody | -9.5831 | |
| 146 | scaffold_102 | 2182179 | genebody | -7.9958 | |
| 147 | scaffold_102 | 2267910 | genebody | -5.4965 | GRB10 |
| 148 | scaffold_104 | 2565407 | intergenic | -17.9426 | |
| 149 | scaffold_104 | 2565409 | intergenic | -11.2440 | |
| 150 | scaffold_104 | 2565430 | intergenic | 7.6555 | |
| 151 | scaffold_104 | 2565706 | intergenic | 17.4688 | |
| 152 | scaffold_104 | 2571505 | intergenic | 15.0000 | |
| 153 | scaffold_104 | 2571528 | intergenic | 13.6364 | |
| 154 | scaffold_104 | 2571559 | intergenic | 7.0707 | |
| 155 | scaffold_104 | 2571575 | intergenic | 7.7947 | |
| 156 | scaffold_104 | 2571576 | intergenic | 20.7071 | |
| 157 | scaffold_104 | 2571653 | intergenic | 6.9319 | |
| 158 | scaffold_104 | 2571678 | intergenic | 18.7500 | |
| 159 | scaffold_104 | 2571706 | intergenic | -13.3333 | |
| 160 | scaffold_104 | 2571714 | intergenic | 29.6970 | |
| 161 | scaffold_104 | 2960365 | intergenic | 10.4615 | |
| 162 | scaffold_105 | 4130222 | genebody | 17.3077 | MBP |
| 163 | scaffold_108 | 1186158 | intergenic | -26.9231 | |
| 164 | scaffold_108 | 1187604 | intergenic | -12.4286 | |
| 165 | scaffold_108 | 1187628 | intergenic | 10.2222 | |
| 166 | scaffold_108 | 1187893 | intergenic | -7.7778 | |
| 167 | scaffold_108 | 1187941 | intergenic | -9.0972 | |
| 168 | scaffold_108 | 1187950 | intergenic | 12.6100 | |
| 169 | scaffold_108 | 1188176 | intergenic | 5.3292 | |
| 170 | scaffold_11 | 4675888 | genebody | -9.7203 | RBM20 |
| 171 | scaffold_11 | 4675922 | genebody | -7.1429 | RBM20 |
| 172 | scaffold_11 | 4676011 | genebody | -20.0000 | RBM20 |
| 173 | scaffold_11 | 13227517 | intergenic | 22.0779 | |
| 174 | scaffold_11 | 13228017 | intergenic | -8.1974 | |
| 175 | scaffold_11 | 13231228 | intergenic | 6.6901 | |
| 176 | scaffold_11 | 13617836 | promoter | 23.7925 | PAX2 |
| 177 | scaffold_11 | 13618114 | promoter | 17.0068 | PAX2 |
| 178 | scaffold_11 | 13618122 | promoter | 7.3864 | PAX2 |

| | | | | | |
|---|---|---|---|---|---|
| 179 | scaffold_11 | 13618152 | promoter | 23.1429 | PAX2 |
| 180 | scaffold_11 | 13770615 | intergenic | -6.4958 | |
| 181 | scaffold_11 | 16434380 | intergenic | -10.4762 | |
| 182 | scaffold_11 | 16436151 | intergenic | -7.8007 | |
| 183 | scaffold_11 | 16436183 | intergenic | -16.7033 | |
| 184 | scaffold_110( | 6337 | intergenic | -22.4803 | |
| 185 | scaffold_110( | 6538 | intergenic | 8.8071 | |
| 186 | scaffold_110( | 6564 | intergenic | 5.9795 | |
| 187 | scaffold_110( | 6570 | intergenic | 7.6318 | |
| 188 | scaffold_111 | 12656 | intergenic | -6.6667 | |
| 189 | scaffold_111 | 12671 | intergenic | -14.7368 | |
| 190 | scaffold_111 | 12679 | intergenic | -5.0280 | |
| 191 | scaffold_111 | 12680 | intergenic | -20.0000 | |
| 192 | scaffold_111 | 12729 | intergenic | -7.6879 | |
| 193 | scaffold_111 | 13704 | intergenic | -14.0196 | |
| 194 | scaffold_111 | 1781402 | intergenic | -5.6075 | |
| 195 | scaffold_111 | 1782351 | intergenic | 11.4695 | |
| 196 | scaffold_111 | 3648074 | intergenic | -6.5027 | |
| 197 | scaffold_111 | 3648112 | intergenic | -7.6203 | |
| 198 | scaffold_111: | 15496 | intergenic | -18.5714 | |
| 199 | scaffold_113 | 1676372 | genebody | 8.3627 | CA5B |
| 200 | scaffold_113 | 3940974 | promoter | -6.0212 | OFD1 |
| 201 | scaffold_113 | 3940975 | promoter | -8.0217 | OFD1 |
| 202 | scaffold_113 | 3940985 | promoter | 19.5641 | OFD1 |
| 203 | scaffold_114 | 701054 | intergenic | -13.6114 | |
| 204 | scaffold_114 | 701145 | intergenic | -6.7724 | |
| 205 | scaffold_115 | 1404704 | genebody | -8.7097 | SETD1B |
| 206 | scaffold_115 | 1404768 | genebody | -7.5391 | SETD1B |
| 207 | scaffold_115 | 1404801 | genebody | 6.4494 | SETD1B |
| 208 | scaffold_115 | 1404807 | genebody | -5.4488 | SETD1B |
| 209 | scaffold_115 | 1405377 | genebody | 8.1081 | SETD1B |
| 210 | scaffold_115 | 1405400 | genebody | 5.7143 | SETD1B |
| 211 | scaffold_115 | 1405404 | genebody | 5.7143 | SETD1B |
| 212 | scaffold_115 | 1405669 | genebody | 12.5000 | SETD1B |
| 213 | scaffold_115 | 1405682 | genebody | -11.1742 | SETD1B |
| 214 | scaffold_115 | 1405689 | genebody | -8.5227 | SETD1B |
| 215 | scaffold_115 | 1405696 | genebody | -11.7647 | SETD1B |
| 216 | scaffold_115 | 1405697 | genebody | 10.7955 | SETD1B |
| 217 | scaffold_115 | 1405949 | genebody | 14.7059 | SETD1B |
| 218 | scaffold_115 | 1405987 | genebody | 5.8986 | SETD1B |
| 219 | scaffold_115 | 1405995 | genebody | -7.5803 | SETD1B |
| 220 | scaffold_115 | 1406044 | genebody | 15.8635 | SETD1B |
| 221 | scaffold_115 | 1406173 | genebody | -6.0574 | SETD1B |
| 222 | scaffold_115 | 1407188 | genebody | -21.4286 | SETD1B |
| 223 | scaffold_115 | 1407203 | genebody | -19.6581 | SETD1B |

| | | | | | |
|---|---|---|---|---|---|
| 224 | scaffold_115 | 1407207 | genebody | 6.8376 | SETD1B |
| 225 | scaffold_115 | 1407238 | genebody | 11.1111 | SETD1B |
| 226 | scaffold_115 | 1407460 | genebody | 21.4286 | SETD1B |
| 227 | scaffold_115 | 1407472 | genebody | 7.1429 | SETD1B |
| 228 | scaffold_115 | 1407475 | genebody | 5.8824 | SETD1B |
| 229 | scaffold_115 | 1407488 | genebody | 8.3333 | SETD1B |
| 230 | scaffold_115 | 1407494 | genebody | 15.9091 | SETD1B |
| 231 | scaffold_115 | 1407569 | genebody | -6.5217 | SETD1B |
| 232 | scaffold_115 | 1407593 | genebody | 5.3931 | SETD1B |
| 233 | scaffold_115 | 1407599 | genebody | 7.6628 | SETD1B |
| 234 | scaffold_115 | 1407612 | genebody | -5.5556 | SETD1B |
| 235 | scaffold_115 | 1407636 | genebody | 7.2601 | SETD1B |
| 236 | scaffold_115 | 1407650 | genebody | 12.7193 | SETD1B |
| 237 | scaffold_115 | 1407725 | genebody | 5.6907 | SETD1B |
| 238 | scaffold_115 | 1407768 | genebody | 5.6420 | SETD1B |
| 239 | scaffold_115 | 1407777 | genebody | -28.1429 | SETD1B |
| 240 | scaffold_115 | 1407826 | genebody | -6.2222 | SETD1B |
| 241 | scaffold_115 | 1455563 | genebody;ge | -12.5999 | TMEM120B |
| 242 | scaffold_115 | 1455574 | genebody;ge | -15.7975 | TMEM120B |
| 243 | scaffold_115 | 1455608 | genebody;ge | -6.4093 | TMEM120B |
| 244 | scaffold_115 | 1548838 | intergenic | 7.7723 | |
| 245 | scaffold_115 | 1548840 | intergenic | -8.4923 | |
| 246 | scaffold_115 | 4048902 | intergenic | -5.4612 | |
| 247 | scaffold_117 | 3775742 | genebody;ge | -7.1982 | UNC5D |
| 248 | scaffold_117 | 3776019 | genebody;ge | -7.8431 | UNC5D |
| 249 | scaffold_117 | 3776067 | genebody;ge | 45.1274 | UNC5D |
| 250 | scaffold_117 | 3776076 | genebody;ge | 6.5967 | UNC5D |
| 251 | scaffold_117 | 3777546 | genebody;ge | -12.3077 | UNC5D |
| 252 | scaffold_119 | 328159 | intergenic | 5.7595 | |
| 253 | scaffold_119 | 1850505 | intergenic | -13.1313 | |
| 254 | scaffold_119 | 1850506 | intergenic | -14.0000 | |
| 255 | scaffold_119 | 1850508 | intergenic | 13.3333 | |
| 256 | scaffold_119 | 1850509 | intergenic | 21.4286 | |
| 257 | scaffold_119 | 1850546 | intergenic | 25.0000 | |
| 258 | scaffold_119 | 1850983 | intergenic | 13.3333 | |
| 259 | scaffold_119 | 1850992 | intergenic | 18.7500 | |
| 260 | scaffold_119 | 1851034 | intergenic | 6.9106 | |
| 261 | scaffold_119 | 1851164 | intergenic | -14.5977 | |
| 262 | scaffold_121 | 19291 | intergenic | 5.8106 | |
| 263 | scaffold_125 | 3044922 | promoter | -7.3748 | C2orf50 |
| 264 | scaffold_125 | 3044950 | promoter | -5.3555 | C2orf50 |
| 265 | scaffold_125 | 3386709 | intergenic | -6.5103 | |
| 266 | scaffold_129 | 1845009 | intergenic | -9.1667 | |
| 267 | scaffold_129 | 3116299 | intergenic | 15.0742 | |
| 268 | scaffold_129 | 3116332 | intergenic | 9.9735 | |

| | | | | | |
|---|---|---|---|---|---|
| 269 | scaffold_13 | 3910516 | promoter | 10.2504 | TGFB2 |
| 270 | scaffold_13 | 4860555 | intergenic | -7.0807 | |
| 271 | scaffold_13 | 4860556 | intergenic | 6.5452 | |
| 272 | scaffold_13 | 4860561 | intergenic | -14.8338 | |
| 273 | scaffold_13 | 4860579 | intergenic | -11.8926 | |
| 274 | scaffold_13 | 4860589 | intergenic | -18.5093 | |
| 275 | scaffold_13 | 4860599 | intergenic | -15.5280 | |
| 276 | scaffold_13 | 4860603 | intergenic | -26.9565 | |
| 277 | scaffold_13 | 6041158 | genebody | 6.4604 | USH2A |
| 278 | scaffold_13 | 6041184 | genebody | 5.0251 | USH2A |
| 279 | scaffold_13 | 6041286 | genebody | 8.0688 | USH2A |
| 280 | scaffold_13 | 6041287 | genebody | 8.8256 | USH2A |
| 281 | scaffold_13 | 6041351 | genebody | 11.5514 | USH2A |
| 282 | scaffold_13 | 6042569 | genebody | -9.4436 | USH2A |
| 283 | scaffold_13 | 6042585 | genebody | -13.7586 | USH2A |
| 284 | scaffold_13 | 7208103 | intergenic | -5.8519 | |
| 285 | scaffold_13 | 7208132 | intergenic | -6.2290 | |
| 286 | scaffold_13 | 17733695 | genebody;ge | 18.3333 | ACIN1 |
| 287 | scaffold_13 | 17733698 | genebody;ge | 21.3636 | ACIN1 |
| 288 | scaffold_13 | 22245032 | intergenic | 5.4166 | |
| 289 | scaffold_13 | 39768207 | intergenic | 6.5990 | |
| 290 | scaffold_130 | 2403177 | promoter | -17.0455 | |
| 291 | scaffold_130 | 2403185 | promoter | -11.3636 | |
| 292 | scaffold_130 | 2405969 | intergenic | 9.8739 | |
| 293 | scaffold_130 | 2406088 | intergenic | 5.1247 | |
| 294 | scaffold_137 | 2017366 | promoter | 11.3979 | ITPKB |
| 295 | scaffold_140 | 738226 | genebody;ge | 6.4968 | TIMP3;SYN3 |
| 296 | scaffold_140 | 985245 | genebody;ge | -21.1310 | LARGE1 |
| 297 | scaffold_140 | 985288 | genebody;ge | 5.7245 | LARGE1 |
| 298 | scaffold_140 | 985308 | genebody;ge | -20.3488 | LARGE1 |
| 299 | scaffold_143 | 83956 | intergenic | 5.1709 | |
| 300 | scaffold_143 | 1150856 | intergenic | 5.5263 | |
| 301 | scaffold_143 | 1151043 | intergenic | -15.7407 | |
| 302 | scaffold_143 | 1151082 | intergenic | 9.3750 | |
| 303 | scaffold_143 | 1151095 | intergenic | 25.2747 | |
| 304 | scaffold_144 | 884849 | intergenic | 9.5534 | |
| 305 | scaffold_144 | 2132696 | intergenic | 9.0391 | |
| 306 | scaffold_147 | 832384 | promoter | 9.7778 | INSYN1 |
| 307 | scaffold_149 | 1019161 | intergenic | -7.9919 | |
| 308 | scaffold_15 | 18434297 | genebody | -9.9435 | STOX1 |
| 309 | scaffold_15 | 18434585 | genebody | 10.7710 | STOX1 |
| 310 | scaffold_15 | 18434586 | genebody | -9.6618 | STOX1 |
| 311 | scaffold_15 | 18434638 | genebody | -11.8481 | STOX1 |
| 312 | scaffold_15 | 18440860 | genebody | 10.7385 | STOX1 |
| 313 | scaffold_15 | 29335091 | intergenic | 23.7132 | |

| | | | | | |
|---|---|---|---|---|---|
| 314 | scaffold_15 | 29335117 | intergenic | -7.2417 | |
| 315 | scaffold_15 | 33165439 | intergenic | -7.6923 | |
| 316 | scaffold_15 | 34393766 | genebody | 9.1119 | TBC1D22B |
| 317 | scaffold_15 | 37878846 | intergenic | 13.8921 | |
| 318 | scaffold_15 | 37878862 | intergenic | -9.3956 | |
| 319 | scaffold_15 | 40638129 | intergenic | -24.5192 | |
| 320 | scaffold_15 | 41538981 | genebody | 5.8227 | n/a |
| 321 | scaffold_150 | 1487385 | genebody | -10.0000 | NKD2 |
| 322 | scaffold_151 | 2093466 | intergenic | 7.2751 | |
| 323 | scaffold_156 | 889591 | genebody | -5.0188 | n/a |
| 324 | scaffold_156 | 2062811 | intergenic | 5.1833 | |
| 325 | scaffold_157 | 448186 | genebody | -6.8744 | n/a |
| 326 | scaffold_157 | 449416 | promoter | -10.0000 | |
| 327 | scaffold_157 | 449425 | promoter | -10.0000 | |
| 328 | scaffold_157 | 449668 | promoter | -7.7376 | |
| 329 | scaffold_157 | 449710 | promoter | 7.2330 | |
| 330 | scaffold_157 | 451189 | intergenic | 6.6667 | |
| 331 | scaffold_157 | 451191 | intergenic | 20.0000 | |
| 332 | scaffold_158 | 1702529 | intergenic | 5.8764 | |
| 333 | scaffold_158 | 1702678 | intergenic | 5.9524 | |
| 334 | scaffold_158 | 1702743 | intergenic | -7.5832 | |
| 335 | scaffold_1592 | 3072 | genebody | -22.7273 | n/a |
| 336 | scaffold_16 | 15313760 | genebody | 7.8938 | GATA3 |
| 337 | scaffold_16 | 15313768 | genebody | -19.0045 | GATA3 |
| 338 | scaffold_166 | 517969 | genebody | 21.3115 | SORCS2 |
| 339 | scaffold_166 | 1209922 | intergenic | 10.0280 | |
| 340 | scaffold_166 | 1366925 | intergenic | 12.7976 | |
| 341 | scaffold_166 | 1366929 | intergenic | 5.2381 | |
| 342 | scaffold_166 | 1367121 | intergenic | 16.0526 | |
| 343 | scaffold_166 | 1397405 | genebody | -23.2456 | DOK7 |
| 344 | scaffold_167 | 215537 | intergenic | -7.1429 | |
| 345 | scaffold_167 | 215587 | intergenic | -5.2217 | |
| 346 | scaffold_168 | 1567710 | genebody | 5.0000 | AKAP8L |
| 347 | scaffold_168 | 1567718 | genebody | -8.3333 | AKAP8L |
| 348 | scaffold_168 | 1567729 | genebody | -14.5833 | AKAP8L |
| 349 | scaffold_168 | 1569602 | genebody | 5.5618 | AKAP8L |
| 350 | scaffold_168 | 1569609 | genebody | 8.5571 | AKAP8L |
| 351 | scaffold_17 | 1455212 | intergenic | -5.4762 | |
| 352 | scaffold_17 | 7090429 | genebody | 5.7765 | FABP6 |
| 353 | scaffold_17 | 7091402 | genebody | -8.9364 | FABP6 |
| 354 | scaffold_17 | 7091416 | genebody | -9.1557 | FABP6 |
| 355 | scaffold_17 | 7091793 | genebody | -6.3799 | FABP6 |
| 356 | scaffold_17 | 37385648 | genebody;ge | -8.7500 | TRAPPC9 |
| 357 | scaffold_17 | 37385675 | genebody;ge | -18.7500 | TRAPPC9 |
| 358 | scaffold_17 | 37385681 | genebody;ge | -15.0000 | TRAPPC9 |

| | | | | | |
|---|---|---|---|---|---|
| 359 | scaffold_17 | 37385703 | genebody;ge | -8.7500 | TRAPPC9 |
| 360 | scaffold_17 | 37385710 | genebody;ge | -8.7500 | TRAPPC9 |
| 361 | scaffold_17 | 37385713 | genebody;ge | 9.0543 | TRAPPC9 |
| 362 | scaffold_17 | 37385717 | genebody;ge | 9.7990 | TRAPPC9 |
| 363 | scaffold_171 | 219389 | genebody | 5.2880 | RFX4 |
| 364 | scaffold_171 | 219390 | genebody | 16.0606 | RFX4 |
| 365 | scaffold_171 | 219440 | genebody | -5.8497 | RFX4 |
| 366 | scaffold_171 | 220999 | genebody | 5.2797 | RFX4 |
| 367 | scaffold_171 | 723712 | intergenic | -5.1750 | |
| 368 | scaffold_171 | 723747 | intergenic | -5.0906 | |
| 369 | scaffold_171 | 724380 | intergenic | 8.9286 | |
| 370 | scaffold_171 | 724392 | intergenic | 5.2273 | |
| 371 | scaffold_171 | 724436 | intergenic | 7.9100 | |
| 372 | scaffold_171 | 724441 | intergenic | 8.2264 | |
| 373 | scaffold_176 | 237869 | promoter | -5.1372 | MGAT4B |
| 374 | scaffold_176 | 237932 | genebody;ge | 10.6195 | MGAT4B |
| 375 | scaffold_18 | 2737971 | intergenic | -6.1624 | |
| 376 | scaffold_18 | 17737318 | intergenic | -6.1315 | |
| 377 | scaffold_18 | 38154126 | genebody; pr | -6.2500 | ;RNF208 |
| 378 | scaffold_18 | 38752273 | intergenic | 7.4286 | |
| 379 | scaffold_18 | 38752299 | intergenic | -7.8431 | |
| 380 | scaffold_18 | 38752495 | intergenic | 28.7407 | |
| 381 | scaffold_18 | 38752547 | intergenic | 17.3280 | |
| 382 | scaffold_18 | 38752726 | intergenic | 5.7937 | |
| 383 | scaffold_184 | 170759 | genebody;ge | -7.3340 | SHANK3 |
| 384 | scaffold_184 | 170790 | genebody;ge | 10.0000 | SHANK3 |
| 385 | scaffold_184 | 173410 | genebody;ge | -18.9991 | SHANK3 |
| 386 | scaffold_184 | 173433 | genebody;ge | -13.1579 | SHANK3 |
| 387 | scaffold_186 | 117181 | intergenic | -7.7915 | |
| 388 | scaffold_186 | 315322 | intergenic | 10.8065 | |
| 389 | scaffold_186 | 315323 | intergenic | -5.1760 | |
| 390 | scaffold_186 | 315326 | intergenic | -8.8710 | |
| 391 | scaffold_187 | 370889 | intergenic | -9.8296 | |
| 392 | scaffold_19 | 10971284 | intergenic | 8.3721 | |
| 393 | scaffold_19 | 10971462 | intergenic | 5.5233 | |
| 394 | scaffold_19 | 10971508 | intergenic | -7.3864 | |
| 395 | scaffold_19 | 11016580 | intergenic | -26.6667 | |
| 396 | scaffold_19 | 12934754 | intergenic | -9.0909 | |
| 397 | scaffold_19 | 14541155 | genebody | 8.7432 | NTM |
| 398 | scaffold_19 | 14561938 | genebody | -6.6294 | NTM |
| 399 | scaffold_19 | 26005068 | intergenic | 8.4615 | |
| 400 | scaffold_19 | 26005098 | intergenic | -16.5385 | |
| 401 | scaffold_19 | 26005456 | intergenic | 7.6087 | |
| 402 | scaffold_19 | 26007437 | intergenic | 13.0435 | |
| 403 | scaffold_19 | 26007474 | intergenic | 17.9348 | |

| | | | | | |
|---|---|---|---|---|---|
| 404 | scaffold_19 | 26007767 | intergenic | -9.4048 | |
| 405 | scaffold_19 | 28468722 | genebody | 5.2841 | NECTIN1 |
| 406 | scaffold_19 | 30245258 | genebody | 6.2791 | FXYD2 |
| 407 | scaffold_19 | 30245445 | genebody | -23.1951 | FXYD2 |
| 408 | scaffold_19 | 30245504 | genebody | 25.9485 | FXYD2 |
| 409 | scaffold_2 | 28475033 | promoter | -15.0000 | RNPC3 |
| 410 | scaffold_2 | 28475078 | promoter | 5.1136 | RNPC3 |
| 411 | scaffold_2 | 28475415 | promoter | 11.6667 | RNPC3 |
| 412 | scaffold_2 | 28509248 | genebody;ge | -5.4054 | |
| 413 | scaffold_2 | 28509273 | genebody;ge | -6.3830 | |
| 414 | scaffold_2 | 28509287 | genebody;ge | 8.8745 | |
| 415 | scaffold_2 | 28509288 | genebody;ge | -6.8836 | |
| 416 | scaffold_2 | 28540993 | genebody | -6.9865 | |
| 417 | scaffold_2 | 28541024 | genebody | -8.2050 | |
| 418 | scaffold_2 | 57547859 | genebody | 6.2356 | GIPC2 |
| 419 | scaffold_2 | 57547919 | genebody | -7.1018 | GIPC2 |
| 420 | scaffold_2 | 67316632 | intergenic | 18.3333 | |
| 421 | scaffold_2 | 67949260 | intergenic | 9.1757 | |
| 422 | scaffold_2 | 77645820 | intergenic | 10.6878 | |
| 423 | scaffold_2 | 77646041 | intergenic | 8.4672 | |
| 424 | scaffold_20 | 14157758 | promoter | 6.2667 | CCDC177 |
| 425 | scaffold_20 | 14157774 | promoter | -7.2196 | CCDC177 |
| 426 | scaffold_20 | 14157778 | promoter | 5.0451 | CCDC177 |
| 427 | scaffold_20 | 14157813 | promoter | -25.9740 | CCDC177 |
| 428 | scaffold_203 | 103315 | genebody | -26.9231 | TPCN1 |
| 429 | scaffold_2033 | 5574 | intergenic | 10.2543 | |
| 430 | scaffold_2033 | 5617 | intergenic | -7.0755 | |
| 431 | scaffold_2033 | 5908 | intergenic | -24.8918 | |
| 432 | scaffold_2033 | 5970 | intergenic | 9.5238 | |
| 433 | scaffold_2035 | 4980 | intergenic | -5.5812 | |
| 434 | scaffold_2035 | 4985 | intergenic | 13.4199 | |
| 435 | scaffold_2035 | 5027 | intergenic | 5.8110 | |
| 436 | scaffold_2035 | 5181 | intergenic | -5.5080 | |
| 437 | scaffold_21 | 24105270 | genebody;ge | 6.5934 | n/a |
| 438 | scaffold_21 | 31749772 | intergenic | 6.1832 | |
| 439 | scaffold_213 | 275950 | intergenic | -5.7143 | |
| 440 | scaffold_22 | 11200532 | intergenic | -5.7720 | |
| 441 | scaffold_22 | 11218832 | intergenic | -7.3152 | |
| 442 | scaffold_226 | 527608 | genebody | 7.5295 | ODAD3 |
| 443 | scaffold_226 | 527620 | genebody | 10.5391 | ODAD3 |
| 444 | scaffold_226 | 527623 | genebody | 10.7979 | ODAD3 |
| 445 | scaffold_226 | 527633 | genebody | -7.7592 | ODAD3 |
| 446 | scaffold_226 | 631257 | intergenic | -7.1429 | |
| 447 | scaffold_226 | 683497 | genebody | -8.3333 | ACP5 |
| 448 | scaffold_226 | 683500 | genebody | 18.1818 | ACP5 |

| | | | | | |
|---|---|---|---|---|---|
| 449 | scaffold_226 | 683542 | genebody | -6.7990 | ACP5 |
| 450 | scaffold_226 | 683602 | genebody | 10.6681 | ACP5 |
| 451 | scaffold_226 | 683617 | genebody | 7.3457 | ACP5 |
| 452 | scaffold_226 | 683751 | genebody | -9.1024 | ACP5 |
| 453 | scaffold_226 | 683756 | genebody | 10.5735 | ACP5 |
| 454 | scaffold_23 | 25112886 | intergenic | -5.6843 | |
| 455 | scaffold_23 | 29612465 | genebody | -6.4219 | LEO1 |
| 456 | scaffold_23 | 29838569 | genebody;ge | 27.7778 | MYO5A |
| 457 | scaffold_24 | 1230202 | genebody | -21.0227 | MGAT4A |
| 458 | scaffold_24 | 1230225 | genebody | 9.6899 | MGAT4A |
| 459 | scaffold_24 | 1230230 | genebody | 5.3426 | MGAT4A |
| 460 | scaffold_24 | 1230273 | genebody | -5.9091 | MGAT4A |
| 461 | scaffold_24 | 1230301 | genebody | 8.6998 | MGAT4A |
| 462 | scaffold_24 | 2682732 | genebody | -6.3058 | FAHD2A |
| 463 | scaffold_24 | 2682733 | genebody | 7.3885 | FAHD2A |
| 464 | scaffold_24 | 2682750 | genebody | 8.5798 | FAHD2A |
| 465 | scaffold_249 | 6064 | intergenic | 8.2850 | |
| 466 | scaffold_249 | 6354 | intergenic | -18.3871 | |
| 467 | scaffold_249 | 6356 | intergenic | -19.3676 | |
| 468 | scaffold_249 | 6390 | intergenic | -18.7135 | |
| 469 | scaffold_249 | 6424 | intergenic | -5.1170 | |
| 470 | scaffold_25 | 11383 | intergenic | -5.9163 | |
| 471 | scaffold_25 | 11424 | intergenic | 14.6859 | |
| 472 | scaffold_25 | 11426 | intergenic | 9.5838 | |
| 473 | scaffold_25 | 1238820 | intergenic | -7.0659 | |
| 474 | scaffold_25 | 7022003 | intergenic | -7.6184 | |
| 475 | scaffold_25 | 7022200 | intergenic | -12.8674 | |
| 476 | scaffold_25 | 7022266 | intergenic | 5.5574 | |
| 477 | scaffold_25 | 7028459 | intergenic | 7.9853 | |
| 478 | scaffold_25 | 8761266 | genebody;ge | 5.2063 | AHDC1 |
| 479 | scaffold_25 | 8761310 | genebody;ge | -5.4144 | AHDC1 |
| 480 | scaffold_25 | 8761470 | genebody;ge | 10.8615 | AHDC1 |
| 481 | scaffold_25 | 11569167 | genebody;ge | -6.6667 | MYOM3 |
| 482 | scaffold_25 | 11569206 | genebody;ge | 7.1053 | MYOM3 |
| 483 | scaffold_25 | 11570287 | genebody;ge | -6.7382 | MYOM3 |
| 484 | scaffold_25 | 11570288 | genebody;ge | 8.0000 | MYOM3 |
| 485 | scaffold_25 | 11570307 | genebody;ge | -6.6411 | MYOM3 |
| 486 | scaffold_25 | 11570920 | genebody;ge | -15.9341 | MYOM3 |
| 487 | scaffold_25 | 11571279 | genebody;ge | -5.3571 | MYOM3 |
| 488 | scaffold_25 | 16670998 | genebody | -31.4286 | IGSF21 |
| 489 | scaffold_25 | 16670999 | genebody | 7.0724 | IGSF21 |
| 490 | scaffold_25 | 16671047 | genebody | 10.9508 | IGSF21 |
| 491 | scaffold_25 | 16996539 | intergenic | 6.2304 | |
| 492 | scaffold_25 | 17887006 | genebody;ge | 5.0000 | EPHA2 |
| 493 | scaffold_25 | 17887025 | genebody;ge | -8.3333 | EPHA2 |

| | | | | | |
|---|---|---|---|---|---|
| 494 | scaffold_25 | 17962742 | intergenic | -10.0390 | |
| 495 | scaffold_25 | 17963135 | intergenic | -5.0000 | |
| 496 | scaffold_25 | 18474719 | genebody | 23.2143 | FHAD1 |
| 497 | scaffold_25 | 23409078 | genebody | -7.0189 | PEX14 |
| 498 | scaffold_25 | 23409107 | genebody | 6.5657 | PEX14 |
| 499 | scaffold_25 | 23409138 | genebody | -6.3278 | PEX14 |
| 500 | scaffold_25 | 24069958 | intergenic | 9.6257 | |
| 501 | scaffold_25 | 24069971 | intergenic | 5.6005 | |
| 502 | scaffold_25 | 24364742 | intergenic | -15.4762 | |
| 503 | scaffold_25 | 26209813 | genebody | -9.7028 | ESPN |
| 504 | scaffold_25 | 26209818 | genebody | -23.7672 | ESPN |
| 505 | scaffold_25 | 26209843 | genebody | -16.2448 | ESPN |
| 506 | scaffold_25 | 26209855 | genebody | -5.4422 | ESPN |
| 507 | scaffold_25 | 26209856 | genebody | -12.0623 | ESPN |
| 508 | scaffold_25 | 26209860 | genebody | -12.0076 | ESPN |
| 509 | scaffold_25 | 26209876 | genebody | -17.8874 | ESPN |
| 510 | scaffold_25 | 26209882 | genebody | -11.8179 | ESPN |
| 511 | scaffold_25 | 26211125 | genebody | 6.8760 | ESPN |
| 512 | scaffold_25 | 26211165 | genebody | 6.5139 | ESPN |
| 513 | scaffold_25 | 26211235 | genebody | 9.0553 | ESPN |
| 514 | scaffold_25 | 26211251 | genebody | 5.9723 | ESPN |
| 515 | scaffold_25 | 26414759 | genebody;ge | -5.0551 | CHD5 |
| 516 | scaffold_25 | 27347361 | genebody | -5.8953 | AJAP1 |
| 517 | scaffold_25 | 27347393 | genebody | 5.1136 | AJAP1 |
| 518 | scaffold_255 | 162093 | intergenic | -5.1982 | |
| 519 | scaffold_255 | 162147 | intergenic | 7.9741 | |
| 520 | scaffold_27 | 569835 | genebody | 8.4572 | MRPS2 |
| 521 | scaffold_27 | 817038 | genebody | 10.5011 | RALGDS |
| 522 | scaffold_27 | 817055 | genebody | 7.8092 | RALGDS |
| 523 | scaffold_27 | 817061 | genebody | 8.6839 | RALGDS |
| 524 | scaffold_27 | 1292872 | genebody | -8.9744 | CFAP77 |
| 525 | scaffold_27 | 1293083 | genebody | -12.5125 | CFAP77 |
| 526 | scaffold_27 | 1293121 | genebody | -12.1588 | CFAP77 |
| 527 | scaffold_27 | 1730450 | intergenic | 17.8571 | |
| 528 | scaffold_27 | 1730493 | intergenic | 10.0000 | |
| 529 | scaffold_27 | 1730725 | intergenic | 6.0821 | |
| 530 | scaffold_27 | 1730781 | intergenic | -8.8580 | |
| 531 | scaffold_27 | 1730796 | intergenic | 9.6283 | |
| 532 | scaffold_27 | 3631606 | genebody | 11.1345 | DYNC2I2 |
| 533 | scaffold_27 | 3631614 | genebody | 16.4286 | DYNC2I2 |
| 534 | scaffold_27 | 3631633 | genebody | 15.7143 | DYNC2I2 |
| 535 | scaffold_27 | 9691097 | genebody | -14.7619 | NEK6 |
| 536 | scaffold_27 | 9691327 | genebody | -23.1250 | CNTFR |
| 537 | scaffold_27 | 9691381 | genebody | -21.5461 | CNTFR |
| 538 | scaffold_27 | 9691385 | genebody | -8.8816 | CNTFR |

| | | | | | |
|---|---|---|---|---|---|
| 539 | scaffold_27 | 14983683 | promoter | 9.1882 | CNTFR |
| 540 | scaffold_27 | 14983883 | promoter | 11.6930 | CNTFR |
| 541 | scaffold_27 | 14983904 | promoter | 9.6126 | CNTFR |
| 542 | scaffold_27 | 17140018 | intergenic | -6.5627 | |
| 543 | scaffold_28 | 6812264 | intergenic | -7.5521 | |
| 544 | scaffold_28 | 14915211 | genebody | 9.3750 | RECQL |
| 545 | scaffold_28 | 21356440 | intergenic | -18.7783 | |
| 546 | scaffold_28 | 21356457 | intergenic | 17.6471 | |
| 547 | scaffold_28 | 21356494 | intergenic | 7.3529 | |
| 548 | scaffold_28 | 21679443 | genebody | 12.7574 | TMPRSS6 |
| 549 | scaffold_28 | 21679569 | genebody | -11.4490 | TMPRSS6 |
| 550 | scaffold_28 | 21891185 | genebody | 5.2632 | |
| 551 | scaffold_28 | 21891230 | genebody | 13.6364 | |
| 552 | scaffold_28 | 21891446 | promoter | -8.6957 | |
| 553 | scaffold_28 | 21891460 | promoter | -18.3333 | |
| 554 | scaffold_28 | 21891491 | promoter | -8.0000 | |
| 555 | scaffold_28 | 22198197 | genebody;ge | 8.3333 | |
| 556 | scaffold_28 | 27357739 | genebody | 6.0520 | EFCAB6 |
| 557 | scaffold_283 | 343141 | genebody | 9.9662 | STAB2 |
| 558 | scaffold_283 | 343327 | genebody | -13.6725 | STAB2 |
| 559 | scaffold_3 | 11303918 | intergenic | -7.8283 | |
| 560 | scaffold_3 | 11329419 | intergenic | -17.1429 | |
| 561 | scaffold_3 | 11329433 | intergenic | 10.9524 | |
| 562 | scaffold_3 | 18813662 | genebody | -11.8644 | GAD1 |
| 563 | scaffold_3 | 18813682 | genebody | -6.2724 | GAD1 |
| 564 | scaffold_3 | 18813705 | genebody | -12.4496 | GAD1 |
| 565 | scaffold_3 | 18814469 | genebody | 6.1905 | GAD1 |
| 566 | scaffold_3 | 18814704 | genebody | -8.9862 | GAD1 |
| 567 | scaffold_3 | 19719998 | promoter | -5.8824 | KLHL23 |
| 568 | scaffold_3 | 19720534 | promoter | 9.5238 | KLHL23 |
| 569 | scaffold_3 | 19720548 | promoter | -9.5238 | KLHL23 |
| 570 | scaffold_3 | 19720884 | promoter | 25.5652 | KLHL23 |
| 571 | scaffold_3 | 19730783 | intergenic | -9.5370 | |
| 572 | scaffold_3 | 19733656 | intergenic | -25.6944 | |
| 573 | scaffold_3 | 19746232 | genebody | -5.8824 | CFAP210 |
| 574 | scaffold_3 | 19746254 | genebody | 6.2500 | CFAP210 |
| 575 | scaffold_3 | 19746272 | genebody | -5.8824 | CFAP210 |
| 576 | scaffold_3 | 19746514 | genebody | -9.5238 | CFAP210 |
| 577 | scaffold_3 | 19746564 | genebody | -19.0476 | CFAP210 |
| 578 | scaffold_3 | 19757807 | genebody | 5.6152 | CFAP210 |
| 579 | scaffold_3 | 19787899 | genebody | 7.5169 | PPIG |
| 580 | scaffold_3 | 19787932 | genebody | -9.5439 | PPIG |
| 581 | scaffold_3 | 51971108 | genebody | 10.4441 | ACMSD |
| 582 | scaffold_3 | 51978415 | genebody | -8.1529 | ACMSD |
| 583 | scaffold_30 | 3585195 | genebody | -9.4886 | MEPCE |

| | | | | | |
|---|---|---|---|---|---|
| 584 | scaffold_30 | 3585212 | genebody | 6.7120 | MEPCE |
| 585 | scaffold_30 | 3585264 | genebody | -16.7153 | MEPCE |
| 586 | scaffold_30 | 3585266 | genebody | -7.6572 | MEPCE |
| 587 | scaffold_30 | 7754868 | intergenic | -17.8571 | |
| 588 | scaffold_30 | 7924709 | intergenic | -6.3218 | |
| 589 | scaffold_304 | 292696 | intergenic | -7.9132 | |
| 590 | scaffold_304 | 292751 | intergenic | -7.5094 | |
| 591 | scaffold_304 | 292752 | intergenic | -5.8028 | |
| 592 | scaffold_31 | 2988859 | intergenic | 5.8824 | |
| 593 | scaffold_31 | 2988932 | intergenic | -8.4311 | |
| 594 | scaffold_31 | 5384849 | intergenic | 30.0395 | |
| 595 | scaffold_31 | 5385130 | intergenic | -27.8431 | |
| 596 | scaffold_31 | 5395688 | intergenic | 18.1818 | |
| 597 | scaffold_31 | 5413472 | genebody | 10.4601 | SOSTDC1 |
| 598 | scaffold_31 | 9017776 | intergenic | 6.7158 | |
| 599 | scaffold_31 | 9017839 | intergenic | 6.1012 | |
| 600 | scaffold_31 | 9029914 | intergenic | 23.2493 | |
| 601 | scaffold_31 | 9029942 | intergenic | 26.6106 | |
| 602 | scaffold_31 | 12011006 | intergenic | -20.8791 | |
| 603 | scaffold_31 | 12011050 | intergenic | 7.6923 | |
| 604 | scaffold_31 | 12011448 | intergenic | 6.5455 | |
| 605 | scaffold_31 | 12014613 | intergenic | 9.9379 | |
| 606 | scaffold_31 | 12029150 | intergenic | -17.2727 | |
| 607 | scaffold_31 | 12035314 | intergenic | -5.2074 | |
| 608 | scaffold_31 | 12075815 | intergenic | 5.3191 | |
| 609 | scaffold_31 | 12075879 | intergenic | -10.9961 | |
| 610 | scaffold_31 | 12080277 | intergenic | -5.3062 | |
| 611 | scaffold_31 | 12080653 | intergenic | -5.9342 | |
| 612 | scaffold_31 | 12080684 | intergenic | -5.9342 | |
| 613 | scaffold_31 | 12094541 | intergenic | 18.4211 | |
| 614 | scaffold_31 | 12094872 | intergenic | 12.1441 | |
| 615 | scaffold_31 | 12095113 | intergenic | 15.0752 | |
| 616 | scaffold_31 | 12103623 | intergenic | 8.3333 | |
| 617 | scaffold_31 | 12121268 | genebody | -5.2177 | |
| 618 | scaffold_31 | 12124123 | genebody | -8.7912 | |
| 619 | scaffold_31 | 12124180 | genebody | 11.8132 | |
| 620 | scaffold_31 | 12137419 | genebody | -6.1538 | |
| 621 | scaffold_31 | 12137429 | genebody | -5.6410 | |
| 622 | scaffold_31 | 14521680 | intergenic | 10.0000 | |
| 623 | scaffold_31 | 14524323 | intergenic | 10.9848 | |
| 624 | scaffold_31 | 14524353 | intergenic | 33.2727 | |
| 625 | scaffold_31 | 14581579 | intergenic | 11.4865 | |
| 626 | scaffold_31 | 14581588 | intergenic | 12.0946 | |
| 627 | scaffold_31 | 14581593 | intergenic | 8.3784 | |
| 628 | scaffold_31 | 14591436 | intergenic | -10.0000 | |

| | | | | | |
|---|---|---|---|---|---|
| 629 | scaffold_31 | 17479190 | intergenic | 9.8116 | |
| 630 | scaffold_31 | 19690036 | promoter | 13.8889 | |
| 631 | scaffold_31 | 19690037 | promoter | 7.5825 | |
| 632 | scaffold_32 | 65168 | intergenic | 17.9348 | |
| 633 | scaffold_32 | 65194 | intergenic | -5.1630 | |
| 634 | scaffold_32 | 19461771 | intergenic | 12.6984 | |
| 635 | scaffold_32 | 19461802 | intergenic | 5.0362 | |
| 636 | scaffold_32 | 19461864 | intergenic | -6.5934 | |
| 637 | scaffold_32 | 19461946 | intergenic | -6.6828 | |
| 638 | scaffold_32 | 20114717 | genebody | -5.2817 | MNT |
| 639 | scaffold_32 | 20114745 | genebody | -6.3648 | MNT |
| 640 | scaffold_32 | 20114786 | genebody | 5.3070 | MNT |
| 641 | scaffold_32 | 20812492 | genebody | 11.7943 | SLC43A2 |
| 642 | scaffold_32 | 20812722 | genebody | 6.7177 | SLC43A2 |
| 643 | scaffold_32 | 20812739 | genebody | 7.4941 | SLC43A2 |
| 644 | scaffold_32 | 23946888 | promoter | 18.1582 | FOXN1 |
| 645 | scaffold_32 | 23947632 | promoter | 6.0269 | FOXN1 |
| 646 | scaffold_33 | 24415612 | intergenic | -7.3214 | |
| 647 | scaffold_33 | 24466781 | intergenic | 12.6812 | |
| 648 | scaffold_334 | 104291 | intergenic | 15.4237 | |
| 649 | scaffold_334 | 104298 | intergenic | 6.0725 | |
| 650 | scaffold_334 | 406154 | intergenic | -13.8761 | |
| 651 | scaffold_334 | 406191 | intergenic | -13.8933 | |
| 652 | scaffold_334 | 406251 | intergenic | 5.5534 | |
| 653 | scaffold_337 | 18200 | intergenic | -6.8831 | |
| 654 | scaffold_34 | 3046261 | intergenic | -8.6957 | |
| 655 | scaffold_34 | 3046280 | intergenic | -6.6667 | |
| 656 | scaffold_34 | 3046489 | intergenic | -12.9032 | |
| 657 | scaffold_34 | 20364508 | genebody | 7.8031 | EPHA3 |
| 658 | scaffold_34 | 20709533 | intergenic | 6.2500 | |
| 659 | scaffold_34 | 20709829 | intergenic | -7.1429 | |
| 660 | scaffold_34 | 20711283 | intergenic | 12.7159 | |
| 661 | scaffold_356 | 220479 | promoter | -6.3955 | |
| 662 | scaffold_356 | 220749 | promoter | -23.8095 | |
| 663 | scaffold_356 | 220753 | promoter | 14.2857 | |
| 664 | scaffold_356 | 220802 | promoter | 26.9872 | |
| 665 | scaffold_356 | 245739 | genebody;ge | 24.3056 | ARID3A |
| 666 | scaffold_356 | 245765 | genebody;ge | 22.2222 | ARID3A |
| 667 | scaffold_356 | 245982 | genebody;ge | 8.3096 | ARID3A |
| 668 | scaffold_36 | 3636877 | intergenic | 6.2500 | |
| 669 | scaffold_368 | 224088 | genebody | 8.5017 | TYK2 |
| 670 | scaffold_38 | 537094 | intergenic | 15.3846 | |
| 671 | scaffold_38 | 6902950 | genebody | -5.6892 | |
| 672 | scaffold_38 | 11069305 | genebody | 16.6667 | DTNBP1 |
| 673 | scaffold_38 | 11069344 | genebody | -9.0909 | DTNBP1 |

| | | | | | |
|---|---|---|---|---|---|
| 674 | scaffold_38 | 11089043 | genebody | 9.3434 | DTNBP1 |
| 675 | scaffold_38 | 11089709 | genebody | 6.5574 | DTNBP1 |
| 676 | scaffold_38 | 11106070 | genebody | -9.4917 | DTNBP1 |
| 677 | scaffold_38 | 15091931 | genebody | 27.5000 | MBOAT1 |
| 678 | scaffold_38 | 15092250 | promoter | 25.0458 | MBOAT1 |
| 679 | scaffold_38 | 15092361 | promoter | 9.5652 | MBOAT1 |
| 680 | scaffold_38 | 15092418 | promoter | 16.9052 | MBOAT1 |
| 681 | scaffold_38 | 15092732 | promoter | -7.7778 | MBOAT1 |
| 682 | scaffold_38 | 21165293 | intergenic | -9.7399 | |
| 683 | scaffold_38 | 21165297 | intergenic | 8.6690 | |
| 684 | scaffold_4 | 1103221 | intergenic | 12.3913 | |
| 685 | scaffold_4 | 6987992 | intergenic | 8.1908 | |
| 686 | scaffold_4 | 7002910 | genebody | 6.3768 | |
| 687 | scaffold_4 | 7007664 | intergenic | 8.2234 | |
| 688 | scaffold_4 | 15490787 | intergenic | -9.3333 | |
| 689 | scaffold_4 | 15490864 | intergenic | -5.2489 | |
| 690 | scaffold_4 | 18836653 | intergenic | 20.9957 | |
| 691 | scaffold_4 | 18836654 | intergenic | -22.5287 | |
| 692 | scaffold_4 | 18837025 | intergenic | -7.7395 | |
| 693 | scaffold_4 | 18837026 | intergenic | 5.9048 | |
| 694 | scaffold_4 | 19064480 | genebody;ge | 8.4167 | ARVCF |
| 695 | scaffold_4 | 19064543 | genebody;ge | 5.0404 | ARVCF |
| 696 | scaffold_4 | 28213454 | intergenic | 5.2344 | |
| 697 | scaffold_4 | 43658158 | genebody | -5.0132 | IL4R |
| 698 | scaffold_4 | 43658197 | genebody | -7.3413 | IL4R |
| 699 | scaffold_4 | 43658555 | genebody | 11.5385 | IL4R |
| 700 | scaffold_4 | 43658576 | genebody | 8.0000 | IL4R |
| 701 | scaffold_4 | 43658589 | genebody | 8.3333 | IL4R |
| 702 | scaffold_4 | 43660321 | genebody | 6.2506 | IL4R |
| 703 | scaffold_4 | 43660358 | genebody | 11.9360 | IL4R |
| 704 | scaffold_4 | 43660450 | genebody | 20.7280 | IL4R |
| 705 | scaffold_4 | 43660515 | genebody | 11.2614 | IL4R |
| 706 | scaffold_4 | 43662021 | genebody | 5.0000 | IL4R |
| 707 | scaffold_4 | 43662295 | genebody | 5.2632 | IL4R |
| 708 | scaffold_4 | 43662460 | genebody | 7.1483 | IL4R |
| 709 | scaffold_4 | 43662517 | genebody | -5.3695 | IL4R |
| 710 | scaffold_4 | 43662690 | genebody | 6.9841 | IL4R |
| 711 | scaffold_4 | 43662802 | genebody | -5.5760 | IL4R |
| 712 | scaffold_4 | 43662895 | genebody | 6.2008 | IL4R |
| 713 | scaffold_4 | 43663628 | genebody | 8.7500 | IL4R |
| 714 | scaffold_4 | 43663975 | genebody | 5.5615 | IL4R |
| 715 | scaffold_4 | 43665148 | intergenic | 10.4356 | |
| 716 | scaffold_4 | 43665301 | intergenic | 6.4950 | |
| 717 | scaffold_4 | 43666602 | intergenic | 6.8132 | |
| 718 | scaffold_4 | 43668936 | intergenic | -10.4167 | |

| | | | | | |
|---|---|---|---|---|---|
| 719 | scaffold_4 | 43668991 | intergenic | 5.0147 | |
| 720 | scaffold_4 | 43669160 | intergenic | 5.8442 | |
| 721 | scaffold_4 | 43669161 | intergenic | 8.8991 | |
| 722 | scaffold_4 | 43671870 | intergenic | 7.1161 | |
| 723 | scaffold_4 | 43671889 | intergenic | -6.4342 | |
| 724 | scaffold_4 | 43671946 | intergenic | 7.1819 | |
| 725 | scaffold_4 | 43671947 | intergenic | -6.0859 | |
| 726 | scaffold_4 | 43672206 | intergenic | 9.2448 | |
| 727 | scaffold_4 | 43672292 | intergenic | -7.4585 | |
| 728 | scaffold_4 | 43672334 | intergenic | 6.3425 | |
| 729 | scaffold_4 | 43672335 | intergenic | -8.8094 | |
| 730 | scaffold_4 | 43672657 | intergenic | 15.1947 | |
| 731 | scaffold_4 | 43672658 | intergenic | 8.0473 | |
| 732 | scaffold_4 | 43672763 | intergenic | -14.5888 | |
| 733 | scaffold_4 | 43672769 | intergenic | -9.4304 | |
| 734 | scaffold_4 | 43672808 | intergenic | -12.6429 | |
| 735 | scaffold_4 | 43672810 | intergenic | -9.0929 | |
| 736 | scaffold_4 | 43672823 | intergenic | -6.9818 | |
| 737 | scaffold_4 | 43672842 | intergenic | 10.1120 | |
| 738 | scaffold_4 | 43672843 | intergenic | -7.0549 | |
| 739 | scaffold_4 | 43672862 | intergenic | 12.3372 | |
| 740 | scaffold_4 | 43672863 | intergenic | -7.7429 | |
| 741 | scaffold_4 | 43672894 | intergenic | 9.7500 | |
| 742 | scaffold_4 | 43672895 | intergenic | -5.7961 | |
| 743 | scaffold_4 | 43672909 | intergenic | -9.3703 | |
| 744 | scaffold_4 | 43672912 | intergenic | -7.4522 | |
| 745 | scaffold_4 | 43674500 | intergenic | -10.9337 | |
| 746 | scaffold_4 | 43674528 | intergenic | 7.5348 | |
| 747 | scaffold_4 | 43674749 | intergenic | -13.6907 | |
| 748 | scaffold_4 | 43678467 | intergenic | 5.7545 | |
| 749 | scaffold_4 | 43678510 | intergenic | 15.7289 | |
| 750 | scaffold_4 | 43679422 | intergenic | -6.1170 | |
| 751 | scaffold_4 | 43679466 | intergenic | 5.4657 | |
| 752 | scaffold_4 | 43679512 | intergenic | -12.6529 | |
| 753 | scaffold_4 | 43679521 | intergenic | -5.2952 | |
| 754 | scaffold_4 | 43679649 | intergenic | 6.8353 | |
| 755 | scaffold_4 | 43701365 | promoter | 12.0346 | IL21R |
| 756 | scaffold_4 | 43704868 | genebody | -12.3684 | IL21R |
| 757 | scaffold_4 | 43706939 | genebody | 8.6486 | IL21R |
| 758 | scaffold_4 | 43706940 | genebody | -5.3571 | IL21R |
| 759 | scaffold_4 | 43707012 | genebody | 5.0595 | IL21R |
| 760 | scaffold_4 | 43707013 | genebody | 5.4736 | IL21R |
| 761 | scaffold_4 | 43707234 | genebody | 12.0061 | IL21R |
| 762 | scaffold_4 | 43707263 | genebody | 11.4995 | IL21R |
| 763 | scaffold_4 | 43716745 | genebody | 7.9743 | IL21R |

| | | | | | |
|---|---|---|---|---|---|
| 764 | scaffold_4 | 43716765 | genebody | 5.3486 | IL21R |
| 765 | scaffold_4 | 43716791 | genebody | 9.3366 | IL21R |
| 766 | scaffold_4 | 43716990 | genebody | 11.4067 | IL21R |
| 767 | scaffold_4 | 43717013 | genebody | 8.4631 | IL21R |
| 768 | scaffold_4 | 43717064 | genebody | -7.3950 | IL21R |
| 769 | scaffold_4 | 43717396 | genebody | -10.2222 | IL21R |
| 770 | scaffold_4 | 43717444 | genebody | 9.3968 | IL21R |
| 771 | scaffold_4 | 43726477 | genebody | 10.3393 | IL21R |
| 772 | scaffold_4 | 43730755 | genebody | 6.5055 | IL21R |
| 773 | scaffold_4 | 43730885 | genebody | -5.2318 | IL21R |
| 774 | scaffold_4 | 43730912 | genebody | -5.2288 | IL21R |
| 775 | scaffold_4 | 43731129 | genebody | -6.1376 | IL21R |
| 776 | scaffold_4 | 43731167 | genebody | 7.0376 | IL21R |
| 777 | scaffold_4 | 43731354 | genebody | -7.1622 | IL21R |
| 778 | scaffold_4 | 43731452 | genebody | -10.0684 | IL21R |
| 779 | scaffold_4 | 43731471 | genebody | -17.2917 | IL21R |
| 780 | scaffold_4 | 43731653 | genebody | 7.4074 | IL21R |
| 781 | scaffold_4 | 43731699 | genebody | -8.3333 | IL21R |
| 782 | scaffold_4 | 43732455 | genebody | -10.0000 | IL21R |
| 783 | scaffold_4 | 43733293 | genebody | 8.5862 | IL21R |
| 784 | scaffold_4 | 43733403 | genebody | 9.0423 | IL21R |
| 785 | scaffold_4 | 43735858 | intergenic | -12.5000 | |
| 786 | scaffold_4 | 43735905 | intergenic | -22.0374 | |
| 787 | scaffold_4 | 43735912 | intergenic | 6.9993 | |
| 788 | scaffold_4 | 43737303 | intergenic | 8.1731 | |
| 789 | scaffold_4 | 43738058 | genebody | -5.7895 | |
| 790 | scaffold_4 | 43738119 | genebody | -6.3158 | |
| 791 | scaffold_4 | 43738424 | genebody | -10.1307 | |
| 792 | scaffold_4 | 43739792 | genebody | 7.8947 | |
| 793 | scaffold_4 | 43740613 | genebody | -21.5881 | |
| 794 | scaffold_4 | 43743314 | genebody | -8.8815 | |
| 795 | scaffold_4 | 43743320 | genebody | 6.3141 | |
| 796 | scaffold_4 | 43743321 | genebody | -8.3333 | |
| 797 | scaffold_4 | 43744878 | genebody | -9.0909 | |
| 798 | scaffold_4 | 43745041 | genebody | -15.0769 | |
| 799 | scaffold_4 | 43745108 | genebody | 6.0606 | |
| 800 | scaffold_4 | 43745817 | genebody | -7.3333 | |
| 801 | scaffold_4 | 43745840 | genebody | -10.0000 | |
| 802 | scaffold_4 | 43966062 | genebody;ge | -10.4575 | KATNIP |
| 803 | scaffold_4 | 43966108 | genebody;ge | -16.6667 | KATNIP |
| 804 | scaffold_4 | 43967152 | genebody;ge | -5.0311 | KATNIP |
| 805 | scaffold_4 | 43976249 | genebody;ge | 7.2896 | KATNIP |
| 806 | scaffold_4 | 43976267 | genebody;ge | 7.7268 | KATNIP |
| 807 | scaffold_4 | 43979316 | genebody;ge | -9.3137 | KATNIP |
| 808 | scaffold_4 | 43979640 | genebody;ge | 6.6667 | KATNIP |

| | | | | | |
|---|---|---|---|---|---|
| 809 | scaffold_4 | 43983740 | genebody;ge | 5.5732 | KATNIP |
| 810 | scaffold_4 | 43983903 | genebody;ge | -5.4622 | KATNIP |
| 811 | scaffold_4 | 43983993 | genebody;ge | 5.3181 | KATNIP |
| 812 | scaffold_4 | 43985712 | genebody;ge | 8.6250 | KATNIP |
| 813 | scaffold_4 | 43987106 | genebody;ge | 13.1579 | KATNIP |
| 814 | scaffold_4 | 43987132 | genebody;ge | 6.4016 | KATNIP |
| 815 | scaffold_4 | 43987161 | genebody;ge | 6.4590 | KATNIP |
| 816 | scaffold_4 | 43987304 | genebody;ge | 11.7647 | KATNIP |
| 817 | scaffold_4 | 43993691 | genebody;ge | 5.9066 | KATNIP |
| 818 | scaffold_4 | 43993736 | genebody;ge | 5.5556 | KATNIP |
| 819 | scaffold_4 | 43993813 | genebody;ge | -6.6722 | KATNIP |
| 820 | scaffold_4 | 43993819 | genebody;ge | -7.5758 | KATNIP |
| 821 | scaffold_4 | 43993831 | genebody;ge | -6.8753 | KATNIP |
| 822 | scaffold_4 | 43993838 | genebody;ge | -6.1560 | KATNIP |
| 823 | scaffold_4 | 43993849 | genebody;ge | -11.5638 | KATNIP |
| 824 | scaffold_4 | 43993868 | genebody;ge | -13.6194 | KATNIP |
| 825 | scaffold_4 | 43993878 | genebody;ge | -5.0792 | KATNIP |
| 826 | scaffold_4 | 43993879 | genebody;ge | -7.8621 | KATNIP |
| 827 | scaffold_4 | 43993940 | genebody;ge | -8.3673 | KATNIP |
| 828 | scaffold_4 | 43993944 | genebody;ge | 9.7565 | KATNIP |
| 829 | scaffold_4 | 43993989 | genebody;ge | -8.8649 | KATNIP |
| 830 | scaffold_4 | 43994015 | genebody;ge | -6.3636 | KATNIP |
| 831 | scaffold_4 | 43994199 | genebody;ge | 22.7642 | KATNIP |
| 832 | scaffold_4 | 43994229 | genebody;ge | 8.3810 | KATNIP |
| 833 | scaffold_4 | 43994234 | genebody;ge | 15.0794 | KATNIP |
| 834 | scaffold_4 | 43994250 | genebody;ge | 17.4603 | KATNIP |
| 835 | scaffold_4 | 43994257 | genebody;ge | 15.0794 | KATNIP |
| 836 | scaffold_4 | 43999404 | genebody;ge | 6.1538 | KATNIP |
| 837 | scaffold_4 | 43999584 | genebody;ge | -8.6761 | KATNIP |
| 838 | scaffold_4 | 44000886 | genebody;ge | -8.3333 | KATNIP |
| 839 | scaffold_4 | 44003196 | genebody;ge | -11.0853 | KATNIP |
| 840 | scaffold_4 | 44004604 | intergenic | 7.9304 | |
| 841 | scaffold_4 | 44005260 | intergenic | 10.0652 | |
| 842 | scaffold_4 | 44005323 | intergenic | 6.0606 | |
| 843 | scaffold_4 | 44005324 | intergenic | 13.6797 | |
| 844 | scaffold_4 | 44006798 | intergenic | 6.6667 | |
| 845 | scaffold_4 | 44006807 | intergenic | 6.6667 | |
| 846 | scaffold_4 | 44007168 | intergenic | 7.8848 | |
| 847 | scaffold_4 | 44007215 | intergenic | -7.5646 | |
| 848 | scaffold_4 | 44007231 | intergenic | -5.6586 | |
| 849 | scaffold_4 | 44007259 | intergenic | 5.8015 | |
| 850 | scaffold_4 | 44007303 | intergenic | -11.8452 | |
| 851 | scaffold_4 | 44007338 | intergenic | -7.0238 | |
| 852 | scaffold_4 | 44010485 | intergenic | 10.7143 | |
| 853 | scaffold_4 | 44010531 | intergenic | 8.9330 | |

| | | | | | |
|---|---|---|---|---|---|
| 854 | scaffold_4 | 44010534 | intergenic | -5.0000 | |
| 855 | scaffold_4 | 44010535 | intergenic | 5.1282 | |
| 856 | scaffold_4 | 44010550 | intergenic | 11.7949 | |
| 857 | scaffold_4 | 44014943 | genebody | 7.0659 | GSG1L |
| 858 | scaffold_4 | 44015783 | genebody | -9.5238 | GSG1L |
| 859 | scaffold_4 | 44015791 | genebody | 7.5758 | GSG1L |
| 860 | scaffold_4 | 44015818 | genebody | 12.6263 | GSG1L |
| 861 | scaffold_4 | 44015839 | genebody | 6.7822 | GSG1L |
| 862 | scaffold_4 | 44016815 | genebody | -6.1151 | GSG1L |
| 863 | scaffold_4 | 44016839 | genebody | 6.8568 | GSG1L |
| 864 | scaffold_4 | 44018185 | genebody | -22.6190 | GSG1L |
| 865 | scaffold_4 | 44018489 | genebody | -9.7619 | GSG1L |
| 866 | scaffold_4 | 44018524 | genebody | 11.6092 | GSG1L |
| 867 | scaffold_4 | 44018664 | genebody | 5.7112 | GSG1L |
| 868 | scaffold_4 | 44019014 | genebody | -16.5157 | GSG1L |
| 869 | scaffold_4 | 44023716 | genebody | 8.6546 | GSG1L |
| 870 | scaffold_4 | 44026377 | genebody | 8.6957 | GSG1L |
| 871 | scaffold_4 | 44026732 | genebody | 11.1111 | GSG1L |
| 872 | scaffold_4 | 44027666 | genebody | 7.8322 | GSG1L |
| 873 | scaffold_4 | 44029760 | genebody | -10.2883 | GSG1L |
| 874 | scaffold_4 | 44029771 | genebody | -5.0311 | GSG1L |
| 875 | scaffold_4 | 44029780 | genebody | -11.5885 | GSG1L |
| 876 | scaffold_4 | 44029905 | genebody | -7.7659 | GSG1L |
| 877 | scaffold_4 | 44029966 | genebody | -6.0759 | GSG1L |
| 878 | scaffold_4 | 44034087 | genebody | -5.8714 | GSG1L |
| 879 | scaffold_4 | 44034417 | genebody | 10.0000 | GSG1L |
| 880 | scaffold_4 | 44036211 | genebody | 7.1429 | GSG1L |
| 881 | scaffold_4 | 44036221 | genebody | -7.1429 | GSG1L |
| 882 | scaffold_4 | 44036639 | genebody | -6.3383 | GSG1L |
| 883 | scaffold_4 | 44058166 | genebody | 5.7692 | GSG1L |
| 884 | scaffold_4 | 44058211 | genebody | -15.7895 | GSG1L |
| 885 | scaffold_4 | 44058545 | genebody | -21.4286 | GSG1L |
| 886 | scaffold_4 | 44059154 | genebody | 9.7877 | GSG1L |
| 887 | scaffold_4 | 44059173 | genebody | 5.2843 | GSG1L |
| 888 | scaffold_4 | 44059201 | genebody | -8.7275 | GSG1L |
| 889 | scaffold_4 | 44059250 | genebody | -5.3108 | GSG1L |
| 890 | scaffold_4 | 44059295 | genebody | 6.2500 | GSG1L |
| 891 | scaffold_4 | 44059341 | genebody | 6.6667 | GSG1L |
| 892 | scaffold_4 | 44059694 | genebody | -6.4780 | GSG1L |
| 893 | scaffold_4 | 44059790 | genebody | -46.0606 | GSG1L |
| 894 | scaffold_4 | 44060172 | genebody | 6.4815 | GSG1L |
| 895 | scaffold_4 | 44060216 | genebody | 9.2593 | GSG1L |
| 896 | scaffold_4 | 44061392 | genebody | -15.0000 | GSG1L |
| 897 | scaffold_4 | 44061442 | genebody | -19.0909 | GSG1L |
| 898 | scaffold_4 | 44061694 | genebody | 9.2105 | GSG1L |

| | | | | | |
|---|---|---|---|---|---|
| 899 | scaffold_4 | 44061695 | genebody | -20.0000 | GSG1L |
| 900 | scaffold_4 | 44061854 | genebody | 34.0659 | GSG1L |
| 901 | scaffold_4 | 44066139 | genebody | 11.3636 | GSG1L |
| 902 | scaffold_4 | 44066191 | genebody | 9.0909 | GSG1L |
| 903 | scaffold_4 | 44066550 | genebody | 7.2398 | GSG1L |
| 904 | scaffold_4 | 44071986 | genebody | 6.8027 | GSG1L |
| 905 | scaffold_4 | 44077405 | genebody | 6.9444 | GSG1L |
| 906 | scaffold_4 | 44082500 | genebody | -9.5238 | GSG1L |
| 907 | scaffold_4 | 44082515 | genebody | -9.5238 | GSG1L |
| 908 | scaffold_4 | 44082535 | genebody | 13.4199 | GSG1L |
| 909 | scaffold_4 | 44083839 | genebody | 9.1749 | GSG1L |
| 910 | scaffold_4 | 44083872 | genebody | 6.5900 | GSG1L |
| 911 | scaffold_4 | 44084051 | genebody | 7.1192 | GSG1L |
| 912 | scaffold_4 | 44090763 | genebody | 7.7139 | GSG1L |
| 913 | scaffold_4 | 44091163 | genebody | -18.1818 | GSG1L |
| 914 | scaffold_4 | 44091209 | genebody | -15.1601 | GSG1L |
| 915 | scaffold_4 | 44096860 | genebody | 29.6053 | GSG1L |
| 916 | scaffold_4 | 44097076 | genebody | -8.2280 | GSG1L |
| 917 | scaffold_4 | 44098145 | genebody | -21.0526 | GSG1L |
| 918 | scaffold_4 | 44098553 | genebody | 12.8676 | GSG1L |
| 919 | scaffold_4 | 44100965 | genebody | 6.5583 | GSG1L |
| 920 | scaffold_4 | 44100991 | genebody | 9.4711 | GSG1L |
| 921 | scaffold_4 | 44100992 | genebody | 5.1810 | GSG1L |
| 922 | scaffold_4 | 44100993 | genebody | 5.3829 | GSG1L |
| 923 | scaffold_4 | 44101031 | genebody | 7.4423 | GSG1L |
| 924 | scaffold_4 | 44101054 | genebody | 10.2752 | GSG1L |
| 925 | scaffold_4 | 44101070 | genebody | 5.6899 | GSG1L |
| 926 | scaffold_4 | 44101126 | genebody | 5.9373 | GSG1L |
| 927 | scaffold_4 | 44103201 | genebody | 13.6667 | GSG1L |
| 928 | scaffold_4 | 44103350 | genebody | 6.0141 | GSG1L |
| 929 | scaffold_4 | 44103416 | genebody | 8.0207 | GSG1L |
| 930 | scaffold_4 | 44103777 | genebody | -14.2857 | GSG1L |
| 931 | scaffold_4 | 44104083 | genebody | -11.4286 | GSG1L |
| 932 | scaffold_4 | 44104124 | genebody | -5.7143 | GSG1L |
| 933 | scaffold_4 | 44104130 | genebody | 18.0952 | GSG1L |
| 934 | scaffold_4 | 44104139 | genebody | 12.3810 | GSG1L |
| 935 | scaffold_4 | 44104915 | genebody | -7.2088 | GSG1L |
| 936 | scaffold_4 | 44105493 | genebody | -8.5788 | GSG1L |
| 937 | scaffold_4 | 44105530 | genebody | -7.0423 | GSG1L |
| 938 | scaffold_4 | 44116274 | genebody | -12.0496 | GSG1L |
| 939 | scaffold_4 | 44116433 | genebody | -9.6555 | GSG1L |
| 940 | scaffold_4 | 44130065 | genebody | -6.0504 | GSG1L |
| 941 | scaffold_4 | 44130073 | genebody | -5.0420 | GSG1L |
| 942 | scaffold_4 | 44132727 | genebody | -6.6667 | GSG1L |
| 943 | scaffold_4 | 44135282 | genebody | -5.5556 | GSG1L |

| | | | | | |
|---|---|---|---|---|---|
| 944 | scaffold_4 | 44136160 | genebody | 5.6250 | GSG1L |
| 945 | scaffold_4 | 44136719 | genebody | -5.0794 | GSG1L |
| 946 | scaffold_4 | 44139385 | genebody | -10.3532 | GSG1L |
| 947 | scaffold_4 | 44139447 | genebody | 5.1531 | GSG1L |
| 948 | scaffold_4 | 44139455 | genebody | 5.9804 | GSG1L |
| 949 | scaffold_4 | 44139531 | genebody | -12.3107 | GSG1L |
| 950 | scaffold_4 | 44139743 | genebody | 9.0332 | GSG1L |
| 951 | scaffold_4 | 44150629 | genebody | 17.6136 | GSG1L |
| 952 | scaffold_4 | 44150679 | genebody | 12.8852 | GSG1L |
| 953 | scaffold_4 | 44151066 | genebody | -24.7024 | GSG1L |
| 954 | scaffold_4 | 44152524 | genebody | -9.4575 | GSG1L |
| 955 | scaffold_4 | 44152570 | genebody | -18.5484 | GSG1L |
| 956 | scaffold_4 | 44160625 | genebody | -8.2921 | GSG1L |
| 957 | scaffold_4 | 44160648 | genebody | -6.6630 | GSG1L |
| 958 | scaffold_4 | 44160686 | genebody | -7.3364 | GSG1L |
| 959 | scaffold_4 | 44162442 | genebody | -7.4617 | GSG1L |
| 960 | scaffold_4 | 44162577 | genebody | -7.2654 | GSG1L |
| 961 | scaffold_4 | 44169360 | genebody | 5.0000 | GSG1L |
| 962 | scaffold_4 | 44170676 | genebody | 7.4220 | GSG1L |
| 963 | scaffold_4 | 44170760 | genebody | 5.7816 | GSG1L |
| 964 | scaffold_4 | 44170766 | genebody | 7.2749 | GSG1L |
| 965 | scaffold_4 | 44174203 | genebody | -6.8783 | GSG1L |
| 966 | scaffold_4 | 44174763 | genebody | -5.1220 | GSG1L |
| 967 | scaffold_4 | 44174813 | genebody | -11.0127 | GSG1L |
| 968 | scaffold_4 | 44174934 | genebody | -6.5217 | GSG1L |
| 969 | scaffold_4 | 44174996 | genebody | 8.7506 | GSG1L |
| 970 | scaffold_4 | 44175094 | genebody | 7.3191 | GSG1L |
| 971 | scaffold_4 | 44175133 | genebody | 9.8485 | GSG1L |
| 972 | scaffold_4 | 44175336 | genebody | -10.6370 | GSG1L |
| 973 | scaffold_4 | 44175383 | genebody | -7.7522 | GSG1L |
| 974 | scaffold_4 | 44175384 | genebody | -11.2886 | GSG1L |
| 975 | scaffold_4 | 44175540 | genebody | 13.8984 | GSG1L |
| 976 | scaffold_4 | 44175583 | genebody | 7.3110 | GSG1L |
| 977 | scaffold_4 | 44175593 | genebody | 5.0699 | GSG1L |
| 978 | scaffold_4 | 44180856 | genebody | -13.3838 | GSG1L |
| 979 | scaffold_4 | 44180894 | genebody | -16.6667 | GSG1L |
| 980 | scaffold_4 | 44180897 | genebody | -17.1182 | GSG1L |
| 981 | scaffold_4 | 44180898 | genebody | -8.3333 | GSG1L |
| 982 | scaffold_4 | 44185490 | genebody | -13.7427 | GSG1L |
| 983 | scaffold_4 | 44189676 | genebody | -8.5885 | GSG1L |
| 984 | scaffold_4 | 44189691 | genebody | 7.1007 | GSG1L |
| 985 | scaffold_4 | 44189880 | genebody | 5.2632 | GSG1L |
| 986 | scaffold_4 | 44189881 | genebody | -13.4992 | GSG1L |
| 987 | scaffold_4 | 44190206 | genebody | -5.2632 | GSG1L |
| 988 | scaffold_4 | 44190228 | genebody | 5.4762 | GSG1L |

| | | | | | |
|---|---|---|---|---|---|
| 989 | scaffold_4 | 44190252 | genebody | -14.0476 | GSG1L |
| 990 | scaffold_4 | 44202602 | genebody | -5.2733 | GSG1L |
| 991 | scaffold_4 | 44202637 | genebody | -16.9848 | GSG1L |
| 992 | scaffold_4 | 44204702 | genebody | -11.1111 | GSG1L |
| 993 | scaffold_4 | 44213627 | genebody | -11.7216 | GSG1L |
| 994 | scaffold_4 | 44213799 | genebody | -6.4481 | GSG1L |
| 995 | scaffold_4 | 44213812 | genebody | 5.0753 | GSG1L |
| 996 | scaffold_4 | 44213813 | genebody | 5.9834 | GSG1L |
| 997 | scaffold_4 | 44214761 | genebody | 23.5088 | GSG1L |
| 998 | scaffold_4 | 44214769 | genebody | -7.0175 | GSG1L |
| 999 | scaffold_4 | 44214795 | genebody | 10.1754 | GSG1L |
| 1000 | scaffold_4 | 44214803 | genebody | -7.0175 | GSG1L |
| 1001 | scaffold_4 | 44214821 | genebody | 18.2456 | GSG1L |
| 1002 | scaffold_4 | 44215103 | genebody | 20.7071 | GSG1L |
| 1003 | scaffold_4 | 44215155 | genebody | 12.1175 | GSG1L |
| 1004 | scaffold_4 | 44215156 | genebody | -19.1919 | GSG1L |
| 1005 | scaffold_4 | 44215368 | genebody | -7.1365 | GSG1L |
| 1006 | scaffold_4 | 44215392 | genebody | -8.0645 | GSG1L |
| 1007 | scaffold_4 | 44215393 | genebody | -11.1513 | GSG1L |
| 1008 | scaffold_4 | 44215775 | genebody | -10.8815 | GSG1L |
| 1009 | scaffold_4 | 44215816 | genebody | 13.6842 | GSG1L |
| 1010 | scaffold_4 | 44215887 | genebody | -9.6717 | GSG1L |
| 1011 | scaffold_4 | 44215963 | genebody | 7.5491 | GSG1L |
| 1012 | scaffold_4 | 44216751 | genebody | -7.1290 | GSG1L |
| 1013 | scaffold_4 | 44216767 | genebody | -9.0374 | GSG1L |
| 1014 | scaffold_4 | 44216930 | genebody | -5.6818 | GSG1L |
| 1015 | scaffold_4 | 44216936 | genebody | -27.2727 | GSG1L |
| 1016 | scaffold_4 | 44216988 | genebody | -11.7647 | GSG1L |
| 1017 | scaffold_4 | 44217296 | genebody | 20.0000 | GSG1L |
| 1018 | scaffold_4 | 44217299 | genebody | -10.0000 | GSG1L |
| 1019 | scaffold_4 | 44217321 | genebody | -6.5015 | GSG1L |
| 1020 | scaffold_4 | 44217363 | genebody | -8.3591 | GSG1L |
| 1021 | scaffold_4 | 44218380 | genebody | -26.3636 | GSG1L |
| 1022 | scaffold_4 | 44218440 | genebody | -25.0000 | GSG1L |
| 1023 | scaffold_4 | 44222486 | genebody | -7.7447 | GSG1L |
| 1024 | scaffold_4 | 44222525 | genebody | -9.2949 | GSG1L |
| 1025 | scaffold_4 | 44223679 | genebody | 10.1250 | GSG1L |
| 1026 | scaffold_4 | 44224744 | genebody | -9.0909 | GSG1L |
| 1027 | scaffold_4 | 44225209 | genebody | 8.3333 | GSG1L |
| 1028 | scaffold_4 | 44227805 | genebody | -5.5556 | GSG1L |
| 1029 | scaffold_4 | 44232516 | genebody | 9.6528 | GSG1L |
| 1030 | scaffold_4 | 44232563 | genebody | 15.4831 | GSG1L |
| 1031 | scaffold_4 | 44232564 | genebody | 5.1973 | GSG1L |
| 1032 | scaffold_4 | 44232584 | genebody | -12.4903 | GSG1L |
| 1033 | scaffold_4 | 44232597 | genebody | -6.1905 | GSG1L |

| | | | | | |
|---|---|---|---|---|---|
| 1034 | scaffold_4 | 44232654 | genebody | -6.0968 | GSG1L |
| 1035 | scaffold_4 | 44232731 | genebody | 6.1898 | GSG1L |
| 1036 | scaffold_4 | 44232733 | genebody | 11.8214 | GSG1L |
| 1037 | scaffold_4 | 44232749 | genebody | 18.9878 | GSG1L |
| 1038 | scaffold_4 | 44232790 | genebody | -6.2609 | GSG1L |
| 1039 | scaffold_4 | 44232840 | genebody | -8.9769 | GSG1L |
| 1040 | scaffold_4 | 44234542 | genebody | 16.5074 | GSG1L |
| 1041 | scaffold_4 | 44234611 | genebody | -6.3542 | GSG1L |
| 1042 | scaffold_4 | 44234613 | genebody | -10.2222 | GSG1L |
| 1043 | scaffold_4 | 44234725 | genebody | 7.4297 | GSG1L |
| 1044 | scaffold_4 | 44234761 | genebody | 8.1453 | GSG1L |
| 1045 | scaffold_4 | 44236368 | promoter | 11.1111 | GSG1L |
| 1046 | scaffold_4 | 44236410 | promoter | 5.5556 | GSG1L |
| 1047 | scaffold_4 | 44236749 | promoter | -10.9524 | GSG1L |
| 1048 | scaffold_4 | 44236750 | promoter | 12.6344 | GSG1L |
| 1049 | scaffold_4 | 44236942 | promoter | 8.9852 | GSG1L |
| 1050 | scaffold_4 | 44236949 | intergenic | 35.2632 | |
| 1051 | scaffold_4 | 44236982 | intergenic | 11.9796 | |
| 1052 | scaffold_4 | 44236993 | intergenic | 27.4283 | |
| 1053 | scaffold_4 | 44243687 | intergenic | -6.5752 | |
| 1054 | scaffold_4 | 44243908 | intergenic | 6.2849 | |
| 1055 | scaffold_4 | 44243945 | intergenic | 7.2972 | |
| 1056 | scaffold_4 | 44243946 | intergenic | 8.5255 | |
| 1057 | scaffold_4 | 44257283 | intergenic | 6.2500 | |
| 1058 | scaffold_4 | 44259240 | intergenic | 5.0526 | |
| 1059 | scaffold_4 | 44287415 | genebody;ge | -5.1028 | XPO6 |
| 1060 | scaffold_4 | 44287555 | genebody;ge | -7.1028 | XPO6 |
| 1061 | scaffold_4 | 44294377 | genebody;ge | -5.3448 | XPO6 |
| 1062 | scaffold_4 | 44294735 | genebody;ge | 12.8571 | XPO6 |
| 1063 | scaffold_4 | 44294737 | genebody;ge | 6.6667 | XPO6 |
| 1064 | scaffold_4 | 44294777 | genebody;ge | 6.6667 | XPO6 |
| 1065 | scaffold_4 | 44373678 | genebody;ge | -7.4286 | XPO6 |
| 1066 | scaffold_4 | 44373687 | genebody;ge | -9.5238 | XPO6 |
| 1067 | scaffold_4 | 44392987 | genebody;ge | -20.0000 | XPO6 |
| 1068 | scaffold_4 | 44393025 | genebody;ge | -5.0000 | XPO6 |
| 1069 | scaffold_4 | 44393466 | genebody;ge | -20.1754 | XPO6 |
| 1070 | scaffold_4 | 44404958 | intergenic | -14.9329 | |
| 1071 | scaffold_4 | 44413373 | intergenic | 12.5167 | |
| 1072 | scaffold_4 | 44413374 | intergenic | 21.4286 | |
| 1073 | scaffold_4 | 44413499 | intergenic | -6.2443 | |
| 1074 | scaffold_4 | 44420064 | intergenic | -33.3333 | |
| 1075 | scaffold_4 | 44442604 | promoter | -10.9176 | SBK1 |
| 1076 | scaffold_4 | 44442637 | promoter | -5.4174 | SBK1 |
| 1077 | scaffold_4 | 44454545 | genebody | 8.8746 | SBK1 |
| 1078 | scaffold_4 | 44454546 | genebody | 9.4329 | SBK1 |

| | | | | | |
|---|---|---|---|---|---|
| 1079 | scaffold_4 | 44454586 | genebody | 9.8767 | SBK1 |
| 1080 | scaffold_4 | 44458007 | genebody | 12.5850 | SBK1 |
| 1081 | scaffold_4 | 44458255 | genebody | -8.1633 | SBK1 |
| 1082 | scaffold_4 | 44460743 | genebody | -8.5857 | SBK1 |
| 1083 | scaffold_4 | 44467038 | promoter | 7.1429 | SBK1 |
| 1084 | scaffold_4 | 44467392 | promoter | 6.6667 | SBK1 |
| 1085 | scaffold_4 | 44468237 | promoter | 6.1224 | SBK1 |
| 1086 | scaffold_4 | 44468547 | promoter | 8.3333 | SBK1 |
| 1087 | scaffold_4 | 44488051 | genebody;ge | -10.2564 | SBK1 |
| 1088 | scaffold_4 | 44488146 | genebody;ge | -21.0526 | SBK1 |
| 1089 | scaffold_4 | 44488155 | genebody;ge | -5.2632 | SBK1 |
| 1090 | scaffold_4 | 44488418 | genebody;ge | -6.2612 | SBK1 |
| 1091 | scaffold_4 | 44488438 | genebody;ge | -6.6935 | SBK1 |
| 1092 | scaffold_4 | 44488455 | genebody;ge | -8.8095 | SBK1 |
| 1093 | scaffold_4 | 44488461 | genebody;ge | -5.5331 | SBK1 |
| 1094 | scaffold_4 | 44488474 | genebody;ge | -9.3277 | SBK1 |
| 1095 | scaffold_4 | 44488483 | genebody;ge | -7.1429 | SBK1 |
| 1096 | scaffold_4 | 44488539 | genebody;ge | -6.8966 | SBK1 |
| 1097 | scaffold_4 | 44488547 | genebody;ge | -6.8966 | SBK1 |
| 1098 | scaffold_4 | 44488558 | genebody;ge | -7.0115 | SBK1 |
| 1099 | scaffold_4 | 44488571 | genebody;ge | 5.0000 | SBK1 |
| 1100 | scaffold_4 | 44488576 | genebody;ge | 19.4737 | SBK1 |
| 1101 | scaffold_4 | 44488578 | genebody;ge | -5.2632 | SBK1 |
| 1102 | scaffold_4 | 44488592 | genebody;ge | -16.5528 | SBK1 |
| 1103 | scaffold_4 | 44488610 | genebody;ge | -12.5000 | SBK1 |
| 1104 | scaffold_4 | 44488639 | genebody;ge | -6.6138 | SBK1 |
| 1105 | scaffold_4 | 44488640 | genebody;ge | -14.7431 | SBK1 |
| 1106 | scaffold_4 | 44488643 | genebody;ge | -31.1576 | SBK1 |
| 1107 | scaffold_4 | 44488650 | genebody;ge | 6.8583 | SBK1 |
| 1108 | scaffold_4 | 44488665 | genebody;ge | -10.0065 | SBK1 |
| 1109 | scaffold_4 | 44488674 | genebody;ge | -10.5806 | SBK1 |
| 1110 | scaffold_4 | 44488684 | genebody;ge | -21.7936 | SBK1 |
| 1111 | scaffold_4 | 44488689 | genebody;ge | -7.4883 | SBK1 |
| 1112 | scaffold_4 | 44488704 | genebody;ge | -8.0543 | SBK1 |
| 1113 | scaffold_4 | 44488709 | genebody;ge | -10.3167 | SBK1 |
| 1114 | scaffold_4 | 44488813 | genebody;ge | 6.6038 | SBK1 |
| 1115 | scaffold_4 | 44488847 | genebody;ge | 6.8403 | SBK1 |
| 1116 | scaffold_4 | 44488871 | genebody;ge | -5.5417 | SBK1 |
| 1117 | scaffold_4 | 44488878 | genebody;ge | 5.0735 | SBK1 |
| 1118 | scaffold_4 | 44488891 | genebody;ge | 16.6667 | SBK1 |
| 1119 | scaffold_4 | 44488892 | genebody;ge | 7.0075 | SBK1 |
| 1120 | scaffold_4 | 44488903 | genebody;ge | -8.3333 | SBK1 |
| 1121 | scaffold_4 | 44488920 | genebody;ge | 6.0012 | SBK1 |
| 1122 | scaffold_4 | 44488971 | genebody;ge | -6.4124 | SBK1 |
| 1123 | scaffold_4 | 44542188 | intergenic | 11.6650 | |

| | | | | | |
|---|---|---|---|---|---|
| 1124 | scaffold_4 | 44542203 | intergenic | 14.4853 | |
| 1125 | scaffold_4 | 44543031 | intergenic | 15.9091 | |
| 1126 | scaffold_4 | 44544803 | intergenic | -36.8421 | |
| 1127 | scaffold_4 | 44544863 | intergenic | -7.7193 | |
| 1128 | scaffold_4 | 44545239 | intergenic | -5.8824 | |
| 1129 | scaffold_4 | 44545255 | intergenic | 7.6923 | |
| 1130 | scaffold_4 | 44552082 | intergenic | -12.3077 | |
| 1131 | scaffold_4 | 44552487 | intergenic | 19.2308 | |
| 1132 | scaffold_4 | 44552902 | intergenic | -20.3202 | |
| 1133 | scaffold_4 | 44552903 | intergenic | 17.4603 | |
| 1134 | scaffold_4 | 44553255 | intergenic | -11.9617 | |
| 1135 | scaffold_4 | 44555520 | intergenic | 5.5944 | |
| 1136 | scaffold_4 | 44555526 | intergenic | 10.4895 | |
| 1137 | scaffold_4 | 44555528 | intergenic | 5.7829 | |
| 1138 | scaffold_4 | 44555529 | intergenic | -6.2937 | |
| 1139 | scaffold_4 | 44561874 | intergenic | -5.5390 | |
| 1140 | scaffold_4 | 44562213 | intergenic | 5.6222 | |
| 1141 | scaffold_4 | 44566107 | intergenic | 9.7403 | |
| 1142 | scaffold_4 | 44566137 | intergenic | -19.0476 | |
| 1143 | scaffold_4 | 44566480 | intergenic | -7.4163 | |
| 1144 | scaffold_4 | 44566983 | intergenic | 14.9123 | |
| 1145 | scaffold_4 | 44567091 | intergenic | -9.0395 | |
| 1146 | scaffold_4 | 44567125 | intergenic | -5.0847 | |
| 1147 | scaffold_4 | 44567194 | intergenic | -12.0000 | |
| 1148 | scaffold_4 | 44567419 | intergenic | -6.0150 | |
| 1149 | scaffold_4 | 44567436 | intergenic | -18.4211 | |
| 1150 | scaffold_4 | 44567459 | intergenic | -41.3534 | |
| 1151 | scaffold_4 | 44567467 | intergenic | 13.1579 | |
| 1152 | scaffold_4 | 44567473 | intergenic | 18.4211 | |
| 1153 | scaffold_4 | 44568513 | intergenic | -13.4424 | |
| 1154 | scaffold_4 | 44568665 | intergenic | -7.7551 | |
| 1155 | scaffold_4 | 44568687 | intergenic | -11.9109 | |
| 1156 | scaffold_4 | 44571722 | intergenic | 9.7475 | |
| 1157 | scaffold_4 | 44571751 | intergenic | 5.5656 | |
| 1158 | scaffold_4 | 44571943 | intergenic | -6.9462 | |
| 1159 | scaffold_4 | 44573285 | intergenic | -12.8750 | |
| 1160 | scaffold_4 | 44573656 | intergenic | 10.9524 | |
| 1161 | scaffold_4 | 44575512 | intergenic | -8.7963 | |
| 1162 | scaffold_4 | 44575515 | intergenic | -5.4111 | |
| 1163 | scaffold_4 | 44578994 | intergenic | -13.0390 | |
| 1164 | scaffold_4 | 44579109 | intergenic | 10.5000 | |
| 1165 | scaffold_4 | 44579904 | intergenic | 9.7580 | |
| 1166 | scaffold_4 | 44586000 | genebody | -5.4148 | LAT |
| 1167 | scaffold_4 | 44591005 | genebody; pr | -6.2500 | SPNS1; LAT |
| 1168 | scaffold_4 | 44591337 | genebody;ge | 5.1948 | SPNS1 |

| | | | | | |
|---|---|---|---|---|---|
| 1169 | scaffold_4 | 44591340 | genebody;ge | -9.0909 | SPNS1 |
| 1170 | scaffold_4 | 44592889 | genebody;ge | -11.5385 | SPNS1 |
| 1171 | scaffold_406 | 152282 | genebody;ge | 5.1411 | FZR1 |
| 1172 | scaffold_406 | 152317 | genebody;ge | 12.0789 | FZR1 |
| 1173 | scaffold_41 | 2468437 | genebody;ge | 9.6774 | L3MBTL3 |
| 1174 | scaffold_41 | 2468481 | genebody;ge | 8.3333 | L3MBTL3 |
| 1175 | scaffold_41 | 8487166 | genebody | 6.5104 | IPCEF1 |
| 1176 | scaffold_42 | 819459 | promoter | 7.8693 | CARD19 |
| 1177 | scaffold_42 | 6604266 | genebody | 7.7778 | SART1 |
| 1178 | scaffold_42 | 6604880 | genebody | -10.6592 | SART1 |
| 1179 | scaffold_42 | 6604902 | genebody | 7.2464 | SART1 |
| 1180 | scaffold_43 | 5403170 | intergenic | 5.2254 | |
| 1181 | scaffold_43 | 5403171 | intergenic | 6.3312 | |
| 1182 | scaffold_43 | 5403185 | intergenic | 5.0533 | |
| 1183 | scaffold_43 | 5403261 | intergenic | 6.6447 | |
| 1184 | scaffold_44 | 15285533 | intergenic | 6.1224 | |
| 1185 | scaffold_44 | 15285570 | intergenic | 21.6667 | |
| 1186 | scaffold_45 | 14356581 | genebody | 10.6306 | PREX1 |
| 1187 | scaffold_45 | 14356647 | genebody | 5.1587 | PREX1 |
| 1188 | scaffold_45 | 15510178 | intergenic | -5.0792 | |
| 1189 | scaffold_45 | 15510267 | intergenic | 8.2431 | |
| 1190 | scaffold_45 | 15510269 | intergenic | 6.4685 | |
| 1191 | scaffold_453 | 6114 | intergenic | -5.2929 | |
| 1192 | scaffold_455 | 151392 | intergenic | -7.5440 | |
| 1193 | scaffold_455 | 151448 | intergenic | -12.0996 | |
| 1194 | scaffold_455 | 157587 | intergenic | -9.8229 | |
| 1195 | scaffold_455 | 157613 | intergenic | -5.3140 | |
| 1196 | scaffold_46 | 13233418 | genebody | -7.2927 | VSIG1 |
| 1197 | scaffold_47 | 14121981 | intergenic | 16.3636 | |
| 1198 | scaffold_47 | 14122376 | intergenic | 5.6034 | |
| 1199 | scaffold_48 | 12755250 | genebody | 5.0926 | CCSER1 |
| 1200 | scaffold_48 | 12755285 | genebody | 7.6389 | CCSER1 |
| 1201 | scaffold_48 | 12755296 | genebody | 31.2821 | CCSER1 |
| 1202 | scaffold_48 | 12757823 | genebody | 6.1309 | CCSER1 |
| 1203 | scaffold_48 | 12808939 | intergenic | 15.6522 | |
| 1204 | scaffold_48 | 12882385 | intergenic | -5.9524 | |
| 1205 | scaffold_48 | 12883243 | intergenic | -5.8668 | |
| 1206 | scaffold_48 | 12884372 | intergenic | -7.5979 | |
| 1207 | scaffold_48 | 12884601 | intergenic | 6.5217 | |
| 1208 | scaffold_49 | 1939847 | intergenic | -6.9876 | |
| 1209 | scaffold_49 | 1939983 | intergenic | 8.5979 | |
| 1210 | scaffold_49 | 2009379 | intergenic | -5.4641 | |
| 1211 | scaffold_49 | 2009518 | intergenic | -7.6190 | |
| 1212 | scaffold_49 | 2011633 | intergenic | -5.3059 | |
| 1213 | scaffold_5 | 5566435 | promoter | -7.1778 | |

| | | | | | |
|---|---|---|---|---|---|
| 1214 | scaffold_5 | 14322216 | genebody;ge | -5.4177 | DCUN1D2 |
| 1215 | scaffold_5 | 14323460 | genebody;ge | -7.1611 | DCUN1D2 |
| 1216 | scaffold_5 | 14323476 | genebody;ge | -17.3913 | DCUN1D2 |
| 1217 | scaffold_5 | 14328198 | genebody;ge | -7.1579 | DCUN1D2 |
| 1218 | scaffold_5 | 14328260 | genebody;ge | -25.0000 | DCUN1D2 |
| 1219 | scaffold_5 | 14348803 | genebody;ge | 9.5217 | |
| 1220 | scaffold_5 | 14352160 | genebody;ge | 5.1645 | |
| 1221 | scaffold_5 | 15844790 | genebody | 5.4945 | COL4A2 |
| 1222 | scaffold_5 | 15844801 | genebody | -5.4945 | COL4A2 |
| 1223 | scaffold_5 | 15844948 | genebody | -6.6563 | COL4A2 |
| 1224 | scaffold_5 | 15845263 | genebody | -12.3412 | COL4A2 |
| 1225 | scaffold_5 | 15845327 | genebody | -13.8028 | COL4A2 |
| 1226 | scaffold_5 | 15845441 | genebody | 7.5839 | COL4A2 |
| 1227 | scaffold_5 | 16388073 | intergenic | -6.5800 | |
| 1228 | scaffold_5 | 16388591 | intergenic | -10.8333 | |
| 1229 | scaffold_5 | 16399872 | intergenic | -6.9257 | |
| 1230 | scaffold_5 | 18950056 | intergenic | 8.3333 | |
| 1231 | scaffold_5 | 23258443 | genebody;ge | 20.9514 | NALCN |
| 1232 | scaffold_5 | 23258461 | genebody;ge | 21.0387 | NALCN |
| 1233 | scaffold_5 | 23258467 | genebody;ge | 22.3784 | NALCN |
| 1234 | scaffold_5 | 23258486 | genebody;ge | 23.6735 | NALCN |
| 1235 | scaffold_5 | 23258488 | genebody;ge | 18.7755 | NALCN |
| 1236 | scaffold_5 | 53722681 | promoter | -6.2500 | PCDH9 |
| 1237 | scaffold_522 | 129268 | intergenic | 10.2982 | |
| 1238 | scaffold_523 | 67998 | intergenic | -7.9622 | |
| 1239 | scaffold_523 | 68004 | intergenic | 7.6923 | |
| 1240 | scaffold_523 | 68210 | intergenic | 6.1456 | |
| 1241 | scaffold_53 | 3965510 | intergenic | -12.3077 | |
| 1242 | scaffold_53 | 4030844 | intergenic | -10.2041 | |
| 1243 | scaffold_53 | 4070350 | intergenic | 7.2363 | |
| 1244 | scaffold_53 | 4104365 | intergenic | 14.8148 | |
| 1245 | scaffold_53 | 4105401 | intergenic | -7.6522 | |
| 1246 | scaffold_53 | 4106036 | intergenic | -9.9548 | |
| 1247 | scaffold_53 | 4106095 | intergenic | 11.3122 | |
| 1248 | scaffold_53 | 4107592 | intergenic | -8.0128 | |
| 1249 | scaffold_53 | 4107601 | intergenic | -15.3846 | |
| 1250 | scaffold_53 | 4107615 | intergenic | -11.1378 | |
| 1251 | scaffold_53 | 4122680 | intergenic | -5.1471 | |
| 1252 | scaffold_53 | 4122730 | intergenic | 14.3382 | |
| 1253 | scaffold_53 | 4122991 | intergenic | -6.8750 | |
| 1254 | scaffold_53 | 4123029 | intergenic | -7.5000 | |
| 1255 | scaffold_53 | 4134701 | intergenic | -6.1211 | |
| 1256 | scaffold_53 | 4161209 | intergenic | -19.5238 | |
| 1257 | scaffold_53 | 4161272 | intergenic | 13.3333 | |
| 1258 | scaffold_53 | 4161277 | intergenic | 17.8571 | |

| | | | | | |
|---|---|---|---|---|---|
| 1259 | scaffold_53 | 4161517 | intergenic | 16.1376 | |
| 1260 | scaffold_53 | 4214062 | intergenic | -9.5238 | |
| 1261 | scaffold_53 | 4214086 | intergenic | -9.5238 | |
| 1262 | scaffold_53 | 4214524 | intergenic | -10.1810 | |
| 1263 | scaffold_53 | 4268886 | intergenic | 6.1999 | |
| 1264 | scaffold_53 | 4268974 | intergenic | 8.0748 | |
| 1265 | scaffold_53 | 4276554 | intergenic | -12.5000 | |
| 1266 | scaffold_53 | 4276574 | intergenic | -6.2500 | |
| 1267 | scaffold_53 | 4276586 | intergenic | -11.3636 | |
| 1268 | scaffold_53 | 4518702 | intergenic | -6.6667 | |
| 1269 | scaffold_53 | 4518724 | intergenic | -7.4510 | |
| 1270 | scaffold_53 | 4518732 | intergenic | 5.8824 | |
| 1271 | scaffold_53 | 4518762 | intergenic | 11.7647 | |
| 1272 | scaffold_53 | 4661197 | intergenic | -8.3333 | |
| 1273 | scaffold_53 | 4661247 | intergenic | -41.6667 | |
| 1274 | scaffold_53 | 4677287 | intergenic | -6.1410 | |
| 1275 | scaffold_53 | 5044306 | intergenic | -5.8824 | |
| 1276 | scaffold_53 | 5044312 | intergenic | 18.1818 | |
| 1277 | scaffold_53 | 5044314 | intergenic | -23.5294 | |
| 1278 | scaffold_53 | 5103286 | intergenic | 10.9528 | |
| 1279 | scaffold_53 | 7199765 | genebody;ge | 16.0665 | CACNA1G |
| 1280 | scaffold_53 | 7199785 | genebody;ge | 5.5043 | CACNA1G |
| 1281 | scaffold_53 | 7199791 | genebody;ge | 11.6967 | CACNA1G |
| 1282 | scaffold_53 | 7199797 | genebody;ge | 10.2564 | CACNA1G |
| 1283 | scaffold_53 | 7199835 | genebody;ge | 5.1282 | CACNA1G |
| 1284 | scaffold_53 | 8143311 | intergenic | 7.2774 | |
| 1285 | scaffold_54 | 7899360 | intergenic | -5.9915 | |
| 1286 | scaffold_54 | 7899409 | intergenic | 6.9431 | |
| 1287 | scaffold_54 | 7900306 | intergenic | -9.3818 | |
| 1288 | scaffold_55 | 2490114 | intergenic | 9.0226 | |
| 1289 | scaffold_55 | 2490139 | intergenic | -5.2632 | |
| 1290 | scaffold_55 | 4072247 | intergenic | -11.5942 | |
| 1291 | scaffold_55 | 4084534 | intergenic | 8.5859 | |
| 1292 | scaffold_55 | 4084932 | intergenic | -21.4706 | |
| 1293 | scaffold_55 | 6737528 | genebody | 6.3830 | ANKH |
| 1294 | scaffold_55 | 9669565 | intergenic | -9.1166 | |
| 1295 | scaffold_55 | 9669738 | intergenic | -7.9276 | |
| 1296 | scaffold_55 | 9669769 | intergenic | 5.4557 | |
| 1297 | scaffold_56 | 4061441 | genebody | -14.4550 | |
| 1298 | scaffold_56 | 4062552 | genebody | -12.5507 | |
| 1299 | scaffold_56 | 4063692 | genebody | 15.9302 | |
| 1300 | scaffold_6 | 477488 | intergenic | -8.9314 | |
| 1301 | scaffold_6 | 477534 | intergenic | -20.6539 | |
| 1302 | scaffold_6 | 6394780 | intergenic | -5.2083 | |
| 1303 | scaffold_6 | 24965245 | intergenic | -10.6481 | |

| | | | | | |
|---|---|---|---|---|---|
| 1304 | scaffold_6 | 28209159 | intergenic | 15.5012 | |
| 1305 | scaffold_6 | 28209224 | intergenic | 9.2207 | |
| 1306 | scaffold_6 | 28209348 | intergenic | -5.3388 | |
| 1307 | scaffold_6 | 28209352 | intergenic | -13.5411 | |
| 1308 | scaffold_6 | 28471187 | intergenic | -5.6378 | |
| 1309 | scaffold_6 | 28797664 | intergenic | 5.4855 | |
| 1310 | scaffold_6 | 28797734 | intergenic | 15.9146 | |
| 1311 | scaffold_6 | 28797813 | intergenic | -12.1951 | |
| 1312 | scaffold_6 | 28797820 | intergenic | -6.9328 | |
| 1313 | scaffold_6 | 28797838 | intergenic | 9.4080 | |
| 1314 | scaffold_6 | 28797844 | intergenic | -14.6783 | |
| 1315 | scaffold_6 | 28797914 | intergenic | 9.6493 | |
| 1316 | scaffold_6 | 29941669 | genebody | 6.9563 | FLT1 |
| 1317 | scaffold_6 | 30601604 | intergenic | -5.6696 | |
| 1318 | scaffold_6 | 36046018 | intergenic | -27.9713 | |
| 1319 | scaffold_61 | 7388040 | genebody;ge | -6.3889 | ARHGEF15 |
| 1320 | scaffold_61 | 7389429 | genebody;ge | -9.5694 | ARHGEF15 |
| 1321 | scaffold_61 | 7389447 | genebody;ge | 6.4593 | ARHGEF15 |
| 1322 | scaffold_61 | 7389469 | genebody;ge | 16.6667 | ARHGEF15 |
| 1323 | scaffold_61 | 7389733 | genebody;ge | -13.8340 | ARHGEF15 |
| 1324 | scaffold_61 | 7389751 | genebody;ge | 5.2701 | ARHGEF15 |
| 1325 | scaffold_61 | 7389793 | genebody;ge | 16.0037 | ARHGEF15 |
| 1326 | scaffold_61 | 7389809 | genebody;ge | 5.2145 | ARHGEF15 |
| 1327 | scaffold_61 | 7389842 | genebody;ge | 6.4103 | ARHGEF15 |
| 1328 | scaffold_61 | 7390047 | genebody;ge | -20.2589 | ARHGEF15 |
| 1329 | scaffold_61 | 7390058 | genebody;ge | -11.6654 | ARHGEF15 |
| 1330 | scaffold_61 | 9069322 | promoter | -11.7647 | CD68 |
| 1331 | scaffold_61 | 9069358 | promoter | -16.6667 | CD68 |
| 1332 | scaffold_61 | 9069359 | promoter | 7.0707 | CD68 |
| 1333 | scaffold_62 | 2481310 | genebody;ge | 6.4616 | ALOX5 |
| 1334 | scaffold_62 | 5317458 | intergenic | 8.7124 | |
| 1335 | scaffold_62 | 5317642 | intergenic | 6.4626 | |
| 1336 | scaffold_62 | 5317662 | intergenic | 5.2721 | |
| 1337 | scaffold_62 | 5317784 | intergenic | 18.7500 | |
| 1338 | scaffold_62 | 5317810 | intergenic | 15.6250 | |
| 1339 | scaffold_62 | 5318148 | intergenic | 6.3987 | |
| 1340 | scaffold_62 | 5321511 | intergenic | 6.9851 | |
| 1341 | scaffold_62 | 5321641 | intergenic | -11.5801 | |
| 1342 | scaffold_62 | 5321709 | intergenic | 7.2801 | |
| 1343 | scaffold_62 | 5322008 | intergenic | 19.0476 | |
| 1344 | scaffold_62 | 5322066 | intergenic | -8.5714 | |
| 1345 | scaffold_62 | 5322071 | intergenic | -10.0000 | |
| 1346 | scaffold_62 | 5323770 | intergenic | 5.1720 | |
| 1347 | scaffold_62 | 5323857 | intergenic | 5.0965 | |
| 1348 | scaffold_625 | 42130 | intergenic | -5.1012 | |

| | | | | | |
|---|---|---|---|---|---|
| 1349 | scaffold_625 | 42162 | intergenic | 6.3596 | |
| 1350 | scaffold_625 | 42163 | intergenic | -5.9109 | |
| 1351 | scaffold_625 | 42166 | intergenic | -8.6623 | |
| 1352 | scaffold_625 | 42198 | intergenic | -5.2365 | |
| 1353 | scaffold_625 | 42216 | intergenic | -5.7018 | |
| 1354 | scaffold_625 | 42318 | intergenic | -16.0131 | |
| 1355 | scaffold_625 | 42366 | intergenic | -8.0495 | |
| 1356 | scaffold_625 | 42370 | intergenic | 17.6471 | |
| 1357 | scaffold_625 | 51932 | intergenic | 19.7090 | |
| 1358 | scaffold_625 | 51954 | intergenic | 29.1444 | |
| 1359 | scaffold_625 | 52267 | intergenic | -17.1946 | |
| 1360 | scaffold_625 | 52290 | intergenic | 21.9457 | |
| 1361 | scaffold_625 | 52304 | intergenic | 31.9005 | |
| 1362 | scaffold_625 | 52337 | intergenic | -11.3122 | |
| 1363 | scaffold_625 | 52340 | intergenic | 21.8750 | |
| 1364 | scaffold_625 | 52373 | intergenic | 25.0000 | |
| 1365 | scaffold_625 | 52390 | intergenic | 18.7500 | |
| 1366 | scaffold_625 | 52396 | intergenic | 15.6250 | |
| 1367 | scaffold_625 | 52492 | intergenic | -11.6256 | |
| 1368 | scaffold_625 | 52507 | intergenic | 17.0443 | |
| 1369 | scaffold_626 | 26922 | intergenic | 9.6469 | |
| 1370 | scaffold_626 | 26945 | intergenic | 5.2394 | |
| 1371 | scaffold_626 | 26973 | intergenic | -12.0145 | |
| 1372 | scaffold_626 | 26974 | intergenic | 6.6087 | |
| 1373 | scaffold_626 | 26976 | intergenic | -25.4743 | |
| 1374 | scaffold_626 | 26999 | intergenic | -5.4287 | |
| 1375 | scaffold_626 | 27090 | intergenic | 10.3618 | |
| 1376 | scaffold_64 | 309515 | intergenic | 5.9297 | |
| 1377 | scaffold_64 | 1767673 | intergenic | -14.7605 | |
| 1378 | scaffold_64 | 1767906 | intergenic | -16.1429 | |
| 1379 | scaffold_64 | 1767953 | intergenic | -39.8621 | |
| 1380 | scaffold_64 | 5209968 | intergenic | 35.5769 | |
| 1381 | scaffold_64 | 5210053 | intergenic | 5.6859 | |
| 1382 | scaffold_66 | 794434 | genebody;ge | 11.3795 | TNS3 |
| 1383 | scaffold_66 | 794516 | genebody;ge | 5.1337 | TNS3 |
| 1384 | scaffold_66 | 795625 | genebody;ge | 6.6138 | TNS3 |
| 1385 | scaffold_66 | 3533338 | genebody;ge | 6.4935 | TPST1 |
| 1386 | scaffold_67 | 7858629 | genebody | 5.3894 | OTOG |
| 1387 | scaffold_67 | 7858871 | genebody | 31.5364 | OTOG |
| 1388 | scaffold_67 | 7858881 | genebody | 9.0836 | OTOG |
| 1389 | scaffold_67 | 7858889 | genebody | 27.5337 | OTOG |
| 1390 | scaffold_67 | 7858923 | genebody | 14.9326 | OTOG |
| 1391 | scaffold_67 | 7860509 | genebody | 5.4878 | OTOG |
| 1392 | scaffold_67 | 9123772 | genebody;ge | -5.4579 | ADAM11 |
| 1393 | scaffold_67 | 9123814 | genebody;ge | -9.9471 | ADAM11 |

| | | | | | |
|---|---|---|---|---|---|
| 1394 | scaffold_67 | 9123815 | genebody;ge | -13.2363 | ADAM11 |
| 1395 | scaffold_67 | 9123847 | genebody;ge | -12.2166 | ADAM11 |
| 1396 | scaffold_67 | 9123856 | genebody;ge | 5.5026 | ADAM11 |
| 1397 | scaffold_67 | 9123883 | genebody;ge | 9.6908 | ADAM11 |
| 1398 | scaffold_67 | 9123892 | genebody;ge | 10.6391 | ADAM11 |
| 1399 | scaffold_67 | 9123893 | genebody;ge | 5.1841 | ADAM11 |
| 1400 | scaffold_67 | 9123900 | genebody;ge | 5.4643 | ADAM11 |
| 1401 | scaffold_7 | 5272006 | intergenic | -5.7931 | |
| 1402 | scaffold_7 | 5272412 | intergenic | -18.5096 | |
| 1403 | scaffold_7 | 18061269 | intergenic | -8.3039 | |
| 1404 | scaffold_7 | 46246098 | genebody | -6.8182 | TGM4 |
| 1405 | scaffold_7 | 47587053 | genebody;ge | -6.6240 | NBEAL2 |
| 1406 | scaffold_7 | 47957239 | intergenic | 11.4909 | |
| 1407 | scaffold_7 | 47957240 | intergenic | 8.8010 | |
| 1408 | scaffold_7 | 47957276 | intergenic | 8.5288 | |
| 1409 | scaffold_7 | 51192435 | promoter | 6.9749 | ETV5 |
| 1410 | scaffold_7 | 51198834 | intergenic | -5.5743 | |
| 1411 | scaffold_7 | 60559906 | genebody | -20.3529 | IQCG |
| 1412 | scaffold_7 | 60560111 | genebody | -8.0514 | |
| 1413 | scaffold_70 | 1150925 | intergenic | -5.9006 | |
| 1414 | scaffold_70 | 6539690 | genebody | -6.2675 | ETFB |
| 1415 | scaffold_70 | 6539703 | genebody | 5.6991 | |
| 1416 | scaffold_70 | 7338737 | genebody | -9.5640 | MYBPC2 |
| 1417 | scaffold_70 | 7500205 | intergenic | -9.1923 | |
| 1418 | scaffold_70 | 7500245 | intergenic | -8.5026 | |
| 1419 | scaffold_72 | 3962647 | intergenic | -8.5442 | |
| 1420 | scaffold_72 | 3962675 | intergenic | -10.4133 | |
| 1421 | scaffold_72 | 4167957 | genebody | -10.5882 | EXT2 |
| 1422 | scaffold_72 | 4168004 | genebody | -15.7895 | EXT2 |
| 1423 | scaffold_72 | 4168332 | genebody | 8.4084 | EXT2 |
| 1424 | scaffold_72 | 4168343 | genebody | 8.3333 | EXT2 |
| 1425 | scaffold_72 | 4168344 | genebody | -5.4054 | EXT2 |
| 1426 | scaffold_72 | 5300917 | intergenic | -6.9226 | |
| 1427 | scaffold_72 | 5426890 | genebody | 6.9892 | MAPK8IP1 |
| 1428 | scaffold_72 | 5426915 | genebody | 10.4167 | MAPK8IP1 |
| 1429 | scaffold_72 | 5426941 | genebody | -6.9444 | MAPK8IP1 |
| 1430 | scaffold_72 | 8384312 | genebody | 9.9608 | ARMC6 |
| 1431 | scaffold_72 | 8384344 | genebody | 8.6275 | ARMC6 |
| 1432 | scaffold_72 | 8384480 | genebody | -8.9744 | ARMC6 |
| 1433 | scaffold_75 | 5612160 | promoter | 5.0398 | ALLC |
| 1434 | scaffold_75 | 5612170 | promoter | -5.6898 | ALLC |
| 1435 | scaffold_75 | 5612980 | genebody | -5.3143 | ALLC |
| 1436 | scaffold_76 | 273224 | genebody | 11.5000 | OTOP2 |
| 1437 | scaffold_76 | 273296 | genebody | 9.0158 | OTOP2 |
| 1438 | scaffold_76 | 342922 | genebody | 5.6250 | GRIN2C |

| | | | | | |
|---|---|---|---|---|---|
| 1439 | scaffold_76 | 343094 | genebody | -5.2422 | GRIN2C |
| 1440 | scaffold_76 | 2370109 | intergenic | 6.4980 | |
| 1441 | scaffold_76 | 2371269 | intergenic | -7.7778 | |
| 1442 | scaffold_76 | 2371301 | intergenic | -21.2181 | |
| 1443 | scaffold_76 | 2517010 | intergenic | -7.6923 | |
| 1444 | scaffold_76 | 2517037 | intergenic | 16.6667 | |
| 1445 | scaffold_76 | 3034449 | promoter | 32.8571 | |
| 1446 | scaffold_76 | 3614172 | genebody | -7.0234 | DNAH17 |
| 1447 | scaffold_76 | 3614197 | genebody | 7.7124 | DNAH17 |
| 1448 | scaffold_76 | 3614202 | genebody | 5.5719 | DNAH17 |
| 1449 | scaffold_8 | 240495 | genebody | -9.4251 | BPIFB4 |
| 1450 | scaffold_8 | 240663 | genebody | 6.8392 | BPIFB4 |
| 1451 | scaffold_8 | 53942159 | intergenic | 11.5217 | |
| 1452 | scaffold_8 | 53942208 | intergenic | 7.1841 | |
| 1453 | scaffold_8 | 53942209 | intergenic | -8.5771 | |
| 1454 | scaffold_8 | 54004803 | intergenic | -17.3077 | |
| 1455 | scaffold_8 | 54012375 | intergenic | 18.2556 | |
| 1456 | scaffold_80 | 4684680 | genebody | -13.4783 | ETHE1 |
| 1457 | scaffold_80 | 5842181 | genebody;ge | -10.7494 | KLC3;ERCC2 |
| 1458 | scaffold_80 | 5842187 | genebody;ge | -8.6353 | KLC3;ERCC2 |
| 1459 | scaffold_80 | 6670981 | genebody | -11.4679 | GNG8 |
| 1460 | scaffold_80 | 6671077 | genebody | -5.8995 | GNG8 |
| 1461 | scaffold_80 | 7468917 | genebody | 9.9895 | EHD2 |
| 1462 | scaffold_80 | 7468933 | genebody | -11.4286 | EHD2 |
| 1463 | scaffold_81 | 401261 | genebody;ge | -6.0000 | ANXA6 |
| 1464 | scaffold_81 | 401323 | genebody;ge | -5.1050 | ANXA6 |
| 1465 | scaffold_81 | 407584 | genebody;ge | -10.5714 | ANXA6 |
| 1466 | scaffold_81 | 407585 | genebody;ge | -13.4625 | ANXA6 |
| 1467 | scaffold_81 | 407644 | genebody;ge | -16.8571 | ANXA6 |
| 1468 | scaffold_81 | 407962 | genebody;ge | -25.8929 | ANXA6 |
| 1469 | scaffold_81 | 570123 | intergenic | 14.2708 | |
| 1470 | scaffold_88 | 6464009 | genebody;ge | -13.7436 | GLIS1 |
| 1471 | scaffold_88 | 6464021 | genebody;ge | -12.6112 | GLIS1 |
| 1472 | scaffold_88 | 6464068 | genebody;ge | -8.5366 | GLIS1 |
| 1473 | scaffold_9 | 1111936 | genebody | -5.8355 | GNPTAB |
| 1474 | scaffold_9 | 1664796 | intergenic | 9.4566 | |
| 1475 | scaffold_9 | 5412245 | genebody;ge | 9.0909 | CFAP54 |
| 1476 | scaffold_9 | 5412265 | genebody;ge | 36.3636 | CFAP54 |
| 1477 | scaffold_9 | 7558583 | genebody | -6.7364 | CRADD |
| 1478 | scaffold_9 | 7569622 | genebody | -5.6025 | CRADD |
| 1479 | scaffold_9 | 8095936 | intergenic | -12.1191 | |
| 1480 | scaffold_9 | 8095952 | intergenic | -10.9025 | |
| 1481 | scaffold_9 | 8095962 | intergenic | -26.5217 | |
| 1482 | scaffold_9 | 8095963 | intergenic | -8.4927 | |
| 1483 | scaffold_9 | 8095993 | intergenic | 14.0993 | |

| | | | | | |
|---|---|---|---|---|---|
| 1484 | scaffold_9 | 8095997 | intergenic | 9.1232 | |
| 1485 | scaffold_9 | 8096007 | intergenic | 11.5174 | |
| 1486 | scaffold_9 | 40091257 | genebody;ge | -5.2810 | AGAP2 |
| 1487 | scaffold_9 | 40091269 | genebody;ge | -15.7036 | AGAP2 |
| 1488 | scaffold_9 | 42030018 | genebody;ge | -6.5723 | SMARCC2 |
| 1489 | scaffold_9 | 42691280 | intergenic | 5.8824 | |
| 1490 | scaffold_9 | 47500249 | genebody | -5.3108 | |
| 1491 | scaffold_9 | 47500288 | genebody | -5.2735 | |
| 1492 | scaffold_9 | 47500289 | genebody | 5.3221 | |
| 1493 | scaffold_9 | 47511054 | genebody | 25.6039 | |
| 1494 | scaffold_9 | 47511078 | genebody | 9.4203 | |
| 1495 | scaffold_9 | 47511089 | genebody | 6.0606 | |
| 1496 | scaffold_9 | 47512021 | genebody | 15.2681 | |
| 1497 | scaffold_9 | 47529490 | genebody | 5.5556 | |
| 1498 | scaffold_9 | 47529491 | genebody | 10.0794 | |
| 1499 | scaffold_90 | 2643210 | genebody;ge | -8.1681 | RSPH1 |
| 1500 | scaffold_90 | 2643233 | genebody;ge | -7.7731 | RSPH1 |
| 1501 | scaffold_90 | 2643467 | genebody;ge | 12.8571 | RSPH1 |
| 1502 | scaffold_90 | 5437056 | intergenic | 6.3566 | |
| 1503 | scaffold_94 | 5701096 | intergenic | -12.7101 | |
| 1504 | scaffold_94 | 5701288 | intergenic | 7.8466 | |
| 1505 | scaffold_94 | 5701289 | intergenic | 14.5032 | |
| 1506 | scaffold_94 | 5701321 | intergenic | -5.8700 | |
| 1507 | scaffold_96 | 319679 | intergenic | 5.3030 | |
| 1508 | scaffold_96 | 319730 | intergenic | 7.1129 | |
| 1509 | scaffold_96 | 319746 | intergenic | 7.1429 | |
| 1510 | scaffold_96 | 319797 | intergenic | -5.2632 | |
| 1511 | scaffold_96 | 3367134 | genebody | 6.2937 | TSHZ2 |
| 1512 | scaffold_96 | 3367157 | genebody | -13.2867 | TSHZ2 |
| 1513 | scaffold_96 | 3367187 | genebody | 26.5734 | TSHZ2 |
| 1514 | scaffold_96 | 5435792 | intergenic | -5.0009 | |
| 1515 | scaffold_96 | 5435875 | intergenic | 8.4411 | |
| 1516 | scaffold_96 | 5435933 | intergenic | 10.7500 | |
| 1517 | scaffold_96 | 5435934 | intergenic | 9.0671 | |
| 1518 | scaffold_96 | 5578812 | intergenic | -5.3902 | |
| 1519 | scaffold_96 | 5699979 | intergenic | -5.6596 | |
| 1520 | scaffold_96 | 5699996 | intergenic | -8.4071 | |
| 1521 | scaffold_96 | 5699997 | intergenic | 10.3505 | |

PFC following ACS exposure
