## Supplemental Table 3 for "Conserved DNA Methylation Signatures in The Prefrontal Cortex of Newborn and Juvenile Guinea Pigs Following Antenatal Corticosteroid Exposure"

**Supplemental Table 3. Gene Set Enrichement Analysis of DMCs identified in the PND1PFC following ACS exposure**

| source | term_name | term_id | adjusted_p_value | negative_log10_of_adjusted_p_value | term_size | query_size | intersection_size |
|---|---|---|---|---|---|---|---|
| GO:MF | molecular_function | GO:0003674 | 6.81E-09 | 8.166544574 | 15751 | 144 | 122 |
| GO:MF | binding | GO:0005488 | 5.20E-08 | 7.284248653 | 11689 | 144 | 101 |
| GO:MF | protein binding | GO:0005515 | 1.40E-07 | 6.853900709 | 6701 | 144 | 71 |
| GO:MF | alpha-1,3-mannosylglycoprotein 4-beta-N-acetylglucosaminyltransferase activity | GO:0008454 | 0.016601065 | 1.779864059 | 2 | 144 | 2 |
| GO:BP | multicellular organismal process | GO:0032501 | 4.82E-08 | 7.316551481 | 5104 | 144 | 62 |
| GO:BP | anatomical structure development | GO:0048856 | 8.08E-08 | 7.092459574 | 3953 | 144 | 53 |
| GO:BP | developmental process | GO:0032502 | 1.01E-07 | 6.997107189 | 4237 | 144 | 55 |
| GO:BP | multicellular organism development | GO:0007275 | 0.000234888 | 3.629139439 | 3162 | 144 | 40 |
| GO:BP | biological_process | GO:0008150 | 0.000325461 | 3.487500936 | 17138 | 144 | 120 |
| GO:BP | negative regulation of cellular process | GO:0048523 | 0.000508899 | 3.293368005 | 3255 | 144 | 40 |
| GO:BP | anatomical structure morphogenesis | GO:0009653 | 0.001146628 | 2.940577366 | 1874 | 144 | 28 |
| GO:BP | organonitrogen compound metabolic process | GO:1901564 | 0.001654098 | 2.781438758 | 4800 | 144 | 50 |
| GO:BP | biological regulation | GO:0065007 | 0.002173618 | 2.662816866 | 9317 | 144 | 78 |
| GO:BP | regulation of cellular process | GO:0050794 | 0.00289382 | 2.538528484 | 8186 | 144 | 71 |
| GO:BP | animal organ development | GO:0048513 | 0.003312304 | 2.479869779 | 2096 | 144 | 29 |
| GO:BP | negative regulation of biological process | GO:0048519 | 0.003455433 | 2.461497481 | 3917 | 144 | 43 |
| GO:BP | cell differentiation | GO:0030154 | 0.003600391 | 2.443650285 | 2722 | 144 | 34 |
| GO:BP | cellular developmental process | GO:0048869 | 0.00434332 | 2.362178123 | 2745 | 144 | 34 |
| GO:BP | protein metabolic process | GO:0019538 | 0.005974854 | 2.223672698 | 4140 | 144 | 44 |
| GO:BP | regulation of biological process | GO:0050789 | 0.006440266 | 2.191096179 | 9032 | 144 | 75 |
| GO:BP | microtubule-based transport | GO:0099111 | 0.00760247 | 2.119045311 | 122 | 144 | 7 |
| GO:BP | cellular process | GO:0009987 | 0.010518805 | 1.978033614 | 15557 | 144 | 109 |
| GO:BP | establishment of localization | GO:0051234 | 0.01290652 | 1.889190829 | 3421 | 144 | 38 |
| GO:BP | tissue development | GO:0009888 | 0.013903734 | 1.856868563 | 1432 | 144 | 22 |
| GO:BP | regulation of nitrogen compound metabolic process | GO:0051171 | 0.014078777 | 1.851435081 | 3710 | 144 | 40 |
| GO:BP | homeostatic process | GO:0042592 | 0.020038066 | 1.6981442 | 1140 | 144 | 19 |
| GO:BP | plasma membrane bounded cell projection assembly | GO:0120031 | 0.022010772 | 1.657364728 | 406 | 144 | 11 |
| GO:BP | localization | GO:0051179 | 0.022323017 | 1.651247119 | 3923 | 144 | 41 |
| GO:BP | cell projection assembly | GO:0030031 | 0.023038511 | 1.637545594 | 408 | 144 | 11 |
| GO:BP | positive regulation of cellular process | GO:0048522 | 0.026327364 | 1.579592619 | 3808 | 144 | 40 |
| GO:BP | transport | GO:0006810 | 0.03330191 | 1.477530854 | 3288 | 144 | 36 |
| GO:BP | regulation of metabolic process | GO:0019222 | 0.038197674 | 1.417963086 | 4742 | 144 | 46 |
| GO:BP | regulation of cellular metabolic process | GO:0031323 | 0.039859726 | 1.399465693 | 3593 | 144 | 38 |
| GO:BP | positive regulation of metabolic process | GO:0009893 | 0.041435614 | 1.382626226 | 2649 | 144 | 31 |
| GO:BP | cerebrospinal fluid circulation | GO:0090660 | 0.041621452 | 1.380682771 | 11 | 144 | 3 |
| GO:BP | regulation of multicellular organismal process | GO:0051239 | 0.041940821 | 1.377363076 | 2015 | 144 | 26 |
| GO:BP | cell adhesion | GO:0007155 | 0.041960997 | 1.377154203 | 890 | 144 | 16 |
| GO:BP | microtubule-based movement | GO:0007018 | 0.043473819 | 1.361772211 | 286 | 144 | 9 |
| GO:BP | positive regulation of nitrogen compound metabolic process | GO:0051173 | 0.044834598 | 1.348386717 | 2147 | 144 | 27 |
| GO:BP | regulation of molecular function | GO:0065009 | 0.045512623 | 1.341868138 | 1663 | 144 | 23 |
| GO:BP | protein modification process | GO:0036211 | 0.045564339 | 1.341374926 | 2662 | 144 | 31 |
| GO:BP | plasma membrane bounded cell projection organization | GO:0120036 | 0.049836041 | 1.302456462 | 1005 | 144 | 17 |
| GO:CC | cellular anatomical entity | GO:0110165 | 1.72E-07 | 6.763511804 | 16449 | 144 | 122 |
| GO:CC | cellular_component | GO:0005575 | 1.42684E-06 | 5.845623904 | 16851 | 144 | 122 |
| GO:CC | cytoplasm | GO:0005737 | 2.73434E-05 | 4.563146972 | 7477 | 144 | 70 |
| GO:CC | cell periphery | GO:0071944 | 3.29247E-05 | 4.48247871 | 3398 | 144 | 42 |
| GO:CC | membrane | GO:0016020 | 0.000936878 | 3.02831675 | 5903 | 144 | 56 |
| GO:CC | cytoskeleton | GO:0005856 | 0.002941728 | 2.531397531 | 1565 | 144 | 23 |
| GO:CC | cell junction | GO:0030054 | 0.004030502 | 2.394640855 | 1261 | 144 | 20 |
| GO:CC | plasma membrane | GO:0005886 | 0.004676794 | 2.330051801 | 3124 | 144 | 35 |
| GO:CC | microtubule cytoskeleton | GO:0015630 | 0.015700206 | 1.80409464 | 951 | 144 | 16 |
| GO:CC | ciliary neurotrophic factor receptor complex | GO:0070110 | 0.02495892 | 1.602774215 | 3 | 144 | 2 |
| GO:CC | synapse | GO:0045202 | 0.032481168 | 1.488368365 | 803 | 144 | 14 |
| GO:CC | nucleoplasm | GO:0005654 | 0.035396789 | 1.451036134 | 2886 | 144 | 31 |
| HP | Phenotypic abnormality | HP:0000118 | 4.24E-09 | 8.372691003 | 4696 | 144 | 61 |
| HP | HP root | HP:0000001 | 4.36E-09 | 8.36042357 | 4699 | 144 | 61 |
| HP | Mode of inheritance | HP:0000005 | 4.73E-08 | 7.325409516 | 4420 | 144 | 57 |
| HP | Abnormality of the face | HP:0000271 | 5.90E-08 | 7.229188181 | 2822 | 144 | 44 |
| HP | Mendelian inheritance | HP:0034345 | 1.07E-07 | 6.971504895 | 4376 | 144 | 56 |
| HP | Abnormality of the head | HP:0000234 | 3.17E-07 | 6.49885631 | 3091 | 144 | 45 |
| HP | Abnormality of head or neck | HP:0000152 | 4.44E-07 | 6.352269784 | 3123 | 144 | 45 |
| HP | Abnormal eyelid morphology | HP:0000492 | 1.04382E-06 | 5.981373883 | 1255 | 144 | 27 |
| HP | Abnormality of the nose | HP:0000366 | 1.80638E-06 | 5.743189893 | 1676 | 144 | 31 |
| HP | Abnormality of the orbital region | HP:0000315 | 2.79523E-06 | 5.553582659 | 1506 | 144 | 29 |
| HP | Abnormal ear morphology | HP:0031703 | 3.60598E-06 | 5.442977038 | 1523 | 144 | 29 |
| HP | Abnormal nervous system physiology | HP:0012638 | 4.11374E-06 | 5.385763356 | 3473 | 144 | 46 |
| HP | Abnormal ocular adnexa morphology | HP:0030669 | 5.78901E-06 | 5.237395819 | 1456 | 144 | 28 |
| HP | Abnormality of the ocular adnexa | HP:0032039 | 7.91616E-06 | 5.101485323 | 1477 | 144 | 28 |
| HP | Abnormality of the nervous system | HP:0000707 | 9.28591E-06 | 5.032175309 | 3692 | 144 | 47 |
| HP | Neurodevelopmental delay | HP:0012758 | 1.02291E-05 | 4.990163934 | 2240 | 144 | 35 |
| HP | Abnormality of immune system physiology | HP:0010978 | 2.65165E-05 | 4.576483817 | 1562 | 144 | 28 |
| HP | Abnormality of the musculoskeletal system | HP:0033127 | 2.82536E-05 | 4.548925877 | 3560 | 144 | 45 |
| HP | Abnormality of the ear | HP:0000598 | 3.0718E-05 | 4.51260643 | 2224 | 144 | 34 |
| HP | Abnormality of the integument | HP:0001574 | 3.4447E-05 | 4.462848383 | 2466 | 144 | 36 |
| HP | Abnormality of higher mental function | HP:0011446 | 4.20752E-05 | 4.375973562 | 2604 | 144 | 37 |
| HP | Abnormality of the skeletal system | HP:0000924 | 6.72123E-05 | 4.172551404 | 3020 | 144 | 40 |
| HP | Abnormality of the pharynx | HP:0000600 | 8.13213E-05 | 4.089795795 | 229 | 144 | 11 |
| HP | Abnormal facial skeleton morphology | HP:0011821 | 9.17852E-05 | 4.037227431 | 1249 | 144 | 24 |
| HP | Neurodevelopmental abnormality | HP:0012759 | 9.79401E-05 | 4.009039622 | 2690 | 144 | 37 |
| HP | Abnormality of the immune system | HP:0002715 | 0.000102995 | 3.987181821 | 1882 | 144 | 30 |
| HP | Abnormal skull morphology | HP:0000929 | 0.000103513 | 3.985003086 | 2221 | 144 | 33 |
| HP | Intellectual disability | HP:0001249 | 0.000105432 | 3.977026956 | 1884 | 144 | 30 |
| HP | Growth abnormality | HP:0001507 | 0.000110821 | 3.955376322 | 2462 | 144 | 35 |
| HP | Delayed speech and language development | HP:0000750 | 0.000115559 | 3.937196734 | 981 | 144 | 21 |
| HP | Abnormality of brain morphology | HP:0012443 | 0.000118861 | 3.924962164 | 2469 | 144 | 35 |
| HP | Abnormality of the skin | HP:0000951 | 0.000124608 | 3.904453503 | 2010 | 144 | 31 |
| HP | Abnormal cerebral cortex morphology | HP:0002538 | 0.000137544 | 3.861557696 | 730 | 144 | 18 |
| HP | Recurrent infections | HP:0002719 | 0.000157777 | 3.801955028 | 909 | 144 | 20 |
| HP | Abnormal axial skeleton morphology | HP:0009121 | 0.000169616 | 3.770534367 | 2505 | 144 | 35 |
| HP | Abnormality of the eye | HP:0000478 | 0.000171099 | 3.766753438 | 2873 | 144 | 38 |
| HP | Recurrent respiratory infections | HP:0002205 | 0.000173177 | 3.761509315 | 581 | 144 | 16 |
| HP | Abnormal cerebral morphology | HP:0002060 | 0.000176814 | 3.752484349 | 1929 | 144 | 30 |
| HP | Language impairment | HP:0002463 | 0.000179101 | 3.746902506 | 1007 | 144 | 21 |
| HP | Abnormal nasal bridge morphology | HP:0000422 | 0.000179101 | 3.746902506 | 1007 | 144 | 21 |
| HP | Abnormal nasopharynx morphology | HP:0001739 | 0.000184219 | 3.73466547 | 195 | 144 | 10 |
| HP | Abnormality of metabolism/homeostasis | HP:0001939 | 0.000207613 | 3.682744995 | 2406 | 144 | 34 |
| HP | Abnormal lung morphology | HP:0002088 | 0.000221164 | 3.655285939 | 1114 | 144 | 22 |
| HP | Abnormal respiratory system morphology | HP:0012252 | 0.000234782 | 3.62933472 | 1314 | 144 | 24 |
| HP | Abnormal nasal morphology | HP:0005105 | 0.000239418 | 3.620843669 | 1417 | 144 | 25 |
| HP | Abnormality of the respiratory system | HP:0002086 | 0.000244526 | 3.611675127 | 1958 | 144 | 30 |
| HP | Abnormal forebrain morphology | HP:0100547 | 0.000247245 | 3.606872714 | 1959 | 144 | 30 |
| HP | Abnormal eye morphology | HP:0012372 | 0.000284513 | 3.545898623 | 2319 | 144 | 33 |
| HP | Abnormal pharynx morphology | HP:0033151 | 0.000336082 | 3.473554928 | 208 | 144 | 10 |
| HP | Unusual infection | HP:0032101 | 0.000452587 | 3.344297542 | 971 | 144 | 20 |
| HP | Abnormality of the upper respiratory tract | HP:0002087 | 0.000546708 | 3.262244513 | 478 | 144 | 14 |
| HP | Respiratory tract infection | HP:0011947 | 0.000550681 | 3.259099963 | 716 | 144 | 17 |
| HP | Abnormal calvaria morphology | HP:0002683 | 0.000650396 | 3.186821805 | 901 | 144 | 19 |
| HP | Abnormal mandible morphology | HP:0000277 | 0.000730835 | 3.136180822 | 1001 | 144 | 20 |
| HP | Seizure | HP:0001250 | 0.00076936 | 3.113870607 | 1837 | 144 | 28 |
| HP | Morphological central nervous system abnormality | HP:0002011 | 0.000876956 | 3.057022137 | 2682 | 144 | 35 |
| HP | Recurrent upper respiratory tract infections | HP:0002788 | 0.00091756 | 3.037365622 | 178 | 144 | 9 |
| HP | Clinical course | HP:0031797 | 0.000920393 | 3.036026925 | 2813 | 144 | 36 |
| HP | Abnormal inflammatory response | HP:0012647 | 0.000941656 | 3.026107515 | 1114 | 144 | 21 |
| HP | Increased inflammatory response | HP:0012649 | 0.000941656 | 3.026107515 | 1114 | 144 | 21 |
| HP | Abnormal nervous system morphology | HP:0012639 | 0.001037261 | 2.984111765 | 2827 | 144 | 36 |
| HP | Abnormal eye physiology | HP:0012373 | 0.001164109 | 2.9340063 | 2345 | 144 | 32 |
| HP | Abnormality of the genitourinary system | HP:0000119 | 0.001268067 | 2.896857748 | 2476 | 144 | 33 |
| HP | Slanting of the palpebral fissure | HP:0200006 | 0.001332115 | 2.875458417 | 677 | 144 | 16 |
| HP | Abnormality of vision | HP:0000504 | 0.001374109 | 2.861978671 | 1241 | 144 | 22 |
| HP | Clinical modifier | HP:0012823 | 0.001392947 | 2.856065525 | 2862 | 144 | 36 |
| HP | Abnormal jaw morphology | HP:0030791 | 0.001407024 | 2.851698561 | 1044 | 144 | 20 |
| HP | Autosomal recessive inheritance | HP:0000007 | 0.001476553 | 2.830751036 | 2869 | 144 | 36 |
| HP | Abnormality of blood and blood-forming tissues | HP:0001871 | 0.001522022 | 2.817579092 | 1352 | 144 | 23 |
| HP | Abnormality of the genital system | HP:0000078 | 0.001740747 | 2.759264436 | 1577 | 144 | 25 |
| HP | Abnormal oral morphology | HP:0031816 | 0.001848859 | 2.733096138 | 2036 | 144 | 29 |
| HP | Abnormal oral cavity morphology | HP:0000163 | 0.001848859 | 2.733096138 | 2036 | 144 | 29 |
| HP | Abnormal fundus morphology | HP:0001098 | 0.002148133 | 2.667938732 | 1275 | 144 | 22 |
| HP | Abnormal skeletal morphology | HP:0011842 | 0.002242742 | 2.64922063 | 2920 | 144 | 36 |
| HP | Abnormal posterior eye segment morphology | HP:0004329 | 0.002261647 | 2.645575236 | 1279 | 144 | 22 |
| HP | Abnormality of the outer ear | HP:0000356 | 0.003429026 | 2.464829201 | 1208 | 144 | 21 |
| HP | Abnormality of the vertebral column | HP:0000925 | 0.003499501 | 2.455993834 | 1420 | 144 | 23 |
| HP | Abnormal retinal morphology | HP:0000479 | 0.003621192 | 2.441148496 | 915 | 144 | 18 |
| HP | Retinal dystrophy | HP:0000556 | 0.003773993 | 2.423198904 | 337 | 144 | 11 |
| HP | Abnormality of the digestive system | HP:0025031 | 0.003781886 | 2.422291565 | 2728 | 144 | 34 |
| HP | Abnormality of the mouth | HP:0000153 | 0.004000751 | 2.397858495 | 2116 | 144 | 29 |
| HP | Abnormal respiratory system physiology | HP:0002795 | 0.005253329 | 2.279565423 | 1347 | 144 | 22 |
| HP | Micrognathia | HP:0000347 | 0.005826376 | 2.23460146 | 759 | 144 | 16 |
| HP | Behavioral abnormality | HP:0000708 | 0.005901981 | 2.229002172 | 1803 | 144 | 26 |
| HP | Neurological speech impairment | HP:0002167 | 0.005920407 | 2.227648441 | 1576 | 144 | 24 |
| HP | Aplasia/Hypoplasia of the mandible | HP:0009118 | 0.006024398 | 2.220086351 | 761 | 144 | 16 |
| HP | Aplasia/Hypoplasia involving bones of the skull | HP:0009116 | 0.007108183 | 2.148241406 | 771 | 144 | 16 |
| HP | Feeding difficulties | HP:0011968 | 0.007244324 | 2.140002141 | 1061 | 144 | 19 |
| HP | Abnormality of the musculature | HP:0003011 | 0.008389319 | 2.076273301 | 2699 | 144 | 33 |
| HP | Abnormality of prenatal development or birth | HP:0001197 | 0.008619084 | 2.064538905 | 974 | 144 | 18 |
| HP | Abnormality of refraction | HP:0000539 | 0.009143467 | 2.038889115 | 608 | 144 | 14 |
| HP | Recurrent otitis media | HP:0000403 | 0.009590626 | 2.018153056 | 178 | 144 | 8 |
| HP | Polycystic kidney dysplasia | HP:0000113 | 0.011241309 | 1.949183111 | 49 | 144 | 5 |
| HP | Abnormality of the palpebral fissures | HP:0008050 | 0.012416749 | 1.905992113 | 806 | 144 | 16 |
| HP | Abnormality of limbs | HP:0040064 | 0.014687712 | 1.833045859 | 2262 | 144 | 29 |
| HP | Abnormal cerebral subcortex morphology | HP:0010993 | 0.015438088 | 1.811406493 | 1119 | 144 | 19 |
| HP | Global developmental delay | HP:0001263 | 0.017106504 | 1.766838733 | 1795 | 144 | 25 |
| HP | Abnormality of body weight | HP:0004323 | 0.019958904 | 1.699863312 | 1694 | 144 | 24 |
| HP | Hypertelorism | HP:0000316 | 0.020345266 | 1.691536625 | 744 | 144 | 15 |
| HP | Abnormal facial shape | HP:0001999 | 0.021268233 | 1.672268581 | 1041 | 144 | 18 |
| HP | Abnormal cerebellum morphology | HP:0001317 | 0.021887717 | 1.659799535 | 942 | 144 | 17 |
| HP | Aplasia/Hypoplasia of the cerebellum | HP:0007360 | 0.023657632 | 1.626028731 | 489 | 144 | 12 |
| HP | Abnormal hindbrain morphology | HP:0011282 | 0.024070767 | 1.618510071 | 949 | 144 | 17 |
| HP | Abnormal metencephalon morphology | HP:0011283 | 0.024070767 | 1.618510071 | 949 | 144 | 17 |
| HP | Onset | HP:0003674 | 0.026660744 | 1.574127734 | 2590 | 144 | 31 |
| HP | Abnormality of limb bone | HP:0040068 | 0.028404346 | 1.546615203 | 1614 | 144 | 23 |
| HP | Abnormality of limb bone morphology | HP:0002813 | 0.028404346 | 1.546615203 | 1614 | 144 | 23 |
| HP | Growth delay | HP:0001510 | 0.028482829 | 1.545416875 | 1731 | 144 | 24 |
| HP | Abnormality of the hand | HP:0001155 | 0.028886681 | 1.539302359 | 1501 | 144 | 22 |
| HP | Abnormal appendicular skeleton morphology | HP:0011844 | 0.030713358 | 1.512672693 | 1739 | 144 | 24 |
| HP | Abnormal muscle physiology | HP:0011804 | 0.031978929 | 1.495136082 | 2357 | 144 | 29 |
| HP | Aplasia/hypoplasia affecting bones of the axial skeleton | HP:0009122 | 0.033214715 | 1.47866947 | 873 | 144 | 16 |
| HP | Increased head circumference | HP:0040194 | 0.03359562 | 1.47371734 | 425 | 144 | 11 |
| HP | Abnormal digit morphology | HP:0011297 | 0.033713997 | 1.472189762 | 1403 | 144 | 21 |
| HP | Abnormality of skin morphology | HP:0011121 | 0.035153921 | 1.454026225 | 1873 | 144 | 25 |
| HP | Abnormal cortical gyration | HP:0002536 | 0.036883175 | 1.433171702 | 352 | 144 | 10 |
| HP | Abnormality of the forehead | HP:0000290 | 0.036903564 | 1.432931691 | 1085 | 144 | 18 |
| HP | Aplasia/Hypoplasia involving the central nervous system | HP:0002977 | 0.03768887 | 1.423786885 | 1761 | 144 | 24 |
| HP | Abnormal pinna morphology | HP:0000377 | 0.038477354 | 1.414794797 | 1195 | 144 | 19 |
| HP | Abnormal ear physiology | HP:0031704 | 0.039887812 | 1.399159787 | 1533 | 144 | 22 |
| HP | Abnormal homeostasis | HP:0012337 | 0.040285622 | 1.394849921 | 1534 | 144 | 22 |
| HP | Abnormality of the cardiovascular system | HP:0001626 | 0.042456053 | 1.37206038 | 2522 | 144 | 30 |
| HP | Inflammatory abnormality of the skin | HP:0011123 | 0.043368623 | 1.36282437 | 437 | 144 | 11 |
| HP | Abnormal reproductive system morphology | HP:0012243 | 0.047186645 | 1.3261809 | 1323 | 144 | 20 |
| HP | Otitis media | HP:0000388 | 0.049788314 | 1.302872582 | 291 | 144 | 9 |

**intersections**
PRICKLE2,NR2F1,UROS,MCPH1,IL6R,TARS3,TNIK,GRB10,MBP,RBM20,PAX2,CA5B,OFD1,SETD1B,UNC5D,TGFB2,USH2A,ACIN1,ITPKB,TIMP3,SYN3,LARGE1,STOX1,TBC1D22B,NKD2,GATA3,SORCS2,DOK7,AKAP8L,FABP6,RFX4,MGAT4B,RNF208,SHANK3,NECTIN1,FXYD2,RNPC3,GIPC2,TPCN1,ACP5,MYO5A,MGAT4A,FAHD2A,AHDC1,MYOM3,EPHA2,FHAD1,PEX14,ESPN,CHD5,AJAP1,MRPS2,RALGDS,DYNC2I2,NEK6,CNTFR,RECQL,TMPRSS6,EFCAB6,STAB2,GAD1,KLHL23,PPIG,ACMSD,MEPCE,SOST
PRICKLE2,NR2F1,MCPH1,IL6R,TARS3,TNIK,GRB10,MBP,RBM20,PAX2,CA5B,OFD1,SETD1B,UNC5D,TGFB2,USH2A,ACIN1,TIMP3,SYN3,LARGE1,STOX1,TBC1D22B,NKD2,GATA3,SORCS2,DOK7,AKAP8L,FABP6,RFX4,RNF208,SHANK3,NECTIN1,RNPC3,GIPC2,TPCN1,ACP5,MYO5A,AHDC1,MYOM3,EPHA2,FHAD1,PEX14,ESPN,CHD5,AJAP1,DYNC2I2,NEK6,CNTFR,RECQL,TMPRSS6,EFCAB6,STAB2,GAD1,KLHL23,PPIG,ACMSD,MEPCE,SOSTDC1,MNT,FOXN1,EPHA3,ARID3A,TYK2,ARVCF,IL4R,IL21R,XPO6,SE
MCPH1,IL6R,GRB10,MBP,RBM20,OFD1,SETD1B,UNC5D,TGFB2,USH2A,TIMP3,TBC1D22B,NKD2,GATA3,SORCS2,DOK7,AKAP8L,SHANK3,NECTIN1,RNPC3,GIPC2,TPCN1,MYO5A,MYOM3,EPHA2,FHAD1,PEX14,ESPN,CHD5,AJAP1,DYNC2I2,NEK6,CNTFR,TMPRSS6,GAD1,KLHL23,SOSTDC1,MNT,EPHA3,ARID3A,TYK2,ARVCF,IL4R,IL21R,XPO6,LAT,FZR1,L3MBTL3,PREX1,DCUN1D2,CACNA1G,FLT1,ALOX5,TNS3,TPST1,OTOG,NBEAL2,IQCG,MYBPC2,EXT2,MAPK8IP1,ARMC6,DNAH17,ETHE1,KLC3,GNG8,EH
MGAT4B,MGAT4A
MCPH1,IL6R,TNIK,GRB10,MBP,RBM20,PAX2,UNC5D,TGFB2,USH2A,ITPKB,INSYN1,STOX1,GATA3,TRAPPC9,RFX4,SHANK3,NECTIN1,ODAD3,ACP5,LEO1,AHDC1,IGSF21,EPHA2,AJAP1,CNTFR,GAD1,SOSTDC1,FOXN1,EPHA3,TYK2,DTNBP1,MBOAT1,ARVCF,IL4R,KATNIP,LAT,FZR1,L3MBTL3,SART1,PREX1,VSIG1,COL4A2,CACNA1G,ANKH,FLT1,ARHGEF15,ALOX5,OTOG,IQCG,EXT2,MAPK8IP1,GRIN2C,KLC3,ERCC2,GNG8,ANXA6,GLIS1,CFAP54,AGAP2,SMARCC2,RSPH1
MCPH1,IL6R,TNIK,MBP,RBM20,PAX2,UNC5D,TGFB2,USH2A,ITPKB,LARGE1,STOX1,GATA3,TRAPPC9,RFX4,SHANK3,NECTIN1,ODAD3,ACP5,LEO1,AHDC1,IGSF21,EPHA2,AJAP1,CNTFR,SOSTDC1,FOXN1,EPHA3,DTNBP1,MBOAT1,ARVCF,FZR1,L3MBTL3,SART1,PREX1,VSIG1,COL4A2,ANKH,FLT1,ARHGEF15,ALOX5,NBEAL2,IQCG,EXT2,ARMC6,KLC3,ERCC2,GNG8,EHD2,CFAP54,AGAP2,SMARCC2,RSPH1
MCPH1,IL6R,TNIK,MBP,RBM20,PAX2,TMEM120B,UNC5D,TGFB2,USH2A,ITPKB,LARGE1,STOX1,GATA3,TRAPPC9,RFX4,SHANK3,NECTIN1,ODAD3,ACP5,LEO1,AHDC1,IGSF21,EPHA2,AJAP1,CNTFR,SOSTDC1,FOXN1,EPHA3,DTNBP1,MBOAT1,ARVCF,FZR1,L3MBTL3,SART1,PREX1,VSIG1,COL4A2,ANKH,FLT1,ARHGEF15,ALOX5,NBEAL2,IQCG,EXT2,ARMC6,KLC3,ERCC2,GNG8,EHD2,GLIS1,CFAP54,AGAP2,SMARCC2,RSPH1
MCPH1,IL6R,TNIK,MBP,RBM20,PAX2,UNC5D,TGFB2,USH2A,ITPKB,STOX1,GATA3,TRAPPC9,RFX4,SHANK3,NECTIN1,ODAD3,ACP5,LEO1,AHDC1,IGSF21,EPHA2,AJAP1,SOSTDC1,FOXN1,EPHA3,DTNBP1,MBOAT1,ARVCF,FZR1,SART1,COL4A2,ANKH,FLT1,ARHGEF15,ALOX5,EXT2,ERCC2,GNG8,SMARCC2
NR2F1,UROS,MCPH1,IL6R,TARS3,TNIK,GRB10,MBP,RBM20,PAX2,CA5B,SETD1B,TMEM120B,UNC5D,TGFB2,USH2A,ITPKB,TIMP3,SYN3,LARGE1,INSYN1,STOX1,NKD2,GATA3,SORCS2,DOK7,AKAP8L,TRAPPC9,RFX4,MGAT4B,RNF208,SHANK3,NECTIN1,FXYD2,TPCN1,ODAD3,ACP5,LEO1,MYO5A,MGAT4A,AHDC1,IGSF21,EPHA2,PEX14,ESPN,CHD5,AJAP1,MRPS2,RALGDS,DYNC2I2,NEK6,CNTFR,RECQL,TMPRSS6,STAB2,GAD1,PPIG,ACMSD,MEPCE,SOSTDC1,MNT,SLC43A2,FOXN1,EPHA3,ARID3A,TY
NR2F1,MCPH1,GRB10,MBP,RBM20,PAX2,TGFB2,USH2A,ITPKB,TIMP3,NKD2,GATA3,SORCS2,RFX4,SHANK3,ACP5,LEO1,EPHA2,PEX14,AJAP1,CNTFR,TMPRSS6,ACMSD,MEPCE,SOSTDC1,MNT,SLC43A2,EPHA3,DTNBP1,FZR1,L3MBTL3,FLT1,ARHGEF15,ALOX5,ETV5,MAPK8IP1,GLIS1,CRADD,AGAP2,SMARCC2
TNIK,MBP,RBM20,PAX2,UNC5D,TGFB2,STOX1,GATA3,SHANK3,NECTIN1,ACP5,LEO1,AHDC1,EPHA2,AJAP1,SOSTDC1,FOXN1,DTNBP1,VSIG1,COL4A2,ANKH,FLT1,ARHGEF15,ALOX5,NBEAL2,EXT2,ERCC2,EHD2
UROS,IL6R,TARS3,TNIK,MBP,PAX2,SETD1B,TGFB2,ITPKB,TIMP3,LARGE1,STOX1,NKD2,GATA3,DOK7,AKAP8L,RFX4,MGAT4B,RNF208,LEO1,MGAT4A,EPHA2,CHD5,MRPS2,NEK6,TMPRSS6,STAB2,GAD1,PPIG,ACMSD,EPHA3,TYK2,DTNBP1,MBOAT1,SBK1,LAT,FZR1,L3MBTL3,DCUN1D2,FLT1,TPST1,ADAM11,TGM4,ETFB,EXT2,MAPK8IP1,ALLC,GRIN2C,GNPTAB,CRADD
NR2F1,MCPH1,IL6R,TNIK,GRB10,MBP,RBM20,PAX2,UNC5D,TGFB2,USH2A,ITPKB,TIMP3,SYN3,INSYN1,STOX1,NKD2,GATA3,SORCS2,DOK7,AKAP8L,TRAPPC9,RFX4,SHANK3,NECTIN1,FXYD2,TPCN1,ODAD3,ACP5,LEO1,EPHA2,PEX14,CHD5,AJAP1,RALGDS,NEK6,CNTFR,TMPRSS6,GAD1,ACMSD,MEPCE,SOSTDC1,MNT,SLC43A2,FOXN1,EPHA3,ARID3A,TYK2,DTNBP1,MBOAT1,IL4R,GSG1L,LAT,SPNS1,FZR1,L3MBTL3,SART1,PREX1,DCUN1D2,COL4A2,CACNA1G,ANKH,FLT1,ARHGEF15,ALOX5,ETV5,EXT
NR2F1,MCPH1,IL6R,TNIK,GRB10,MBP,RBM20,PAX2,UNC5D,TGFB2,USH2A,ITPKB,TIMP3,INSYN1,STOX1,NKD2,GATA3,SORCS2,DOK7,AKAP8L,TRAPPC9,RFX4,SHANK3,NECTIN1,FXYD2,TPCN1,ODAD3,ACP5,LEO1,EPHA2,PEX14,CHD5,AJAP1,RALGDS,NEK6,CNTFR,TMPRSS6,ACMSD,MEPCE,SOSTDC1,MNT,SLC43A2,FOXN1,EPHA3,ARID3A,TYK2,DTNBP1,MBOAT1,IL4R,GSG1L,LAT,FZR1,L3MBTL3,SART1,PREX1,COL4A2,CACNA1G,FLT1,ARHGEF15,ALOX5,ETV5,MAPK8IP1,GRIN2C,ERCC2,GNG8,EHD2,A
MCPH1,IL6R,RBM20,PAX2,TGFB2,USH2A,LARGE1,STOX1,GATA3,TRAPPC9,RFX4,SHANK3,NECTIN1,ODAD3,ACP5,AHDC1,EPHA2,AJAP1,CNTFR,SOSTDC1,FOXN1,FZR1,ANKH,ARHGEF15,ALOX5,EXT2,ERCC2,GNG8,AGAP2
NR2F1,MCPH1,GRB10,MBP,RBM20,PAX2,TGFB2,USH2A,ITPKB,TIMP3,STOX1,NKD2,GATA3,SORCS2,RFX4,SHANK3,ACP5,LEO1,EPHA2,PEX14,AJAP1,CNTFR,TMPRSS6,ACMSD,MEPCE,SOSTDC1,MNT,SLC43A2,EPHA3,DTNBP1,FZR1,L3MBTL3,COL4A2,FLT1,ARHGEF15,ALOX5,ETV5,MAPK8IP1,GRIN2C,GLIS1,CRADD,AGAP2,SMARCC2
TNIK,TMEM120B,UNC5D,TGFB2,USH2A,ITPKB,GATA3,TRAPPC9,SHANK3,LEO1,EPHA2,AJAP1,SOSTDC1,FOXN1,EPHA3,DTNBP1,MBOAT1,FZR1,L3MBTL3,SART1,PREX1,VSIG1,COL4A2,ALOX5,NBEAL2,IQCG,EXT2,ARMC6,KLC3,ERCC2,EHD2,GLIS1,CFAP54,RSPH1
TNIK,TMEM120B,UNC5D,TGFB2,USH2A,ITPKB,GATA3,TRAPPC9,SHANK3,LEO1,EPHA2,AJAP1,SOSTDC1,FOXN1,EPHA3,DTNBP1,MBOAT1,FZR1,L3MBTL3,SART1,PREX1,VSIG1,COL4A2,ALOX5,NBEAL2,IQCG,EXT2,ARMC6,KLC3,ERCC2,EHD2,GLIS1,CFAP54,RSPH1
IL6R,TARS3,TNIK,MBP,PAX2,SETD1B,TGFB2,ITPKB,TIMP3,LARGE1,STOX1,NKD2,GATA3,DOK7,AKAP8L,RFX4,MGAT4B,RNF208,LEO1,MGAT4A,EPHA2,CHD5,MRPS2,NEK6,TMPRSS6,GAD1,PPIG,EPHA3,TYK2,DTNBP1,SBK1,LAT,FZR1,L3MBTL3,DCUN1D2,FLT1,TPST1,ADAM11,TGM4,EXT2,MAPK8IP1,GRIN2C,GNPTAB,CRADD
NR2F1,MCPH1,IL6R,TNIK,GRB10,MBP,RBM20,PAX2,UNC5D,TGFB2,USH2A,ITPKB,TIMP3,SYN3,INSYN1,STOX1,NKD2,GATA3,SORCS2,DOK7,AKAP8L,TRAPPC9,RFX4,SHANK3,NECTIN1,FXYD2,TPCN1,ODAD3,ACP5,LEO1,EPHA2,PEX14,CHD5,AJAP1,RALGDS,NEK6,CNTFR,TMPRSS6,ACMSD,MEPCE,SOSTDC1,MNT,SLC43A2,FOXN1,EPHA3,ARID3A,TYK2,DTNBP1,MBOAT1,IL4R,GSG1L,LAT,FZR1,L3MBTL3,SART1,PREX1,DCUN1D2,COL4A2,CACNA1G,ANKH,FLT1,ARHGEF15,ALOX5,ETV5,MAPK8IP1,GRIN
ODAD3,PEX14,DYNC2I2,DTNBP1,KATNIP,KLC3,CFAP54
NR2F1,UROS,MCPH1,IL6R,TARS3,TNIK,GRB10,MBP,RBM20,PAX2,SETD1B,TMEM120B,UNC5D,TGFB2,USH2A,ITPKB,TIMP3,SYN3,INSYN1,STOX1,NKD2,GATA3,SORCS2,DOK7,AKAP8L,TRAPPC9,RFX4,SHANK3,NECTIN1,FXYD2,TPCN1,ODAD3,ACP5,LEO1,MYO5A,IGSF21,EPHA2,PEX14,ESPN,CHD5,AJAP1,MRPS2,RALGDS,DYNC2I2,NEK6,CNTFR,RECQL,TMPRSS6,STAB2,GAD1,PPIG,ACMSD,MEPCE,SOSTDC1,MNT,SLC43A2,FOXN1,EPHA3,ARID3A,TYK2,DTNBP1,MBOAT1,ARVCF,IL4R,KATNIP,GSG1L,X
MCPH1,GRB10,TGFB2,USH2A,SYN3,NKD2,GATA3,SORCS2,AKAP8L,SHANK3,NECTIN1,FXYD2,TPCN1,ODAD3,MYO5A,PEX14,DYNC2I2,STAB2,SLC43A2,EPHA3,DTNBP1,KATNIP,GSG1L,XPO6,SPNS1,NALCN,CACNA1G,ANKH,ALOX5,EXT2,MAPK8IP1,OTOP2,GRIN2C,KLC3,EHD2,ANXA6,GNPTAB,CFAP54
IL6R,PAX2,TGFB2,USH2A,STOX1,GATA3,SHANK3,NECTIN1,LEO1,AHDC1,EPHA2,AJAP1,SOSTDC1,FOXN1,FZR1,VSIG1,COL4A2,ANKH,ALOX5,EXT2,ERCC2,AGAP2
NR2F1,MCPH1,IL6R,TNIK,MBP,RBM20,PAX2,TGFB2,ITPKB,TIMP3,STOX1,NKD2,GATA3,DOK7,AKAP8L,TRAPPC9,RFX4,ACP5,LEO1,PEX14,CHD5,TMPRSS6,ACMSD,MEPCE,MNT,FOXN1,ARID3A,DTNBP1,LAT,FZR1,L3MBTL3,DCUN1D2,FLT1,ETV5,MAPK8IP1,GRIN2C,GLIS1,CRADD,AGAP2,SMARCC2
GRB10,MBP,USH2A,ITPKB,LARGE1,GATA3,SHANK3,ACP5,TMPRSS6,FOXN1,LAT,SPNS1,L3MBTL3,VSIG1,ANKH,ALOX5,EXT2,ERCC2,ANXA6
TNIK,RFX4,ODAD3,EPHA2,ESPN,DYNC2I2,IQCG,DNAH17,KLC3,CFAP54,RSPH1
MCPH1,TNIK,GRB10,TGFB2,USH2A,SYN3,NKD2,GATA3,SORCS2,AKAP8L,SHANK3,NECTIN1,FXYD2,TPCN1,ODAD3,MYO5A,EPHA2,PEX14,DYNC2I2,STAB2,MEPCE,SLC43A2,EPHA3,DTNBP1,KATNIP,GSG1L,XPO6,SPNS1,NALCN,CACNA1G,ANKH,ALOX5,EXT2,MAPK8IP1,OTOP2,GRIN2C,KLC3,EHD2,ANXA6,GNPTAB,CFAP54
TNIK,RFX4,ODAD3,EPHA2,ESPN,DYNC2I2,IQCG,DNAH17,KLC3,CFAP54,RSPH1
IL6R,TNIK,GRB10,RBM20,PAX2,TGFB2,ITPKB,TIMP3,STOX1,NKD2,GATA3,DOK7,AKAP8L,RFX4,SHANK3,TPCN1,LEO1,EPHA2,CNTFR,TMPRSS6,MEPCE,FOXN1,EPHA3,ARID3A,TYK2,DTNBP1,LAT,FZR1,SART1,PREX1,FLT1,ARHGEF15,ALOX5,ETV5,MAPK8IP1,EHD2,GLIS1,CRADD,AGAP2,SMARCC2
GRB10,TGFB2,SYN3,NKD2,GATA3,SORCS2,AKAP8L,SHANK3,NECTIN1,FXYD2,TPCN1,ODAD3,MYO5A,PEX14,DYNC2I2,STAB2,SLC43A2,EPHA3,DTNBP1,KATNIP,GSG1L,XPO6,SPNS1,NALCN,CACNA1G,ANKH,ALOX5,EXT2,MAPK8IP1,OTOP2,GRIN2C,KLC3,EHD2,ANXA6,GNPTAB,CFAP54
NR2F1,MCPH1,IL6R,TNIK,GRB10,MBP,RBM20,PAX2,TGFB2,ITPKB,TIMP3,STOX1,NKD2,GATA3,DOK7,AKAP8L,TRAPPC9,RFX4,TPCN1,ACP5,LEO1,EPHA2,PEX14,CHD5,TMPRSS6,ACMSD,MEPCE,MNT,FOXN1,ARID3A,TYK2,DTNBP1,LAT,FZR1,L3MBTL3,DCUN1D2,FLT1,ALOX5,ETV5,MAPK8IP1,GRIN2C,GLIS1,CRADD,AGAP2,SMARCC2,TSHZ2
NR2F1,MCPH1,IL6R,TNIK,GRB10,RBM20,PAX2,TGFB2,ITPKB,TIMP3,STOX1,GATA3,DOK7,AKAP8L,TRAPPC9,RFX4,TPCN1,ACP5,LEO1,PEX14,CHD5,TMPRSS6,ACMSD,MEPCE,MNT,FOXN1,ARID3A,DTNBP1,LAT,FZR1,L3MBTL3,FLT1,ALOX5,ETV5,MAPK8IP1,GLIS1,AGAP2,SMARCC2
IL6R,TNIK,GRB10,MBP,RBM20,PAX2,TGFB2,STOX1,NKD2,GATA3,DOK7,AKAP8L,RFX4,TPCN1,LEO1,TMPRSS6,MEPCE,FOXN1,ARID3A,TYK2,DTNBP1,LAT,FZR1,L3MBTL3,DCUN1D2,FLT1,ETV5,GLIS1,CRADD,AGAP2,SMARCC2
ODAD3,KATNIP,CFAP54
IL6R,GRB10,MBP,TGFB2,ITPKB,STOX1,GATA3,SHANK3,ACP5,LEO1,EPHA2,AJAP1,SOSTDC1,FOXN1,TYK2,LAT,SART1,COL4A2,CACNA1G,ANKH,FLT1,ALOX5,MAPK8IP1,ANXA6,AGAP2,SMARCC2
MBP,UNC5D,TGFB2,ITPKB,GATA3,NECTIN1,EPHA2,AJAP1,STAB2,EPHA3,TYK2,ARVCF,SART1,PREX1,PCDH9,ALOX5
ODAD3,PEX14,DYNC2I2,DTNBP1,KATNIP,IQCG,DNAH17,KLC3,CFAP54
IL6R,TNIK,MBP,RBM20,PAX2,TGFB2,STOX1,NKD2,GATA3,DOK7,AKAP8L,RFX4,LEO1,TMPRSS6,MEPCE,FOXN1,ARID3A,LAT,FZR1,L3MBTL3,DCUN1D2,FLT1,ETV5,GLIS1,CRADD,AGAP2,SMARCC2
MCPH1,MBP,PAX2,TGFB2,TIMP3,STOX1,GATA3,DOK7,TRAPPC9,SHANK3,FXYD2,EPHA2,PEX14,MEPCE,EPHA3,DTNBP1,LAT,FZR1,PREX1,FLT1,ARHGEF15,ERCC2,CRADD
IL6R,TNIK,SETD1B,TGFB2,ITPKB,LARGE1,STOX1,GATA3,DOK7,AKAP8L,MGAT4B,RNF208,LEO1,MGAT4A,EPHA2,CHD5,NEK6,GAD1,PPIG,EPHA3,TYK2,DTNBP1,SBK1,LAT,FZR1,DCUN1D2,FLT1,TPST1,TGM4,EXT2,MAPK8IP1
TNIK,UNC5D,GATA3,RFX4,SHANK3,ODAD3,EPHA2,ESPN,DYNC2I2,EPHA3,DTNBP1,MBOAT1,IQCG,DNAH17,KLC3,CFAP54,RSPH1
PRICKLE2,NR2F1,IL6R,TARS3,TNIK,GRB10,MBP,RBM20,PAX2,CA5B,OFD1,SETD1B,TMEM120B,UNC5D,TGFB2,USH2A,ACIN1,ITPKB,TIMP3,SYN3,LARGE1,INSYN1,STOX1,NKD2,GATA3,SORCS2,DOK7,AKAP8L,TRAPPC9,RFX4,RNF208,SHANK3,NECTIN1,FXYD2,RNPC3,TPCN1,ODAD3,ACP5,LEO1,MYO5A,AHDC1,MYOM3,IGSF21,EPHA2,PEX14,ESPN,CHD5,AJAP1,MRPS2,DYNC2I2,NEK6,CNTFR,RECQL,TMPRSS6,EFCAB6,STAB2,GAD1,PPIG,ACMSD,MEPCE,SOSTDC1,MNT,SLC43A2,EPHA3,ARID3A,TYK2,
PRICKLE2,NR2F1,IL6R,TARS3,TNIK,GRB10,MBP,RBM20,PAX2,CA5B,OFD1,SETD1B,TMEM120B,UNC5D,TGFB2,USH2A,ACIN1,ITPKB,TIMP3,SYN3,LARGE1,INSYN1,STOX1,NKD2,GATA3,SORCS2,DOK7,AKAP8L,TRAPPC9,RFX4,RNF208,SHANK3,NECTIN1,FXYD2,RNPC3,TPCN1,ODAD3,ACP5,LEO1,MYO5A,AHDC1,MYOM3,IGSF21,EPHA2,PEX14,ESPN,CHD5,AJAP1,MRPS2,DYNC2I2,NEK6,CNTFR,RECQL,TMPRSS6,EFCAB6,STAB2,GAD1,PPIG,ACMSD,MEPCE,SOSTDC1,MNT,SLC43A2,EPHA3,ARID3A,TYK2,
PRICKLE2,NR2F1,TARS3,TNIK,GRB10,RBM20,CA5B,USH2A,ACIN1,ITPKB,SYN3,LARGE1,STOX1,NKD2,SORCS2,DOK7,AKAP8L,TRAPPC9,RNF208,TPCN1,ODAD3,ACP5,MYO5A,MYOM3,PEX14,CHD5,MRPS2,DYNC2I2,NEK6,STAB2,GAD1,PPIG,ACMSD,EPHA3,ARID3A,TYK2,DTNBP1,MBOAT1,ARVCF,XPO6,LAT,SPNS1,CARD19,SART1,PREX1,DCUN1D2,CACNA1G,ARHGEF15,ALOX5,TNS3,TPST1,OTOG,TGM4,NBEAL2,IQCG,ETFB,EXT2,MAPK8IP1,ARMC6,DNAH17,BPIFB4,ETHE1,KLC3,ERCC2,EHD2,ANXA6
IL6R,TNIK,UNC5D,TGFB2,USH2A,ACIN1,TIMP3,STOX1,NKD2,SORCS2,SHANK3,NECTIN1,IGSF21,EPHA2,AJAP1,CNTFR,TMPRSS6,STAB2,GAD1,SLC43A2,EPHA3,DTNBP1,ARVCF,IL4R,GSG1L,XPO6,LAT,VSIG1,COL4A2,NALCN,PCDH9,CACNA1G,ANKH,FLT1,CD68,OTOG,TGM4,MAPK8IP1,OTOP2,GRIN2C,GNG8,EHD2
IL6R,TNIK,TMEM120B,UNC5D,USH2A,ACIN1,ITPKB,SYN3,LARGE1,NKD2,SORCS2,SHANK3,NECTIN1,FXYD2,TPCN1,MYO5A,IGSF21,EPHA2,PEX14,AJAP1,CNTFR,TMPRSS6,STAB2,ACMSD,SLC43A2,EPHA3,ARID3A,TYK2,DTNBP1,MBOAT1,ARVCF,IL4R,IL21R,GSG1L,XPO6,LAT,FZR1,VSIG1,COL4A2,NALCN,PCDH9,CACNA1G,ANKH,FLT1,CD68,ALOX5,TPST1,ADAM11,EXT2,MAPK8IP1,OTOP2,GRIN2C,GNG8,EHD2,ANXA6,GNPTAB
TNIK,PAX2,OFD1,USH2A,ITPKB,STOX1,ODAD3,LEO1,MYO5A,DYNC2I2,NEK6,EPHA3,TYK2,DTNBP1,IL4R,FLT1,IQCG,DNAH17,BPIFB4,KLC3,ERCC2,EHD2,RSPH1
SYN3,INSYN1,SORCS2,SHANK3,NECTIN1,IGSF21,EPHA2,AJAP1,GAD1,DTNBP1,ARVCF,GSG1L,LAT,PCDH9,CACNA1G,ARHGEF15,TNS3,MAPK8IP1,GRIN2C,AGAP2
IL6R,TNIK,UNC5D,USH2A,ACIN1,NKD2,SORCS2,SHANK3,NECTIN1,IGSF21,EPHA2,AJAP1,CNTFR,TMPRSS6,STAB2,SLC43A2,EPHA3,DTNBP1,ARVCF,IL4R,GSG1L,XPO6,LAT,VSIG1,NALCN,PCDH9,CACNA1G,ANKH,FLT1,CD68,MAPK8IP1,OTOP2,GRIN2C,GNG8,EHD2
PAX2,OFD1,USH2A,STOX1,ODAD3,LEO1,DYNC2I2,NEK6,DTNBP1,IL4R,IQCG,DNAH17,KLC3,ERCC2,EHD2,RSPH1
IL6R,CNTFR
SYN3,INSYN1,SORCS2,SHANK3,NECTIN1,IGSF21,GAD1,DTNBP1,GSG1L,CACNA1G,ARHGEF15,MAPK8IP1,GRIN2C,AGAP2
NR2F1,TNIK,PAX2,SETD1B,ACIN1,STOX1,GATA3,DOK7,AKAP8L,RNF208,RNPC3,LEO1,CHD5,NEK6,RECQL,EFCAB6,PPIG,MNT,EPHA3,ARID3A,IL4R,XPO6,FZR1,L3MBTL3,SART1,ALOX5,ETV5,ETHE1,ERCC2,AGAP2,SMARCC2
NR2F1,UROS,MCPH1,IL6R,TNIK,GRB10,RBM20,PAX2,OFD1,SETD1B,TGFB2,USH2A,TIMP3,LARGE1,STOX1,GATA3,DOK7,TRAPPC9,SHANK3,NECTIN1,FXYD2,RNPC3,ODAD3,ACP5,MYO5A,AHDC1,EPHA2,PEX14,ESPN,CHD5,MRPS2,DYNC2I2,TMPRSS6,GAD1,FOXN1,TYK2,DTNBP1,ARVCF,IL4R,IL21R,KATNIP,LAT,FZR1,COL4A2,NALCN,CACNA1G,ANKH,FLT1,ALOX5,OTOG,NBEAL2,ETFB,EXT2,MAPK8IP1,DNAH17,ETHE1,ERCC2,GNPTAB,CRADD,SMARCC2,RSPH1
NR2F1,UROS,MCPH1,IL6R,TNIK,GRB10,RBM20,PAX2,OFD1,SETD1B,TGFB2,USH2A,TIMP3,LARGE1,STOX1,GATA3,DOK7,TRAPPC9,SHANK3,NECTIN1,FXYD2,RNPC3,ODAD3,ACP5,MYO5A,AHDC1,EPHA2,PEX14,ESPN,CHD5,MRPS2,DYNC2I2,TMPRSS6,GAD1,FOXN1,TYK2,DTNBP1,ARVCF,IL4R,IL21R,KATNIP,LAT,FZR1,COL4A2,NALCN,CACNA1G,ANKH,FLT1,ALOX5,OTOG,NBEAL2,ETFB,EXT2,MAPK8IP1,DNAH17,ETHE1,ERCC2,GNPTAB,CRADD,SMARCC2,RSPH1
NR2F1,UROS,MCPH1,IL6R,TNIK,RBM20,PAX2,OFD1,SETD1B,TGFB2,USH2A,TIMP3,LARGE1,STOX1,GATA3,DOK7,TRAPPC9,SHANK3,NECTIN1,FXYD2,RNPC3,ODAD3,ACP5,MYO5A,AHDC1,EPHA2,PEX14,ESPN,CHD5,MRPS2,DYNC2I2,TMPRSS6,GAD1,FOXN1,TYK2,DTNBP1,IL4R,IL21R,KATNIP,LAT,COL4A2,NALCN,CACNA1G,ANKH,ALOX5,OTOG,NBEAL2,ETFB,EXT2,MAPK8IP1,DNAH17,ETHE1,ERCC2,GNPTAB,CRADD,SMARCC2,RSPH1
NR2F1,UROS,MCPH1,IL6R,GRB10,PAX2,OFD1,SETD1B,TGFB2,USH2A,LARGE1,GATA3,DOK7,TRAPPC9,SHANK3,NECTIN1,RNPC3,ODAD3,ACP5,MYO5A,AHDC1,PEX14,ESPN,CHD5,MRPS2,DYNC2I2,GAD1,FOXN1,DTNBP1,ARVCF,IL4R,IL21R,KATNIP,FZR1,NALCN,CACNA1G,ANKH,NBEAL2,ETFB,EXT2,ERCC2,GNPTAB,SMARCC2,RSPH1
NR2F1,UROS,MCPH1,IL6R,TNIK,RBM20,PAX2,OFD1,SETD1B,TGFB2,USH2A,TIMP3,LARGE1,GATA3,DOK7,TRAPPC9,SHANK3,NECTIN1,FXYD2,RNPC3,ODAD3,ACP5,MYO5A,AHDC1,EPHA2,PEX14,ESPN,CHD5,MRPS2,DYNC2I2,TMPRSS6,GAD1,FOXN1,TYK2,DTNBP1,IL4R,IL21R,KATNIP,LAT,COL4A2,NALCN,CACNA1G,ANKH,ALOX5,OTOG,NBEAL2,ETFB,EXT2,MAPK8IP1,DNAH17,ETHE1,ERCC2,GNPTAB,CRADD,SMARCC2,RSPH1
NR2F1,UROS,MCPH1,IL6R,GRB10,PAX2,OFD1,SETD1B,TGFB2,USH2A,LARGE1,GATA3,DOK7,TRAPPC9,SHANK3,NECTIN1,RNPC3,ODAD3,ACP5,MYO5A,AHDC1,PEX14,ESPN,CHD5,MRPS2,DYNC2I2,GAD1,FOXN1,DTNBP1,ARVCF,IL4R,IL21R,KATNIP,FZR1,NALCN,CACNA1G,ANKH,NBEAL2,ETFB,EXT2,ERCC2,GNPTAB,CRADD,SMARCC2,RSPH1
NR2F1,UROS,MCPH1,IL6R,GRB10,PAX2,OFD1,SETD1B,TGFB2,USH2A,LARGE1,GATA3,DOK7,TRAPPC9,SHANK3,NECTIN1,RNPC3,ODAD3,ACP5,MYO5A,AHDC1,PEX14,ESPN,CHD5,MRPS2,DYNC2I2,GAD1,FOXN1,DTNBP1,ARVCF,IL4R,IL21R,KATNIP,FZR1,NALCN,CACNA1G,ANKH,NBEAL2,ETFB,EXT2,ERCC2,GNPTAB,CRADD,SMARCC2,RSPH1
NR2F1,UROS,MCPH1,OFD1,SETD1B,TGFB2,DOK7,TRAPPC9,SHANK3,NECTIN1,MYO5A,AHDC1,PEX14,CHD5,MRPS2,DYNC2I2,GAD1,ARVCF,FZR1,NALCN,CACNA1G,ANKH,ETFB,EXT2,ERCC2,GNPTAB,SMARCC2
NR2F1,IL6R,OFD1,USH2A,DOK7,TRAPPC9,SHANK3,NECTIN1,ODAD3,ACP5,AHDC1,PEX14,CHD5,DYNC2I2,GAD1,FOXN1,DTNBP1,ARVCF,IL4R,IL21R,KATNIP,NALCN,CACNA1G,ANKH,NBEAL2,ETFB,EXT2,ERCC2,GNPTAB,SMARCC2,RSPH1
NR2F1,UROS,MCPH1,PAX2,OFD1,SETD1B,TGFB2,DOK7,TRAPPC9,SHANK3,NECTIN1,MYO5A,AHDC1,PEX14,CHD5,MRPS2,DYNC2I2,GAD1,ARVCF,KATNIP,FZR1,NALCN,CACNA1G,ANKH,ETFB,EXT2,ERCC2,GNPTAB,SMARCC2
NR2F1,IL6R,GRB10,OFD1,USH2A,LARGE1,DOK7,TRAPPC9,SHANK3,NECTIN1,ODAD3,ACP5,AHDC1,PEX14,ESPN,CHD5,MRPS2,GAD1,ARVCF,IL21R,KATNIP,NALCN,CACNA1G,ANKH,ETFB,EXT2,ERCC2,GNPTAB,RSPH1
NR2F1,UROS,MCPH1,TNIK,GRB10,PAX2,OFD1,SETD1B,TGFB2,USH2A,LARGE1,STOX1,GATA3,DOK7,TRAPPC9,SHANK3,NECTIN1,FXYD2,ODAD3,ACP5,MYO5A,AHDC1,PEX14,ESPN,CHD5,MRPS2,DYNC2I2,TMPRSS6,GAD1,ARVCF,KATNIP,FZR1,COL4A2,NALCN,CACNA1G,ANKH,FLT1,OTOG,ETFB,EXT2,ETHE1,ERCC2,GNPTAB,CRADD,SMARCC2,RSPH1
NR2F1,UROS,MCPH1,OFD1,SETD1B,TGFB2,DOK7,TRAPPC9,SHANK3,NECTIN1,MYO5A,AHDC1,PEX14,CHD5,MRPS2,DYNC2I2,GAD1,ARVCF,KATNIP,FZR1,NALCN,CACNA1G,ANKH,ETFB,EXT2,ERCC2,GNPTAB,SMARCC2
NR2F1,UROS,MCPH1,OFD1,SETD1B,TGFB2,DOK7,TRAPPC9,SHANK3,NECTIN1,MYO5A,AHDC1,PEX14,CHD5,MRPS2,DYNC2I2,GAD1,ARVCF,KATNIP,FZR1,NALCN,CACNA1G,ANKH,ETFB,EXT2,ERCC2,GNPTAB,SMARCC2
NR2F1,UROS,MCPH1,TNIK,GRB10,PAX2,OFD1,SETD1B,TGFB2,USH2A,LARGE1,STOX1,GATA3,DOK7,TRAPPC9,SHANK3,NECTIN1,FXYD2,RNPC3,ODAD3,ACP5,MYO5A,AHDC1,PEX14,ESPN,CHD5,MRPS2,DYNC2I2,TMPRSS6,GAD1,ARVCF,KATNIP,FZR1,COL4A2,NALCN,CACNA1G,ANKH,FLT1,OTOG,ETFB,EXT2,ETHE1,ERCC2,GNPTAB,CRADD,SMARCC2,RSPH1
NR2F1,MCPH1,TNIK,GRB10,PAX2,OFD1,SETD1B,LARGE1,DOK7,TRAPPC9,SHANK3,NECTIN1,ODAD3,ACP5,MYO5A,AHDC1,PEX14,ESPN,CHD5,MRPS2,DYNC2I2,GAD1,ARVCF,KATNIP,FZR1,COL4A2,NALCN,CACNA1G,EXT2,ETHE1,ERCC2,GNPTAB,CRADD,SMARCC2,RSPH1
UROS,IL6R,GRB10,PAX2,OFD1,TGFB2,STOX1,GATA3,SHANK3,NECTIN1,ODAD3,ACP5,MYO5A,DYNC2I2,FOXN1,TYK2,ARVCF,IL4R,IL21R,KATNIP,LAT,CACNA1G,FLT1,ALOX5,ERCC2,GNPTAB,SMARCC2,RSPH1
NR2F1,UROS,MCPH1,IL6R,GRB10,RBM20,PAX2,OFD1,SETD1B,TGFB2,USH2A,LARGE1,GATA3,DOK7,TRAPPC9,SHANK3,NECTIN1,FXYD2,RNPC3,ODAD3,ACP5,MYO5A,AHDC1,PEX14,ESPN,CHD5,MRPS2,DYNC2I2,GAD1,ARVCF,IL21R,KATNIP,FZR1,COL4A2,NALCN,CACNA1G,ANKH,ETFB,EXT2,ETHE1,ERCC2,GNPTAB,CRADD,SMARCC2,RSPH1
NR2F1,IL6R,GRB10,RBM20,PAX2,OFD1,USH2A,LARGE1,GATA3,DOK7,TRAPPC9,SHANK3,NECTIN1,ODAD3,ACP5,AHDC1,PEX14,ESPN,CHD5,MRPS2,GAD1,ARVCF,IL21R,KATNIP,NALCN,CACNA1G,ANKH,OTOG,ETFB,EXT2,ERCC2,GNPTAB,SMARCC2,RSPH1
NR2F1,UROS,IL6R,GRB10,RBM20,PAX2,OFD1,SETD1B,TGFB2,USH2A,GATA3,DOK7,TRAPPC9,SHANK3,NECTIN1,ACP5,MYO5A,PEX14,CHD5,MRPS2,TMPRSS6,GAD1,FOXN1,DTNBP1,ARVCF,IL4R,KATNIP,NALCN,CACNA1G,NBEAL2,ETFB,EXT2,ETHE1,ERCC2,GNPTAB,SMARCC2
NR2F1,MCPH1,TNIK,GRB10,PAX2,OFD1,SETD1B,USH2A,LARGE1,DOK7,TRAPPC9,SHANK3,NECTIN1,ODAD3,ACP5,MYO5A,AHDC1,PEX14,ESPN,CHD5,MRPS2,DYNC2I2,TMPRSS6,GAD1,ARVCF,KATNIP,FZR1,NALCN,CACNA1G,ETFB,EXT2,ETHE1,ERCC2,GNPTAB,CRADD,SMARCC2,RSPH1
NR2F1,UROS,MCPH1,GRB10,PAX2,OFD1,SETD1B,TGFB2,USH2A,LARGE1,GATA3,DOK7,TRAPPC9,SHANK3,NECTIN1,FXYD2,RNPC3,ODAD3,ACP5,MYO5A,AHDC1,PEX14,ESPN,CHD5,DYNC2I2,GAD1,ARVCF,IL21R,KATNIP,FZR1,NALCN,CACNA1G,ANKH,ETFB,EXT2,ERCC2,GNPTAB,CRADD,SMARCC2,RSPH1
IL6R,OFD1,ACP5,FOXN1,ARVCF,IL21R,KATNIP,ANKH,GNPTAB,SMARCC2,RSPH1
NR2F1,GRB10,OFD1,TGFB2,LARGE1,DOK7,SHANK3,NECTIN1,ODAD3,ACP5,AHDC1,PEX14,CHD5,DYNC2I2,GAD1,ARVCF,IL21R,NALCN,CACNA1G,ANKH,EXT2,ERCC2,GNPTAB,RSPH1
NR2F1,MCPH1,TNIK,GRB10,PAX2,OFD1,SETD1B,USH2A,LARGE1,DOK7,TRAPPC9,SHANK3,NECTIN1,ODAD3,ACP5,MYO5A,AHDC1,PEX14,ESPN,CHD5,MRPS2,DYNC2I2,TMPRSS6,GAD1,ARVCF,KATNIP,FZR1,COL4A2,NALCN,CACNA1G,EXT2,ETHE1,ERCC2,GNPTAB,CRADD,SMARCC2,RSPH1
UROS,IL6R,GRB10,RBM20,PAX2,OFD1,TGFB2,STOX1,GATA3,SHANK3,NECTIN1,ODAD3,ACP5,MYO5A,DYNC2I2,FOXN1,TYK2,ARVCF,IL4R,IL21R,KATNIP,LAT,CACNA1G,FLT1,ALOX5,NBEAL2,ERCC2,GNPTAB,SMARCC2,RSPH1
NR2F1,MCPH1,GRB10,OFD1,SETD1B,TGFB2,LARGE1,DOK7,TRAPPC9,SHANK3,NECTIN1,RNPC3,ODAD3,ACP5,AHDC1,PEX14,CHD5,DYNC2I2,GAD1,ARVCF,IL21R,KATNIP,FZR1,NALCN,CACNA1G,ANKH,ETFB,EXT2,ERCC2,GNPTAB,CRADD,SMARCC2,RSPH1
NR2F1,MCPH1,TNIK,PAX2,OFD1,SETD1B,USH2A,LARGE1,TRAPPC9,SHANK3,NECTIN1,ACP5,MYO5A,AHDC1,ESPN,CHD5,MRPS2,TMPRSS6,GAD1,ARVCF,KATNIP,FZR1,NALCN,CACNA1G,EXT2,ETHE1,ERCC2,GNPTAB,CRADD,SMARCC2
NR2F1,UROS,MCPH1,GRB10,PAX2,OFD1,TGFB2,USH2A,LARGE1,STOX1,GATA3,DOK7,TRAPPC9,SHANK3,NECTIN1,RNPC3,ACP5,AHDC1,PEX14,MRPS2,DYNC2I2,ARVCF,IL21R,KATNIP,LAT,FZR1,COL4A2,NALCN,FLT1,EXT2,MAPK8IP1,ETHE1,ERCC2,GNPTAB,SMARCC2
NR2F1,TNIK,GRB10,OFD1,SETD1B,LARGE1,TRAPPC9,SHANK3,NECTIN1,ODAD3,AHDC1,CHD5,DYNC2I2,FZR1,NALCN,EXT2,ERCC2,GNPTAB,CRADD,SMARCC2,RSPH1
NR2F1,MCPH1,GRB10,PAX2,OFD1,TGFB2,USH2A,LARGE1,GATA3,DOK7,TRAPPC9,SHANK3,RNPC3,ODAD3,ACP5,MYO5A,AHDC1,PEX14,ESPN,DYNC2I2,GAD1,ARVCF,KATNIP,FZR1,COL4A2,NALCN,CACNA1G,ETFB,EXT2,ETHE1,ERCC2,GNPTAB,CRADD,SMARCC2,RSPH1
NR2F1,UROS,IL6R,GRB10,RBM20,PAX2,OFD1,TGFB2,USH2A,GATA3,DOK7,SHANK3,NECTIN1,ACP5,MYO5A,PEX14,MRPS2,TMPRSS6,FOXN1,DTNBP1,ARVCF,IL4R,NALCN,CACNA1G,NBEAL2,ETFB,EXT2,ETHE1,ERCC2,GNPTAB,SMARCC2
MCPH1,OFD1,USH2A,LARGE1,GATA3,TRAPPC9,ACP5,MYO5A,AHDC1,PEX14,ESPN,KATNIP,COL4A2,ETFB,ERCC2,GNPTAB,CRADD,SMARCC2
UROS,IL6R,OFD1,STOX1,SHANK3,NECTIN1,ODAD3,ACP5,MYO5A,DYNC2I2,FOXN1,TYK2,IL21R,KATNIP,LAT,FLT1,ERCC2,GNPTAB,SMARCC2,RSPH1
NR2F1,UROS,MCPH1,GRB10,PAX2,OFD1,SETD1B,TGFB2,LARGE1,DOK7,TRAPPC9,SHANK3,NECTIN1,RNPC3,ODAD3,ACP5,AHDC1,PEX14,CHD5,DYNC2I2,GAD1,ARVCF,IL21R,KATNIP,FZR1,NALCN,CACNA1G,ANKH,ETFB,EXT2,ERCC2,GNPTAB,CRADD,SMARCC2,RSPH1
NR2F1,UROS,PAX2,OFD1,TGFB2,USH2A,TIMP3,LARGE1,STOX1,GATA3,DOK7,TRAPPC9,SHANK3,ODAD3,MYO5A,AHDC1,EPHA2,PEX14,ESPN,MRPS2,DYNC2I2,GAD1,DTNBP1,ARVCF,KATNIP,FZR1,COL4A2,NALCN,CACNA1G,ANKH,FLT1,ETFB,EXT2,ETHE1,ERCC2,GNPTAB,SMARCC2,RSPH1
IL6R,OFD1,NECTIN1,ODAD3,ACP5,MYO5A,DYNC2I2,FOXN1,TYK2,IL21R,KATNIP,LAT,ERCC2,GNPTAB,SMARCC2,RSPH1
NR2F1,MCPH1,OFD1,TGFB2,USH2A,LARGE1,GATA3,TRAPPC9,SHANK3,RNPC3,ACP5,MYO5A,AHDC1,PEX14,ESPN,DYNC2I2,GAD1,ARVCF,KATNIP,FZR1,COL4A2,NALCN,CACNA1G,ETFB,EXT2,ETHE1,ERCC2,GNPTAB,CRADD,SMARCC2
NR2F1,TNIK,GRB10,OFD1,SETD1B,LARGE1,TRAPPC9,SHANK3,NECTIN1,ODAD3,AHDC1,CHD5,DYNC2I2,FZR1,NALCN,EXT2,ERCC2,GNPTAB,CRADD,SMARCC2,RSPH1
NR2F1,OFD1,USH2A,DOK7,TRAPPC9,SHANK3,NECTIN1,AHDC1,PEX14,CHD5,DYNC2I2,GAD1,ARVCF,KATNIP,NALCN,ANKH,ETFB,EXT2,ERCC2,GNPTAB,SMARCC2
IL6R,OFD1,ACP5,FOXN1,IL21R,KATNIP,ANKH,GNPTAB,SMARCC2,RSPH1
UROS,MCPH1,IL6R,GRB10,RBM20,PAX2,OFD1,USH2A,LARGE1,STOX1,GATA3,DOK7,TRAPPC9,SHANK3,FXYD2,ACP5,MYO5A,PEX14,MRPS2,DYNC2I2,TMPRSS6,TYK2,ARVCF,IL4R,IL21R,LAT,NALCN,ANKH,FLT1,ETFB,MAPK8IP1,ETHE1,ERCC2,GNPTAB
IL6R,PAX2,OFD1,TGFB2,DOK7,NECTIN1,ODAD3,ACP5,MYO5A,DYNC2I2,FOXN1,TYK2,ARVCF,IL21R,KATNIP,LAT,ETFB,EXT2,ERCC2,GNPTAB,SMARCC2,RSPH1
IL6R,PAX2,OFD1,TGFB2,DOK7,NECTIN1,ODAD3,ACP5,MYO5A,AHDC1,DYNC2I2,FOXN1,TYK2,ARVCF,IL21R,KATNIP,LAT,ANKH,ETFB,EXT2,ERCC2,GNPTAB,SMARCC2,RSPH1
NR2F1,OFD1,USH2A,DOK7,TRAPPC9,SHANK3,NECTIN1,ODAD3,ACP5,AHDC1,PEX14,CHD5,DYNC2I2,GAD1,ARVCF,KATNIP,NALCN,CACNA1G,ANKH,ETFB,EXT2,ERCC2,GNPTAB,SMARCC2,RSPH1
IL6R,GRB10,PAX2,OFD1,TGFB2,LARGE1,DOK7,NECTIN1,ODAD3,ACP5,MYO5A,AHDC1,PEX14,DYNC2I2,FOXN1,TYK2,ARVCF,IL4R,IL21R,KATNIP,LAT,NALCN,ANKH,ALOX5,ETFB,EXT2,ERCC2,GNPTAB,SMARCC2,RSPH1
NR2F1,MCPH1,OFD1,TGFB2,USH2A,LARGE1,GATA3,TRAPPC9,SHANK3,RNPC3,ACP5,MYO5A,AHDC1,PEX14,ESPN,DYNC2I2,GAD1,ARVCF,KATNIP,FZR1,COL4A2,NALCN,CACNA1G,ETFB,EXT2,ETHE1,ERCC2,GNPTAB,CRADD,SMARCC2
NR2F1,UROS,PAX2,OFD1,TGFB2,USH2A,TIMP3,LARGE1,GATA3,DOK7,TRAPPC9,SHANK3,ODAD3,MYO5A,AHDC1,EPHA2,PEX14,ESPN,DYNC2I2,GAD1,DTNBP1,ARVCF,KATNIP,FZR1,NALCN,CACNA1G,ANKH,ETFB,EXT2,ETHE1,ERCC2,GNPTAB,RSPH1
IL6R,OFD1,ACP5,FOXN1,IL21R,KATNIP,ANKH,GNPTAB,SMARCC2,RSPH1
UROS,IL6R,OFD1,STOX1,SHANK3,NECTIN1,ODAD3,ACP5,MYO5A,DYNC2I2,FOXN1,TYK2,IL21R,KATNIP,LAT,FLT1,ERCC2,GNPTAB,SMARCC2,RSPH1
IL6R,OFD1,ODAD3,ACP5,AHDC1,DYNC2I2,FOXN1,ARVCF,IL21R,KATNIP,ANKH,GNPTAB,SMARCC2,RSPH1
IL6R,PAX2,OFD1,NECTIN1,ODAD3,ACP5,MYO5A,DYNC2I2,FOXN1,TYK2,IL21R,KATNIP,LAT,ERCC2,GNPTAB,SMARCC2,RSPH1
GRB10,OFD1,SETD1B,TGFB2,TRAPPC9,SHANK3,RNPC3,ACP5,AHDC1,PEX14,CHD5,DYNC2I2,ARVCF,KATNIP,NALCN,ANKH,EXT2,ERCC2,GNPTAB
NR2F1,GRB10,OFD1,TGFB2,LARGE1,DOK7,SHANK3,NECTIN1,AHDC1,PEX14,CHD5,DYNC2I2,GAD1,ARVCF,NALCN,CACNA1G,ANKH,EXT2,ERCC2,GNPTAB
NR2F1,MCPH1,PAX2,OFD1,SETD1B,LARGE1,GATA3,TRAPPC9,SHANK3,NECTIN1,FXYD2,ACP5,MYO5A,AHDC1,PEX14,CHD5,GAD1,ARVCF,KATNIP,FZR1,COL4A2,NALCN,ANKH,EXT2,ETHE1,ERCC2,CRADD,SMARCC2
NR2F1,MCPH1,GRB10,PAX2,OFD1,TGFB2,USH2A,LARGE1,GATA3,DOK7,TRAPPC9,SHANK3,RNPC3,ODAD3,ACP5,MYO5A,AHDC1,PEX14,ESPN,DYNC2I2,GAD1,ARVCF,KATNIP,FZR1,COL4A2,NALCN,CACNA1G,ETFB,EXT2,ETHE1,ERCC2,GNPTAB,CRADD,SMARCC2,RSPH1
IL6R,OFD1,ACP5,FOXN1,IL21R,KATNIP,GNPTAB,SMARCC2,RSPH1
UROS,IL6R,TNIK,RBM20,PAX2,OFD1,TIMP3,LARGE1,STOX1,GATA3,DOK7,TRAPPC9,SHANK3,NECTIN1,ODAD3,ACP5,MYO5A,AHDC1,PEX14,CHD5,MRPS2,GAD1,IL21R,KATNIP,LAT,NALCN,CACNA1G,ANKH,NBEAL2,ETFB,EXT2,MAPK8IP1,DNAH17,ETHE1,ERCC2,GNPTAB
UROS,IL6R,GRB10,PAX2,OFD1,TGFB2,GATA3,SHANK3,NECTIN1,ODAD3,ACP5,FOXN1,ARVCF,IL4R,IL21R,LAT,CACNA1G,ERCC2,GNPTAB,SMARCC2,RSPH1
UROS,IL6R,GRB10,PAX2,OFD1,TGFB2,GATA3,SHANK3,NECTIN1,ODAD3,ACP5,FOXN1,ARVCF,IL4R,IL21R,LAT,CACNA1G,ERCC2,GNPTAB,SMARCC2,RSPH1
NR2F1,MCPH1,GRB10,PAX2,OFD1,TGFB2,USH2A,LARGE1,GATA3,DOK7,TRAPPC9,SHANK3,RNPC3,ODAD3,ACP5,MYO5A,AHDC1,PEX14,ESPN,DYNC2I2,GAD1,ARVCF,KATNIP,FZR1,COL4A2,NALCN,CACNA1G,ANKH,ETFB,EXT2,ETHE1,ERCC2,GNPTAB,CRADD,SMARCC2,RSPH1
NR2F1,UROS,PAX2,OFD1,TGFB2,USH2A,TIMP3,LARGE1,STOX1,GATA3,DOK7,TRAPPC9,SHANK3,MYO5A,AHDC1,EPHA2,PEX14,ESPN,MRPS2,DTNBP1,ARVCF,KATNIP,FZR1,COL4A2,NALCN,CACNA1G,ANKH,FLT1,EXT2,ERCC2,GNPTAB,SMARCC2
UROS,MCPH1,GRB10,PAX2,OFD1,USH2A,LARGE1,STOX1,GATA3,DOK7,SHANK3,NECTIN1,FXYD2,ODAD3,ACP5,PEX14,MRPS2,DYNC2I2,GAD1,ARVCF,KATNIP,NALCN,CACNA1G,FLT1,NBEAL2,ETFB,EXT2,DNAH17,ETHE1,ERCC2,GNPTAB,SMARCC2,RSPH1
NR2F1,MCPH1,OFD1,SETD1B,TGFB2,DOK7,NECTIN1,AHDC1,PEX14,CHD5,MRPS2,GAD1,ARVCF,FZR1,NALCN,CACNA1G
NR2F1,UROS,PAX2,OFD1,USH2A,TIMP3,LARGE1,STOX1,DOK7,TRAPPC9,SHANK3,MYO5A,AHDC1,PEX14,ESPN,DTNBP1,FZR1,CACNA1G,ANKH,FLT1,ERCC2,SMARCC2
UROS,IL6R,TNIK,RBM20,PAX2,OFD1,TIMP3,LARGE1,STOX1,GATA3,DOK7,TRAPPC9,SHANK3,NECTIN1,ODAD3,ACP5,MYO5A,AHDC1,PEX14,CHD5,MRPS2,GAD1,IL21R,KATNIP,LAT,NALCN,CACNA1G,ANKH,NBEAL2,ETFB,EXT2,MAPK8IP1,DNAH17,ETHE1,ERCC2,GNPTAB
NR2F1,GRB10,OFD1,TGFB2,LARGE1,DOK7,SHANK3,NECTIN1,AHDC1,PEX14,CHD5,DYNC2I2,GAD1,ARVCF,NALCN,CACNA1G,ANKH,EXT2,ERCC2,GNPTAB
UROS,MCPH1,IL6R,TNIK,USH2A,LARGE1,DOK7,TRAPPC9,NECTIN1,RNPC3,ODAD3,ACP5,MYO5A,PEX14,ESPN,MRPS2,DYNC2I2,TMPRSS6,GAD1,FOXN1,TYK2,DTNBP1,IL21R,KATNIP,LAT,NALCN,OTOG,NBEAL2,ETFB,EXT2,DNAH17,ETHE1,ERCC2,GNPTAB,CRADD,RSPH1
UROS,IL6R,RBM20,PAX2,TGFB2,STOX1,GATA3,ACP5,PEX14,TMPRSS6,FOXN1,TYK2,DTNBP1,ARVCF,IL4R,IL21R,LAT,COL4A2,FLT1,NBEAL2,EXT2,ETHE1,ERCC2
GRB10,PAX2,OFD1,USH2A,LARGE1,STOX1,GATA3,DOK7,NECTIN1,ODAD3,PEX14,DYNC2I2,GAD1,ARVCF,KATNIP,NALCN,CACNA1G,FLT1,NBEAL2,ETFB,EXT2,DNAH17,ERCC2,SMARCC2,RSPH1
NR2F1,UROS,MCPH1,GRB10,OFD1,SETD1B,TGFB2,USH2A,LARGE1,GATA3,DOK7,TRAPPC9,SHANK3,NECTIN1,ACP5,PEX14,ESPN,CHD5,DYNC2I2,GAD1,ARVCF,KATNIP,FZR1,NALCN,ANKH,EXT2,ERCC2,GNPTAB,SMARCC2
NR2F1,UROS,MCPH1,GRB10,OFD1,SETD1B,TGFB2,USH2A,LARGE1,GATA3,DOK7,TRAPPC9,SHANK3,NECTIN1,ACP5,PEX14,ESPN,CHD5,DYNC2I2,GAD1,ARVCF,KATNIP,FZR1,NALCN,ANKH,EXT2,ERCC2,GNPTAB,SMARCC2
NR2F1,PAX2,OFD1,USH2A,TIMP3,LARGE1,GATA3,TRAPPC9,ODAD3,MYO5A,EPHA2,PEX14,ESPN,DYNC2I2,DTNBP1,ARVCF,FZR1,NALCN,ETHE1,ERCC2,GNPTAB,RSPH1
NR2F1,UROS,MCPH1,GRB10,PAX2,OFD1,SETD1B,TGFB2,LARGE1,DOK7,TRAPPC9,SHANK3,NECTIN1,FXYD2,RNPC3,ODAD3,ACP5,AHDC1,PEX14,CHD5,DYNC2I2,GAD1,ARVCF,IL21R,KATNIP,FZR1,NALCN,CACNA1G,ANKH,ETFB,EXT2,ERCC2,GNPTAB,CRADD,SMARCC2,RSPH1
NR2F1,PAX2,OFD1,USH2A,TIMP3,LARGE1,GATA3,TRAPPC9,ODAD3,MYO5A,EPHA2,PEX14,ESPN,DYNC2I2,DTNBP1,ARVCF,FZR1,NALCN,ETHE1,ERCC2,GNPTAB,RSPH1
NR2F1,GRB10,OFD1,LARGE1,DOK7,TRAPPC9,SHANK3,NECTIN1,ACP5,AHDC1,PEX14,CHD5,MRPS2,GAD1,ARVCF,KATNIP,NALCN,CACNA1G,ETFB,EXT2,ERCC2
UROS,GRB10,PAX2,OFD1,TGFB2,LARGE1,DOK7,TRAPPC9,SHANK3,NECTIN1,ACP5,AHDC1,DYNC2I2,GAD1,ARVCF,KATNIP,NALCN,CACNA1G,ANKH,EXT2,ERCC2,GNPTAB,SMARCC2
PAX2,OFD1,USH2A,TIMP3,LARGE1,GATA3,TRAPPC9,ODAD3,MYO5A,PEX14,ESPN,DYNC2I2,ARVCF,FZR1,ETHE1,ERCC2,GNPTAB,RSPH1
OFD1,USH2A,TIMP3,LARGE1,GATA3,TRAPPC9,ODAD3,MYO5A,PEX14,DYNC2I2,RSPH1
NR2F1,UROS,GRB10,PAX2,OFD1,TGFB2,LARGE1,STOX1,DOK7,SHANK3,NECTIN1,ODAD3,ACP5,AHDC1,PEX14,DYNC2I2,TMPRSS6,GAD1,ARVCF,IL21R,KATNIP,LAT,FZR1,NALCN,CACNA1G,FLT1,NBEAL2,ETFB,EXT2,ETHE1,ERCC2,GNPTAB,SMARCC2,RSPH1
NR2F1,UROS,MCPH1,GRB10,OFD1,SETD1B,TGFB2,USH2A,LARGE1,GATA3,DOK7,TRAPPC9,SHANK3,NECTIN1,ACP5,PEX14,ESPN,CHD5,DYNC2I2,GAD1,ARVCF,KATNIP,FZR1,NALCN,ANKH,EXT2,ERCC2,GNPTAB,SMARCC2
GRB10,PAX2,OFD1,TGFB2,LARGE1,DOK7,NECTIN1,ODAD3,ACP5,AHDC1,PEX14,DYNC2I2,ARVCF,IL4R,KATNIP,LAT,NALCN,ALOX5,ETFB,ERCC2,GNPTAB,RSPH1
GRB10,OFD1,LARGE1,DOK7,SHANK3,NECTIN1,AHDC1,PEX14,CHD5,DYNC2I2,GAD1,ARVCF,NALCN,EXT2,ERCC2,GNPTAB
NR2F1,TNIK,GRB10,PAX2,OFD1,SETD1B,USH2A,LARGE1,STOX1,DOK7,TRAPPC9,SHANK3,NECTIN1,AHDC1,PEX14,ESPN,CHD5,ARVCF,FZR1,NALCN,CACNA1G,FLT1,EXT2,ERCC2,GNPTAB,SMARCC2
NR2F1,TNIK,GRB10,OFD1,SETD1B,LARGE1,TRAPPC9,SHANK3,NECTIN1,ODAD3,AHDC1,CHD5,MRPS2,DYNC2I2,ARVCF,FZR1,NALCN,CACNA1G,EXT2,ERCC2,GNPTAB,CRADD,SMARCC2,RSPH1
GRB10,OFD1,LARGE1,DOK7,SHANK3,NECTIN1,AHDC1,PEX14,CHD5,DYNC2I2,GAD1,ARVCF,NALCN,EXT2,ERCC2,GNPTAB
GRB10,OFD1,LARGE1,DOK7,SHANK3,NECTIN1,AHDC1,PEX14,CHD5,DYNC2I2,GAD1,ARVCF,NALCN,EXT2,ERCC2,GNPTAB
NR2F1,GRB10,PAX2,OFD1,LARGE1,SHANK3,NECTIN1,AHDC1,PEX14,DYNC2I2,ARVCF,KATNIP,FZR1,NALCN,EXT2,ETHE1,ERCC2,GNPTAB,SMARCC2
NR2F1,UROS,GRB10,RBM20,OFD1,TGFB2,LARGE1,GATA3,DOK7,TRAPPC9,SHANK3,NECTIN1,FXYD2,ACP5,MYO5A,AHDC1,PEX14,CHD5,MRPS2,GAD1,ARVCF,KATNIP,FZR1,COL4A2,NALCN,CACNA1G,ANKH,ETFB,EXT2,ETHE1,ERCC2,GNPTAB,SMARCC2
UROS,PAX2,OFD1,TGFB2,STOX1,DOK7,NECTIN1,ODAD3,PEX14,DYNC2I2,ARVCF,FZR1,NALCN,FLT1,EXT2,ERCC2,GNPTAB,RSPH1
NR2F1,PAX2,OFD1,USH2A,LARGE1,SHANK3,MYO5A,AHDC1,EPHA2,ESPN,CACNA1G,ERCC2,GNPTAB,SMARCC2
IL6R,OFD1,NECTIN1,ODAD3,ACP5,IL21R,GNPTAB,RSPH1
OFD1,GATA3,SHANK3,ARVCF,ETFB
NR2F1,MCPH1,OFD1,SETD1B,TGFB2,DOK7,NECTIN1,AHDC1,PEX14,CHD5,MRPS2,GAD1,ARVCF,FZR1,NALCN,CACNA1G
NR2F1,UROS,GRB10,RBM20,OFD1,SETD1B,TGFB2,LARGE1,DOK7,TRAPPC9,SHANK3,NECTIN1,ODAD3,ACP5,PEX14,MRPS2,DYNC2I2,GAD1,ARVCF,KATNIP,FZR1,NALCN,CACNA1G,ANKH,EXT2,ERCC2,GNPTAB,SMARCC2,RSPH1
NR2F1,MCPH1,OFD1,LARGE1,GATA3,TRAPPC9,SHANK3,ACP5,AHDC1,DYNC2I2,GAD1,KATNIP,FZR1,NALCN,CACNA1G,ETHE1,ERCC2,GNPTAB,SMARCC2
NR2F1,MCPH1,GRB10,OFD1,SETD1B,LARGE1,TRAPPC9,SHANK3,ACP5,MYO5A,AHDC1,PEX14,ESPN,MRPS2,GAD1,ARVCF,KATNIP,FZR1,COL4A2,NALCN,CACNA1G,EXT2,ETHE1,ERCC2,GNPTAB
GRB10,PAX2,OFD1,USH2A,STOX1,TRAPPC9,SHANK3,NECTIN1,RNPC3,AHDC1,PEX14,MRPS2,DYNC2I2,ARVCF,IL21R,LAT,FZR1,NALCN,FLT1,MAPK8IP1,ETHE1,ERCC2,GNPTAB,SMARCC2
OFD1,TGFB2,DOK7,TRAPPC9,SHANK3,AHDC1,GAD1,ARVCF,KATNIP,NALCN,CACNA1G,ANKH,EXT2,ERCC2,GNPTAB
NR2F1,GRB10,PAX2,OFD1,SETD1B,TRAPPC9,NECTIN1,AHDC1,PEX14,GAD1,ARVCF,KATNIP,NALCN,ETFB,EXT2,ERCC2,GNPTAB,SMARCC2
PAX2,OFD1,USH2A,LARGE1,DOK7,TRAPPC9,SHANK3,MYO5A,AHDC1,ESPN,DYNC2I2,GAD1,KATNIP,NALCN,CACNA1G,ERCC2,SMARCC2
OFD1,USH2A,LARGE1,DOK7,TRAPPC9,MYO5A,ESPN,DYNC2I2,KATNIP,CACNA1G,ERCC2,SMARCC2
PAX2,OFD1,USH2A,LARGE1,DOK7,TRAPPC9,SHANK3,MYO5A,AHDC1,ESPN,DYNC2I2,GAD1,KATNIP,NALCN,CACNA1G,ERCC2,SMARCC2
PAX2,OFD1,USH2A,LARGE1,DOK7,TRAPPC9,SHANK3,MYO5A,AHDC1,ESPN,DYNC2I2,GAD1,KATNIP,NALCN,CACNA1G,ERCC2,SMARCC2
UROS,IL6R,TNIK,RBM20,PAX2,OFD1,TIMP3,LARGE1,STOX1,GATA3,DOK7,TRAPPC9,SHANK3,NECTIN1,ODAD3,ACP5,MYO5A,AHDC1,CHD5,MRPS2,GAD1,IL21R,KATNIP,NALCN,ANKH,EXT2,MAPK8IP1,DNAH17,ETHE1,ERCC2,GNPTAB
NR2F1,GRB10,OFD1,SETD1B,TGFB2,LARGE1,DOK7,TRAPPC9,SHANK3,NECTIN1,ODAD3,ACP5,DYNC2I2,ARVCF,KATNIP,NALCN,CACNA1G,ANKH,EXT2,ERCC2,GNPTAB,SMARCC2,RSPH1
NR2F1,GRB10,OFD1,SETD1B,TGFB2,LARGE1,DOK7,TRAPPC9,SHANK3,NECTIN1,ODAD3,ACP5,DYNC2I2,ARVCF,KATNIP,NALCN,CACNA1G,ANKH,EXT2,ERCC2,GNPTAB,SMARCC2,RSPH1
NR2F1,UROS,MCPH1,GRB10,PAX2,OFD1,LARGE1,STOX1,GATA3,DOK7,RNPC3,ACP5,PEX14,DYNC2I2,ARVCF,KATNIP,FZR1,COL4A2,NALCN,FLT1,EXT2,ERCC2,GNPTAB,SMARCC2
NR2F1,UROS,GRB10,RBM20,OFD1,SETD1B,TGFB2,LARGE1,DOK7,TRAPPC9,SHANK3,NECTIN1,ACP5,PEX14,DYNC2I2,ARVCF,KATNIP,NALCN,EXT2,ERCC2,GNPTAB,SMARCC2
NR2F1,GRB10,OFD1,SETD1B,TGFB2,LARGE1,DOK7,TRAPPC9,SHANK3,NECTIN1,ODAD3,ACP5,PEX14,DYNC2I2,ARVCF,KATNIP,NALCN,CACNA1G,ANKH,EXT2,ERCC2,GNPTAB,SMARCC2,RSPH1
NR2F1,RBM20,OFD1,LARGE1,GATA3,DOK7,TRAPPC9,SHANK3,FXYD2,ACP5,MYO5A,AHDC1,PEX14,CHD5,MRPS2,GAD1,ARVCF,KATNIP,FZR1,COL4A2,NALCN,CACNA1G,ANKH,ETFB,EXT2,ETHE1,ERCC2,GNPTAB,SMARCC2
GRB10,OFD1,LARGE1,DOK7,SHANK3,NECTIN1,AHDC1,PEX14,CHD5,DYNC2I2,GAD1,ARVCF,NALCN,EXT2,ERCC2,GNPTAB
GRB10,OFD1,LARGE1,SHANK3,PEX14,DYNC2I2,GAD1,ANKH,ETFB,EXT2,CRADD
NR2F1,GRB10,OFD1,SETD1B,TGFB2,DOK7,TRAPPC9,SHANK3,NECTIN1,ODAD3,ACP5,DYNC2I2,ARVCF,KATNIP,NALCN,CACNA1G,EXT2,ERCC2,GNPTAB,SMARCC2,RSPH1
NR2F1,UROS,GRB10,RBM20,PAX2,OFD1,TGFB2,USH2A,DOK7,SHANK3,NECTIN1,ACP5,MYO5A,PEX14,MRPS2,TMPRSS6,DTNBP1,ARVCF,NALCN,NBEAL2,ETFB,EXT2,ETHE1,ERCC2,GNPTAB
MCPH1,OFD1,LARGE1,AHDC1,PEX14,KATNIP,COL4A2,ETFB,CRADD,SMARCC2
MCPH1,GRB10,OFD1,LARGE1,TRAPPC9,RNPC3,ACP5,PEX14,CHD5,DYNC2I2,KATNIP,FZR1,NALCN,ETFB,EXT2,ERCC2,GNPTAB,SMARCC2
NR2F1,MCPH1,OFD1,USH2A,LARGE1,DOK7,TRAPPC9,SHANK3,RNPC3,MYO5A,AHDC1,PEX14,ESPN,DYNC2I2,GAD1,ARVCF,KATNIP,FZR1,NALCN,CACNA1G,EXT2,ERCC2,GNPTAB,SMARCC2
NR2F1,OFD1,LARGE1,DOK7,TRAPPC9,SHANK3,NECTIN1,ACP5,AHDC1,PEX14,CHD5,MRPS2,GAD1,ARVCF,KATNIP,NALCN,CACNA1G,ETFB,ERCC2
NR2F1,RBM20,PAX2,OFD1,USH2A,GATA3,SHANK3,NECTIN1,ODAD3,AHDC1,PEX14,ESPN,MRPS2,ARVCF,NALCN,CACNA1G,ANKH,OTOG,ERCC2,GNPTAB,SMARCC2,RSPH1
UROS,IL6R,GRB10,PAX2,OFD1,USH2A,STOX1,GATA3,DOK7,TRAPPC9,SHANK3,ACP5,MRPS2,DYNC2I2,NALCN,ANKH,FLT1,ETFB,MAPK8IP1,ETHE1,ERCC2,GNPTAB
UROS,GRB10,RBM20,PAX2,OFD1,TGFB2,USH2A,STOX1,GATA3,SHANK3,NECTIN1,ODAD3,ACP5,MYO5A,PEX14,ESPN,DYNC2I2,DTNBP1,ARVCF,KATNIP,LAT,COL4A2,FLT1,NBEAL2,EXT2,ETHE1,ERCC2,GNPTAB,SMARCC2,RSPH1
UROS,IL6R,GATA3,SHANK3,ACP5,FOXN1,ARVCF,IL4R,CACNA1G,ERCC2,SMARCC2
GRB10,PAX2,OFD1,USH2A,LARGE1,STOX1,GATA3,DOK7,NECTIN1,PEX14,DYNC2I2,GAD1,ARVCF,KATNIP,NALCN,FLT1,EXT2,DNAH17,ERCC2,SMARCC2
IL6R,OFD1,NECTIN1,ODAD3,ACP5,ARVCF,IL21R,GNPTAB,RSPH1
