## Supplemental Table 4 for "Conserved DNA Methylation Signatures in The Prefrontal Cortex of Newborn and Juvenile Guinea Pigs Following Antenatal Corticosteroid Exposure"

**Supplementary Table 4. List of DMCs identified within the binding site for PLAGL1 in the PND1PFC following ACS exposure**

| seqnames | loc | MethDiff | Gene name |
|---|---|---|---|
| scaffold_0 | 3190833 | 14.9099 | |
| scaffold_0 | 3190835 | 14.5480 | |
| scaffold_0 | 4938109 | 19.7743 | |
| scaffold_0 | 81521644 | 12.5325 | |
| scaffold_0 | 81521703 | 10.3668 | |
| scaffold_0 | 81521735 | 25.9259 | |
| scaffold_0 | 81521750 | 6.7669 | |
| scaffold_0 | 81521768 | -8.5299 | |
| scaffold_1 | 12188172 | 14.4483 | NR2F1 |
| scaffold_1 | 12189570 | 30.0676 | NR2F1 |
| scaffold_1 | 12189573 | 25.7353 | NR2F1 |
| scaffold_1 | 12189574 | 28.8007 | NR2F1 |
| scaffold_1 | 12189576 | 18.1373 | NR2F1 |
| scaffold_1 | 12189615 | 20.5882 | NR2F1 |
| scaffold_1 | 22073022 | -9.9795 | |
| scaffold_1 | 22073209 | 43.3566 | |
| scaffold_1 | 22073210 | -7.0011 | |
| scaffold_1 | 67622611 | 14.9232 | MCPH1 |
| scaffold_102 | 2182048 | -11.9675 | |
| scaffold_102 | 2182054 | -15.4158 | |
| scaffold_102 | 2182163 | -5.4740 | |
| scaffold_102 | 2267910 | -5.4965 | GRB10 |
| scaffold_104 | 2571575 | 7.7947 | |
| scaffold_104 | 2571576 | 20.7071 | |
| scaffold_104 | 2960365 | 10.4615 | |
| scaffold_11 | 4675922 | -7.1429 | RBM20 |
| scaffold_11 | 13227517 | 22.0779 | |
| scaffold_11 | 13618122 | 7.3864 | PAX2 |
| scaffold_115 | 1405404 | 5.7143 | SETD1B |
| scaffold_115 | 1405949 | 14.7059 | SETD1B |
| scaffold_115 | 1405995 | -7.5803 | SETD1B |
| scaffold_115 | 1407188 | -21.4286 | SETD1B |
| scaffold_115 | 1407203 | -19.6581 | SETD1B |
| scaffold_115 | 1407612 | -5.5556 | SETD1B |
| scaffold_115 | 1407826 | -6.2222 | SETD1B |
| scaffold_117 | 3777546 | -12.3077 | UNC5D |
| scaffold_119 | 1850983 | 13.3333 | |
| scaffold_119 | 1850992 | 18.7500 | |
| scaffold_129 | 3116299 | 15.0742 | |
| scaffold_13 | 4860579 | -11.8926 | |
| scaffold_13 | 7208132 | -6.2290 | |
| scaffold_15 | 18434585 | 10.7710 | STOX1 |
| scaffold_15 | 18434586 | -9.6618 | STOX1 |
| scaffold_157 | 451189 | 6.6667 | |
| scaffold_157 | 451191 | 20.0000 | |
| scaffold_16 | 15313760 | 7.8938 | GATA3 |
| scaffold_166 | 1367121 | 16.0526 | |
| scaffold_17 | 1455212 | -5.4762 | |
| scaffold_17 | 37385648 | -8.7500 | TRAPPC9 |
| scaffold_17 | 37385681 | -15.0000 | TRAPPC9 |
| scaffold_171 | 219389 | 5.2880 | RFX4 |
| scaffold_171 | 219390 | 16.0606 | RFX4 |
| scaffold_171 | 220999 | 5.2797 | RFX4 |
| scaffold_171 | 724380 | 8.9286 | |
| scaffold_171 | 724392 | 5.2273 | |
| scaffold_176 | 237932 | 10.6195 | MGAT4B |

| | | | |
|---|---|---|---|
| scaffold_18 | 38154126 | -6.2500 | ;RNF208 |
| scaffold_18 | 38752726 | 5.7937 | |
| scaffold_184 | 170790 | 10.0000 | SHANK3 |
| scaffold_187 | 370889 | -9.8296 | |
| scaffold_19 | 14541155 | 8.7432 | NTM |
| scaffold_19 | 26005068 | 8.4615 | |
| scaffold_19 | 30245445 | -23.1951 | FXYD2 |
| scaffold_2 | 28509248 | -5.4054 | |
| scaffold_2 | 67316632 | 18.3333 | |
| scaffold_22 | 11218832 | -7.3152 | |
| scaffold_226 | 683617 | 7.3457 | ACP5 |
| scaffold_25 | 8761310 | -5.4144 | AHDC1 |
| scaffold_25 | 11570287 | -6.7382 | MYOM3 |
| scaffold_25 | 11570288 | 8.0000 | MYOM3 |
| scaffold_25 | 16996539 | 6.2304 | |
| scaffold_25 | 26209876 | -17.8874 | ESPN |
| scaffold_25 | 26209882 | -11.8179 | ESPN |
| scaffold_25 | 26211125 | 6.8760 | ESPN |
| scaffold_25 | 26211251 | 5.9723 | ESPN |
| scaffold_27 | 1730450 | 17.8571 | |
| scaffold_27 | 1730725 | 6.0821 | |
| scaffold_27 | 3631606 | 11.1345 | DYNC2I2 |
| scaffold_27 | 3631633 | 15.7143 | DYNC2I2 |
| scaffold_27 | 9691327 | -23.1250 | CNTFR |
| scaffold_27 | 9691381 | -21.5461 | CNTFR |
| scaffold_27 | 9691385 | -8.8816 | CNTFR |
| scaffold_27 | 14983904 | 9.6126 | CNTFR |
| scaffold_283 | 343141 | 9.9662 | STAB2 |
| scaffold_3 | 11303918 | -7.8283 | |
| scaffold_3 | 19719998 | -5.8824 | KLHL23 |
| scaffold_3 | 19757807 | 5.6152 | CFAP210 |
| scaffold_30 | 3585195 | -9.4886 | MEPCE |
| scaffold_31 | 9029914 | 23.2493 | |
| scaffold_31 | 12080653 | -5.9342 | |
| scaffold_31 | 14524353 | 33.2727 | |
| scaffold_32 | 65194 | -5.1630 | |
| scaffold_32 | 19461946 | -6.6828 | |
| scaffold_32 | 20114717 | -5.2817 | MNT |
| scaffold_32 | 20114786 | 5.3070 | MNT |
| scaffold_334 | 104291 | 15.4237 | |
| scaffold_356 | 220749 | -23.8095 | |
| scaffold_356 | 220753 | 14.2857 | |
| scaffold_356 | 245739 | 24.3056 | ARID3A |
| scaffold_356 | 245765 | 22.2222 | ARID3A |
| scaffold_38 | 537094 | 15.3846 | |
| scaffold_4 | 18837025 | -7.7395 | |
| scaffold_4 | 18837026 | 5.9048 | |
| scaffold_4 | 43658158 | -5.0132 | IL4R |
| scaffold_4 | 43658197 | -7.3413 | IL4R |
| scaffold_4 | 43662895 | 6.2008 | IL4R |
| scaffold_4 | 43665301 | 6.4950 | |
| scaffold_4 | 43668936 | -10.4167 | |
| scaffold_4 | 43669160 | 5.8442 | |
| scaffold_4 | 43669161 | 8.8991 | |
| scaffold_4 | 43671889 | -6.4342 | |
| scaffold_4 | 43672334 | 6.3425 | |
| scaffold_4 | 43672823 | -6.9818 | |
| scaffold_4 | 43672912 | -7.4522 | |

| | | | |
|---|---|---|---|
| scaffold_4 | 43674500 | -10.9337 | |
| scaffold_4 | 43674749 | -13.6907 | |
| scaffold_4 | 43679422 | -6.1170 | |
| scaffold_4 | 43679466 | 5.4657 | |
| scaffold_4 | 43679521 | -5.2952 | |
| scaffold_4 | 43701365 | 12.0346 | IL21R |
| scaffold_4 | 43704868 | -12.3684 | IL21R |
| scaffold_4 | 43706939 | 8.6486 | IL21R |
| scaffold_4 | 43706940 | -5.3571 | IL21R |
| scaffold_4 | 43707012 | 5.0595 | IL21R |
| scaffold_4 | 43707013 | 5.4736 | IL21R |
| scaffold_4 | 43717396 | -10.2222 | IL21R |
| scaffold_4 | 43726477 | 10.3393 | IL21R |
| scaffold_4 | 43730755 | 6.5055 | IL21R |
| scaffold_4 | 43735905 | -22.0374 | |
| scaffold_4 | 43737303 | 8.1731 | |
| scaffold_4 | 43743320 | 6.3141 | |
| scaffold_4 | 43743321 | -8.3333 | |
| scaffold_4 | 43745108 | 6.0606 | |
| scaffold_4 | 43745817 | -7.3333 | |
| scaffold_4 | 43983740 | 5.5732 | KATNIP |
| scaffold_4 | 43983993 | 5.3181 | KATNIP |
| scaffold_4 | 43993878 | -5.0792 | KATNIP |
| scaffold_4 | 43993879 | -7.8621 | KATNIP |
| scaffold_4 | 43993944 | 9.7565 | KATNIP |
| scaffold_4 | 43999584 | -8.6761 | KATNIP |
| scaffold_4 | 44005323 | 6.0606 | |
| scaffold_4 | 44005324 | 13.6797 | |
| scaffold_4 | 44007338 | -7.0238 | |
| scaffold_4 | 44010485 | 10.7143 | |
| scaffold_4 | 44010550 | 11.7949 | |
| scaffold_4 | 44018185 | -22.6190 | GSG1L |
| scaffold_4 | 44019014 | -16.5157 | GSG1L |
| scaffold_4 | 44029966 | -6.0759 | GSG1L |
| scaffold_4 | 44034087 | -5.8714 | GSG1L |
| scaffold_4 | 44059173 | 5.2843 | GSG1L |
| scaffold_4 | 44071986 | 6.8027 | GSG1L |
| scaffold_4 | 44082500 | -9.5238 | GSG1L |
| scaffold_4 | 44084051 | 7.1192 | GSG1L |
| scaffold_4 | 44090763 | 7.7139 | GSG1L |
| scaffold_4 | 44097076 | -8.2280 | GSG1L |
| scaffold_4 | 44101070 | 5.6899 | GSG1L |
| scaffold_4 | 44104139 | 12.3810 | GSG1L |
| scaffold_4 | 44116433 | -9.6555 | GSG1L |
| scaffold_4 | 44136160 | 5.6250 | GSG1L |
| scaffold_4 | 44139743 | 9.0332 | GSG1L |
| scaffold_4 | 44150629 | 17.6136 | GSG1L |
| scaffold_4 | 44152570 | -18.5484 | GSG1L |
| scaffold_4 | 44160686 | -7.3364 | GSG1L |
| scaffold_4 | 44162577 | -7.2654 | GSG1L |
| scaffold_4 | 44180856 | -13.3838 | GSG1L |
| scaffold_4 | 44189676 | -8.5885 | GSG1L |
| scaffold_4 | 44232790 | -6.2609 | GSG1L |
| scaffold_4 | 44234611 | -6.3542 | GSG1L |
| scaffold_4 | 44234613 | -10.2222 | GSG1L |
| scaffold_4 | 44236368 | 11.1111 | GSG1L |
| scaffold_4 | 44236942 | 8.9852 | GSG1L |
| scaffold_4 | 44236949 | 35.2632 | |

| | | | |
|---|---|---|---|
| scaffold_4 | 44236993 | 27.4283 | |
| scaffold_4 | 44243687 | -6.5752 | |
| scaffold_4 | 44243908 | 6.2849 | |
| scaffold_4 | 44373687 | -9.5238 | XPO6 |
| scaffold_4 | 44393466 | -20.1754 | XPO6 |
| scaffold_4 | 44467392 | 6.6667 | SBK1 |
| scaffold_4 | 44468547 | 8.3333 | SBK1 |
| scaffold_4 | 44488438 | -6.6935 | SBK1 |
| scaffold_4 | 44488558 | -7.0115 | SBK1 |
| scaffold_4 | 44488674 | -10.5806 | SBK1 |
| scaffold_4 | 44488684 | -21.7936 | SBK1 |
| scaffold_4 | 44488704 | -8.0543 | SBK1 |
| scaffold_4 | 44488709 | -10.3167 | SBK1 |
| scaffold_4 | 44488847 | 6.8403 | SBK1 |
| scaffold_4 | 44488871 | -5.5417 | SBK1 |
| scaffold_4 | 44488878 | 5.0735 | SBK1 |
| scaffold_4 | 44488891 | 16.6667 | SBK1 |
| scaffold_4 | 44488892 | 7.0075 | SBK1 |
| scaffold_4 | 44545255 | 7.6923 | |
| scaffold_4 | 44555526 | 10.4895 | |
| scaffold_4 | 44555528 | 5.7829 | |
| scaffold_4 | 44555529 | -6.2937 | |
| scaffold_4 | 44567091 | -9.0395 | |
| scaffold_4 | 44578994 | -13.0390 | |
| scaffold_4 | 44591337 | 5.1948 | SPNS1 |
| scaffold_4 | 44591340 | -9.0909 | SPNS1 |
| scaffold_406 | 152282 | 5.1411 | FZR1 |
| scaffold_41 | 8487166 | 6.5104 | IPCEF1 |
| scaffold_48 | 12757823 | 6.1309 | CCSER1 |
| scaffold_48 | 12882385 | -5.9524 | |
| scaffold_5 | 5566435 | -7.1778 | |
| scaffold_5 | 14328260 | -25.0000 | DCUN1D2 |
| scaffold_5 | 14352160 | 5.1645 | |
| scaffold_5 | 23258443 | 20.9514 | NALCN |
| scaffold_5 | 23258488 | 18.7755 | NALCN |
| scaffold_53 | 4107592 | -8.0128 | |
| scaffold_53 | 4107601 | -15.3846 | |
| scaffold_53 | 4107615 | -11.1378 | |
| scaffold_53 | 4161209 | -19.5238 | |
| scaffold_55 | 4084534 | 8.5859 | |
| scaffold_6 | 6394780 | -5.2083 | |
| scaffold_6 | 28209159 | 15.5012 | |
| scaffold_61 | 7389793 | 16.0037 | ARHGEF15 |
| scaffold_61 | 7390058 | -11.6654 | ARHGEF15 |
| scaffold_62 | 5317458 | 8.7124 | |
| scaffold_62 | 5322066 | -8.5714 | |
| scaffold_62 | 5322071 | -10.0000 | |
| scaffold_64 | 309515 | 5.9297 | |
| scaffold_67 | 9123772 | -5.4579 | ADAM11 |
| scaffold_67 | 9123900 | 5.4643 | ADAM11 |
| scaffold_7 | 46246098 | -6.8182 | TGM4 |
| scaffold_7 | 51192435 | 6.9749 | ETV5 |
| scaffold_70 | 1150925 | -5.9006 | |
| scaffold_70 | 7338737 | -9.5640 | MYBPC2 |
| scaffold_72 | 3962647 | -8.5442 | |
| scaffold_72 | 5300917 | -6.9226 | |
| scaffold_72 | 8384344 | 8.6275 | ARMC6 |
| scaffold_76 | 273296 | 9.0158 | OTOP2 |

| | | | |
|---|---|---|---|
| scaffold_76 | 2517037 | 16.6667 | |
| scaffold_80 | 6670981 | -11.4679 | GNG8 |
| scaffold_88 | 6464009 | -13.7436 | GLIS1 |
| scaffold_9 | 8095993 | 14.0993 | |
| scaffold_90 | 2643210 | -8.1681 | RSPH1 |
| scaffold_94 | 5701288 | 7.8466 | |
| scaffold_96 | 319730 | 7.1129 | |
| scaffold_96 | 5435934 | 9.0671 | |
