## Supplemental Table 5 for "Conserved DNA Methylation Signatures in The Prefrontal Cortex of Newborn and Juvenile Guinea Pigs Following Antenatal Corticosteroid Exposure"

**Supplementary Table 5. List of DMCs identified within the binding site for TFAP2C in the PND1PFC following ACS exposure**

| seqnames | loc | MethDiff | Gene name |
|---|---|---|---|
| scaffold_0 | 4938208 | 8.9113 | |
| scaffold_0 | 81521658 | 9.6180 | |
| scaffold_0 | 81521679 | 14.7116 | |
| scaffold_0 | 81521686 | 13.3857 | |
| scaffold_0 | 81521694 | 11.8648 | |
| scaffold_0 | 81521703 | 10.3668 | |
| scaffold_0 | 81521735 | 25.9259 | |
| scaffold_0 | 81521750 | 6.7669 | |
| scaffold_0 | 81521759 | -8.3485 | |
| scaffold_1 | 12189503 | 26.1029 | NR2F1 |
| scaffold_1 | 12189506 | 19.4989 | NR2F1 |
| scaffold_1 | 12189512 | 20.2381 | NR2F1 |
| scaffold_1 | 12189515 | 28.1746 | NR2F1 |
| scaffold_1 | 12189518 | 13.1250 | NR2F1 |
| scaffold_1 | 12189521 | 36.3636 | NR2F1 |
| scaffold_1 | 12189524 | 35.8586 | NR2F1 |
| scaffold_1 | 12189529 | 19.1919 | NR2F1 |
| scaffold_1 | 12189558 | 24.4881 | NR2F1 |
| scaffold_1 | 12189570 | 30.0676 | NR2F1 |
| scaffold_1 | 12189615 | 20.5882 | NR2F1 |
| scaffold_1 | 22073022 | -9.9795 | |
| scaffold_1 | 22073043 | -13.6364 | |
| scaffold_1 | 22073209 | 43.3566 | |
| scaffold_1 | 22073210 | -7.0011 | |
| scaffold_1 | 67622582 | 10.7028 | MCPH1 |
| scaffold_1 | 67622586 | 18.7042 | MCPH1 |
| scaffold_1 | 67622588 | 12.7957 | MCPH1 |
| scaffold_10 | 15327363 | 13.0848 | IL6R |
| scaffold_10 | 15327371 | 14.2178 | IL6R |
| scaffold_10 | 15327420 | 12.7178 | IL6R |
| scaffold_10 | 32971348 | -14.0687 | |
| scaffold_10 | 45822346 | -18.8235 | |
| scaffold_10 | 47593463 | -18.8811 | TARS3 |
| scaffold_102 | 2182012 | 45.0000 | n/a |
| scaffold_102 | 2182014 | 42.7273 | |
| scaffold_102 | 2182021 | -13.3333 | |
| scaffold_102 | 2182022 | 45.0000 | |
| scaffold_102 | 2182027 | -15.4158 | |
| scaffold_102 | 2182029 | -15.4158 | |
| scaffold_102 | 2182044 | -21.2982 | |
| scaffold_102 | 2182048 | -11.9675 | |
| scaffold_102 | 2182054 | -15.4158 | |
| scaffold_102 | 2182163 | -5.4740 | |
| scaffold_104 | 2571528 | 13.6364 | |
| scaffold_104 | 2571678 | 18.7500 | |
| scaffold_108 | 1187941 | -9.0972 | |
| scaffold_108 | 1187950 | 12.6100 | |
| scaffold_11 | 13617836 | 23.7925 | PAX2 |
| scaffold_111 | 12656 | -6.6667 | |
| scaffold_111 | 1782351 | 11.4695 | |
| scaffold_113 | 3940974 | -6.0212 | OFD1 |
| scaffold_113 | 3940975 | -8.0217 | OFD1 |
| scaffold_115 | 1405377 | 8.1081 | SETD1B |
| scaffold_115 | 1405400 | 5.7143 | SETD1B |
| scaffold_115 | 1405404 | 5.7143 | SETD1B |
| scaffold_115 | 1405949 | 14.7059 | SETD1B |

| | | | |
|---|---|---|---|
| scaffold_115 | 1405987 | 5.8986 | SETD1B |
| scaffold_115 | 1405995 | -7.5803 | SETD1B |
| scaffold_115 | 1407203 | -19.6581 | SETD1B |
| scaffold_115 | 1407612 | -5.5556 | SETD1B |
| scaffold_115 | 1407650 | 12.7193 | SETD1B |
| scaffold_119 | 1850983 | 13.3333 | |
| scaffold_119 | 1850992 | 18.7500 | |
| scaffold_125 | 3044950 | -5.3555 | C2orf50 |
| scaffold_13 | 4860603 | -26.9565 | |
| scaffold_13 | 7208132 | -6.2290 | |
| scaffold_140 | 985308 | -20.3488 | LARGE1 |
| scaffold_15 | 29335091 | 23.7132 | |
| scaffold_15 | 33165439 | -7.6923 | |
| scaffold_157 | 448186 | -6.8744 | n/a |
| scaffold_157 | 451189 | 6.6667 | |
| scaffold_157 | 451191 | 20.0000 | |
| scaffold_158 | 1702529 | 5.8764 | |
| scaffold_158 | 1702678 | 5.9524 | |
| scaffold_16 | 15313760 | 7.8938 | GATA3 |
| scaffold_166 | 1367121 | 16.0526 | |
| scaffold_167 | 215537 | -7.1429 | |
| scaffold_17 | 1455212 | -5.4762 | |
| scaffold_17 | 37385675 | -18.7500 | TRAPPC9 |
| scaffold_17 | 37385681 | -15.0000 | TRAPPC9 |
| scaffold_17 | 37385703 | -8.7500 | TRAPPC9 |
| scaffold_17 | 37385710 | -8.7500 | TRAPPC9 |
| scaffold_171 | 724380 | 8.9286 | |
| scaffold_171 | 724441 | 8.2264 | |
| scaffold_184 | 170790 | 10.0000 | SHANK3 |
| scaffold_187 | 370889 | -9.8296 | |
| scaffold_19 | 30245445 | -23.1951 | FXYD2 |
| scaffold_2 | 28509248 | -5.4054 | |
| scaffold_2 | 28541024 | -8.2050 | |
| scaffold_20 | 14157758 | 6.2667 | CCDC177 |
| scaffold_20 | 14157774 | -7.2196 | CCDC177 |
| scaffold_20 | 14157778 | 5.0451 | CCDC177 |
| scaffold_20 | 14157813 | -25.9740 | CCDC177 |
| scaffold_2035 | 5181 | -5.5080 | |
| scaffold_226 | 527620 | 10.5391 | ODAD3 |
| scaffold_226 | 683617 | 7.3457 | ACP5 |
| scaffold_23 | 25112886 | -5.6843 | |
| scaffold_23 | 29612465 | -6.4219 | LEO1 |
| scaffold_24 | 1230273 | -5.9091 | MGAT4A |
| scaffold_24 | 1230301 | 8.6998 | MGAT4A |
| scaffold_25 | 8761470 | 10.8615 | AHDC1 |
| scaffold_25 | 16670998 | -31.4286 | IGSF21 |
| scaffold_25 | 17887025 | -8.3333 | EPHA2 |
| scaffold_25 | 17963135 | -5.0000 | |
| scaffold_25 | 26209843 | -16.2448 | ESPN |
| scaffold_25 | 26211125 | 6.8760 | ESPN |
| scaffold_25 | 27347393 | 5.1136 | AJAP1 |
| scaffold_255 | 162093 | -5.1982 | |
| scaffold_255 | 162147 | 7.9741 | |
| scaffold_27 | 569835 | 8.4572 | MRPS2 |
| scaffold_27 | 1730450 | 17.8571 | |
| scaffold_27 | 3631606 | 11.1345 | DYNC2I2 |
| scaffold_27 | 3631614 | 16.4286 | DYNC2I2 |
| scaffold_27 | 9691097 | -14.7619 | NEK6 |

| | | | |
|---|---|---|---|
| scaffold_27 | 9691327 | -23.1250 | CNTFR |
| scaffold_27 | 14983883 | 11.6930 | CNTFR |
| scaffold_28 | 21679569 | -11.4490 | TMPRSS6 |
| scaffold_28 | 21891446 | -8.6957 | |
| scaffold_28 | 22198197 | 8.3333 | |
| scaffold_28 | 27357739 | 6.0520 | EFCAB6 |
| scaffold_3 | 18813662 | -11.8644 | GAD1 |
| scaffold_3 | 18813682 | -6.2724 | GAD1 |
| scaffold_3 | 19719998 | -5.8824 | KLHL23 |
| scaffold_3 | 19746232 | -5.8824 | CFAP210 |
| scaffold_3 | 19746514 | -9.5238 | CFAP210 |
| scaffold_3 | 19746564 | -19.0476 | CFAP210 |
| scaffold_3 | 19757807 | 5.6152 | CFAP210 |
| scaffold_3 | 51971108 | 10.4441 | ACMSD |
| scaffold_31 | 12137429 | -5.6410 | |
| scaffold_31 | 14524323 | 10.9848 | |
| scaffold_31 | 14524353 | 33.2727 | |
| scaffold_31 | 14581579 | 11.4865 | |
| scaffold_31 | 14581588 | 12.0946 | |
| scaffold_32 | 19461802 | 5.0362 | |
| scaffold_32 | 20114745 | -6.3648 | MNT |
| scaffold_32 | 20114786 | 5.3070 | MNT |
| scaffold_32 | 20812739 | 7.4941 | SLC43A2 |
| scaffold_334 | 104291 | 15.4237 | |
| scaffold_337 | 18200 | -6.8831 | |
| scaffold_356 | 220749 | -23.8095 | |
| scaffold_356 | 220753 | 14.2857 | |
| scaffold_356 | 220802 | 26.9872 | |
| scaffold_356 | 245739 | 24.3056 | ARID3A |
| scaffold_38 | 537094 | 15.3846 | |
| scaffold_38 | 6902950 | -5.6892 | |
| scaffold_38 | 15092361 | 9.5652 | MBOAT1 |
| scaffold_4 | 19064543 | 5.0404 | ARVCF |
| scaffold_4 | 43660358 | 11.9360 | IL4R |
| scaffold_4 | 43665301 | 6.4950 | |
| scaffold_4 | 43668936 | -10.4167 | |
| scaffold_4 | 43669161 | 8.8991 | |
| scaffold_4 | 43671889 | -6.4342 | |
| scaffold_4 | 43671946 | 7.1819 | |
| scaffold_4 | 43671947 | -6.0859 | |
| scaffold_4 | 43672657 | 15.1947 | |
| scaffold_4 | 43672823 | -6.9818 | |
| scaffold_4 | 43672909 | -9.3703 | |
| scaffold_4 | 43672912 | -7.4522 | |
| scaffold_4 | 43674500 | -10.9337 | |
| scaffold_4 | 43674749 | -13.6907 | |
| scaffold_4 | 43679422 | -6.1170 | |
| scaffold_4 | 43679466 | 5.4657 | |
| scaffold_4 | 43704868 | -12.3684 | IL21R |
| scaffold_4 | 43706939 | 8.6486 | IL21R |
| scaffold_4 | 43706940 | -5.3571 | IL21R |
| scaffold_4 | 43707263 | 11.4995 | IL21R |
| scaffold_4 | 43717396 | -10.2222 | IL21R |
| scaffold_4 | 43717444 | 9.3968 | IL21R |
| scaffold_4 | 43726477 | 10.3393 | IL21R |
| scaffold_4 | 43730755 | 6.5055 | IL21R |
| scaffold_4 | 43731167 | 7.0376 | IL21R |
| scaffold_4 | 43731452 | -10.0684 | IL21R |

| | | | |
|---|---|---|---|
| scaffold_4 | 43732455 | -10.0000 | IL21R |
| scaffold_4 | 43737303 | 8.1731 | |
| scaffold_4 | 43738424 | -10.1307 | |
| scaffold_4 | 43743321 | -8.3333 | |
| scaffold_4 | 43745817 | -7.3333 | |
| scaffold_4 | 43967152 | -5.0311 | KATNIP |
| scaffold_4 | 43979316 | -9.3137 | KATNIP |
| scaffold_4 | 43983740 | 5.5732 | KATNIP |
| scaffold_4 | 43983903 | -5.4622 | KATNIP |
| scaffold_4 | 43987132 | 6.4016 | KATNIP |
| scaffold_4 | 43987304 | 11.7647 | KATNIP |
| scaffold_4 | 43993940 | -8.3673 | KATNIP |
| scaffold_4 | 43993944 | 9.7565 | KATNIP |
| scaffold_4 | 43994250 | 17.4603 | KATNIP |
| scaffold_4 | 43994257 | 15.0794 | KATNIP |
| scaffold_4 | 44005260 | 10.0652 | |
| scaffold_4 | 44010485 | 10.7143 | |
| scaffold_4 | 44014943 | 7.0659 | GSG1L |
| scaffold_4 | 44029760 | -10.2883 | GSG1L |
| scaffold_4 | 44059154 | 9.7877 | GSG1L |
| scaffold_4 | 44059341 | 6.6667 | GSG1L |
| scaffold_4 | 44060216 | 9.2593 | GSG1L |
| scaffold_4 | 44066550 | 7.2398 | GSG1L |
| scaffold_4 | 44084051 | 7.1192 | GSG1L |
| scaffold_4 | 44097076 | -8.2280 | GSG1L |
| scaffold_4 | 44100965 | 6.5583 | GSG1L |
| scaffold_4 | 44101126 | 5.9373 | GSG1L |
| scaffold_4 | 44105493 | -8.5788 | GSG1L |
| scaffold_4 | 44116433 | -9.6555 | GSG1L |
| scaffold_4 | 44150629 | 17.6136 | GSG1L |
| scaffold_4 | 44152570 | -18.5484 | GSG1L |
| scaffold_4 | 44160686 | -7.3364 | GSG1L |
| scaffold_4 | 44162577 | -7.2654 | GSG1L |
| scaffold_4 | 44169360 | 5.0000 | GSG1L |
| scaffold_4 | 44170676 | 7.4220 | GSG1L |
| scaffold_4 | 44175593 | 5.0699 | GSG1L |
| scaffold_4 | 44180856 | -13.3838 | GSG1L |
| scaffold_4 | 44189676 | -8.5885 | GSG1L |
| scaffold_4 | 44202637 | -16.9848 | GSG1L |
| scaffold_4 | 44204702 | -11.1111 | GSG1L |
| scaffold_4 | 44213812 | 5.0753 | GSG1L |
| scaffold_4 | 44213813 | 5.9834 | GSG1L |
| scaffold_4 | 44214795 | 10.1754 | GSG1L |
| scaffold_4 | 44217296 | 20.0000 | GSG1L |
| scaffold_4 | 44232790 | -6.2609 | GSG1L |
| scaffold_4 | 44234611 | -6.3542 | GSG1L |
| scaffold_4 | 44234613 | -10.2222 | GSG1L |
| scaffold_4 | 44236368 | 11.1111 | GSG1L |
| scaffold_4 | 44236410 | 5.5556 | GSG1L |
| scaffold_4 | 44294777 | 6.6667 | XPO6 |
| scaffold_4 | 44373678 | -7.4286 | XPO6 |
| scaffold_4 | 44393466 | -20.1754 | XPO6 |
| scaffold_4 | 44467392 | 6.6667 | SBK1 |
| scaffold_4 | 44468547 | 8.3333 | SBK1 |
| scaffold_4 | 44488558 | -7.0115 | SBK1 |
| scaffold_4 | 44488592 | -16.5528 | SBK1 |
| scaffold_4 | 44488610 | -12.5000 | SBK1 |
| scaffold_4 | 44488684 | -21.7936 | SBK1 |

| | | | |
|---|---|---|---|
| scaffold_4 | 44488689 | -7.4883 | SBK1 |
| scaffold_4 | 44488871 | -5.5417 | SBK1 |
| scaffold_4 | 44488878 | 5.0735 | SBK1 |
| scaffold_4 | 44488891 | 16.6667 | SBK1 |
| scaffold_4 | 44488892 | 7.0075 | SBK1 |
| scaffold_4 | 44488920 | 6.0012 | SBK1 |
| scaffold_4 | 44544803 | -36.8421 | |
| scaffold_4 | 44567125 | -5.0847 | |
| scaffold_4 | 44567436 | -18.4211 | |
| scaffold_4 | 44568513 | -13.4424 | |
| scaffold_4 | 44573656 | 10.9524 | |
| scaffold_4 | 44579109 | 10.5000 | |
| scaffold_4 | 44579904 | 9.7580 | |
| scaffold_4 | 44591337 | 5.1948 | SPNS1 |
| scaffold_4 | 44591340 | -9.0909 | SPNS1 |
| scaffold_406 | 152282 | 5.1411 | FZR1 |
| scaffold_47 | 14121981 | 16.3636 | |
| scaffold_48 | 12882385 | -5.9524 | |
| scaffold_49 | 1939847 | -6.9876 | |
| scaffold_5 | 5566435 | -7.1778 | |
| scaffold_5 | 14352160 | 5.1645 | |
| scaffold_5 | 15845441 | 7.5839 | COL4A2 |
| scaffold_5 | 16388591 | -10.8333 | |
| scaffold_5 | 23258443 | 20.9514 | NALCN |
| scaffold_5 | 23258486 | 23.6735 | NALCN |
| scaffold_5 | 23258488 | 18.7755 | NALCN |
| scaffold_523 | 68004 | 7.6923 | |
| scaffold_53 | 4104365 | 14.8148 | |
| scaffold_53 | 4105401 | -7.6522 | |
| scaffold_53 | 4106036 | -9.9548 | |
| scaffold_53 | 4107615 | -11.1378 | |
| scaffold_53 | 4214524 | -10.1810 | |
| scaffold_53 | 4276554 | -12.5000 | |
| scaffold_53 | 4276574 | -6.2500 | |
| scaffold_53 | 5044306 | -5.8824 | |
| scaffold_53 | 5044312 | 18.1818 | |
| scaffold_53 | 5044314 | -23.5294 | |
| scaffold_54 | 7899360 | -5.9915 | |
| scaffold_55 | 4084534 | 8.5859 | |
| scaffold_6 | 477488 | -8.9314 | |
| scaffold_6 | 6394780 | -5.2083 | |
| scaffold_6 | 28209159 | 15.5012 | |
| scaffold_6 | 28797664 | 5.4855 | |
| scaffold_6 | 28797914 | 9.6493 | |
| scaffold_61 | 7389429 | -9.5694 | ARHGEF15 |
| scaffold_61 | 7389469 | 16.6667 | ARHGEF15 |
| scaffold_61 | 7389733 | -13.8340 | ARHGEF15 |
| scaffold_62 | 5321709 | 7.2801 | |
| scaffold_62 | 5322008 | 19.0476 | |
| scaffold_625 | 42318 | -16.0131 | |
| scaffold_625 | 52492 | -11.6256 | |
| scaffold_67 | 9123772 | -5.4579 | ADAM11 |
| scaffold_67 | 9123856 | 5.5026 | ADAM11 |
| scaffold_67 | 9123900 | 5.4643 | ADAM11 |
| scaffold_7 | 51192435 | 6.9749 | ETV5 |
| scaffold_70 | 6539703 | 5.6991 | |
| scaffold_70 | 7338737 | -9.5640 | MYBPC2 |
| scaffold_72 | 4167957 | -10.5882 | EXT2 |

| | | | |
|---|---|---|---|
| scaffold_72 | 5300917 | -6.9226 | |
| scaffold_72 | 5426941 | -6.9444 | MAPK8IP1 |
| scaffold_72 | 8384344 | 8.6275 | ARMC6 |
| scaffold_72 | 8384480 | -8.9744 | ARMC6 |
| scaffold_75 | 5612160 | 5.0398 | ALLC |
| scaffold_76 | 2517037 | 16.6667 | |
| scaffold_8 | 53942159 | 11.5217 | |
| scaffold_8 | 53942208 | 7.1841 | |
| scaffold_8 | 53942209 | -8.5771 | |
| scaffold_80 | 5842187 | -8.6353 | KLC3;ERCC2 |
| scaffold_80 | 6670981 | -11.4679 | GNG8 |
| scaffold_80 | 6671077 | -5.8995 | GNG8 |
| scaffold_81 | 407584 | -10.5714 | ANXA6 |
| scaffold_81 | 407585 | -13.4625 | ANXA6 |
| scaffold_88 | 6464009 | -13.7436 | GLIS1 |
| scaffold_88 | 6464021 | -12.6112 | GLIS1 |
| scaffold_9 | 8095993 | 14.0993 | |
| scaffold_9 | 8095997 | 9.1232 | |
| scaffold_9 | 8096007 | 11.5174 | |
| scaffold_9 | 47500249 | -5.3108 | |
| scaffold_9 | 47511089 | 6.0606 | |
| scaffold_94 | 5701288 | 7.8466 | |
| scaffold_94 | 5701289 | 14.5032 | |
