## Supplemental Table 6 for "Conserved DNA Methylation Signatures in The Prefrontal Cortex of Newborn and Juvenile Guinea Pigs Following Antenatal Corticosteroid Exposure"

**Supplementary Table 6. List of DMCs identified within the binding site for EGR1 in the PND1PFC following ACS exposure**

| seqnames | start | MethDiff | Gene name |
|---|---|---|---|
| scaffold_0 | 62694884 | -54.8485 | PRICKLE2 |
| scaffold_0 | 81521604 | 6.4516 | |
| scaffold_0 | 81521644 | 12.5325 | |
| scaffold_0 | 81521703 | 10.3668 | |
| scaffold_0 | 81521735 | 25.9259 | |
| scaffold_0 | 81521743 | 7.2595 | |
| scaffold_0 | 81521768 | -8.5299 | |
| scaffold_0 | 81521795 | 5.2632 | |
| scaffold_0 | 81521799 | -15.4265 | |
| scaffold_1 | 12188172 | 14.4483 | NR2F1 |
| scaffold_1 | 12189503 | 26.1029 | NR2F1 |
| scaffold_1 | 12189506 | 19.4989 | NR2F1 |
| scaffold_1 | 12189512 | 20.2381 | NR2F1 |
| scaffold_1 | 12189515 | 28.1746 | NR2F1 |
| scaffold_1 | 12189518 | 13.1250 | NR2F1 |
| scaffold_1 | 12189521 | 36.3636 | NR2F1 |
| scaffold_1 | 12189524 | 35.8586 | NR2F1 |
| scaffold_1 | 12189615 | 20.5882 | NR2F1 |
| scaffold_1 | 73765934 | 9.7214 | |
| scaffold_1 | 73765935 | 12.1448 | |
| scaffold_10 | 35166327 | 50.8772 | |
| scaffold_10 | 45822409 | -17.6923 | |
| scaffold_102 | 2182021 | -13.3333 | |
| scaffold_102 | 2182022 | 45.0000 | |
| scaffold_102 | 2182027 | -15.4158 | |
| scaffold_102 | 2182029 | -15.4158 | |
| scaffold_102 | 2182033 | -15.4158 | |
| scaffold_102 | 2182048 | -11.9675 | |
| scaffold_102 | 2182054 | -15.4158 | |
| scaffold_104 | 2571706 | -13.3333 | |
| scaffold_104 | 2571714 | 29.6970 | |
| scaffold_108 | 1186158 | -26.9231 | |
| scaffold_11 | 13618122 | 7.3864 | PAX2 |
| scaffold_11 | 16436183 | -16.7033 | |
| scaffold_111 | 1781402 | -5.6075 | |
| scaffold_111 | 3648074 | -6.5027 | |
| scaffold_115 | 1405669 | 12.5000 | SETD1B |
| scaffold_115 | 1407472 | 7.1429 | SETD1B |
| scaffold_115 | 1407475 | 5.8824 | SETD1B |
| scaffold_115 | 1407569 | -6.5217 | SETD1B |
| scaffold_115 | 1407725 | 5.6907 | SETD1B |
| scaffold_115 | 4048902 | -5.4612 | |
| scaffold_119 | 1850983 | 13.3333 | |
| scaffold_119 | 1850992 | 18.7500 | |
| scaffold_125 | 3044950 | -5.3555 | C2orf50 |
| scaffold_13 | 3910516 | 10.2504 | TGFB2 |
| scaffold_13 | 4860555 | -7.0807 | |
| scaffold_13 | 4860556 | 6.5452 | |
| scaffold_13 | 6041158 | 6.4604 | USH2A |
| scaffold_13 | 7208132 | -6.2290 | |
| scaffold_140 | 985245 | -21.1310 | LARGE1 |
| scaffold_15 | 29335091 | 23.7132 | |
| scaffold_15 | 40638129 | -24.5192 | |
| scaffold_158 | 1702678 | 5.9524 | |
| scaffold_166 | 517969 | 21.3115 | SORCS2 |
| scaffold_166 | 1366925 | 12.7976 | |

| | | | |
|---|---|---|---|
| scaffold_166 | 1366929 | 5.2381 | |
| scaffold_166 | 1397405 | -23.2456 | DOK7 |
| scaffold_168 | 1567710 | 5.0000 | AKAP8L |
| scaffold_168 | 1569602 | 5.5618 | AKAP8L |
| scaffold_17 | 37385675 | -18.7500 | TRAPPC9 |
| scaffold_17 | 37385681 | -15.0000 | TRAPPC9 |
| scaffold_17 | 37385710 | -8.7500 | TRAPPC9 |
| scaffold_17 | 37385713 | 9.0543 | TRAPPC9 |
| scaffold_17 | 37385717 | 9.7990 | TRAPPC9 |
| scaffold_184 | 170790 | 10.0000 | SHANK3 |
| scaffold_186 | 315322 | 10.8065 | |
| scaffold_186 | 315323 | -5.1760 | |
| scaffold_186 | 315326 | -8.8710 | |
| scaffold_19 | 14541155 | 8.7432 | NTM |
| scaffold_19 | 30245445 | -23.1951 | FXYD2 |
| scaffold_20 | 14157758 | 6.2667 | CCDC177 |
| scaffold_2035 | 5181 | -5.5080 | |
| scaffold_21 | 31749772 | 6.1832 | |
| scaffold_23 | 25112886 | -5.6843 | |
| scaffold_2495 | 6354 | -18.3871 | |
| scaffold_2495 | 6356 | -19.3676 | |
| scaffold_25 | 11383 | -5.9163 | |
| scaffold_25 | 11570287 | -6.7382 | MYOM3 |
| scaffold_25 | 11570288 | 8.0000 | MYOM3 |
| scaffold_25 | 17887006 | 5.0000 | EPHA2 |
| scaffold_25 | 23409107 | 6.5657 | PEX14 |
| scaffold_25 | 26209855 | -5.4422 | ESPN |
| scaffold_25 | 26209856 | -12.0623 | ESPN |
| scaffold_25 | 26209860 | -12.0076 | ESPN |
| scaffold_25 | 26209876 | -17.8874 | ESPN |
| scaffold_25 | 26209882 | -11.8179 | ESPN |
| scaffold_255 | 162147 | 7.9741 | |
| scaffold_27 | 569835 | 8.4572 | MRPS2 |
| scaffold_27 | 1293083 | -12.5125 | CFAP77 |
| scaffold_27 | 1730781 | -8.8580 | |
| scaffold_27 | 3631606 | 11.1345 | DYNC2I2 |
| scaffold_27 | 9691327 | -23.1250 | CNTFR |
| scaffold_27 | 9691381 | -21.5461 | CNTFR |
| scaffold_27 | 9691385 | -8.8816 | CNTFR |
| scaffold_27 | 14983883 | 11.6930 | CNTFR |
| scaffold_28 | 21679569 | -11.4490 | TMPRSS6 |
| scaffold_28 | 21891185 | 5.2632 | |
| scaffold_28 | 21891446 | -8.6957 | |
| scaffold_3 | 18813682 | -6.2724 | GAD1 |
| scaffold_3 | 19719998 | -5.8824 | KLHL23 |
| scaffold_31 | 5384849 | 30.0395 | |
| scaffold_31 | 14581588 | 12.0946 | |
| scaffold_31 | 14591436 | -10.0000 | |
| scaffold_32 | 20114745 | -6.3648 | MNT |
| scaffold_32 | 23946888 | 18.1582 | FOXN1 |
| scaffold_32 | 23947632 | 6.0269 | FOXN1 |
| scaffold_33 | 24466781 | 12.6812 | |
| scaffold_334 | 104291 | 15.4237 | |
| scaffold_334 | 104298 | 6.0725 | |
| scaffold_356 | 220802 | 26.9872 | |
| scaffold_356 | 245739 | 24.3056 | ARID3A |
| scaffold_368 | 224088 | 8.5017 | TYK2 |
| scaffold_38 | 537094 | 15.3846 | |

| | | | |
|---|---|---|---|
| scaffold_4 | 18836653 | 20.9957 | |
| scaffold_4 | 18836654 | -22.5287 | |
| scaffold_4 | 19064543 | 5.0404 | ARVCF |
| scaffold_4 | 43658158 | -5.0132 | IL4R |
| scaffold_4 | 43658576 | 8.0000 | IL4R |
| scaffold_4 | 43665148 | 10.4356 | |
| scaffold_4 | 43665301 | 6.4950 | |
| scaffold_4 | 43671870 | 7.1161 | |
| scaffold_4 | 43672763 | -14.5888 | |
| scaffold_4 | 43672769 | -9.4304 | |
| scaffold_4 | 43672808 | -12.6429 | |
| scaffold_4 | 43672810 | -9.0929 | |
| scaffold_4 | 43672842 | 10.1120 | |
| scaffold_4 | 43672843 | -7.0549 | |
| scaffold_4 | 43672909 | -9.3703 | |
| scaffold_4 | 43672912 | -7.4522 | |
| scaffold_4 | 43674500 | -10.9337 | |
| scaffold_4 | 43679466 | 5.4657 | |
| scaffold_4 | 43679512 | -12.6529 | |
| scaffold_4 | 43701365 | 12.0346 | IL21R |
| scaffold_4 | 43706939 | 8.6486 | IL21R |
| scaffold_4 | 43706940 | -5.3571 | IL21R |
| scaffold_4 | 43726477 | 10.3393 | IL21R |
| scaffold_4 | 43735905 | -22.0374 | |
| scaffold_4 | 43735912 | 6.9993 | |
| scaffold_4 | 43738424 | -10.1307 | |
| scaffold_4 | 43743320 | 6.3141 | |
| scaffold_4 | 43743321 | -8.3333 | |
| scaffold_4 | 43745817 | -7.3333 | |
| scaffold_4 | 43976249 | 7.2896 | KATNIP |
| scaffold_4 | 43979316 | -9.3137 | KATNIP |
| scaffold_4 | 43983740 | 5.5732 | KATNIP |
| scaffold_4 | 43983993 | 5.3181 | KATNIP |
| scaffold_4 | 43993691 | 5.9066 | KATNIP |
| scaffold_4 | 43993838 | -6.1560 | KATNIP |
| scaffold_4 | 44000886 | -8.3333 | KATNIP |
| scaffold_4 | 44005323 | 6.0606 | |
| scaffold_4 | 44005324 | 13.6797 | |
| scaffold_4 | 44007259 | 5.8015 | |
| scaffold_4 | 44010550 | 11.7949 | |
| scaffold_4 | 44016815 | -6.1151 | GSG1L |
| scaffold_4 | 44018489 | -9.7619 | GSG1L |
| scaffold_4 | 44019014 | -16.5157 | GSG1L |
| scaffold_4 | 44026732 | 11.1111 | GSG1L |
| scaffold_4 | 44029905 | -7.7659 | GSG1L |
| scaffold_4 | 44029966 | -6.0759 | GSG1L |
| scaffold_4 | 44060216 | 9.2593 | GSG1L |
| scaffold_4 | 44061694 | 9.2105 | GSG1L |
| scaffold_4 | 44061695 | -20.0000 | GSG1L |
| scaffold_4 | 44082500 | -9.5238 | GSG1L |
| scaffold_4 | 44101126 | 5.9373 | GSG1L |
| scaffold_4 | 44104124 | -5.7143 | GSG1L |
| scaffold_4 | 44104130 | 18.0952 | GSG1L |
| scaffold_4 | 44104915 | -7.2088 | GSG1L |
| scaffold_4 | 44116433 | -9.6555 | GSG1L |
| scaffold_4 | 44136719 | -5.0794 | GSG1L |
| scaffold_4 | 44139455 | 5.9804 | GSG1L |
| scaffold_4 | 44150629 | 17.6136 | GSG1L |

| | | | |
|---|---|---|---|
| scaffold_4 | 44160648 | -6.6630 | GSG1L |
| scaffold_4 | 44169360 | 5.0000 | GSG1L |
| scaffold_4 | 44174763 | -5.1220 | GSG1L |
| scaffold_4 | 44175133 | 9.8485 | GSG1L |
| scaffold_4 | 44190252 | -14.0476 | GSG1L |
| scaffold_4 | 44202637 | -16.9848 | GSG1L |
| scaffold_4 | 44214795 | 10.1754 | GSG1L |
| scaffold_4 | 44222486 | -7.7447 | GSG1L |
| scaffold_4 | 44232563 | 15.4831 | GSG1L |
| scaffold_4 | 44232564 | 5.1973 | GSG1L |
| scaffold_4 | 44234613 | -10.2222 | GSG1L |
| scaffold_4 | 44234725 | 7.4297 | GSG1L |
| scaffold_4 | 44236368 | 11.1111 | GSG1L |
| scaffold_4 | 44236410 | 5.5556 | GSG1L |
| scaffold_4 | 44236942 | 8.9852 | GSG1L |
| scaffold_4 | 44236949 | 35.2632 | |
| scaffold_4 | 44393466 | -20.1754 | XPO6 |
| scaffold_4 | 44454586 | 9.8767 | SBK1 |
| scaffold_4 | 44467038 | 7.1429 | SBK1 |
| scaffold_4 | 44467392 | 6.6667 | SBK1 |
| scaffold_4 | 44468547 | 8.3333 | SBK1 |
| scaffold_4 | 44488455 | -8.8095 | SBK1 |
| scaffold_4 | 44488571 | 5.0000 | SBK1 |
| scaffold_4 | 44488576 | 19.4737 | SBK1 |
| scaffold_4 | 44488578 | -5.2632 | SBK1 |
| scaffold_4 | 44488674 | -10.5806 | SBK1 |
| scaffold_4 | 44488684 | -21.7936 | SBK1 |
| scaffold_4 | 44488689 | -7.4883 | SBK1 |
| scaffold_4 | 44488813 | 6.6038 | SBK1 |
| scaffold_4 | 44488871 | -5.5417 | SBK1 |
| scaffold_4 | 44488878 | 5.0735 | SBK1 |
| scaffold_4 | 44488891 | 16.6667 | SBK1 |
| scaffold_4 | 44488892 | 7.0075 | SBK1 |
| scaffold_4 | 44488920 | 6.0012 | SBK1 |
| scaffold_4 | 44555520 | 5.5944 | |
| scaffold_4 | 44555526 | 10.4895 | |
| scaffold_4 | 44555528 | 5.7829 | |
| scaffold_4 | 44555529 | -6.2937 | |
| scaffold_4 | 44567436 | -18.4211 | |
| scaffold_4 | 44571722 | 9.7475 | |
| scaffold_4 | 44571943 | -6.9462 | |
| scaffold_4 | 44579904 | 9.7580 | |
| scaffold_4 | 44592889 | -11.5385 | SPNS1 |
| scaffold_406 | 152282 | 5.1411 | FZR1 |
| scaffold_45 | 15510267 | 8.2431 | |
| scaffold_45 | 15510269 | 6.4685 | |
| scaffold_453 | 6114 | -5.2929 | |
| scaffold_49 | 1939847 | -6.9876 | |
| scaffold_5 | 15844801 | -5.4945 | COL4A2 |
| scaffold_5 | 15844948 | -6.6563 | COL4A2 |
| scaffold_5 | 15845441 | 7.5839 | COL4A2 |
| scaffold_5 | 23258461 | 21.0387 | NALCN |
| scaffold_5 | 23258467 | 22.3784 | NALCN |
| scaffold_5 | 23258488 | 18.7755 | NALCN |
| scaffold_523 | 67998 | -7.9622 | |
| scaffold_53 | 4518762 | 11.7647 | |
| scaffold_53 | 4677287 | -6.1410 | |
| scaffold_53 | 5044306 | -5.8824 | |

| | | | |
|---|---|---|---|
| scaffold_53 | 5044312 | 18.1818 | |
| scaffold_53 | 5044314 | -23.5294 | |
| scaffold_53 | 7199765 | 16.0665 | CACNA1G |
| scaffold_54 | 7899360 | -5.9915 | |
| scaffold_55 | 4084534 | 8.5859 | |
| scaffold_6 | 6394780 | -5.2083 | |
| scaffold_6 | 28209159 | 15.5012 | |
| scaffold_6 | 28209224 | 9.2207 | |
| scaffold_6 | 28797838 | 9.4080 | |
| scaffold_6 | 28797844 | -14.6783 | |
| scaffold_6 | 30601604 | -5.6696 | |
| scaffold_61 | 7389429 | -9.5694 | ARHGEF15 |
| scaffold_61 | 7389793 | 16.0037 | ARHGEF15 |
| scaffold_625 | 51932 | 19.7090 | |
| scaffold_626 | 26922 | 9.6469 | |
| scaffold_626 | 26973 | -12.0145 | |
| scaffold_626 | 26974 | 6.6087 | |
| scaffold_626 | 26976 | -25.4743 | |
| scaffold_64 | 1767906 | -16.1429 | |
| scaffold_67 | 9123900 | 5.4643 | ADAM11 |
| scaffold_7 | 47587053 | -6.6240 | NBEAL2 |
| scaffold_70 | 6539703 | 5.6991 | |
| scaffold_72 | 5426915 | 10.4167 | MAPK8IP1 |
| scaffold_76 | 343094 | -5.2422 | GRIN2C |
| scaffold_76 | 2371269 | -7.7778 | |
| scaffold_76 | 2371301 | -21.2181 | |
| scaffold_76 | 2517010 | -7.6923 | |
| scaffold_76 | 3034449 | 32.8571 | |
| scaffold_80 | 5842181 | -10.7494 | KLC3;ERCC2 |
| scaffold_9 | 1664796 | 9.4566 | |
| scaffold_90 | 5437056 | 6.3566 | |
| scaffold_96 | 319797 | -5.2632 | |
| scaffold_96 | 5699996 | -8.4071 | |
| scaffold_96 | 5699997 | 10.3505 | |
