## Supplemental Table 7 for "Conserved DNA Methylation Signatures in The Prefrontal Cortex of Newborn and Juvenile Guinea Pigs Following Antenatal Corticosteroid Exposure"

**Supplementary Table 7. List of DMCs identified within the binding site for SP1 in the PND1PFC following ACS exposure**

| seqnames | start | MethDiff | Gene name |
|---|---|---|---|
| scaffold_0 | 4938157 | 21.2032 | |
| scaffold_0 | 4938208 | 8.9113 | |
| scaffold_0 | 62694884 | -54.8485 | PRICKLE2 |
| scaffold_0 | 81521703 | 10.3668 | |
| scaffold_0 | 81521735 | 25.9259 | |
| scaffold_0 | 81521768 | -8.5299 | |
| scaffold_1 | 12188172 | 14.4483 | NR2F1 |
| scaffold_1 | **12189503** | 26.1029 | NR2F1 |
| scaffold_1 | 12189506 | 19.4989 | NR2F1 |
| scaffold_1 | 12189512 | 20.2381 | NR2F1 |
| scaffold_1 | 12189515 | 28.1746 | NR2F1 |
| scaffold_1 | 12189518 | 13.1250 | NR2F1 |
| scaffold_1 | 12189521 | 36.3636 | NR2F1 |
| scaffold_1 | 12189524 | 35.8586 | NR2F1 |
| scaffold_1 | 12189600 | 21.5686 | NR2F1 |
| scaffold_1 | 12189603 | 11.5196 | NR2F1 |
| scaffold_1 | 22073022 | -9.9795 | |
| scaffold_1 | 22073043 | -13.6364 | |
| scaffold_1 | 22073062 | -12.8938 | |
| scaffold_1 | 73765934 | 9.7214 | |
| scaffold_10 | 15327363 | 13.0848 | IL6R |
| scaffold_10 | 15327383 | 11.3562 | IL6R |
| scaffold_10 | 15327447 | 11.2500 | IL6R |
| scaffold_104 | 2565407 | -17.9426 | |
| scaffold_104 | 2565430 | 7.6555 | |
| scaffold_104 | 2571559 | 7.0707 | |
| scaffold_11 | 13617836 | 23.7925 | PAX2 |
| scaffold_1100 | 6538 | 8.8071 | |
| scaffold_111 | 12729 | -7.6879 | |
| scaffold_111 | 1782351 | 11.4695 | |
| scaffold_115 | 1404768 | -7.5391 | SETD1B |
| scaffold_115 | 1405669 | 12.5000 | SETD1B |
| scaffold_115 | 1405682 | -11.1742 | SETD1B |
| scaffold_115 | 1407238 | 11.1111 | SETD1B |
| scaffold_115 | 1407488 | 8.3333 | SETD1B |
| scaffold_115 | 1407494 | 15.9091 | SETD1B |
| scaffold_115 | 4048902 | -5.4612 | |
| scaffold_117 | 3775742 | -7.1982 | UNC5D |
| scaffold_119 | 328159 | 5.7595 | |
| scaffold_119 | 1850992 | 18.7500 | |
| scaffold_13 | 4860579 | -11.8926 | |
| scaffold_13 | 6041158 | 6.4604 | USH2A |
| scaffold_13 | 6042569 | -9.4436 | USH2A |
| scaffold_13 | 7208132 | -6.2290 | |
| scaffold_137 | 2017366 | 11.3979 | ITPKB |
| scaffold_140 | 985245 | -21.1310 | LARGE1 |
| scaffold_143 | 83956 | 5.1709 | |
| scaffold_15 | 29335091 | 23.7132 | |
| scaffold_157 | 449710 | 7.2330 | |
| scaffold_158 | 1702678 | 5.9524 | |

| | | | |
|---|---|---|---|
| scaffold_158 | 1702743 | -7.5832 | |
| scaffold_16 | 15313760 | 7.8938 | GATA3 |
| scaffold_168 | 1567710 | 5.0000 | AKAP8L |
| scaffold_17 | 37385681 | -15.0000 | TRAPPC9 |
| scaffold_171 | 219440 | -5.8497 | RFX4 |
| scaffold_18 | 38752726 | 5.7937 | |
| scaffold_184 | 170790 | 10.0000 | SHANK3 |
| scaffold_19 | 30245258 | 6.2791 | FXYD2 |
| scaffold_19 | 30245445 | -23.1951 | FXYD2 |
| scaffold_19 | 30245504 | 25.9485 | FXYD2 |
| scaffold_2 | 28509248 | -5.4054 | |
| scaffold_2 | 28541024 | -8.2050 | |
| scaffold_2035 | 5181 | -5.5080 | |
| scaffold_25 | 11570287 | -6.7382 | MYOM3 |
| scaffold_25 | 17962742 | -10.0390 | |
| scaffold_25 | 18474719 | 23.2143 | FHAD1 |
| scaffold_25 | 23409107 | 6.5657 | PEX14 |
| scaffold_25 | 26209855 | -5.4422 | ESPN |
| scaffold_25 | 26209856 | -12.0623 | ESPN |
| scaffold_25 | 26209860 | -12.0076 | ESPN |
| scaffold_25 | 26209876 | -17.8874 | ESPN |
| scaffold_25 | 26209882 | -11.8179 | ESPN |
| scaffold_25 | 27347393 | 5.1136 | AJAP1 |
| scaffold_255 | 162147 | 7.9741 | |
| scaffold_27 | 1730781 | -8.8580 | |
| scaffold_27 | 9691327 | -23.1250 | CNTFR |
| scaffold_27 | 14983883 | 11.6930 | CNTFR |
| scaffold_28 | 21356457 | 17.6471 | |
| scaffold_3 | 19746232 | -5.8824 | CFAP210 |
| scaffold_30 | 3585195 | -9.4886 | MEPCE |
| scaffold_304 | 292751 | -7.5094 | |
| scaffold_31 | 12094872 | 12.1441 | |
| scaffold_31 | 14524323 | 10.9848 | |
| scaffold_31 | 14524353 | 33.2727 | |
| scaffold_31 | 14581588 | 12.0946 | |
| scaffold_31 | 17479190 | 9.8116 | |
| scaffold_32 | 20114786 | 5.3070 | MNT |
| scaffold_32 | 23946888 | 18.1582 | FOXN1 |
| scaffold_334 | 104291 | 15.4237 | |
| scaffold_334 | 104298 | 6.0725 | |
| scaffold_334 | 406251 | 5.5534 | |
| scaffold_356 | 220749 | -23.8095 | |
| scaffold_4 | 43658158 | -5.0132 | IL4R |
| scaffold_4 | 43658576 | 8.0000 | IL4R |
| scaffold_4 | 43658589 | 8.3333 | IL4R |
| scaffold_4 | 43662295 | 5.2632 | IL4R |
| scaffold_4 | 43671870 | 7.1161 | |
| scaffold_4 | 43672862 | 12.3372 | |
| scaffold_4 | 43672863 | -7.7429 | |
| scaffold_4 | 43672894 | 9.7500 | |
| scaffold_4 | 43672895 | -5.7961 | |
| scaffold_4 | 43674500 | -10.9337 | |

| | | | |
|---|---|---|---|
| scaffold_4 | 43674749 | -13.6907 | |
| scaffold_4 | 43679466 | 5.4657 | |
| scaffold_4 | 43679521 | -5.2952 | |
| scaffold_4 | 43731471 | -17.2917 | IL21R |
| scaffold_4 | 43732455 | -10.0000 | IL21R |
| scaffold_4 | 43738119 | -6.3158 | |
| scaffold_4 | 43745108 | 6.0606 | |
| scaffold_4 | 43976249 | 7.2896 | KATNIP |
| scaffold_4 | 43979316 | -9.3137 | KATNIP |
| scaffold_4 | 43983993 | 5.3181 | KATNIP |
| scaffold_4 | 43994229 | 8.3810 | KATNIP |
| scaffold_4 | 43994234 | 15.0794 | KATNIP |
| scaffold_4 | 43999584 | -8.6761 | KATNIP |
| scaffold_4 | 44010550 | 11.7949 | |
| scaffold_4 | 44018489 | -9.7619 | GSG1L |
| scaffold_4 | 44034417 | 10.0000 | GSG1L |
| scaffold_4 | 44060216 | 9.2593 | GSG1L |
| scaffold_4 | 44061694 | 9.2105 | GSG1L |
| scaffold_4 | 44061695 | -20.0000 | GSG1L |
| scaffold_4 | 44084051 | 7.1192 | GSG1L |
| scaffold_4 | 44101031 | 7.4423 | GSG1L |
| scaffold_4 | 44101126 | 5.9373 | GSG1L |
| scaffold_4 | 44103777 | -14.2857 | GSG1L |
| scaffold_4 | 44116433 | -9.6555 | GSG1L |
| scaffold_4 | 44136719 | -5.0794 | GSG1L |
| scaffold_4 | 44139447 | 5.1531 | GSG1L |
| scaffold_4 | 44139455 | 5.9804 | GSG1L |
| scaffold_4 | 44139743 | 9.0332 | GSG1L |
| scaffold_4 | 44160648 | -6.6630 | GSG1L |
| scaffold_4 | 44169360 | 5.0000 | GSG1L |
| scaffold_4 | 44202637 | -16.9848 | GSG1L |
| scaffold_4 | 44204702 | -11.1111 | GSG1L |
| scaffold_4 | 44215816 | 13.6842 | GSG1L |
| scaffold_4 | 44232563 | 15.4831 | GSG1L |
| scaffold_4 | 44232564 | 5.1973 | GSG1L |
| scaffold_4 | 44232584 | -12.4903 | GSG1L |
| scaffold_4 | 44232749 | 18.9878 | GSG1L |
| scaffold_4 | 44236942 | 8.9852 | GSG1L |
| scaffold_4 | 44236949 | 35.2632 | |
| scaffold_4 | 44373678 | -7.4286 | XPO6 |
| scaffold_4 | 44413373 | 12.5167 | |
| scaffold_4 | 44442604 | -10.9176 | SBK1 |
| scaffold_4 | 44442637 | -5.4174 | SBK1 |
| scaffold_4 | 44454586 | 9.8767 | SBK1 |
| scaffold_4 | 44460743 | -8.5857 | SBK1 |
| scaffold_4 | 44488610 | -12.5000 | SBK1 |
| scaffold_4 | 44488674 | -10.5806 | SBK1 |
| scaffold_4 | 44488689 | -7.4883 | SBK1 |
| scaffold_4 | 44488878 | 5.0735 | SBK1 |
| scaffold_4 | 44488891 | 16.6667 | SBK1 |
| scaffold_4 | 44488892 | 7.0075 | SBK1 |
| scaffold_4 | 44555526 | 10.4895 | |

| | | | |
|---|---|---|---|
| scaffold_4 | 44555528 | 5.7829 | |
| scaffold_4 | 44555529 | -6.2937 | |
| scaffold_4 | 44566137 | -19.0476 | |
| scaffold_4 | 44567419 | -6.0150 | |
| scaffold_4 | 44567436 | -18.4211 | |
| scaffold_4 | 44573656 | 10.9524 | |
| scaffold_406 | 152282 | 5.1411 | FZR1 |
| scaffold_49 | 1939983 | 8.5979 | |
| scaffold_49 | 2011633 | -5.3059 | |
| scaffold_53 | 4161209 | -19.5238 | |
| scaffold_53 | 4518762 | 11.7647 | |
| scaffold_55 | 4084534 | 8.5859 | |
| scaffold_61 | 7389429 | -9.5694 | ARHGEF15 |
| scaffold_61 | 7389793 | 16.0037 | ARHGEF15 |
| scaffold_61 | 7390058 | -11.6654 | ARHGEF15 |
| scaffold_62 | 5321709 | 7.2801 | |
| scaffold_62 | 5323770 | 5.1720 | |
| scaffold_626 | 26922 | 9.6469 | |
| scaffold_64 | 1767673 | -14.7605 | |
| scaffold_67 | 7858889 | 27.5337 | OTOG |
| scaffold_67 | 9123772 | -5.4579 | ADAM11 |
| scaffold_67 | 9123847 | -12.2166 | ADAM11 |
| scaffold_7 | 47587053 | -6.6240 | NBEAL2 |
| scaffold_70 | 6539703 | 5.6991 | |
| scaffold_72 | 3962647 | -8.5442 | |
| scaffold_72 | 4167957 | -10.5882 | EXT2 |
| scaffold_76 | 273224 | 11.5000 | OTOP2 |
| scaffold_76 | 2517037 | 16.6667 | |
| scaffold_76 | 3034449 | 32.8571 | |
| scaffold_76 | 3614197 | 7.7124 | DNAH17 |
| scaffold_76 | 3614202 | 5.5719 | DNAH17 |
| scaffold_88 | 6464021 | -12.6112 | GLIS1 |
| scaffold_9 | 40091257 | -5.2810 | AGAP2 |
| scaffold_90 | 5437056 | 6.3566 | |
| scaffold_96 | 319797 | -5.2632 | |
