## Supplemental Table 8 for "Conserved DNA Methylation Signatures in The Prefrontal Cortex of Newborn and Juvenile Guinea Pigs Following Antenatal Corticosteroid Exposure"

**Supplementary Table 8. List of DMCs identified within the binding site for ZNF263 in the PND1PFC following ACS exposure**

| seqnames | start | MethDiff | Gene name |
|---|---|---|---|
| scaffold_0 | 4938109 | 19.7743 | |
| scaffold_0 | 4938208 | 8.9113 | |
| scaffold_0 | 81521735 | 25.9259 | |
| scaffold_0 | 81521768 | -8.5299 | |
| scaffold_1 | 12189558 | 24.4881 | NR2F1 |
| scaffold_1 | 12189615 | 20.5882 | NR2F1 |
| scaffold_1 | 12189617 | 17.4020 | NR2F1 |
| scaffold_1 | 42320161 | 22.1803 | UROS |
| scaffold_1 | 67622577 | 11.1722 | MCPH1 |
| scaffold_1 | 67622582 | 10.7028 | MCPH1 |
| scaffold_10 | 15327363 | 13.0848 | IL6R |
| scaffold_10 | 15327371 | 14.2178 | IL6R |
| scaffold_10 | 35166327 | 50.8772 | |
| scaffold_10 | 45822346 | -18.8235 | |
| scaffold_10 | 47593463 | -18.8811 | TARS3 |
| scaffold_104 | 2565409 | -11.2440 | |
| scaffold_104 | 2571505 | 15.0000 | |
| scaffold_104 | 2571575 | 7.7947 | |
| scaffold_104 | 2571576 | 20.7071 | |
| scaffold_104 | 2571678 | 18.7500 | |
| scaffold_104 | 2571714 | 29.6970 | |
| scaffold_11 | 13617836 | 23.7925 | PAX2 |
| caffold_1112 | 15496 | -18.5714 | |
| scaffold_115 | 1407238 | 11.1111 | SETD1B |
| scaffold_115 | 1407593 | 5.3931 | SETD1B |
| scaffold_115 | 1407599 | 7.6628 | SETD1B |
| scaffold_119 | 1850983 | 13.3333 | |
| scaffold_121 | 19291 | 5.8106 | |
| scaffold_13 | 4860599 | -15.5280 | |
| scaffold_13 | 4860603 | -26.9565 | |
| scaffold_13 | 6041158 | 6.4604 | USH2A |
| scaffold_13 | 7208132 | -6.2290 | |
| scaffold_143 | 83956 | 5.1709 | |
| scaffold_15 | 33165439 | -7.6923 | |
| scaffold_15 | 34393766 | 9.1119 | TBC1D22B |
| scaffold_17 | 37385710 | -8.7500 | TRAPPC9 |
| scaffold_17 | 37385713 | 9.0543 | TRAPPC9 |
| scaffold_171 | 219389 | 5.2880 | RFX4 |
| scaffold_171 | 219390 | 16.0606 | RFX4 |
| scaffold_171 | 219440 | -5.8497 | RFX4 |
| scaffold_18 | 38752273 | 7.4286 | |
| scaffold_184 | 170790 | 10.0000 | SHANK3 |
| scaffold_19 | 30245258 | 6.2791 | FXYD2 |
| scaffold_19 | 30245445 | -23.1951 | FXYD2 |
| scaffold_19 | 30245504 | 25.9485 | FXYD2 |
| scaffold_203 | 103315 | -26.9231 | TPCN1 |
| caffold_2035 | 5027 | 5.8110 | |
| caffold_2035 | 5181 | -5.5080 | |
| scaffold_226 | 527608 | 7.5295 | ODAD3 |
| scaffold_226 | 527633 | -7.7592 | ODAD3 |
| scaffold_226 | 631257 | -7.1429 | |
| scaffold_226 | 683602 | 10.6681 | ACP5 |
| scaffold_226 | 683751 | -9.1024 | ACP5 |
| scaffold_23 | 25112886 | -5.6843 | |
| scaffold_25 | 23409107 | 6.5657 | PEX14 |
| scaffold_25 | 26209855 | -5.4422 | ESPN |

| | | | |
|---|---|---|---|
| scaffold_25 | 26209856 | -12.0623 | ESPN |
| scaffold_25 | 26209860 | -12.0076 | ESPN |
| scaffold_25 | 26209876 | -17.8874 | ESPN |
| scaffold_25 | 26211125 | 6.8760 | ESPN |
| scaffold_25 | 27347393 | 5.1136 | AJAP1 |
| scaffold_255 | 162147 | 7.9741 | |
| scaffold_27 | 1730781 | -8.8580 | |
| scaffold_27 | 9691097 | -14.7619 | NEK6 |
| scaffold_3 | 18814469 | 6.1905 | GAD1 |
| scaffold_3 | 19746232 | -5.8824 | CFAP210 |
| scaffold_3 | 19757807 | 5.6152 | CFAP210 |
| scaffold_304 | 292751 | -7.5094 | |
| scaffold_304 | 292752 | -5.8028 | |
| scaffold_31 | 5395688 | 18.1818 | |
| scaffold_31 | 9017839 | 6.1012 | |
| scaffold_31 | 12094872 | 12.1441 | |
| scaffold_31 | 14524323 | 10.9848 | |
| scaffold_31 | 14524353 | 33.2727 | |
| scaffold_31 | 14581579 | 11.4865 | |
| scaffold_31 | 14581588 | 12.0946 | |
| scaffold_32 | 19461802 | 5.0362 | |
| scaffold_33 | 24466781 | 12.6812 | |
| scaffold_334 | 104291 | 15.4237 | |
| scaffold_334 | 104298 | 6.0725 | |
| scaffold_334 | 406251 | 5.5534 | |
| scaffold_34 | 3046489 | -12.9032 | |
| scaffold_38 | 6902950 | -5.6892 | |
| scaffold_4 | 43658158 | -5.0132 | IL4R |
| scaffold_4 | 43658576 | 8.0000 | IL4R |
| scaffold_4 | 43671870 | 7.1161 | |
| scaffold_4 | 43672912 | -7.4522 | |
| scaffold_4 | 43674749 | -13.6907 | |
| scaffold_4 | 43678510 | 15.7289 | |
| scaffold_4 | 43731167 | 7.0376 | IL21R |
| scaffold_4 | 43731354 | -7.1622 | IL21R |
| scaffold_4 | 43732455 | -10.0000 | IL21R |
| scaffold_4 | 43737303 | 8.1731 | |
| scaffold_4 | 43738424 | -10.1307 | |
| scaffold_4 | 43976249 | 7.2896 | KATNIP |
| scaffold_4 | 43979316 | -9.3137 | KATNIP |
| scaffold_4 | 43983903 | -5.4622 | KATNIP |
| scaffold_4 | 43987132 | 6.4016 | KATNIP |
| scaffold_4 | 44007168 | 7.8848 | |
| scaffold_4 | 44018489 | -9.7619 | GSG1L |
| scaffold_4 | 44036221 | -7.1429 | GSG1L |
| scaffold_4 | 44091163 | -18.1818 | GSG1L |
| scaffold_4 | 44101031 | 7.4423 | GSG1L |
| scaffold_4 | 44101126 | 5.9373 | GSG1L |
| scaffold_4 | 44103777 | -14.2857 | GSG1L |
| scaffold_4 | 44104124 | -5.7143 | GSG1L |
| scaffold_4 | 44169360 | 5.0000 | GSG1L |
| scaffold_4 | 44174934 | -6.5217 | GSG1L |
| scaffold_4 | 44175593 | 5.0699 | GSG1L |
| scaffold_4 | 44180856 | -13.3838 | GSG1L |
| scaffold_4 | 44202602 | -5.2733 | GSG1L |
| scaffold_4 | 44202637 | -16.9848 | GSG1L |
| scaffold_4 | 44204702 | -11.1111 | GSG1L |
| scaffold_4 | 44214795 | 10.1754 | GSG1L |

| | | | |
|---|---|---|---|
| scaffold_4 | 44214803 | -7.0175 | GSG1L |
| scaffold_4 | 44215816 | 13.6842 | GSG1L |
| scaffold_4 | 44216751 | -7.1290 | GSG1L |
| scaffold_4 | 44217321 | -6.5015 | GSG1L |
| scaffold_4 | 44232516 | 9.6528 | GSG1L |
| scaffold_4 | 44232563 | 15.4831 | GSG1L |
| scaffold_4 | 44232564 | 5.1973 | GSG1L |
| scaffold_4 | 44232597 | -6.1905 | GSG1L |
| scaffold_4 | 44232654 | -6.0968 | GSG1L |
| scaffold_4 | 44236749 | -10.9524 | GSG1L |
| scaffold_4 | 44236942 | 8.9852 | GSG1L |
| scaffold_4 | 44236949 | 35.2632 | |
| scaffold_4 | 44294777 | 6.6667 | XPO6 |
| scaffold_4 | 44373678 | -7.4286 | XPO6 |
| scaffold_4 | 44413373 | 12.5167 | |
| scaffold_4 | 44413374 | 21.4286 | |
| scaffold_4 | 44467392 | 6.6667 | SBK1 |
| scaffold_4 | 44468547 | 8.3333 | SBK1 |
| scaffold_4 | 44488610 | -12.5000 | SBK1 |
| scaffold_4 | 44488878 | 5.0735 | SBK1 |
| scaffold_4 | 44567125 | -5.0847 | |
| scaffold_4 | 44567436 | -18.4211 | |
| scaffold_4 | 44578994 | -13.0390 | |
| scaffold_406 | 152282 | 5.1411 | FZR1 |
| scaffold_45 | 15510178 | -5.0792 | |
| scaffold_48 | 12757823 | 6.1309 | CCSER1 |
| scaffold_5 | 14328198 | -7.1579 | DCUN1D2 |
| scaffold_53 | 3965510 | -12.3077 | |
| scaffold_53 | 7199765 | 16.0665 | CACNA1G |
| scaffold_6 | 24965245 | -10.6481 | |
| scaffold_6 | 28797664 | 5.4855 | |
| scaffold_61 | 7389447 | 6.4593 | ARHGEF15 |
| scaffold_61 | 7389793 | 16.0037 | ARHGEF15 |
| scaffold_626 | 26922 | 9.6469 | |
| scaffold_626 | 26973 | -12.0145 | |
| scaffold_626 | 26974 | 6.6087 | |
| scaffold_626 | 26976 | -25.4743 | |
| scaffold_64 | 1767673 | -14.7605 | |
| scaffold_67 | 7858889 | 27.5337 | OTOG |
| scaffold_7 | 47587053 | -6.6240 | NBEAL2 |
| scaffold_7 | 51192435 | 6.9749 | ETV5 |
| scaffold_70 | 1150925 | -5.9006 | |
| scaffold_70 | 7500245 | -8.5026 | |
| scaffold_72 | 4167957 | -10.5882 | EXT2 |
| scaffold_72 | 8384344 | 8.6275 | ARMC6 |
| scaffold_76 | 343094 | -5.2422 | GRIN2C |
| scaffold_76 | 3614172 | -7.0234 | DNAH17 |
| scaffold_76 | 3614197 | 7.7124 | DNAH17 |
| scaffold_80 | 5842187 | -8.6353 | KLC3;ERCC2 |
| scaffold_81 | 570123 | 14.2708 | |
| scaffold_88 | 6464068 | -8.5366 | GLIS1 |
| scaffold_9 | 8095962 | -26.5217 | |
| scaffold_9 | 8095963 | -8.4927 | |
| scaffold_9 | 42030018 | -6.5723 | SMARCC2 |
| scaffold_90 | 5437056 | 6.3566 | |
| scaffold_94 | 5701096 | -12.7101 | |
| scaffold_96 | 319797 | -5.2632 | |
| scaffold_96 | 5578812 | -5.3902 | |
