## Supplemental Table 9 for "Conserved DNA Methylation Signatures in The Prefrontal Cortex of Newborn and Juvenile Guinea Pigs Following Antenatal Corticosteroid Exposure"

**Supplemental Table 9. Full list of differentially methylated cytosines in the guinea pig PND14PFC following ACS exposure**

| | seqnames | start | MethDiff | geneName |
|---|---|---|---|---|
| 1 | scaffold_0 | 3665236 | -15.88 | |
| 2 | scaffold_0 | 3665268 | -25.68 | |
| 3 | scaffold_0 | 3684576 | 12.67 | RIMS2 |
| 4 | scaffold_0 | 3684590 | 19.40 | RIMS2 |
| 5 | scaffold_0 | 3684617 | 13.45 | RIMS2 |
| 6 | scaffold_0 | 3684618 | 15.75 | RIMS2 |
| 7 | scaffold_0 | 3684621 | 11.24 | RIMS2 |
| 8 | scaffold_0 | 3684622 | 18.31 | RIMS2 |
| 9 | scaffold_0 | 3684666 | 12.69 | RIMS2 |
| 10 | scaffold_0 | 3684678 | 18.17 | RIMS2 |
| 11 | scaffold_0 | 5589377 | -13.89 | |
| 12 | scaffold_0 | 5589386 | -13.38 | |
| 13 | scaffold_0 | 5589421 | -15.85 | |
| 14 | scaffold_0 | 5589427 | -24.23 | |
| 15 | scaffold_0 | 5589428 | -28.50 | |
| 16 | scaffold_0 | 9497540 | 32.14 | |
| 17 | scaffold_0 | 9497546 | -15.79 | |
| 18 | scaffold_0 | 9497547 | 15.18 | |
| 19 | scaffold_0 | 9497559 | -19.62 | |
| 20 | scaffold_0 | 9497560 | -13.59 | |
| 21 | scaffold_0 | 9497571 | -15.95 | |
| 22 | scaffold_0 | 9497572 | -21.48 | |
| 23 | scaffold_0 | 9497625 | -21.43 | |
| 24 | scaffold_0 | 87064889 | 5.47 | |
| 25 | scaffold_0 | 87064923 | 14.97 | |
| 26 | scaffold_0 | 87064991 | 19.96 | |
| 27 | scaffold_0 | 87065033 | 26.70 | |
| 28 | scaffold_0 | 87065049 | 32.31 | |
| 29 | scaffold_0 | 87065058 | 13.78 | |
| 30 | scaffold_0 | 87065059 | 34.71 | |
| 31 | scaffold_0 | 87065072 | 12.61 | |
| 32 | scaffold_0 | 87065090 | 14.27 | |
| 33 | scaffold_0 | 87065091 | 9.58 | |
| 34 | scaffold_0 | 87065126 | 12.42 | |
| 35 | scaffold_0 | 87065160 | 7.77 | |
| 36 | scaffold_1 | 10198632 | 47.22 | |
| 37 | scaffold_1 | 10198634 | 26.11 | |
| 38 | scaffold_1 | 10198650 | 27.78 | |
| 39 | scaffold_1 | 10198664 | 47.78 | |
| 40 | scaffold_1 | 65104682 | -45.45 | |
| 41 | scaffold_1 | 65104691 | -51.14 | |
| 42 | scaffold_1 | 65104706 | -12.50 | |
| 43 | scaffold_1 | 65104720 | -51.14 | |
| 44 | scaffold_10 | 48577576 | -17.86 | APBA2 |
| 45 | scaffold_104 | 2632168 | 17.88 | |
| 46 | scaffold_104 | 2632179 | 5.45 | |
| 47 | scaffold_104 | 2632216 | 18.00 | |
| 48 | scaffold_104 | 2635279 | -7.50 | |
| 49 | scaffold_109 | 2866369 | -13.77 | PLXND1 |
| 50 | scaffold_109 | 2866406 | 7.98 | PLXND1 |
| 51 | scaffold_109 | 2866416 | -5.98 | PLXND1 |
| 52 | scaffold_109 | 2866417 | -10.59 | PLXND1 |
| 53 | scaffold_11 | 5053384 | -21.07 | |
| 54 | scaffold_11 | 5059043 | 12.07 | |
| 55 | scaffold_11 | 5059627 | -10.09 | |
| 56 | scaffold_11 | 5059653 | -10.43 | |

| | | | | |
|---|---|---|---|---|
| 57 | scaffold_11 | 5059839 | 6.45 | |
| 58 | scaffold_11 | 5062648 | 11.01 | |
| 59 | scaffold_11 | 5062946 | -7.20 | |
| 60 | scaffold_11 | 14181692 | -16.67 | |
| 61 | scaffold_11 | 14181700 | 16.67 | |
| 62 | scaffold_114 | 2615152 | -7.14 | |
| 63 | scaffold_115 | 2716887 | -5.86 | CUX2 |
| 64 | scaffold_125 | 3159214 | -9.79 | ATP6V1C2 |
| 65 | scaffold_125 | 3159545 | 19.12 | ATP6V1C2 |
| 66 | scaffold_125 | 3159554 | -16.95 | ATP6V1C2 |
| 67 | scaffold_13 | 9282875 | -15.00 | |
| 68 | scaffold_13 | 9282914 | 5.09 | |
| 69 | scaffold_13 | 9284009 | -10.43 | |
| 70 | scaffold_13 | 36256829 | -27.27 | |
| 71 | scaffold_13 | 36257091 | -6.25 | |
| 72 | scaffold_132 | 672314 | -8.33 | RBM10 |
| 73 | scaffold_135 | 1226802 | -8.15 | |
| 74 | scaffold_135 | 1226803 | -10.38 | |
| 75 | scaffold_135 | 1226867 | -11.32 | |
| 76 | scaffold_135 | 1226935 | 7.39 | |
| 77 | scaffold_14 | 40432675 | -6.43 | |
| 78 | scaffold_140 | 746133 | 8.25 | TIMP3 |
| 79 | scaffold_140 | 746180 | 26.76 | TIMP3 |
| 80 | scaffold_140 | 746181 | 5.27 | TIMP3 |
| 81 | scaffold_140 | 746200 | 18.18 | TIMP3 |
| 82 | scaffold_140 | 746201 | -5.39 | TIMP3 |
| 83 | scaffold_140 | 746246 | -6.17 | TIMP3 |
| 84 | scaffold_140 | 746256 | 9.43 | TIMP3 |
| 85 | scaffold_140 | 746417 | -8.40 | TIMP3 |
| 86 | scaffold_140 | 746461 | -6.58 | TIMP3 |
| 87 | scaffold_140 | 746909 | -9.01 | TIMP3 |
| 88 | scaffold_140 | 746984 | -5.39 | TIMP3 |
| 89 | scaffold_140 | 747019 | 6.12 | TIMP3 |
| 90 | scaffold_140 | 747038 | 10.01 | TIMP3 |
| 91 | scaffold_140 | 747309 | -8.24 | TIMP3 |
| 92 | scaffold_140 | 747375 | -21.00 | TIMP3 |
| 93 | scaffold_140 | 747592 | 17.50 | TIMP3 |
| 94 | scaffold_15 | 23763883 | -6.53 | |
| 95 | scaffold_15 | 32550138 | 20.24 | |
| 96 | scaffold_15 | 32550317 | 17.85 | |
| 97 | scaffold_15 | 32550353 | -13.42 | |
| 98 | scaffold_15 | 32550354 | 9.26 | |
| 99 | scaffold_15 | 32550359 | -11.26 | |
| 100 | scaffold_15 | 32550380 | 15.71 | |
| 101 | scaffold_15 | 32550395 | 31.85 | |
| 102 | scaffold_15 | 32550409 | 11.85 | |
| 103 | scaffold_15 | 32550413 | 33.01 | |
| 104 | scaffold_15 | 33089033 | -18.33 | |
| 105 | scaffold_15 | 33089086 | 16.19 | |
| 106 | scaffold_15 | 33089111 | -9.49 | |
| 107 | scaffold_15 | 34265179 | 18.37 | FGD2 |
| 108 | scaffold_15 | 38692364 | 14.01 | |
| 109 | scaffold_155 | 1146693 | -6.37 | |
| 110 | scaffold_155 | 1146720 | -5.56 | |
| 111 | scaffold_155 | 1146733 | 5.87 | |
| 112 | scaffold_155 | 1146827 | 5.52 | |
| 113 | scaffold_155 | 1146843 | -7.59 | |
| 114 | scaffold_155 | 1146922 | 5.36 | |

| | | | | |
|---|---|---|---|---|
| 115 | scaffold_155 | 1147655 | -6.98 | |
| 116 | scaffold_155 | 1147857 | -5.99 | |
| 117 | scaffold_155 | 1147870 | -6.86 | |
| 118 | scaffold_159 | 636934 | -11.54 | |
| 119 | scaffold_159 | 636982 | -11.67 | |
| 120 | scaffold_159 | 637027 | -5.66 | |
| 121 | scaffold_159 | 637028 | 6.02 | |
| 122 | scaffold_159 | 637035 | -5.56 | |
| 123 | scaffold_160 | 267844 | -10.71 | |
| 124 | scaffold_160 | 267890 | -7.14 | |
| 125 | scaffold_166 | 578641 | -23.56 | SORCS2 |
| 126 | scaffold_166 | 1409905 | -5.56 | DOK7 |
| 127 | scaffold_166 | 1410143 | 20.15 | DOK7 |
| 128 | scaffold_171 | 777263 | 9.71 | |
| 129 | scaffold_171 | 777299 | 6.13 | |
| 130 | scaffold_171 | 777304 | -6.23 | |
| 131 | scaffold_171 | 777314 | -5.40 | |
| 132 | scaffold_171 | 777315 | -11.81 | |
| 133 | scaffold_171 | 1555507 | 9.31 | CHST11 |
| 134 | scaffold_171 | 1555817 | 7.86 | CHST11 |
| 135 | scaffold_171 | 1556283 | -5.04 | CHST11 |
| 136 | scaffold_178 | 1472271 | 6.36 | |
| 137 | scaffold_178 | 1472285 | 7.65 | |
| 138 | scaffold_178 | 1472287 | 8.12 | |
| 139 | scaffold_178 | 1472335 | 8.06 | |
| 140 | scaffold_178 | 1472399 | 6.82 | |
| 141 | scaffold_178 | 1472622 | -10.00 | |
| 142 | scaffold_18 | 2776827 | 8.72 | |
| 143 | scaffold_18 | 2777029 | -9.61 | |
| 144 | scaffold_18 | 39948740 | 11.54 | |
| 145 | scaffold_18 | 39949155 | 9.10 | |
| 146 | scaffold_19 | 30377606 | 7.50 | DSCAML1 |
| 147 | scaffold_19 | 30378554 | -8.80 | DSCAML1 |
| 148 | scaffold_19 | 36033256 | -18.14 | PPP2R1B |
| 149 | scaffold_19 | 36033296 | 9.91 | PPP2R1B |
| 150 | scaffold_19 | 36033423 | -7.32 | PPP2R1B |
| 151 | scaffold_2 | 36381253 | -16.56 | KCND3 |
| 152 | scaffold_2 | 36381254 | -6.57 | KCND3 |
| 153 | scaffold_2 | 36381297 | -11.35 | KCND3 |
| 154 | scaffold_2 | 36381304 | -5.76 | KCND3 |
| 155 | scaffold_2 | 46708451 | -18.31 | |
| 156 | scaffold_2 | 46708545 | -15.01 | |
| 157 | scaffold_2 | 46708591 | -7.21 | |
| 158 | scaffold_2 | 46708593 | -12.15 | |
| 159 | scaffold_2 | 46708610 | -10.13 | |
| 160 | scaffold_2 | 61538899 | 8.82 | |
| 161 | scaffold_2 | 65848849 | 6.50 | GNG12 |
| 162 | scaffold_2 | 67751748 | -8.05 | CACHD1 |
| 163 | scaffold_2 | 67751749 | -5.17 | CACHD1 |
| 164 | scaffold_2 | 67751763 | -10.99 | CACHD1 |
| 165 | scaffold_2 | 67752016 | -6.25 | CACHD1 |
| 166 | scaffold_2 | 67752038 | -13.54 | CACHD1 |
| 167 | scaffold_2 | 78013496 | 6.67 | |
| 168 | scaffold_2 | 78015134 | 10.38 | |
| 169 | scaffold_2 | 78017266 | 5.64 | |
| 170 | scaffold_2 | 78017478 | 7.91 | |
| 171 | scaffold_2 | 78017753 | -12.77 | |
| 172 | scaffold_2 | 78017820 | 10.29 | |

| | | | | |
|---|---|---|---|---|
| 173 | scaffold_2 | 78018521 | -6.25 | |
| 174 | scaffold_2 | 78018616 | -6.44 | |
| 175 | scaffold_2 | 78018691 | -9.43 | |
| 176 | scaffold_20 | 3718371 | -5.94 | |
| 177 | scaffold_20 | 3718381 | -6.89 | |
| 178 | scaffold_20 | 3719478 | 5.14 | |
| 179 | scaffold_24 | 2100133 | 9.18 | |
| 180 | scaffold_24 | 2101487 | 6.54 | |
| 181 | scaffold_24 | 2101600 | -16.59 | |
| 182 | scaffold_24 | 8136134 | 8.70 | |
| 183 | scaffold_24 | 8136414 | 8.75 | |
| 184 | scaffold_24 | 8136437 | -6.11 | |
| 185 | scaffold_24 | 8136779 | 13.53 | |
| 186 | scaffold_25 | 11324404 | -10.27 | NIPAL3 |
| 187 | scaffold_25 | 11325953 | -14.29 | NIPAL3 |
| 188 | scaffold_25 | 11325987 | -15.38 | NIPAL3 |
| 189 | scaffold_25 | 11326241 | -8.00 | NIPAL3 |
| 190 | scaffold_25 | 11326247 | -5.10 | NIPAL3 |
| 191 | scaffold_25 | 16284723 | -12.94 | |
| 192 | scaffold_25 | 23048688 | 15.30 | |
| 193 | scaffold_25 | 23048689 | -6.77 | |
| 194 | scaffold_25 | 23048694 | 6.98 | |
| 195 | scaffold_25 | 23048726 | 15.38 | |
| 196 | scaffold_25 | 26474908 | 5.51 | |
| 197 | scaffold_25 | 26474953 | -7.01 | |
| 198 | scaffold_25 | 26476541 | 30.05 | |
| 199 | scaffold_27 | 909219 | -7.11 | |
| 200 | scaffold_27 | 909575 | 9.09 | |
| 201 | scaffold_27 | 909610 | -8.33 | |
| 202 | scaffold_27 | 2634722 | -17.25 | HMCN2 |
| 203 | scaffold_27 | 11748547 | -29.55 | |
| 204 | scaffold_28 | 22471980 | -5.74 | BAIAP2L2 |
| 205 | scaffold_28 | 22472036 | 5.91 | BAIAP2L2 |
| 206 | scaffold_28 | 22472080 | -12.61 | BAIAP2L2 |
| 207 | scaffold_28 | 22472103 | -9.09 | BAIAP2L2 |
| 208 | scaffold_28 | 22472104 | -7.03 | BAIAP2L2 |
| 209 | scaffold_28 | 22472132 | -6.25 | BAIAP2L2 |
| 210 | scaffold_28 | 26831468 | -17.93 | PACSIN2 |
| 211 | scaffold_28 | 27122030 | 9.52 | |
| 212 | scaffold_28 | 27122047 | 5.65 | |
| 213 | scaffold_3 | 3415512 | 9.88 | |
| 214 | scaffold_3 | 3415516 | -5.25 | |
| 215 | scaffold_3 | 3434523 | -10.00 | |
| 216 | scaffold_3 | 3434564 | -7.50 | |
| 217 | scaffold_3 | 3434828 | -6.36 | |
| 218 | scaffold_3 | 3482571 | 8.64 | |
| 219 | scaffold_3 | 3482824 | -15.14 | |
| 220 | scaffold_304 | 317594 | -9.70 | |
| 221 | scaffold_304 | 317603 | -5.51 | |
| 222 | scaffold_304 | 317650 | -5.26 | |
| 223 | scaffold_309 | 242033 | -21.79 | |
| 224 | scaffold_309 | 242048 | 11.85 | |
| 225 | scaffold_309 | 242298 | 8.82 | |
| 226 | scaffold_32 | 21988863 | -9.71 | |
| 227 | scaffold_32 | 21988869 | -9.05 | |
| 228 | scaffold_32 | 21988910 | -7.48 | |
| 229 | scaffold_32 | 21988924 | -9.49 | |
| 230 | scaffold_32 | 21990745 | -11.78 | |

| | | | | |
|---|---|---|---|---|
| 231 | scaffold_32 | 21990988 | -14.01 | |
| 232 | scaffold_32 | 21990994 | -17.55 | |
| 233 | scaffold_32 | 21991018 | -20.74 | |
| 234 | scaffold_32 | 21991094 | -5.89 | |
| 235 | scaffold_32 | 21997047 | -7.32 | |
| 236 | scaffold_32 | 22002541 | 5.39 | |
| 237 | scaffold_32 | 22013674 | 7.93 | |
| 238 | scaffold_32 | 22013745 | -6.90 | |
| 239 | scaffold_37 | 20578870 | -5.37 | |
| 240 | scaffold_37 | 20578934 | -6.25 | |
| 241 | scaffold_37 | 20598529 | 7.63 | |
| 242 | scaffold_38 | 2547860 | 5.17 | FARS2 |
| 243 | scaffold_38 | 2547891 | 5.06 | FARS2 |
| 244 | scaffold_38 | 4432803 | -7.31 | TXNDC5 |
| 245 | scaffold_38 | 4432832 | 11.10 | TXNDC5 |
| 246 | scaffold_38 | 4433701 | 5.83 | TXNDC5 |
| 247 | scaffold_39 | 11246291 | -31.82 | PARD3 |
| 248 | scaffold_39 | 11246309 | -25.38 | PARD3 |
| 249 | scaffold_39 | 15067966 | 10.99 | |
| 250 | scaffold_39 | 15067967 | -8.85 | |
| 251 | scaffold_42 | 5821615 | -6.42 | |
| 252 | scaffold_42 | 5821667 | -16.07 | |
| 253 | scaffold_42 | 5821717 | 16.43 | |
| 254 | scaffold_42 | 5825126 | 15.55 | ARL2 |
| 255 | scaffold_42 | 5825142 | -5.01 | ARL2 |
| 256 | scaffold_42 | 5825157 | 5.45 | ARL2 |
| 257 | scaffold_42 | 5826412 | -10.71 | ARL2 |
| 258 | scaffold_42 | 6052456 | 10.95 | CDC42EP2 |
| 259 | scaffold_42 | 6052504 | 9.52 | CDC42EP2 |
| 260 | scaffold_42 | 6052506 | 8.97 | CDC42EP2 |
| 261 | scaffold_42 | 6052508 | 14.32 | CDC42EP2 |
| 262 | scaffold_42 | 6052622 | -22.43 | CDC42EP2 |
| 263 | scaffold_42 | 6052638 | -9.28 | CDC42EP2 |
| 264 | scaffold_42 | 6052734 | -15.30 | CDC42EP2 |
| 265 | scaffold_42 | 6052887 | -6.93 | CDC42EP2 |
| 266 | scaffold_42 | 6052908 | 23.52 | CDC42EP2 |
| 267 | scaffold_42 | 6053215 | -30.82 | CDC42EP2 |
| 268 | scaffold_42 | 6056824 | -6.08 | CDC42EP2 |
| 269 | scaffold_42 | 6096998 | 9.50 | SLC25A45 |
| 270 | scaffold_42 | 6097359 | 11.76 | SLC25A45 |
| 271 | scaffold_42 | 6137631 | 5.83 | |
| 272 | scaffold_434 | 22073 | 5.34 | |
| 273 | scaffold_46 | 616270 | 14.59 | |
| 274 | scaffold_46 | 616273 | 22.92 | |
| 275 | scaffold_46 | 616276 | -20.00 | |
| 276 | scaffold_46 | 616337 | 8.33 | |
| 277 | scaffold_46 | 616345 | -5.23 | |
| 278 | scaffold_46 | 616362 | -20.84 | |
| 279 | scaffold_46 | 616366 | -5.00 | |
| 280 | scaffold_46 | 616430 | -7.11 | |
| 281 | scaffold_5 | 47426708 | 11.90 | |
| 282 | scaffold_509 | 65292 | -9.05 | |
| 283 | scaffold_509 | 65302 | -29.46 | |
| 284 | scaffold_509 | 65322 | 12.50 | |
| 285 | scaffold_509 | 65458 | -5.68 | |
| 286 | scaffold_511 | 116633 | 8.57 | KDM4B |
| 287 | scaffold_511 | 116638 | 5.21 | KDM4B |
| 288 | scaffold_511 | 116639 | 5.11 | KDM4B |

| | | | | |
|---|---|---|---|---|
| 289 | scaffold_511 | 116646 | -6.56 | KDM4B |
| 290 | scaffold_511 | 133859 | -6.25 | |
| 291 | scaffold_53 | 2049886 | -6.37 | |
| 292 | scaffold_53 | 2049889 | -5.29 | |
| 293 | scaffold_53 | 2049890 | -9.03 | |
| 294 | scaffold_53 | 5918184 | -8.41 | |
| 295 | scaffold_53 | 6034626 | -8.47 | CA10 |
| 296 | scaffold_53 | 6034671 | -16.38 | CA10 |
| 297 | scaffold_53 | 6044291 | -7.19 | CA10 |
| 298 | scaffold_53 | 6062219 | -7.91 | CA10 |
| 299 | scaffold_53 | 6130993 | 5.45 | CA10 |
| 300 | scaffold_53 | 6130997 | -5.59 | CA10 |
| 301 | scaffold_53 | 6162318 | 7.66 | CA10 |
| 302 | scaffold_53 | 6162946 | 17.65 | CA10 |
| 303 | scaffold_53 | 6170257 | -10.00 | CA10 |
| 304 | scaffold_53 | 6184604 | -6.52 | CA10 |
| 305 | scaffold_53 | 6270377 | 9.26 | CA10 |
| 306 | scaffold_53 | 6270684 | 7.49 | CA10 |
| 307 | scaffold_53 | 6272353 | -8.17 | CA10 |
| 308 | scaffold_53 | 6300313 | -13.31 | CA10 |
| 309 | scaffold_53 | 6300773 | 7.23 | CA10 |
| 310 | scaffold_53 | 6342952 | 8.17 | CA10 |
| 311 | scaffold_53 | 6354310 | -12.54 | CA10 |
| 312 | scaffold_53 | 6354477 | 5.76 | CA10 |
| 313 | scaffold_53 | 6354521 | 7.35 | CA10 |
| 314 | scaffold_53 | 6355748 | 17.79 | CA10 |
| 315 | scaffold_53 | 6355760 | -10.58 | CA10 |
| 316 | scaffold_53 | 6359467 | -11.11 | CA10 |
| 317 | scaffold_53 | 6359470 | -14.81 | CA10 |
| 318 | scaffold_53 | 6359704 | -15.76 | CA10 |
| 319 | scaffold_53 | 6359759 | -15.33 | CA10 |
| 320 | scaffold_53 | 6371415 | -5.67 | CA10 |
| 321 | scaffold_53 | 6371662 | 10.69 | CA10 |
| 322 | scaffold_53 | 6445603 | 11.11 | CA10 |
| 323 | scaffold_53 | 6445948 | 10.00 | CA10 |
| 324 | scaffold_53 | 6449659 | -6.99 | CA10 |
| 325 | scaffold_53 | 6449789 | -15.00 | CA10 |
| 326 | scaffold_53 | 6449790 | 5.18 | CA10 |
| 327 | scaffold_53 | 6449827 | -21.30 | CA10 |
| 328 | scaffold_53 | 6452553 | -7.11 | CA10 |
| 329 | scaffold_53 | 6461759 | 6.07 | CA10 |
| 330 | scaffold_53 | 6461832 | -17.49 | CA10 |
| 331 | scaffold_53 | 6464276 | -6.55 | CA10 |
| 332 | scaffold_53 | 6475532 | -8.03 | |
| 333 | scaffold_53 | 6475666 | 12.27 | |
| 334 | scaffold_53 | 6522656 | -5.39 | |
| 335 | scaffold_53 | 6525402 | 19.05 | |
| 336 | scaffold_53 | 6525403 | -7.14 | |
| 337 | scaffold_53 | 6525465 | -10.13 | |
| 338 | scaffold_53 | 6525786 | 10.66 | |
| 339 | scaffold_53 | 6525792 | -5.37 | |
| 340 | scaffold_53 | 6525839 | -7.20 | |
| 341 | scaffold_53 | 6525904 | -8.86 | |
| 342 | scaffold_53 | 6525905 | 10.12 | |
| 343 | scaffold_53 | 6526027 | 8.95 | |
| 344 | scaffold_53 | 6526087 | 7.06 | |
| 345 | scaffold_53 | 6677362 | 8.37 | |
| 346 | scaffold_53 | 6680218 | -32.29 | |

| | | | | |
|---|---|---|---|---|
| 347 | scaffold_53 | 6680262 | -10.16 | |
| 348 | scaffold_53 | 6680490 | 27.13 | |
| 349 | scaffold_53 | 6680536 | 19.03 | |
| 350 | scaffold_53 | 6681572 | 7.33 | |
| 351 | scaffold_53 | 6681603 | -15.68 | |
| 352 | scaffold_53 | 6681604 | -20.58 | |
| 353 | scaffold_53 | 6681742 | -19.81 | |
| 354 | scaffold_53 | 6687281 | -8.17 | |
| 355 | scaffold_53 | 6687292 | -20.77 | |
| 356 | scaffold_53 | 6687538 | -14.74 | |
| 357 | scaffold_53 | 6691573 | 6.25 | |
| 358 | scaffold_53 | 6697469 | -10.42 | |
| 359 | scaffold_53 | 6705034 | -6.28 | |
| 360 | scaffold_53 | 6705098 | 6.66 | |
| 361 | scaffold_53 | 6710048 | 15.95 | |
| 362 | scaffold_53 | 6713876 | 5.89 | |
| 363 | scaffold_53 | 6713879 | -5.47 | |
| 364 | scaffold_53 | 6728527 | 10.12 | UTP18 |
| 365 | scaffold_53 | 6728554 | 10.12 | UTP18 |
| 366 | scaffold_53 | 6732958 | 5.06 | UTP18 |
| 367 | scaffold_53 | 6733144 | -8.21 | UTP18 |
| 368 | scaffold_53 | 6738976 | 6.13 | UTP18 |
| 369 | scaffold_53 | 6754504 | 9.34 | UTP18 |
| 370 | scaffold_53 | 6754574 | -9.40 | UTP18 |
| 371 | scaffold_53 | 6777315 | 7.80 | MBTD1 |
| 372 | scaffold_53 | 6785080 | 6.62 | MBTD1 |
| 373 | scaffold_53 | 6785364 | -6.28 | MBTD1 |
| 374 | scaffold_53 | 6787804 | -6.06 | MBTD1 |
| 375 | scaffold_53 | 6795687 | -6.91 | MBTD1 |
| 376 | scaffold_53 | 6803960 | -5.51 | MBTD1 |
| 377 | scaffold_53 | 6808561 | 10.39 | MBTD1 |
| 378 | scaffold_53 | 6808583 | 6.26 | MBTD1 |
| 379 | scaffold_53 | 6811289 | -12.95 | MBTD1 |
| 380 | scaffold_53 | 6811333 | -15.62 | MBTD1 |
| 381 | scaffold_53 | 6811342 | -7.70 | MBTD1 |
| 382 | scaffold_53 | 6811349 | -31.16 | MBTD1 |
| 383 | scaffold_53 | 6811377 | -5.75 | MBTD1 |
| 384 | scaffold_53 | 6811420 | -8.18 | MBTD1 |
| 385 | scaffold_53 | 6812984 | 6.96 | NME2 |
| 386 | scaffold_53 | 6813049 | -7.07 | NME2 |
| 387 | scaffold_53 | 6816414 | -8.91 | NME2 |
| 388 | scaffold_53 | 6816429 | -5.26 | NME2 |
| 389 | scaffold_53 | 6816713 | -8.36 | NME2 |
| 390 | scaffold_53 | 6816715 | -5.30 | NME2 |
| 391 | scaffold_53 | 6816719 | 8.86 | NME2 |
| 392 | scaffold_53 | 6816726 | -9.20 | NME2 |
| 393 | scaffold_53 | 6816823 | -13.05 | NME2 |
| 394 | scaffold_53 | 6817911 | -8.27 | |
| 395 | scaffold_53 | 6818091 | 13.92 | NME1 |
| 396 | scaffold_53 | 6822588 | -6.68 | |
| 397 | scaffold_53 | 6824900 | -10.14 | |
| 398 | scaffold_53 | 6828383 | -10.16 | |
| 399 | scaffold_53 | 6834866 | 18.57 | |
| 400 | scaffold_53 | 6835582 | -6.40 | |
| 401 | scaffold_53 | 6835594 | -12.18 | |
| 402 | scaffold_53 | 6836807 | -18.98 | |
| 403 | scaffold_53 | 6837017 | 6.67 | |
| 404 | scaffold_53 | 6842804 | -7.27 | |

| | | | | |
|---|---|---|---|---|
| 405 | scaffold_53 | 6842817 | -16.79 | |
| 406 | scaffold_53 | 6842824 | -10.53 | |
| 407 | scaffold_53 | 6843198 | -7.36 | |
| 408 | scaffold_53 | 6843199 | -14.29 | |
| 409 | scaffold_53 | 6843205 | -8.18 | |
| 410 | scaffold_53 | 6843245 | -13.93 | |
| 411 | scaffold_53 | 6846525 | -10.53 | SPAG9 |
| 412 | scaffold_53 | 6853796 | -5.05 | SPAG9 |
| 413 | scaffold_53 | 6854022 | 8.02 | SPAG9 |
| 414 | scaffold_53 | 6854023 | -9.62 | SPAG9 |
| 415 | scaffold_53 | 6859814 | 5.68 | SPAG9 |
| 416 | scaffold_53 | 6859815 | 11.11 | SPAG9 |
| 417 | scaffold_53 | 6859840 | 7.93 | SPAG9 |
| 418 | scaffold_53 | 6859878 | 9.94 | SPAG9 |
| 419 | scaffold_53 | 6859879 | 21.70 | SPAG9 |
| 420 | scaffold_53 | 6859886 | 14.34 | SPAG9 |
| 421 | scaffold_53 | 6859889 | -8.72 | SPAG9 |
| 422 | scaffold_53 | 6859928 | -26.19 | SPAG9 |
| 423 | scaffold_53 | 6859945 | -7.80 | SPAG9 |
| 424 | scaffold_53 | 6859952 | 5.12 | SPAG9 |
| 425 | scaffold_53 | 6860003 | 35.00 | SPAG9 |
| 426 | scaffold_53 | 6860022 | 15.00 | SPAG9 |
| 427 | scaffold_53 | 6860024 | 25.00 | SPAG9 |
| 428 | scaffold_53 | 6860031 | 15.00 | SPAG9 |
| 429 | scaffold_53 | 6860060 | -9.47 | SPAG9 |
| 430 | scaffold_53 | 6861837 | -11.38 | SPAG9 |
| 431 | scaffold_53 | 6862130 | 16.67 | SPAG9 |
| 432 | scaffold_53 | 6863186 | 11.06 | SPAG9 |
| 433 | scaffold_53 | 6863236 | -8.33 | SPAG9 |
| 434 | scaffold_53 | 6863243 | 18.33 | SPAG9 |
| 435 | scaffold_53 | 6863244 | -13.83 | SPAG9 |
| 436 | scaffold_53 | 6863251 | 23.20 | SPAG9 |
| 437 | scaffold_53 | 6874461 | 5.95 | SPAG9 |
| 438 | scaffold_53 | 6874498 | 5.27 | SPAG9 |
| 439 | scaffold_53 | 6874517 | 6.65 | SPAG9 |
| 440 | scaffold_53 | 6874644 | -6.37 | SPAG9 |
| 441 | scaffold_53 | 6874645 | -17.34 | SPAG9 |
| 442 | scaffold_53 | 6874714 | 5.47 | SPAG9 |
| 443 | scaffold_53 | 6882412 | 9.15 | SPAG9 |
| 444 | scaffold_53 | 6882642 | 5.17 | SPAG9 |
| 445 | scaffold_53 | 6887921 | -6.01 | SPAG9 |
| 446 | scaffold_53 | 6890298 | -5.91 | SPAG9 |
| 447 | scaffold_53 | 6890527 | 5.67 | SPAG9 |
| 448 | scaffold_53 | 6891496 | -5.56 | SPAG9 |
| 449 | scaffold_53 | 6891747 | 16.00 | SPAG9 |
| 450 | scaffold_53 | 6891782 | 9.03 | SPAG9 |
| 451 | scaffold_53 | 6891798 | 5.32 | SPAG9 |
| 452 | scaffold_53 | 6895071 | 6.45 | SPAG9 |
| 453 | scaffold_53 | 6896837 | -9.82 | SPAG9 |
| 454 | scaffold_53 | 6899246 | 9.17 | SPAG9 |
| 455 | scaffold_53 | 6899278 | -10.87 | SPAG9 |
| 456 | scaffold_53 | 6899555 | 13.31 | SPAG9 |
| 457 | scaffold_53 | 6899590 | 6.37 | SPAG9 |
| 458 | scaffold_53 | 6900918 | -11.11 | SPAG9 |
| 459 | scaffold_53 | 6900965 | -7.83 | SPAG9 |
| 460 | scaffold_53 | 6904389 | -17.28 | SPAG9 |
| 461 | scaffold_53 | 6904393 | -11.03 | SPAG9 |
| 462 | scaffold_53 | 6904405 | -10.29 | SPAG9 |

| | | | | |
|---|---|---|---|---|
| 463 | scaffold_53 | 6904657 | -7.63 | SPAG9 |
| 464 | scaffold_53 | 6904671 | 10.53 | SPAG9 |
| 465 | scaffold_53 | 6904695 | 17.86 | SPAG9 |
| 466 | scaffold_53 | 6904720 | 8.10 | SPAG9 |
| 467 | scaffold_53 | 6908591 | 11.48 | SPAG9 |
| 468 | scaffold_53 | 6915620 | 11.31 | SPAG9 |
| 469 | scaffold_53 | 6915848 | -17.31 | SPAG9 |
| 470 | scaffold_53 | 6915864 | -17.73 | SPAG9 |
| 471 | scaffold_53 | 6917942 | 9.52 | SPAG9 |
| 472 | scaffold_53 | 6917998 | 18.10 | SPAG9 |
| 473 | scaffold_53 | 6918094 | -14.29 | SPAG9 |
| 474 | scaffold_53 | 6918126 | -9.52 | SPAG9 |
| 475 | scaffold_53 | 6918132 | 17.14 | SPAG9 |
| 476 | scaffold_53 | 6918432 | -8.00 | SPAG9 |
| 477 | scaffold_53 | 6921919 | -5.98 | SPAG9 |
| 478 | scaffold_53 | 6921964 | 6.70 | SPAG9 |
| 479 | scaffold_53 | 6922046 | 35.78 | SPAG9 |
| 480 | scaffold_53 | 6923835 | -7.94 | SPAG9 |
| 481 | scaffold_53 | 6923970 | -7.29 | SPAG9 |
| 482 | scaffold_53 | 6927508 | 7.44 | SPAG9 |
| 483 | scaffold_53 | 6927516 | 25.38 | SPAG9 |
| 484 | scaffold_53 | 6927551 | 14.81 | SPAG9 |
| 485 | scaffold_53 | 6927552 | 6.15 | SPAG9 |
| 486 | scaffold_53 | 6927567 | 15.50 | SPAG9 |
| 487 | scaffold_53 | 6927575 | -5.63 | SPAG9 |
| 488 | scaffold_53 | 6927581 | -5.38 | SPAG9 |
| 489 | scaffold_53 | 6927582 | -7.17 | SPAG9 |
| 490 | scaffold_53 | 6930829 | 8.33 | SPAG9 |
| 491 | scaffold_53 | 6930841 | 7.58 | SPAG9 |
| 492 | scaffold_53 | 6930842 | 20.00 | SPAG9 |
| 493 | scaffold_53 | 6930853 | -9.09 | SPAG9 |
| 494 | scaffold_53 | 6930854 | 8.33 | SPAG9 |
| 495 | scaffold_53 | 6930868 | 8.18 | SPAG9 |
| 496 | scaffold_53 | 6930869 | 7.69 | SPAG9 |
| 497 | scaffold_53 | 6930887 | -6.67 | SPAG9 |
| 498 | scaffold_53 | 6930892 | 7.69 | SPAG9 |
| 499 | scaffold_53 | 6933115 | 27.27 | SPAG9 |
| 500 | scaffold_53 | 6933142 | 9.09 | SPAG9 |
| 501 | scaffold_53 | 6933398 | 22.86 | SPAG9 |
| 502 | scaffold_53 | 6933442 | -7.14 | SPAG9 |
| 503 | scaffold_53 | 6934996 | -11.40 | SPAG9 |
| 504 | scaffold_53 | 6935005 | -5.85 | SPAG9 |
| 505 | scaffold_53 | 6935017 | -6.14 | SPAG9 |
| 506 | scaffold_53 | 6935057 | -7.08 | SPAG9 |
| 507 | scaffold_53 | 6935295 | 5.97 | SPAG9 |
| 508 | scaffold_53 | 6935296 | 11.11 | SPAG9 |
| 509 | scaffold_53 | 6935337 | -5.69 | SPAG9 |
| 510 | scaffold_53 | 6935359 | -6.14 | SPAG9 |
| 511 | scaffold_53 | 6935487 | -9.30 | SPAG9 |
| 512 | scaffold_53 | 6935691 | 9.76 | SPAG9 |
| 513 | scaffold_53 | 6935720 | -9.56 | SPAG9 |
| 514 | scaffold_53 | 6935730 | -6.45 | SPAG9 |
| 515 | scaffold_53 | 6938170 | -6.81 | SPAG9 |
| 516 | scaffold_53 | 6941163 | -15.18 | SPAG9 |
| 517 | scaffold_53 | 6941677 | -7.69 | SPAG9 |
| 518 | scaffold_53 | 6945205 | 9.42 | SPAG9 |
| 519 | scaffold_53 | 6945245 | 6.22 | SPAG9 |
| 520 | scaffold_53 | 6945429 | 10.53 | SPAG9 |

| | | | | |
|---|---|---|---|---|
| 521 | scaffold_53 | 6945487 | 6.54 | SPAG9 |
| 522 | scaffold_53 | 6945869 | 14.62 | SPAG9 |
| 523 | scaffold_53 | 6945873 | 13.16 | SPAG9 |
| 524 | scaffold_53 | 6946899 | 9.35 | SPAG9 |
| 525 | scaffold_53 | 6946970 | -6.67 | SPAG9 |
| 526 | scaffold_53 | 6947025 | 20.94 | SPAG9 |
| 527 | scaffold_53 | 6947068 | 20.39 | SPAG9 |
| 528 | scaffold_53 | 6947075 | 29.08 | SPAG9 |
| 529 | scaffold_53 | 6947078 | 18.26 | SPAG9 |
| 530 | scaffold_53 | 6947081 | 14.79 | SPAG9 |
| 531 | scaffold_53 | 6947144 | -8.08 | SPAG9 |
| 532 | scaffold_53 | 6947172 | -9.54 | SPAG9 |
| 533 | scaffold_53 | 6947175 | -8.47 | SPAG9 |
| 534 | scaffold_53 | 6947197 | -6.99 | SPAG9 |
| 535 | scaffold_53 | 6949766 | -7.78 | SPAG9 |
| 536 | scaffold_53 | 6950099 | 15.91 | SPAG9 |
| 537 | scaffold_53 | 6950101 | 12.50 | SPAG9 |
| 538 | scaffold_53 | 6950107 | -5.09 | SPAG9 |
| 539 | scaffold_53 | 6950108 | -5.68 | SPAG9 |
| 540 | scaffold_53 | 6950151 | -5.68 | SPAG9 |
| 541 | scaffold_53 | 6950168 | 5.60 | SPAG9 |
| 542 | scaffold_53 | 6950173 | -5.36 | SPAG9 |
| 543 | scaffold_53 | 6950228 | 14.97 | SPAG9 |
| 544 | scaffold_53 | 6952661 | 15.64 | SPAG9 |
| 545 | scaffold_53 | 6952678 | 22.85 | SPAG9 |
| 546 | scaffold_53 | 6952711 | 19.22 | SPAG9 |
| 547 | scaffold_53 | 6952825 | -9.09 | SPAG9 |
| 548 | scaffold_53 | 6952832 | -6.64 | SPAG9 |
| 549 | scaffold_53 | 6952844 | -11.32 | SPAG9 |
| 550 | scaffold_53 | 6952859 | -18.06 | SPAG9 |
| 551 | scaffold_53 | 6952864 | -7.74 | SPAG9 |
| 552 | scaffold_53 | 6952881 | -13.50 | SPAG9 |
| 553 | scaffold_53 | 6952897 | -13.22 | SPAG9 |
| 554 | scaffold_53 | 6952898 | -6.52 | SPAG9 |
| 555 | scaffold_53 | 6952908 | -10.34 | SPAG9 |
| 556 | scaffold_53 | 6952920 | -11.38 | SPAG9 |
| 557 | scaffold_53 | 6952931 | -15.40 | SPAG9 |
| 558 | scaffold_53 | 6952933 | -17.18 | SPAG9 |
| 559 | scaffold_53 | 6952957 | -20.96 | SPAG9 |
| 560 | scaffold_53 | 6952964 | -16.00 | SPAG9 |
| 561 | scaffold_53 | 6952967 | -8.96 | SPAG9 |
| 562 | scaffold_53 | 6953101 | -5.07 | SPAG9 |
| 563 | scaffold_53 | 6953102 | 13.84 | SPAG9 |
| 564 | scaffold_53 | 6953158 | 13.75 | |
| 565 | scaffold_53 | 6953168 | -12.07 | |
| 566 | scaffold_53 | 6953353 | -19.48 | |
| 567 | scaffold_53 | 6953569 | -5.52 | |
| 568 | scaffold_53 | 6953676 | 5.68 | |
| 569 | scaffold_53 | 6953722 | 8.00 | |
| 570 | scaffold_53 | 6953769 | 11.67 | |
| 571 | scaffold_53 | 6954235 | -5.59 | |
| 572 | scaffold_53 | 6954367 | -9.02 | |
| 573 | scaffold_53 | 6954388 | -8.42 | |
| 574 | scaffold_53 | 6954399 | -10.68 | |
| 575 | scaffold_53 | 6954414 | -5.21 | |
| 576 | scaffold_53 | 6955972 | 14.29 | |
| 577 | scaffold_53 | 6956005 | 22.73 | |
| 578 | scaffold_53 | 6956395 | -10.00 | |

| | | | |
|---|---|---|---|
| 579 | scaffold_53 | 6956787 | 11.43 |
| 580 | scaffold_53 | 6958311 | 29.08 |
| 581 | scaffold_53 | 6958361 | -7.25 |
| 582 | scaffold_53 | 6958384 | -11.81 |
| 583 | scaffold_53 | 6958457 | 6.56 |
| 584 | scaffold_53 | 6958743 | 6.10 |
| 585 | scaffold_53 | 6958751 | 8.65 |
| 586 | scaffold_53 | 6958779 | 18.82 |
| 587 | scaffold_53 | 6958792 | 10.83 |
| 588 | scaffold_53 | 6958793 | 9.22 |
| 589 | scaffold_53 | 6958812 | 6.81 |
| 590 | scaffold_53 | 6958827 | -10.61 |
| 591 | scaffold_53 | 6958834 | -5.40 |
| 592 | scaffold_53 | 6958873 | 31.82 |
| 593 | scaffold_53 | 6958877 | 23.64 |
| 594 | scaffold_53 | 6961757 | -7.44 |
| 595 | scaffold_53 | 6961887 | 7.14 |
| 596 | scaffold_53 | 6961893 | -33.77 |
| 597 | scaffold_53 | 6961946 | -16.82 |
| 598 | scaffold_53 | 6961947 | 17.67 |
| 599 | scaffold_53 | 6961979 | -11.00 |
| 600 | scaffold_53 | 6961988 | -13.40 |
| 601 | scaffold_53 | 6962101 | 33.52 |
| 602 | scaffold_53 | 6962158 | 21.25 |
| 603 | scaffold_53 | 6962325 | 12.34 |
| 604 | scaffold_53 | 6962346 | 18.33 |
| 605 | scaffold_53 | 6962393 | 20.00 |
| 606 | scaffold_53 | 6962420 | 20.96 |
| 607 | scaffold_53 | 6962694 | 9.75 |
| 608 | scaffold_53 | 6964594 | -38.46 |
| 609 | scaffold_53 | 6964603 | -27.47 |
| 610 | scaffold_53 | 6964671 | 6.77 |
| 611 | scaffold_53 | 6965557 | -5.56 |
| 612 | scaffold_53 | 6965579 | 8.30 |
| 613 | scaffold_53 | 6965588 | 7.92 |
| 614 | scaffold_53 | 6965621 | 5.33 |
| 615 | scaffold_53 | 6965646 | 10.58 |
| 616 | scaffold_53 | 6965678 | -12.98 |
| 617 | scaffold_53 | 6965682 | 14.90 |
| 618 | scaffold_53 | 6965731 | 32.38 |
| 619 | scaffold_53 | 6965747 | 25.71 |
| 620 | scaffold_53 | 6965760 | 41.43 |
| 621 | scaffold_53 | 6965782 | 22.86 |
| 622 | scaffold_53 | 6965786 | 16.67 |
| 623 | scaffold_53 | 6965792 | 6.19 |
| 624 | scaffold_53 | 6965942 | -6.54 |
| 625 | scaffold_53 | 6966868 | -11.43 |
| 626 | scaffold_53 | 6966928 | -9.23 |
| 627 | scaffold_53 | 6967180 | 5.05 |
| 628 | scaffold_53 | 6967523 | -7.42 |
| 629 | scaffold_53 | 6967550 | 11.51 |
| 630 | scaffold_53 | 6968048 | -8.09 |
| 631 | scaffold_53 | 6968049 | -5.09 |
| 632 | scaffold_53 | 6968089 | 16.77 |
| 633 | scaffold_53 | 6968212 | 8.33 |
| 634 | scaffold_53 | 6968224 | -8.33 |
| 635 | scaffold_53 | 6968268 | -8.67 |
| 636 | scaffold_53 | 6968315 | -5.54 |

| | | | | |
|---|---|---|---|---|
| 637 | scaffold_53 | 6972893 | 5.20 | |
| 638 | scaffold_53 | 6972930 | -6.52 | |
| 639 | scaffold_53 | 6972938 | -8.60 | |
| 640 | scaffold_53 | 6972971 | 11.04 | |
| 641 | scaffold_53 | 6973057 | 6.53 | |
| 642 | scaffold_53 | 6973086 | 8.21 | |
| 643 | scaffold_53 | 6973121 | -6.26 | |
| 644 | scaffold_53 | 6973136 | 8.70 | |
| 645 | scaffold_53 | 6973240 | -11.35 | |
| 646 | scaffold_53 | 6973300 | 9.69 | |
| 647 | scaffold_53 | 6973381 | -13.56 | |
| 648 | scaffold_53 | 6973390 | -5.65 | |
| 649 | scaffold_53 | 6973419 | -11.17 | |
| 650 | scaffold_53 | 6973738 | 19.46 | |
| 651 | scaffold_53 | 6973781 | 5.17 | |
| 652 | scaffold_53 | 6973807 | 6.38 | |
| 653 | scaffold_53 | 6973812 | -5.00 | |
| 654 | scaffold_53 | 6973819 | -10.00 | |
| 655 | scaffold_53 | 6973830 | -7.69 | |
| 656 | scaffold_53 | 6973834 | -7.89 | |
| 657 | scaffold_53 | 6973846 | -7.40 | |
| 658 | scaffold_53 | 6973892 | -17.48 | |
| 659 | scaffold_53 | 6973981 | 8.40 | |
| 660 | scaffold_53 | 6974032 | -6.60 | |
| 661 | scaffold_53 | 6974046 | 11.76 | |
| 662 | scaffold_53 | 6974062 | -12.30 | |
| 663 | scaffold_53 | 6974089 | -6.67 | |
| 664 | scaffold_53 | 6974194 | -5.13 | |
| 665 | scaffold_53 | 6975938 | -13.04 | |
| 666 | scaffold_53 | 6976098 | -16.85 | |
| 667 | scaffold_53 | 6976157 | -17.49 | |
| 668 | scaffold_53 | 6977162 | -6.31 | |
| 669 | scaffold_53 | 6977209 | -9.83 | |
| 670 | scaffold_53 | 6977494 | 5.51 | |
| 671 | scaffold_53 | 6977495 | -7.77 | |
| 672 | scaffold_53 | 6977705 | -16.65 | |
| 673 | scaffold_53 | 6977755 | -10.74 | |
| 674 | scaffold_53 | 6977756 | 8.61 | |
| 675 | scaffold_53 | 6977785 | -10.96 | |
| 676 | scaffold_55 | 292366 | 12.85 | ADCY2 |
| 677 | scaffold_55 | 292367 | -16.67 | ADCY2 |
| 678 | scaffold_55 | 292382 | -6.71 | ADCY2 |
| 679 | scaffold_55 | 8480831 | -16.83 | MYO10 |
| 680 | scaffold_56 | 7511258 | -14.29 | |
| 681 | scaffold_56 | 7511273 | 7.27 | |
| 682 | scaffold_56 | 7511294 | -18.18 | |
| 683 | scaffold_56 | 7511308 | 5.11 | |
| 684 | scaffold_56 | 7511318 | 9.32 | |
| 685 | scaffold_56 | 7511325 | 10.66 | |
| 686 | scaffold_6 | 32457316 | -18.66 | |
| 687 | scaffold_6 | 32457329 | -5.74 | |
| 688 | scaffold_6 | 43172321 | 21.25 | |
| 689 | scaffold_6 | 43172339 | 31.34 | |
| 690 | scaffold_6 | 43172340 | 28.75 | |
| 691 | scaffold_6 | 43172359 | -10.77 | |
| 692 | scaffold_6 | 43172687 | -11.64 | |
| 693 | scaffold_61 | 10023665 | -12.63 | PFN1 |
| 694 | scaffold_61 | 10023666 | 32.14 | PFN1 |

| | | | | |
|---|---|---|---|---|
| 695 | scaffold_62 | 5834647 | 16.67 | WAPL |
| 696 | scaffold_62 | 5834648 | -10.89 | WAPL |
| 697 | scaffold_62 | 5837206 | -8.70 | WAPL |
| 698 | scaffold_62 | 5837492 | 8.51 | WAPL |
| 699 | scaffold_65 | 5437941 | -7.19 | |
| 700 | scaffold_65 | 5445131 | 10.80 | |
| 701 | scaffold_65 | 5447216 | 10.86 | |
| 702 | scaffold_65 | 5448403 | 7.04 | |
| 703 | scaffold_67 | 5551547 | -9.29 | |
| 704 | scaffold_67 | 9124061 | -5.41 | ADAM11 |
| 705 | scaffold_67 | 9124070 | -7.84 | ADAM11 |
| 706 | scaffold_67 | 9124079 | -6.14 | ADAM11 |
| 707 | scaffold_687 | 42370 | 7.02 | |
| 708 | scaffold_687 | 42382 | -5.98 | |
| 709 | scaffold_687 | 42396 | 14.24 | |
| 710 | scaffold_687 | 42402 | 17.56 | |
| 711 | scaffold_687 | 42406 | 10.34 | |
| 712 | scaffold_687 | 42412 | 16.73 | |
| 713 | scaffold_687 | 42460 | 5.50 | |
| 714 | scaffold_687 | 42515 | -10.01 | |
| 715 | scaffold_687 | 42540 | 14.88 | |
| 716 | scaffold_687 | 42553 | 10.14 | |
| 717 | scaffold_687 | 42567 | -6.77 | |
| 718 | scaffold_687 | 42586 | 15.54 | |
| 719 | scaffold_687 | 42626 | 11.58 | |
| 720 | scaffold_687 | 42681 | -9.80 | |
| 721 | scaffold_687 | 42697 | -11.94 | |
| 722 | scaffold_687 | 42752 | -9.22 | |
| 723 | scaffold_7 | 46862584 | 11.67 | FYCO1 |
| 724 | scaffold_7 | 58076344 | 5.85 | |
| 725 | scaffold_7 | 58076412 | 15.79 | |
| 726 | scaffold_7 | 58076493 | 7.04 | |
| 727 | scaffold_7 | 58076506 | -5.49 | |
| 728 | scaffold_7 | 58076527 | -7.97 | |
| 729 | scaffold_7 | 58076842 | -13.43 | |
| 730 | scaffold_70 | 7240731 | -5.56 | SYT3 |
| 731 | scaffold_72 | 3969892 | -27.65 | |
| 732 | scaffold_72 | 3969924 | 13.52 | |
| 733 | scaffold_72 | 3969966 | 14.34 | |
| 734 | scaffold_72 | 3969980 | 11.49 | |
| 735 | scaffold_72 | 3970124 | -5.03 | |
| 736 | scaffold_72 | 3970126 | -8.16 | |
| 737 | scaffold_72 | 3970129 | -16.27 | |
| 738 | scaffold_72 | 3970168 | 10.91 | |
| 739 | scaffold_72 | 3970180 | -23.64 | |
| 740 | scaffold_72 | 3971709 | -6.28 | |
| 741 | scaffold_72 | 3971752 | 14.73 | |
| 742 | scaffold_72 | 3977596 | 8.04 | |
| 743 | scaffold_72 | 3977900 | -36.11 | |
| 744 | scaffold_72 | 3977919 | 5.56 | |
| 745 | scaffold_72 | 3977922 | 11.11 | |
| 746 | scaffold_72 | 3977932 | -8.33 | |
| 747 | scaffold_72 | 3978080 | 6.06 | |
| 748 | scaffold_72 | 4939302 | -7.22 | |
| 749 | scaffold_72 | 4939303 | -19.29 | |
| 750 | scaffold_72 | 4939369 | -16.39 | |
| 751 | scaffold_72 | 4939415 | 5.09 | |
| 752 | scaffold_72 | 4943361 | -10.63 | |

| | | | | |
|---|---|---|---|---|
| 753 | scaffold_72 | 4943448 | 5.09 | |
| 754 | scaffold_72 | 4943921 | 9.09 | |
| 755 | scaffold_81 | 7197256 | -5.17 | |
| 756 | scaffold_81 | 7197270 | 15.00 | |
| 757 | scaffold_82 | 4072515 | -6.86 | LAMA1 |
| 758 | scaffold_85 | 562195 | -6.88 | EVC2 |
| 759 | scaffold_85 | 564384 | -8.55 | EVC2 |
| 760 | scaffold_85 | 564393 | -7.37 | EVC2 |
| 761 | scaffold_85 | 564480 | 5.76 | EVC2 |
| 762 | scaffold_85 | 564527 | -6.86 | EVC2 |
| 763 | scaffold_87 | 4373256 | 5.88 | ITGB5 |
| 764 | scaffold_9 | 5820541 | -8.43 | HAL |
| 765 | scaffold_9 | 5824998 | 5.48 | HAL |
| 766 | scaffold_9 | 6720445 | -14.64 | TMCC3 |
| 767 | scaffold_9 | 6720456 | -21.43 | TMCC3 |
| 768 | scaffold_9 | 6720630 | -28.57 | TMCC3 |
| 769 | scaffold_9 | 6720693 | -10.96 | TMCC3 |
| 770 | scaffold_9 | 6723112 | -5.15 | TMCC3 |
| 771 | scaffold_9 | 6723158 | 5.22 | TMCC3 |
| 772 | scaffold_90 | 6041529 | -13.73 | CLDN14 |
| 773 | scaffold_90 | 6041540 | -8.79 | CLDN14 |
| 774 | scaffold_90 | 6041545 | -40.00 | CLDN14 |
| 775 | scaffold_90 | 6041561 | -16.25 | CLDN14 |
| 776 | scaffold_912 | 12984 | -13.13 | |
