## Supplemental Table 10 for "Conserved DNA Methylation Signatures in The Prefrontal Cortex of Newborn and Juvenile Guinea Pigs Following Antenatal Corticosteroid Exposure"

**Supplemental Table 10. Gene Set Enrichment Analysis of DMCs identified in the PND14PFC following ACS exposure**

| | source | term_name | term_id | adjusted_p_value | negative_log10_of_adjusted_p_value | term_size | query_size | intersection_size | effective_domain_size | intersections | | | | | | |
|---|---|---|---|---|---|---|---|---|---|---|---|---|---|---|---|---|
| 1 | GO:MF | molecular_function | GO:0003674 | 0.00109947 | 2.95881586 | 15743 | 43 | 39 | 26855 | PARD3 | SORCS2 | CLDN14 | FGD2 | PACSIN2 | APBA2 | HMCN2 |
| 2 | GO:MF | protein binding | GO:0005515 | 0.00734869 | 2.13379 | 6695 | 29 | 18 | 26855 | PARD3 | SORCS2 | PACSIN2 | APBA2 | MYO10 | RIMS2 | NME1 |
| 3 | GO:MF | binding | GO:0005488 | 0.01877155 | 1.726499962 | 11662 | 42 | 31 | 26855 | PARD3 | SORCS2 | FGD2 | PACSIN2 | APBA2 | HMCN2 | MYO10 |
| 4 | GO:MF | lipid binding | GO:0008289 | 0.03272234 | 1.485155642 | 528 | 25 | 5 | 26855 | FGD2 | PACSIN2 | PFN1 | LAMA1 | BAIAP2L2 | | |
| 5 | GO:MF | nucleoside diphosphate kinase activity | GO:0004550 | 0.03670461 | 1.435279356 | 14 | 32 | 2 | 26855 | NME1 | NME2 | | | | | |
| 6 | GO:BP | regulation of cellular process | GO:0050794 | 0.00021855 | 3.660447535 | 8191 | 45 | 31 | 26855 | SORCS2 | FGD2 | PACSIN2 | APBA2 | MYO10 | RIMS2 | NME1 |
| 7 | GO:BP | regulation of biological process | GO:0050789 | 0.00222817 | 2.652052023 | 8994 | 45 | 31 | 26855 | SORCS2 | FGD2 | PACSIN2 | APBA2 | MYO10 | RIMS2 | NME1 |
| 8 | GO:BP | signaling | GO:0023052 | 0.00537762 | 2.269409766 | 4916 | 42 | 21 | 26855 | SORCS2 | FGD2 | PACSIN2 | APBA2 | MYO10 | RIMS2 | KCND3 |
| 9 | GO:BP | anterograde trans-synaptic signaling | GO:0098916 | 0.00769092 | 2.11402145 | 375 | 9 | 4 | 26855 | SORCS2 | | PACSIN2 | APBA2 | | RIMS2 | |
| 10 | GO:BP | chemical synaptic transmission | GO:0007268 | 0.00769092 | 2.11402145 | 375 | 9 | 4 | 26855 | SORCS2 | | PACSIN2 | APBA2 | | RIMS2 | |
| 11 | GO:BP | trans-synaptic signaling | GO:0099537 | 0.00810501 | 2.091246658 | 380 | 9 | 4 | 26855 | SORCS2 | | PACSIN2 | APBA2 | | RIMS2 | |
| 12 | GO:BP | biological regulation | GO:0065007 | 0.00834293 | 2.07868117 | 9500 | 45 | 31 | 26855 | SORCS2 | FGD2 | PACSIN2 | APBA2 | MYO10 | RIMS2 | NME1 |
| 13 | GO:BP | synaptic signaling | GO:0099536 | 0.00935265 | 2.029065498 | 394 | 9 | 4 | 26855 | SORCS2 | | PACSIN2 | APBA2 | | RIMS2 | |
| 14 | GO:BP | transport | GO:0006810 | 0.02404007 | 1.619064361 | 3252 | 15 | 9 | 26855 | SORCS2 | PACSIN2 | APBA2 | RIMS2 | NME1 | FYCO1 | SLC25A45 |
| 15 | GO:BP | cell communication | GO:0007154 | 0.02746667 | 1.561194023 | 4965 | 42 | 20 | 26855 | SORCS2 | FGD2 | PACSIN2 | APBA2 | MYO10 | RIMS2 | LAMA1 |
| 16 | GO:BP | modulation of chemical synaptic transmission | GO:0050804 | 0.02958416 | 1.52894079 | 258 | 6 | 3 | 26855 | SORCS2 | | PACSIN2 | APBA2 | | | |
| 17 | GO:BP | regulation of trans-synaptic signaling | GO:0099177 | 0.02992832 | 1.523917609 | 259 | 6 | 3 | 26855 | SORCS2 | | PACSIN2 | APBA2 | | | |
| 18 | GO:BP | positive regulation of cellular process | GO:0048522 | 0.03081051 | 1.511301163 | 3793 | 45 | 18 | 26855 | FGD2 | MYO10 | NME1 | FYCO1 | PFN1 | RBM10 | DOK7 |
| 19 | GO:BP | establishment of localization | GO:0051234 | 0.0334015 | 1.476234014 | 3384 | 15 | 9 | 26855 | SORCS2 | PACSIN2 | APBA2 | RIMS2 | NME1 | FYCO1 | SLC25A45 |
| 20 | GO:BP | UTP biosynthetic process | GO:0006228 | 0.04961936 | 1.304348863 | 7 | 32 | 2 | 26855 | NME1 | NME2 | | | | | |
| 21 | GO:CC | cell junction | GO:0030054 | 4.29E-07 | 6.368003456 | 1152 | 25 | 11 | 26855 | PARD3 | SORCS2 | CLDN14 | PACSIN2 | APBA2 | HMCN2 | RIMS2 |
| 22 | GO:CC | cellular anatomical entity | GO:0110165 | 5.4781E-05 | 4.261372752 | 17018 | 46 | 44 | 26855 | PARD3 | SORCS2 | CLDN14 | FGD2 | PACSIN2 | APBA2 | HMCN2 |
| 23 | GO:CC | membrane | GO:0016020 | 5.5452E-05 | 4.256083615 | 6572 | 42 | 26 | 26855 | PARD3 | SORCS2 | CLDN14 | PACSIN2 | MYO10 | RIMS2 | NME1 |
| 24 | GO:CC | cellular_component | GO:0005575 | 8.0603E-05 | 4.093647457 | 17180 | 46 | 44 | 26855 | PARD3 | SORCS2 | CLDN14 | FGD2 | PACSIN2 | APBA2 | HMCN2 |
| 25 | GO:CC | cell periphery | GO:0071944 | 0.00017793 | 3.749753413 | 3450 | 41 | 18 | 26855 | PARD3 | SORCS2 | CLDN14 | PACSIN2 | | HMCN2 | MYO10 |
| 26 | GO:CC | synapse | GO:0045202 | 0.00034773 | 3.458751643 | 689 | 16 | 6 | 26855 | SORCS2 | | PACSIN2 | APBA2 | | RIMS2 | KCND3 |
| 27 | GO:CC | cell-cell junction | GO:0005911 | 0.00179876 | 2.74502733 | 314 | 25 | 5 | 26855 | PARD3 | | CLDN14 | PACSIN2 | | | |
| 28 | GO:CC | recycling endosome membrane | GO:0055038 | 0.00397161 | 2.40103289 | 39 | 5 | 2 | 26855 | SORCS2 | PACSIN2 | | | | | |
| 29 | GO:CC | bicellular tight junction | GO:0005923 | 0.00422987 | 2.373673431 | 73 | 3 | 2 | 26855 | PARD3 | | CLDN14 | | | | |
| 30 | GO:CC | tight junction | GO:0070160 | 0.00521387 | 2.282840199 | 81 | 3 | 2 | 26855 | PARD3 | | CLDN14 | | | | |
| 31 | GO:CC | cytoplasmic vesicle | GO:0031410 | 0.00638978 | 2.194513828 | 1052 | 6 | 4 | 26855 | SORCS2 | FGD2 | PACSIN2 | APBA2 | | | |
| 32 | GO:CC | intracellular vesicle | GO:0097708 | 0.00643779 | 2.191262987 | 1054 | 6 | 4 | 26855 | SORCS2 | FGD2 | PACSIN2 | APBA2 | | | |
| 33 | GO:CC | apical junction complex | GO:0043296 | 0.00644347 | 2.190879928 | 90 | 3 | 2 | 26855 | PARD3 | | CLDN14 | | | | |
| 34 | GO:CC | endomembrane system | GO:0012505 | 0.00673073 | 2.17193773 | 2541 | 12 | 7 | 26855 | PARD3 | SORCS2 | FGD2 | PACSIN2 | APBA2 | TMCC3 | FYCO1 |
| 35 | GO:CC | anchoring junction | GO:0070161 | 0.00771393 | 2.112724559 | 430 | 5 | 3 | 26855 | PARD3 | | CLDN14 | PACSIN2 | | | |
| 36 | GO:CC | vesicle | GO:0031982 | 0.00904319 | 2.043678406 | 1149 | 6 | 4 | 26855 | SORCS2 | FGD2 | PACSIN2 | APBA2 | | | |
| 37 | GO:CC | endosome | GO:0005768 | 0.01485653 | 1.828082486 | 552 | 12 | 4 | 26855 | SORCS2 | FGD2 | PACSIN2 | FYCO1 | | | |
| 38 | GO:CC | basement membrane | GO:0005604 | 0.02061837 | 1.685745644 | 60 | 41 | 3 | 26855 | HMCN2 | LAMA1 | TIMP3 | | | | |
| 39 | GO:CC | PAR polarity complex | GO:0120157 | 0.02164943 | 1.664553604 | 3 | 1 | 1 | 26855 | PARD3 | | | | | | |
| 40 | GO:CC | recycling endosome | GO:0055037 | 0.03024169 | 1.519393954 | 107 | 5 | 2 | 26855 | SORCS2 | PACSIN2 | | | | | |
| 41 | GO:CC | cytoplasm | GO:0005737 | 0.0434658 | 1.361852372 | 7394 | 45 | 24 | 26855 | SORCS2 | FGD2 | PACSIN2 | APBA2 | HMCN2 | MYO10 | NME1 |
| 42 | GO:CC | cell leading edge | GO:0031252 | 0.04693422 | 1.328510421 | 258 | 32 | 4 | 26855 | FGD2 | NME1 | PLXND1 | NME2 | | | |
| 43 | HP | Overlapping fingers | HP:0010557 | 0.00951542 | 2.021571952 | 32 | 33 | 3 | 26855 | DOK7 | EVC2 | CHST11 | | | | |
| 44 | HP | Aplasia/Hypoplasia of the lungs | HP:0006703 | 0.0245332 | 1.610245887 | 142 | 31 | 4 | 26855 | RBM10 | DOK7 | PLXND1 | EVC2 | | | |

| | | | | | | | | | | | | | | | | | | |
|---|---|---|---|---|---|---|---|---|---|---|---|---|---|---|---|---|---|---|
| MYO10 | RIMS2 | NME1 | TMCC3 | FYCO1 | NIPAL3 | KCND3 | PFN1 | RBM10 | DOK7 | LAMA1 | GNG12 | ADAM11 | ITGB5 | CUX2 | BAIAP2L2 | PLXND1 | MBTD1 | PPP2R1B |
| TMCC3 | KCND3 | PFN1 | RBM10 | DOK7 | LAMA1 | GNG12 | BAIAP2L2 | PLXND1 | MBTD1 | PPP2R1B | | | | | | | | |
| RIMS2 | NME1 | TMCC3 | FYCO1 | KCND3 | PFN1 | RBM10 | DOK7 | LAMA1 | GNG12 | CUX2 | BAIAP2L2 | PLXND1 | MBTD1 | PPP2R1B | FARS2 | NME2 | UTP18 | KDM4B |
| FYCO1 | KCND3 | PFN1 | RBM10 | DOK7 | LAMA1 | GNG12 | ITGB5 | CUX2 | PLXND1 | SYT3 | MBTD1 | PPP2R1B | EVC2 | NME2 | CHST11 | ADCY2 | KDM4B | ATP6V1C2 |
| FYCO1 | KCND3 | PFN1 | RBM10 | DOK7 | LAMA1 | GNG12 | ITGB5 | CUX2 | PLXND1 | SYT3 | MBTD1 | PPP2R1B | EVC2 | NME2 | CHST11 | ADCY2 | KDM4B | ATP6V1C2 |
| | LAMA1 | GNG12 | ITGB5 | CUX2 | PLXND1 | PPP2R1B | EVC2 | NME2 | CHST11 | ADCY2 | ATP6V1C2 | CDC42EP2 | TIMP3 | SPAG9 | | | | |
| FYCO1 | KCND3 | PFN1 | RBM10 | DOK7 | LAMA1 | GNG12 | ITGB5 | CUX2 | PLXND1 | SYT3 | MBTD1 | PPP2R1B | EVC2 | NME2 | CHST11 | ADCY2 | KDM4B | ATP6V1C2 |
| NIPAL3 | KCND3 | | | | | | | | | | | | | | | | | |
| GNG12 | ITGB5 | CUX2 | PLXND1 | PPP2R1B | EVC2 | NME2 | CHST11 | ADCY2 | ATP6V1C2 | CDC42EP2 | TIMP3 | SPAG9 | | | | | | |
| CUX2 | SYT3 | MBTD1 | PPP2R1B | NME2 | ATP6V1C2 | CDC42EP2 | TIMP3 | SPAG9 | WAPL | ARL2 | | | | | | | | |
| NIPAL3 | KCND3 | | | | | | | | | | | | | | | | | |
| | KCND3 | PFN1 | LAMA1 | BAIAP2L2 | | | | | | | | | | | | | | |
| MYO10 | RIMS2 | NME1 | TMCC3 | FYCO1 | SLC25A45 | NIPAL3 | KCND3 | PFN1 | RBM10 | DOK7 | LAMA1 | GNG12 | ADAM11 | ITGB5 | CUX2 | BAIAP2L2 | PLXND1 | SYT3 |
| TMCC3 | SLC25A45 | NIPAL3 | KCND3 | LAMA1 | GNG12 | ADAM11 | ITGB5 | BAIAP2L2 | PLXND1 | SYT3 | PPP2R1B | EVC2 | CHST11 | ADCY2 | UTP18 | ATP6V1C2 | CDC42EP2 | SPAG9 |
| MYO10 | RIMS2 | NME1 | TMCC3 | FYCO1 | SLC25A45 | NIPAL3 | KCND3 | PFN1 | RBM10 | DOK7 | LAMA1 | GNG12 | ADAM11 | ITGB5 | CUX2 | BAIAP2L2 | PLXND1 | SYT3 |
| NME1 | KCND3 | PFN1 | LAMA1 | | GNG12 | ITGB5 | PLXND1 | EVC2 | NME2 | ADCY2 | CDC42EP2 | TIMP3 | | | | | | |
| PFN1 | | | | | | | | | | | | | | | | | | |
| | | | LAMA1 | BAIAP2L2 | | | | | | | | | | | | | | |
| TMCC3 | FYCO1 | PFN1 | DOK7 | BAIAP2L2 | SYT3 | FARS2 | NME2 | CHST11 | ADCY2 | TXNDC5 | KDM4B | CDC42EP2 | SPAG9 | HAL | WAPL | ARL2 | | |

| | | | | | | | | | | | | |
|---|---|---|---|---|---|---|---|---|---|---|---|---|
| FARS2 | NME2 | CHST11 | ADCY2 | UTP18 | TXNDC5 | KDM4B | CA10 | ATP6V1C2 | CDC42EP2 | TIMP3 | SPAG9 | HAL |
| CA10 | ATP6V1C2 | CDC42EP2 | TIMP3 | SPAG9 | | | | | | | | |

| | | | | |
|---|---|---|---|---|
| CDC42EP2 | TIMP3 | SPAG9 | WAPL | ARL2 |
| CDC42EP2 | TIMP3 | SPAG9 | WAPL | ARL2 |

| | | | | |
|---|---|---|---|---|
| CDC42EP2 | TIMP3 | SPAG9 | WAPL | ARL2 |

| | | | | | | | | | | | | | | | | | |
|---|---|---|---|---|---|---|---|---|---|---|---|---|---|---|---|---|---|
| MBTD1 | PPP2R1B | FARS2 | EVC2 | NME2 | CHST11 | ADCY2 | UTP18 | TXNDC5 | KDM4B | ATP6V1C2 | CDC42EP2 | TIMP3 | SPAG9 | HAL | WAPL | ARL2 | DSCAML1 |
| MBTD1 | PPP2R1B | FARS2 | EVC2 | NME2 | CHST11 | ADCY2 | UTP18 | TXNDC5 | KDM4B | ATP6V1C2 | CDC42EP2 | TIMP3 | SPAG9 | HAL | WAPL | ARL2 | DSCAML1 |
