## Supplemental Table 11 for "Conserved DNA Methylation Signatures in The Prefrontal Cortex of Newborn and Juvenile Guinea Pigs Following Antenatal Corticosteroid Exposure"

**Supplementary Table 11. List of DMCs identified within the binding site for PLAGL1 in the PND14PFC following ACS exposure**

| seqnames | start | MethDiff | geneName |
|---|---|---|---|
| scaffold_0 | 5589377 | -13.89 | |
| scaffold_0 | 5589421 | -15.85 | |
| scaffold_0 | 5589427 | -24.23 | |
| scaffold_0 | 5589428 | -28.50 | |
| scaffold_1 | 10198664 | 47.78 | |
| scaffold_10 | 48577576 | -17.86 | APBA2 |
| scaffold_104 | 2635279 | -7.50 | |
| scaffold_109 | 2866416 | -5.98 | PLXND1 |
| scaffold_109 | 2866417 | -10.59 | PLXND1 |
| scaffold_11 | 5059839 | 6.45 | |
| scaffold_11 | 5062946 | -7.20 | |
| scaffold_13 | 9282875 | -15.00 | |
| scaffold_132 | 672314 | -8.33 | RBM10 |
| scaffold_140 | 746133 | 8.25 | TIMP3 |
| scaffold_140 | 747309 | -8.24 | TIMP3 |
| scaffold_15 | 32550317 | 17.85 | |
| scaffold_15 | 32550395 | 31.85 | |
| scaffold_15 | 33089086 | 16.19 | |
| scaffold_15 | 34265179 | 18.37 | FGD2 |
| scaffold_155 | 1146733 | 5.87 | |
| scaffold_159 | 636982 | -11.67 | |
| scaffold_160 | 267844 | -10.71 | |
| scaffold_160 | 267890 | -7.14 | |
| scaffold_166 | 578641 | -23.56 | SORCS2 |
| scaffold_171 | 777299 | 6.13 | |
| scaffold_178 | 1472285 | 7.65 | |
| scaffold_178 | 1472287 | 8.12 | |
| scaffold_178 | 1472335 | 8.06 | |
| scaffold_18 | 2777029 | -9.61 | |
| scaffold_19 | 30378554 | -8.80 | DSCAML1 |
| scaffold_19 | 36033256 | -18.14 | PPP2R1B |
| scaffold_2 | 36381253 | -16.56 | KCND3 |
| scaffold_2 | 36381254 | -6.57 | KCND3 |
| scaffold_2 | 36381297 | -11.35 | KCND3 |
| scaffold_2 | 78013496 | 6.67 | |
| scaffold_2 | 78018616 | -6.44 | |
| scaffold_20 | 3718371 | -5.94 | |
| scaffold_20 | 3719478 | 5.14 | |
| scaffold_25 | 16284723 | -12.94 | |
| scaffold_25 | 23048726 | 15.38 | |
| scaffold_27 | 909219 | -7.11 | |
| scaffold_28 | 22472036 | 5.91 | BAIAP2L2 |
| scaffold_309 | 242033 | -21.79 | |
| scaffold_309 | 242298 | 8.82 | |
| scaffold_37 | 20578870 | -5.37 | |
| scaffold_38 | 2547860 | 5.17 | FARS2 |
| scaffold_38 | 4432832 | 11.10 | TXNDC5 |
| scaffold_42 | 5821667 | -16.07 | |
| scaffold_42 | 5826412 | -10.71 | ARL2 |
| scaffold_42 | 6052504 | 9.52 | CDC42EP2 |
| scaffold_42 | 6052506 | 8.97 | CDC42EP2 |
| scaffold_42 | 6052508 | 14.32 | CDC42EP2 |
| scaffold_42 | 6052638 | -9.28 | CDC42EP2 |
| scaffold_42 | 6052734 | -15.30 | CDC42EP2 |
| scaffold_42 | 6096998 | 9.50 | SLC25A45 |
| scaffold_46 | 616270 | 14.59 | |

| | | | |
|---|---|---|---|
| scaffold_46 | 616273 | 22.92 | |
| scaffold_53 | 6342952 | 8.17 | CA10 |
| scaffold_53 | 6359759 | -15.33 | CA10 |
| scaffold_53 | 6445603 | 11.11 | CA10 |
| scaffold_53 | 6449789 | -15.00 | CA10 |
| scaffold_53 | 6449790 | 5.18 | CA10 |
| scaffold_53 | 6461832 | -17.49 | CA10 |
| scaffold_53 | 6525402 | 19.05 | |
| scaffold_53 | 6525403 | -7.14 | |
| scaffold_53 | 6525786 | 10.66 | |
| scaffold_53 | 6525792 | -5.37 | |
| scaffold_53 | 6681603 | -15.68 | |
| scaffold_53 | 6681604 | -20.58 | |
| scaffold_53 | 6710048 | 15.95 | |
| scaffold_53 | 6754504 | 9.34 | UTP18 |
| scaffold_53 | 6754574 | -9.40 | UTP18 |
| scaffold_53 | 6811349 | -31.16 | MBTD1 |
| scaffold_53 | 6811377 | -5.75 | MBTD1 |
| scaffold_53 | 6816726 | -9.20 | NME2 |
| scaffold_53 | 6835594 | -12.18 | |
| scaffold_53 | 6846525 | -10.53 | SPAG9 |
| scaffold_53 | 6859952 | 5.12 | SPAG9 |
| scaffold_53 | 6862130 | 16.67 | SPAG9 |
| scaffold_53 | 6863186 | 11.06 | SPAG9 |
| scaffold_53 | 6904671 | 10.53 | SPAG9 |
| scaffold_53 | 6927508 | 7.44 | SPAG9 |
| scaffold_53 | 6933398 | 22.86 | SPAG9 |
| scaffold_53 | 6935295 | 5.97 | SPAG9 |
| scaffold_53 | 6935296 | 11.11 | SPAG9 |
| scaffold_53 | 6938170 | -6.81 | SPAG9 |
| scaffold_53 | 6945205 | 9.42 | SPAG9 |
| scaffold_53 | 6945245 | 6.22 | SPAG9 |
| scaffold_53 | 6945429 | 10.53 | SPAG9 |
| scaffold_53 | 6947081 | 14.79 | SPAG9 |
| scaffold_53 | 6950228 | 14.97 | SPAG9 |
| scaffold_53 | 6952661 | 15.64 | SPAG9 |
| scaffold_53 | 6952957 | -20.96 | SPAG9 |
| scaffold_53 | 6953168 | -12.07 | |
| scaffold_53 | 6954414 | -5.21 | |
| scaffold_53 | 6956787 | 11.43 | |
| scaffold_53 | 6958361 | -7.25 | |
| scaffold_53 | 6961893 | -33.77 | |
| scaffold_53 | 6962101 | 33.52 | |
| scaffold_53 | 6965557 | -5.56 | |
| scaffold_53 | 6965786 | 16.67 | |
| scaffold_53 | 6965792 | 6.19 | |
| scaffold_53 | 6966868 | -11.43 | |
| scaffold_53 | 6967550 | 11.51 | |
| scaffold_53 | 6968089 | 16.77 | |
| scaffold_53 | 6968224 | -8.33 | |
| scaffold_53 | 6972893 | 5.20 | |
| scaffold_53 | 6972930 | -6.52 | |
| scaffold_53 | 6973738 | 19.46 | |
| scaffold_53 | 6973781 | 5.17 | |
| scaffold_53 | 6977162 | -6.31 | |
| scaffold_56 | 7511308 | 5.11 | |
| scaffold_6 | 32457316 | -18.66 | |
| scaffold_6 | 32457329 | -5.74 | |

| | | | |
|---|---|---|---|
| scaffold_62 | 5837492 | 8.51 | WAPL |
| scaffold_67 | 5551547 | -9.29 | |
| scaffold_67 | 9124061 | -5.41 | ADAM11 |
| scaffold_67 | 9124070 | -7.84 | ADAM11 |
| scaffold_687 | 42540 | 14.88 | |
| scaffold_687 | 42553 | 10.14 | |
| scaffold_7 | 58076842 | -13.43 | |
| scaffold_70 | 7240731 | -5.56 | SYT3 |
| scaffold_72 | 3969892 | -27.65 | |
| scaffold_72 | 3970124 | -5.03 | |
| scaffold_72 | 3970126 | -8.16 | |
| scaffold_72 | 3970180 | -23.64 | |
| scaffold_72 | 3977596 | 8.04 | |
| scaffold_72 | 4939369 | -16.39 | |
| scaffold_72 | 4943921 | 9.09 | |
