## Supplemental Table 12 for "Conserved DNA Methylation Signatures in The Prefrontal Cortex of Newborn and Juvenile Guinea Pigs Following Antenatal Corticosteroid Exposure"

**Supplementary Table 12. List of DMCs identified within the binding site for TFAP2C in the PND14PFC following ACS exposure**

| seqnames | start | MethDiff | geneName |
|---|---|---|---|
| scaffold_0 | 3665268 | -25.68 | |
| scaffold_0 | 5589427 | -24.23 | |
| scaffold_0 | 5589428 | -28.50 | |
| scaffold_1 | 10198664 | 47.78 | |
| scaffold_10 | 48577576 | -17.86 | APBA2 |
| scaffold_11 | 5062946 | -7.20 | |
| scaffold_11 | 14181692 | -16.67 | |
| scaffold_115 | 2716887 | -5.86 | CUX2 |
| scaffold_13 | 9282875 | -15.00 | |
| scaffold_140 | 746180 | 26.76 | TIMP3 |
| scaffold_140 | 746181 | 5.27 | TIMP3 |
| scaffold_15 | 23763883 | -6.53 | |
| scaffold_15 | 32550409 | 11.85 | |
| scaffold_155 | 1146733 | 5.87 | |
| scaffold_155 | 1147857 | -5.99 | |
| scaffold_155 | 1147870 | -6.86 | |
| scaffold_160 | 267890 | -7.14 | |
| scaffold_178 | 1472285 | 7.65 | |
| scaffold_178 | 1472287 | 8.12 | |
| scaffold_18 | 2777029 | -9.61 | |
| scaffold_19 | 36033423 | -7.32 | PPP2R1B |
| scaffold_27 | 909610 | -8.33 | |
| scaffold_28 | 22472103 | -9.09 | BAIAP2L2 |
| scaffold_28 | 22472104 | -7.03 | BAIAP2L2 |
| scaffold_304 | 317594 | -9.70 | |
| scaffold_309 | 242033 | -21.79 | |
| scaffold_32 | 21997047 | -7.32 | |
| scaffold_38 | 4432832 | 11.10 | TXNDC5 |
| scaffold_39 | 11246309 | -25.38 | PARD3 |
| scaffold_42 | 5821667 | -16.07 | |
| scaffold_42 | 6052734 | -15.30 | CDC42EP2 |
| scaffold_46 | 616270 | 14.59 | |
| scaffold_46 | 616273 | 22.92 | |
| scaffold_46 | 616276 | -20.00 | |
| scaffold_53 | 6034626 | -8.47 | CA10 |
| scaffold_53 | 6272353 | -8.17 | CA10 |
| scaffold_53 | 6354521 | 7.35 | CA10 |
| scaffold_53 | 6475532 | -8.03 | |
| scaffold_53 | 6525402 | 19.05 | |
| scaffold_53 | 6525403 | -7.14 | |
| scaffold_53 | 6680536 | 19.03 | |
| scaffold_53 | 6754504 | 9.34 | UTP18 |
| scaffold_53 | 6754574 | -9.40 | UTP18 |
| scaffold_53 | 6816429 | -5.26 | NME2 |
| scaffold_53 | 6818091 | 13.92 | NME1 |
| scaffold_53 | 6842804 | -7.27 | |
| scaffold_53 | 6862130 | 16.67 | SPAG9 |
| scaffold_53 | 6899246 | 9.17 | SPAG9 |
| scaffold_53 | 6927508 | 7.44 | SPAG9 |
| scaffold_53 | 6933398 | 22.86 | SPAG9 |
| scaffold_53 | 6938170 | -6.81 | SPAG9 |
| scaffold_53 | 6941677 | -7.69 | SPAG9 |
| scaffold_53 | 6945429 | 10.53 | SPAG9 |
| scaffold_53 | 6947081 | 14.79 | SPAG9 |
| scaffold_53 | 6950228 | 14.97 | SPAG9 |
| scaffold_53 | 6953353 | -19.48 | |

| | | | |
|---|---|---|---|
| scaffold_53 | 6958834 | -5.40 | |
| scaffold_53 | 6962101 | 33.52 | |
| scaffold_53 | 6965942 | -6.54 | |
| scaffold_53 | 6966868 | -11.43 | |
| scaffold_53 | 6968089 | 16.77 | |
| scaffold_53 | 6973419 | -11.17 | |
| scaffold_53 | 6973738 | 19.46 | |
| scaffold_53 | 6974062 | -12.30 | |
| scaffold_53 | 6975938 | -13.04 | |
| scaffold_53 | 6977162 | -6.31 | |
| scaffold_53 | 6977755 | -10.74 | |
| scaffold_53 | 6977756 | 8.61 | |
| scaffold_62 | 5837492 | 8.51 | WAPL |
| scaffold_67 | 9124070 | -7.84 | ADAM11 |
| scaffold_687 | 42370 | 7.02 | |
| scaffold_7 | 58076842 | -13.43 | |
| scaffold_72 | 3977596 | 8.04 | |
| scaffold_72 | 3977919 | 5.56 | |
| scaffold_72 | 4943921 | 9.09 | |
