## Supplemental Table 13 for "Conserved DNA Methylation Signatures in The Prefrontal Cortex of Newborn and Juvenile Guinea Pigs Following Antenatal Corticosteroid Exposure"

**Supplementary Table 13. List of DMCs identified within the binding site for SP1 in the PND14PFC following ACS exposure**

| seqnames | start | MethDiff | geneName |
|---|---|---|---|
| scaffold_0 | 5589427 | -24.23 | |
| scaffold_0 | 5589428 | -28.50 | |
| scaffold_0 | 9497625 | -21.43 | |
| scaffold_0 | 87065049 | 32.31 | |
| scaffold_1 | 10198664 | 47.78 | |
| scaffold_11 | 5062648 | 11.01 | |
| scaffold_114 | 2615152 | -7.14 | |
| scaffold_125 | 3159554 | -16.95 | ATP6V1C2 |
| scaffold_132 | 672314 | -8.33 | RBM10 |
| scaffold_15 | 32550353 | -13.42 | |
| scaffold_15 | 32550354 | 9.26 | |
| scaffold_15 | 32550359 | -11.26 | |
| scaffold_155 | 1146922 | 5.36 | |
| scaffold_166 | 578641 | -23.56 | SORCS2 |
| scaffold_166 | 1409905 | -5.56 | DOK7 |
| scaffold_178 | 1472285 | 7.65 | |
| scaffold_178 | 1472335 | 8.06 | |
| scaffold_178 | 1472622 | -10.00 | |
| scaffold_19 | 30378554 | -8.80 | DSCAML1 |
| scaffold_2 | 36381253 | -16.56 | KCND3 |
| scaffold_2 | 36381254 | -6.57 | KCND3 |
| scaffold_2 | 65848849 | 6.50 | GNG12 |
| scaffold_2 | 78017478 | 7.91 | |
| scaffold_20 | 3718371 | -5.94 | |
| scaffold_24 | 2100133 | 9.18 | |
| scaffold_25 | 16284723 | -12.94 | |
| scaffold_27 | 2634722 | -17.25 | HMCN2 |
| scaffold_32 | 21990994 | -17.55 | |
| scaffold_37 | 20598529 | 7.63 | |
| scaffold_38 | 2547860 | 5.17 | FARS2 |
| scaffold_42 | 6052734 | -15.30 | CDC42EP2 |
| scaffold_509 | 65302 | -29.46 | |
| scaffold_53 | 2049886 | -6.37 | |
| scaffold_53 | 2049889 | -5.29 | |
| scaffold_53 | 6270684 | 7.49 | CA10 |
| scaffold_53 | 6354521 | 7.35 | CA10 |
| scaffold_53 | 6445603 | 11.11 | CA10 |
| scaffold_53 | 6525839 | -7.20 | |
| scaffold_53 | 6680218 | -32.29 | |
| scaffold_53 | 6687281 | -8.17 | |
| scaffold_53 | 6754574 | -9.40 | UTP18 |
| scaffold_53 | 6842804 | -7.27 | |
| scaffold_53 | 6862130 | 16.67 | SPAG9 |
| scaffold_53 | 6882412 | 9.15 | SPAG9 |
| scaffold_53 | 6899278 | -10.87 | SPAG9 |
| scaffold_53 | 6899590 | 6.37 | SPAG9 |
| scaffold_53 | 6927508 | 7.44 | SPAG9 |
| scaffold_53 | 6927516 | 25.38 | SPAG9 |
| scaffold_53 | 6935017 | -6.14 | SPAG9 |
| scaffold_53 | 6935730 | -6.45 | SPAG9 |
| scaffold_53 | 6938170 | -6.81 | SPAG9 |
| scaffold_53 | 6945429 | 10.53 | SPAG9 |
| scaffold_53 | 6947075 | 29.08 | SPAG9 |
| scaffold_53 | 6947078 | 18.26 | SPAG9 |
| scaffold_53 | 6947081 | 14.79 | SPAG9 |
| scaffold_53 | 6950228 | 14.97 | SPAG9 |

| | | | |
|---|---|---|---|
| scaffold_53 | 6954414 | -5.21 | |
| scaffold_53 | 6958457 | 6.56 | |
| scaffold_53 | 6962393 | 20.00 | |
| scaffold_53 | 6965682 | 14.90 | |
| scaffold_53 | 6965731 | 32.38 | |
| scaffold_53 | 6966928 | -9.23 | |
| scaffold_53 | 6968224 | -8.33 | |
| scaffold_53 | 6968268 | -8.67 | |
| scaffold_53 | 6972893 | 5.20 | |
| scaffold_53 | 6973240 | -11.35 | |
| scaffold_53 | 6973390 | -5.65 | |
| scaffold_53 | 6973846 | -7.40 | |
| scaffold_53 | 6977494 | 5.51 | |
| scaffold_53 | 6977495 | -7.77 | |
| scaffold_67 | 9124070 | -7.84 | ADAM11 |
| scaffold_687 | 42460 | 5.50 | |
| scaffold_687 | 42626 | 11.58 | |
| scaffold_7 | 58076412 | 15.79 | |
| scaffold_7 | 58076842 | -13.43 | |
| scaffold_72 | 3969966 | 14.34 | |
| scaffold_72 | 3970168 | 10.91 | |
| scaffold_72 | 4939369 | -16.39 | |
| scaffold_72 | 4943361 | -10.63 | |
| scaffold_85 | 564393 | -7.37 | EVC2 |
| scaffold_9 | 6720456 | -21.43 | TMCC3 |
