## Supplemental Table 14 for "Conserved DNA Methylation Signatures in The Prefrontal Cortex of Newborn and Juvenile Guinea Pigs Following Antenatal Corticosteroid Exposure"

**Supplementary Table 14. List of DMCs identified within the binding site for ZNF263 in the PND14PFC following ACS exposure**

| seqnames | start | MethDiff | geneName |
|---|---|---|---|
| scaffold_0 | 3684666 | 12.69 | RIMS2 |
| scaffold_0 | 9497625 | -21.43 | |
| scaffold_1 | 65104682 | -45.45 | |
| scaffold_1 | 65104691 | -51.14 | |
| scaffold_104 | 2635279 | -7.50 | |
| scaffold_109 | 2866416 | -5.98 | PLXND1 |
| scaffold_109 | 2866417 | -10.59 | PLXND1 |
| scaffold_11 | 5062648 | 11.01 | |
| scaffold_11 | 14181700 | 16.67 | |
| scaffold_125 | 3159545 | 19.12 | ATP6V1C2 |
| scaffold_135 | 1226867 | -11.32 | |
| scaffold_140 | 746246 | -6.17 | TIMP3 |
| scaffold_140 | 746256 | 9.43 | TIMP3 |
| scaffold_15 | 32550354 | 9.26 | |
| scaffold_15 | 32550359 | -11.26 | |
| scaffold_15 | 32550380 | 15.71 | |
| scaffold_15 | 32550409 | 11.85 | |
| scaffold_155 | 1147655 | -6.98 | |
| scaffold_155 | 1147857 | -5.99 | |
| scaffold_178 | 1472271 | 6.36 | |
| scaffold_2 | 36381253 | -16.56 | KCND3 |
| scaffold_2 | 36381254 | -6.57 | KCND3 |
| scaffold_2 | 46708451 | -18.31 | |
| scaffold_2 | 67751748 | -8.05 | CACHD1 |
| scaffold_2 | 67751749 | -5.17 | CACHD1 |
| scaffold_2 | 78015134 | 10.38 | |
| scaffold_20 | 3718371 | -5.94 | |
| scaffold_24 | 8136779 | 13.53 | |
| scaffold_25 | 16284723 | -12.94 | |
| scaffold_27 | 909575 | 9.09 | |
| scaffold_27 | 909610 | -8.33 | |
| scaffold_27 | 11748547 | -29.55 | |
| scaffold_28 | 27122047 | 5.65 | |
| scaffold_38 | 2547891 | 5.06 | FARS2 |
| scaffold_46 | 616270 | 14.59 | |
| scaffold_46 | 616273 | 22.92 | |
| scaffold_46 | 616430 | -7.11 | |
| scaffold_53 | 6184604 | -6.52 | CA10 |
| scaffold_53 | 6300773 | 7.23 | CA10 |
| scaffold_53 | 6342952 | 8.17 | CA10 |
| scaffold_53 | 6354521 | 7.35 | CA10 |
| scaffold_53 | 6525786 | 10.66 | |
| scaffold_53 | 6681742 | -19.81 | |
| scaffold_53 | 6754504 | 9.34 | UTP18 |
| scaffold_53 | 6818091 | 13.92 | NME1 |
| scaffold_53 | 6842817 | -16.79 | |
| scaffold_53 | 6859945 | -7.80 | SPAG9 |
| scaffold_53 | 6860003 | 35.00 | SPAG9 |
| scaffold_53 | 6874498 | 5.27 | SPAG9 |
| scaffold_53 | 6874517 | 6.65 | SPAG9 |
| scaffold_53 | 6887921 | -6.01 | SPAG9 |
| scaffold_53 | 6899246 | 9.17 | SPAG9 |
| scaffold_53 | 6917998 | 18.10 | SPAG9 |
| scaffold_53 | 6921964 | 6.70 | SPAG9 |
| scaffold_53 | 6922046 | 35.78 | SPAG9 |
| scaffold_53 | 6935017 | -6.14 | SPAG9 |

| | | | |
|---|---|---|---|
| scaffold_53 | 6935487 | -9.30 | SPAG9 |
| scaffold_53 | 6945245 | 6.22 | SPAG9 |
| scaffold_53 | 6956395 | -10.00 | |
| scaffold_53 | 6958384 | -11.81 | |
| scaffold_53 | 6958457 | 6.56 | |
| scaffold_53 | 6961979 | -11.00 | |
| scaffold_53 | 6965682 | 14.90 | |
| scaffold_53 | 6965731 | 32.38 | |
| scaffold_53 | 6966868 | -11.43 | |
| scaffold_53 | 6966928 | -9.23 | |
| scaffold_53 | 6967550 | 11.51 | |
| scaffold_53 | 6968224 | -8.33 | |
| scaffold_53 | 6968268 | -8.67 | |
| scaffold_53 | 6972893 | 5.20 | |
| scaffold_53 | 6972938 | -8.60 | |
| scaffold_53 | 6973136 | 8.70 | |
| scaffold_53 | 6973781 | 5.17 | |
| scaffold_53 | 6973819 | -10.00 | |
| scaffold_53 | 6973846 | -7.40 | |
| scaffold_67 | 9124070 | -7.84 | ADAM11 |
| scaffold_67 | 9124079 | -6.14 | ADAM11 |
| scaffold_687 | 42540 | 14.88 | |
| scaffold_687 | 42626 | 11.58 | |
| scaffold_7 | 58076412 | 15.79 | |
| scaffold_7 | 58076842 | -13.43 | |
| scaffold_72 | 3970180 | -23.64 | |
| scaffold_72 | 3977596 | 8.04 | |
| scaffold_72 | 3977932 | -8.33 | |
| scaffold_72 | 4943448 | 5.09 | |
| scaffold_85 | 564384 | -8.55 | EVC2 |
| scaffold_85 | 564393 | -7.37 | EVC2 |
